# *In Vivo* Dual RNA-Seq Analysis Reveals the Basis for Differential Tissue Tropism of Clinical Isolates of *Streptococcus pneumoniae*

**DOI:** 10.1101/862755

**Authors:** Vikrant Minhas, Rieza Aprianto, Lauren J. McAllister, Hui Wang, Shannon C. David, Kimberley T. McLean, Iain Comerford, Shaun R. McColl, James C. Paton, Jan-Willem Veening, Claudia Trappetti

**Affiliations:** Research Centre for Infectious Diseases, Department of Molecular and Biomedical Science, University of Adelaide, Adelaide, 5005, Australia; Department of Fundamental Microbiology, Faculty of Biology and Medicine, University of Lausanne, CH-1015 Lausanne, Switzerland; Department of Molecular and Biomedical Science, University of Adelaide, Adelaide, 5005, Australia

## Abstract

*Streptococcus pneumoniae* is a genetically diverse human-adapted pathogen commonly carried asymptomatically in the nasopharynx. We have recently shown that a single nucleotide polymorphism (SNP) in the raffinose pathway regulatory gene *rafR* accounts for a significant difference in the capacity of clonally-related strains to cause localised versus systemic infection. Here we have used dual RNA-seq to show that this SNP extensively impacts both bacterial and host transcriptomes in infected lungs. It affects expression of bacterial genes encoding multiple sugar transporters, and fine-tunes carbohydrate metabolism, along with extensive rewiring of host transcriptional responses to infection, particularly expression of genes encoding cytokine and chemokine ligands and receptors. The dual RNA-seq data predicted a crucial role for differential neutrophil recruitment in the distinct virulence profiles of the infecting strains and single cell analysis revealed that while reduced expression of the RafR regulon driven by a single *rafR* SNP provides a clear advantage for pneumococci to colonize the ear, in the lung it leads to massive recruitment of neutrophils and bacterial clearance. Importantly, the observed disease outcomes were confirmed by *in vivo* neutrophil depletion showing that early detection of bacteria by the host in the lung environment is crucial for effective clearance. Thus, dual RNA-seq provides a powerful tool for understanding complex host-pathogen interactions and revealed how a single bacterial SNP can drive differential disease outcomes.

## INTRODUCTION

*Streptococcus pneumoniae* is a major human pathogen responsible for massive global morbidity and mortality. Despite this, the pneumococcus makes up part of the commensal human nasopharyngeal flora, colonizing up to 65% of individuals (Kadioglu et al., 2008) (Weiser et al., 2018). *S. pneumoniae* can invade from this nasopharyngeal reservoir to cause disease, for example, by aspiration into the lungs to cause pneumonia, by direct or indirect invasion of the blood (bacteremia) or central nervous system (meningitis), or by ascension of the eustachian tube to access the middle ear and cause the localised disease otitis media (OM) (Kadioglu et al., 2008)(Weiser et al., 2018). *S. pneumoniae* is an extremely heterogeneous species, comprising at least 98 capsular serotypes and over 12,000 clonal lineages (sequence types; ST) recognisable by multi-locus sequence typing (Enright and Spratt, 1998; van Tonder et al., 2019). Unsurprisingly, *S. pneumoniae* strains differ markedly in their capacity to progress from carriage to disease and/or the nature of the disease that they cause (Kadioglu et al., 2008; Weiser et al., 2018).

We have previously reported marked differences in virulence in a murine intranasal (IN) challenge model between *S. pneumoniae* strains belonging to the same serotype and ST, which correlated with clinical isolation site in humans (ear versus blood). In serotype 3 ST180, ST232 and ST233, and in serotype 14 ST15, human ear isolates had greater capacity to cause OM in mice relative to their respective serotype-/ST-matched blood isolates, while blood isolates preferentially caused pneumonia or sepsis in mice, suggesting stable niche adaptation within a clonal lineage (Amin et al., 2015; Trappetti et al., 2013). Recently, we have shown that the distinct virulence phenotypes correlated with single nucleotide polymorphisms (SNPs) in genes encoding uptake and utilization of the sugar raffinose. In serotype 14 ST15, the SNP was in the raffinose pathway regulatory gene *rafR*, while in serotype 3 ST180, the SNP was in *rafK*, which encodes an ATPase that energises the raffinose uptake ABC transporter (Minhas et al., 2019). Both SNPs result in non-conservative amino acid changes in functionally critical domains of the respective gene product (D249G for RafR; I227T for RafK). Moreover, in both serotypes/lineages, ear isolates had *in vitro* growth defects in a chemically-defined medium with raffinose as the sole carbon source, correlating with defective transcription of raffinose pathway operons. Remarkably, in serotype 14 ST15, exchanging the *rafR* alleles between blood and ear isolates reversed both the *in vitro* and *in vivo* phenotypes (Minhas et al., 2019). Thus, the single D249G SNP in *rafR* appears to be the determinant of differential virulence phenotype between the blood and ear isolates, which may reflect differential engagement of innate host defences and/or differential bacterial nutritional fitness in distinct host niches (**Figure 1**).

**Figure 1.**
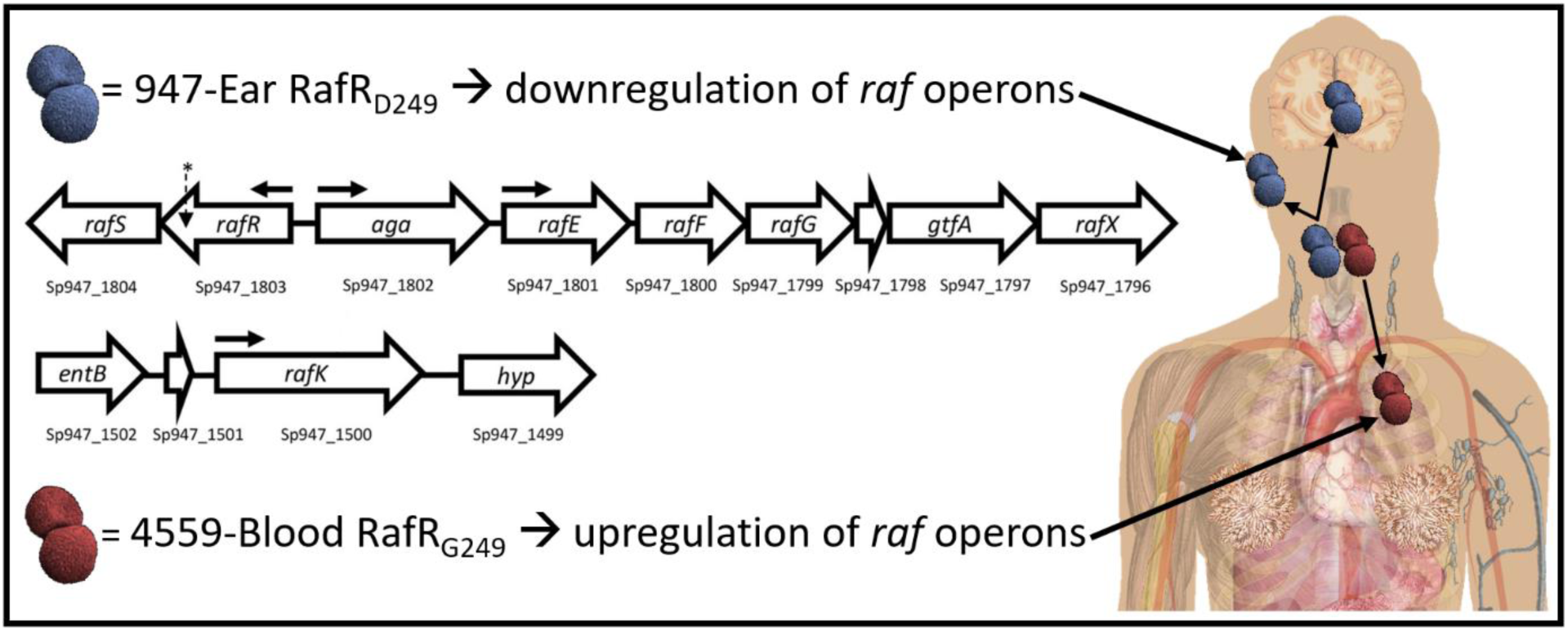
A SNP in *rafR* between the serotype 14 sequence type 15 clonal isolates 4559-Blood and 947-Ear leads to a non-conservative G249D amino acid substitution in the raffinose pathway regulator RafR. RafR_G249_ results in upregulation of *raf* operons (horizontal arrows denote transcriptional start sites) in 4559-Blood relative to 947-Ear, favouring persistence in the lung after intranasal challenge. Lower *raf* pathway expression mediated by RafR_D249_ facilitates clearance of 947-Ear from the lung, but promotes spread to and/or persistence in the ear and brain. The location of the SNP in *rafR* is indicated by an asterisk (Minhas et al., 2019).

Dual RNA-seq applies deep sequencing to simultaneously quantify genome-wide transcriptional responses of host and pathogen (Westermann et al., 2017; Wolf et al., 2018). This approach offers higher efficiency and more restricted technical bias compared to conventional approaches, such as assaying single species or array-based methods. In the present study, we have used dual RNA-seq analysis to examine host-pathogen transcriptional cross-talk in the blood and ear isolates and *rafR*-swapped derivatives thereof, during the early stages of infection. Our data strongly suggest that the *rafR* SNP interacts with the pneumococcal genetic background in the different clinical isolates, which in turn, induces variegated transcriptional responses in the pathogen; this response, in turn, initiates a diverging host response that determines the outcome of infection.

## RESULTS AND DISCUSSION

### Comparative Host/Pathogen Transcriptomics

Our previous studies have shown that at 6 h after IN challenge with serotype 14 ST15 *S. pneumoniae*, the numbers of blood and ear isolates (strains 4559-Blood and 9-47-Ear, respectively) present in murine lungs are similar (10^6^ – 10^7^ CFU per lung). However, by 24 h, the ear isolate had been cleared from the lungs, instead spreading to the ear and brain. In contrast, the blood isolate persisted in the lungs at 24 h, but did not spread to the ear or brain (Amin et al., 2015). Thus, 6 h post infection is a critical decision point in the pathogenic process, and the similarity of bacterial loads in the lung at this time enables examination of the transcriptional cross-talk between the pneumococcus and its host without the complication of bacterial dose effects. Accordingly, groups of 12 mice were anaesthetized and challenged IN with 10^8^ CFU of either 4559-Blood, 9-47-Ear or their respective *rafR*-swapped mutants (**Table 1**); at 6 h, mice were euthanized and total RNA was extracted from perfused lungs and purified. RNA extracted from lungs of 4 mice were pooled into 1 sample for subsequent Dual RNA-seq analysis in triplicate (see Methods).

**Table 1.** Resources Table, describing bacterial strains, antibodies, chemicals and commercial assays used in this study.

| **Bacterial Strains** | **Source** | **Identifier** |
| --- | --- | --- |
| *Streptococcus pneumoniae*: Clinical blood isolate capsular serotype 14, Multi Locus Sequence Type 15: 4559-Blood | Minhas *et al.* | N/A |
| *Streptococcus pneumoniae*: Clinical isolate ear capsular serotype 14, Multi Locus Sequence Type 15: 9-47-Ear | Minhas *et al.* | N/A |
| *Streptococcus pneumoniae*: Clinical isolate ear capsular serotype 14, Multi Locus Sequence Type 15: 4559M (4559-Blood containing *rafR* of 9-47-Ear) | Minhas *et al.* | N/A |
| *Streptococcus pneumoniae*: Clinical isolate ear capsular serotype 14, Multi Locus Sequence Type 15: 9-47M (947 containing *rafR* of 4559-Blood) | Minhas *et al.* | N/A |
| **Antibodies** | **Source** | **Identifier** |
| Anti-mouse/human CD11b-PE (clone M1/70) | BioLegend | Cat# 101208, RRID: AB_312791 |
| Anti-mouse CD11c-BV786 (clone HL3) | BD Biosciences | Cat# 563735, RRID: AB_2738394 |
| Anti-mouse CD24-BV711 (clone M1/69) | BD Biosciences | Cat# 563450, RRID: AB_2738213 |
| Anti-mouse CD45-FITC (clone 30-F11) | BioLegend | Cat# 103107, RRID: AB_312972 |
| Anti-mouse CD64-BV421 (clone X54-5/7.1) | BioLegend | Cat# 139309, RRID: AB_2562694 |
| Anti-mouse Ly6C-PerCP/Cy5.5 (clone HK1.4) | BioLegend | Cat# 128011, RRID: AB_1659242 |
| Anti-mouse Ly6G-BUV395 (clone 1A8) | BD Biosciences | Cat# 563978, RRID: AB_2716852 |
| Anti-mouse I-A/I-E-BV650 (clone M5/114.15.2) | BD Biosciences | Cat# 563415, RRID: AB_2738192 |
| Anti-mouse Ly6G (clone 1A8) | Bio X Cell | Cat# BE0075-1, RRID: AB_1107721 |
| Rat IgG2A Isotype Control (clone 54447) | R and D Systems | Cat# MAB006, RRID: AB_357349 |
| Anti-mouse Ly-6G, Ly-6C-Biotin (clone RB6-8C5) | BD Biosciences | Cat# 553125, RRID: AB_394641 |
| Anti-mouse IL-17A (clone 17F3) | Bio X Cell | Cat# BE0173, RRID: AB_10950102 |
| Mouse IgG1 Isotype Control (clone MOPC-21) | Bio X Cell | Cat# BE0083, RRID: AB_1107784 |
| **Chemicals** | **Source** | **Identifier** |
| Zombie NIR™ Fixable Viability Kit | BioLegend | Cat# 423105 |
| BV421 Streptavidin | BD Biosciences | Cat# 563259 |
| Horse serum, heat inactivated | Thermo Fisher Scientific | Cat# 26050088 |
| LAB LEMCO Nutrient Broth + 10% horse serum (serum broth) | Adelaide University Technical Services Unit | Custom Synthesis |
| Columbia blood agar base (dehydrated) | Thermo Scientific Oxoid | Cat# CM0331T |
| Defibrinated horse blood | Australian Ethical Biologicals | Cat# PDHB500 |
| *Pentobarbital Sodium Anaesthetic Injection* | *Illium* | N/A |
| Acid-Phenol:Chloroform, pH 4.5 (with IAA, 125:24:1) | Thermo Fisher Scientific | Cat# AM9722 |
| SuperScript® III Platinum® One-Step qRT-PCR Kit | Thermo Fisher Scientific | Cat# 11736059 |
| TRIzol® Reagent | Thermo Fisher Scientific | Cat# 15596026 |
| DMEM + HEPES | Thermo Fisher Scientific | Cat # 12430054 |
| Fetal Bovine Serum | Corning | Cat# 35076CV |
| Collagenase from *Clostridium histolyticum* | Sigma-Aldrich | Cat# C9891 |
| Dnase I | Sigma-Aldrich | Cat# 11284932001 |
| Sodium azide solution | Sigma-Aldrich | Cat# 08591 |
| Paraformaldehyde | Sigma-Aldrich | Cat# P6148 |
| Bovine Serum Albumin Fraction V | Sigma-Aldrich | Cat# 10735 |
| 10x RBC lysis buffer | eBioscience | Cat# 00430054 |
| **Commercial Assays** | **Source** | **Identifier** |
| RNeasy Mini Kit | Qiagen | Cat# 74106 |
| Ribo-Zero Murine rRNA depletion kit | Illumina | N/A |
| Ribo-Zero Gram positive bacteria rRNA depletion kit | Illumina | N/A |
| Stranded cDNA library preparation kit | Illumina | N/A |

Within the sequencing libraries, an overwhelming majority of reads originate from the host genome (average: 99.5%, range: 99.1 to 99.7%), which translates into an average depth of 1.3 times (range: 0.8 to 1.8 times). Conversely, 0.52% of the total reads originated from the pathogen genome (0.33 to 0.93%). Of these pneumococcal reads, 64.5% mapped onto the genes encoding ribosomal RNAs (61.4 to 67.3%) and 35.5% reads mapped onto non-rRNA encoding genes (32.7 to 38.6%). Previous data indicate that non-depleted libraries only contain 5% non-ribosomal RNA reads; thus, this treatment enriched the non-ribosomal RNAs sevenfold. Non-ribosomal reads depth was 2.7 times for the pathogen genome, ranging from 1.4 to 4.6 times. Further downstream analysis, including differential gene expression, excluded ribosomal reads from the pathogen library. Tables **S1-S6** list pneumococcal genes that are significantly differentially expressed (fold change (FC) >2, *p* <0.05) for each of the six pairwise comparisons between the four strains. Tables **S7-S12** list murine genes that are significantly differentially expressed (FC >1.5, *p* <0.05) for the same pairwise comparisons.

### Fine Tuning of Carbohydrate Metabolism Driven by RafR During Pneumococcal Infection

In order to directly compare pathogen transcriptional responses in murine lung, we listed homologous genes between the two wild type ear and blood isolates and used these genes to visualize the transcriptional response in a principal component analysis (PCA) plot (**Figure 2a**). Here, the pneumococcal transcriptional response of the ear isolate (strain 9-47-Ear, dark orange) to murine lung infection diverges considerably from the response of the blood isolate (strain 4559-Blood, dark purple). Specifically, 76 homologous genes are significantly upregulated in the ear isolate, while 40 genes are upregulated in the blood isolate in the murine lung. Upregulated genes in strain 9-47-Ear include genes involved in carbohydrate metabolism, general stress response and nutrient transporters, while upregulated genes in the blood isolate include genes encoding permeases for small molecules and nisin biosynthesis orthologous proteins.

**Figure 2.**
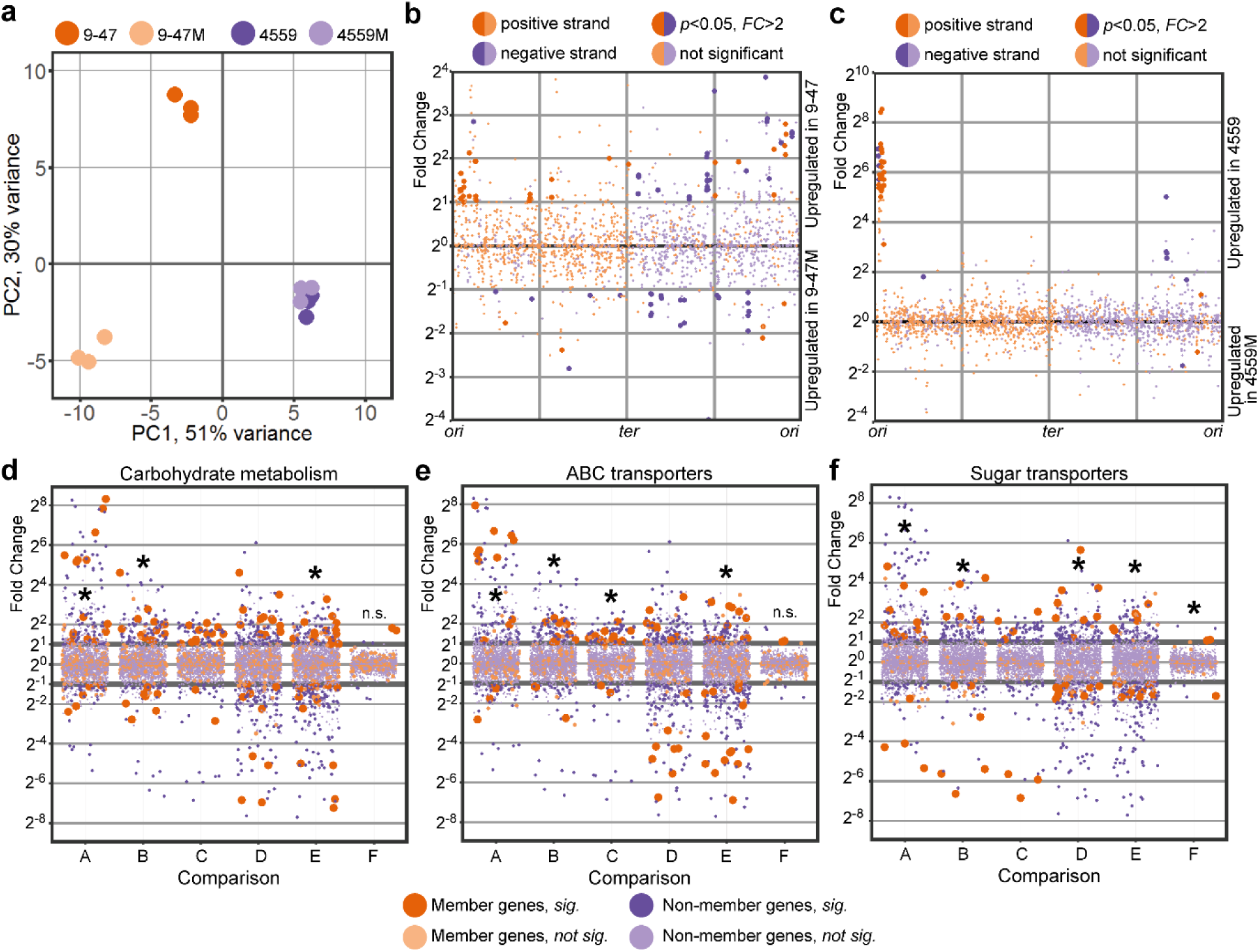
Pathogen transcriptional responses in murine lung. **(a)**. PCA plot showing divergence of transcriptional response to lung infection within the ear (9-47-Ear) and blood isolates (4559-Blood). *rafR* swap (9-47M) rewires pneumococcal transcriptional response only in the ear isolate background but not in the blood isolate. (**b)**. Differential expression due to the *rafR* swap in the ear isolate background is spread throughout the pneumococcal genome, while in the blood isolate background, differential expression due to the *rafR* swap is limited to a genomic island (**c**). Functional enrichment showed specific function being differentially expressed, including carbohydrate metabolism (**d**), ABC transporters (**e**) and sugar transporters (**F**). A: comparison of 9-47-Ear to 9-47M; B: 9-47-Ear to 4559-Blood; C: 9-47-Ear to 4559M; D: 9-47M to 4559-Blood; E: 9-47M to 4559M and F: 4559-Blood to 4559M. * denotes statistically significant functional enrichment for the indicated strain-strain comparison.

Furthermore, replacing the *rafR* of the ear isolate 9-47-Ear with the allele from the blood isolate 4559-Blood (designated strain 9-47M) dissociates its transcriptional response considerably from its parental 9-47-Ear strain (**Figure 2a**, 9-47-Ear, dark orange to 9-47M, light orange). Specifically, 87 genes are upregulated in the wild type strain (9-47-Ear) while 36 genes are upregulated in the *rafR* swap strain (9-47M, **Figure 2b**), with differentially expressed genes being spread across the pneumococcal genome. Presence of the blood isolate *rafR* allele in 9-47-Ear activates the expression of major genes pertaining to carbohydrate metabolism, including *adhA* (alcohol dehydrogenase) and *spxB* (pyruvate oxidase); and genes encoding permeases, including *glnH6P6* (transporting arginine, cysteine) and *ycjOP-yesO* (transporting multiple sugars). Also, a subset of genes with function in carbohydrate metabolism are repressed in the *rafR*-swap strain, such as glycogen synthesis (*glgACD*) and sucrose metabolism (*scrB*) and ribulose metabolism (*ulaDEF*). Expression of seven genes encoding subunits of ATPase (*ntpABCDEGK*) and genes coding for iron (*piuB*) and sugar (*scrA, satABC, gadEW*) permeases are also repressed.

On the other hand, replacing the blood isolate *rafR* with the ear allele (designated strain 4559M) does not noticeably interrupt pneumococcal transcriptional response to murine lung (**Figure 2a**, 4559-Blood, dark purple to 4559M, light purple). Essentially, the *rafR* swap activates only two genes: *yxlF*, encoding a putative subunit of an ABC transporter and *phoU1* encoding a phosphate transporter; and represses 35 genes, mostly contained in a single genomic island (**Figure 2c**, upregulated in 4559-Blood). The genomic island consists of 28 consecutive genes encoding subunits of bacteriophage(s), interspaced by *dnaC*, encoding a DNA replication protein and *lytA*, encoding autolysin. The activation of bacteriophage-associated genes indicates that the original isolate (strain 4559-Blood) endures host-derived stress, unlike the *rafR* swap mutant (strain 4559M). Other genes repressed in 4559M include *adhAE* (alcohol dehydrogenases), *gtfA* (sucrose phosphorylase) and *rafEG* (raffinose sugar transporter). Taken together, the single D249G SNP in *rafR* interferes with global gene expression within the already transcriptionally-distinct parental clinical isolates. This effect is more pronounced in the ear isolate (9-47-Ear) than the blood isolate (4559-Blood).

Next, we performed quantified enrichment analyses on specific gene functions. Carbohydrate metabolism is enriched in the differentially expressed genes between the pneumococcal strains, particularly when comparing the ear isolate to its cognate *rafR* swap (**Figure 2d**, *comparison A*, 9-47-Ear vs 9-47M, *p* = 0.03), comparing the two clinical isolates (*comparison B*, 9-47-Ear vs 4559-Blood, *p* = 0.017) and comparing the swap cognates (*comparison E*, 9-47M vs. 4559M, *p* = 0.041). Another function, ABC transporters, is also enriched in the comparison within the ear isolates (**Figure 2e**, *comparison A*, 9-47-Ear vs. 9-47M, *p* = 0.049), between the original isolates (*comparison B*, 9-47-Ear vs 4559-Blood, *p* = 0.014), between ear isolate and *rafR* 746G in blood isolate (*comparison C*, 9-47-Ear vs. 4559M, *p* = 0.01) and between the *rafR* cognates (*comparison E*, 9-47M vs. 4559M, *p* = 1.8 ×10^−4^).

Additionally, since the pneumococcal genome has an exceptionally high number of sugar transporters (Bidossi et al., 2012), we quantified enrichment for this function (**Figure 2f**). Sugar transporters are enriched in almost all comparisons (except between 4559-Blood and 4559M), highlighting the role of *rafR* in the widespread regulation of pneumococcal sugar importers. Specifically, ear and blood isolates behave differently in regard to sugar transporter expression (9-47-Ear vs. 4559-Blood, **Figure 2f**, *comparison B*). The ear isolate upregulates *scrA* (encoding a mannose and trehalose transporter) and *ulaA* (ascorbate transporter), while the blood isolate upregulates *ycjOP-yesO* (alternative sugar transporters), *rafE* (raffinose transporter) and *malFG* (maltose transporter). Furthermore, *rafR* swap in the ear isolate background (9-47-Ear vs. 9-47M, **Figure 2f**, *comparison A*) reduces the expression of *gadEW* (encoding sorbose and mannose transporter), *satABC* (arabinose and lactose transporter), *ulaAC* and *glpF* (glycerol transporter), while the swap activates the expression of *ycjOP-yesO* and *bguD* (encoding complex polysaccharide transporters). In contrast, *rafR* swap in the blood isolate background (4559-Blood vs. 4559M, **Figure 2f**, *comparison F*) downregulates the expression of *rafEG* and *malD* (maltose transporter). The enrichment analysis reveals that the D249G SNP in *rafR* directly and indirectly affects the expression of genes encoding sugar transporters, other (ABC) transporters and carbohydrate metabolism.

We also identified genes that were commonly up or down regulated between the strains that persisted in murine lungs (4559-Blood and 9-47M) or the strains that were cleared from the lungs by 24 h post-infection (9-47-Ear and 4559M), as these may be determinants of the distinct virulence phenotypes of the blood and ear isolates. *adhP* (Sp947_00279) was significantly upregulated in the strains that persisted in the lungs, 9-47M and 4559-Blood (9-47-Ear vs 9-47M, FC = 0.48, *p* = 0.004; 9-47-Ear vs 4559-Blood, FC = 0.32, *p* = 5.38 ×10^−7^; 9-47M vs 4559M = 2.39, *p* = 0.00018; 4559-Blood vs 4559M, FC = 3.53, *p* = 2.23×10^−8^). Additionally, an operon containing two permeases, Sp947_01595 and Sp947_01596, and a putative beta-D-galactosidase Sp947_01598, involved in the import of sialic acid and N-Acetylmannosamine, were highly down regulated by 947-Ear, and less so by 9-47M (9-47-Ear vs 9-47M, FC’s = 0.05, 0.06 and 0.06, *p* = 1.12×10^−6^, 1.70×10^−7^ and 7.36×10^−5^; 9-47-Ear vs 4559-Blood. FC’s = 0.02, 0.02 and 0.02, *p* = 9.79×10^−12^, 2.70×10^−13^ and 2.11×10^−8^; 9-47M vs 4559M, FC’s = 0.32, 0.35 and 0.36, *p* = 1.02×10^−7^, 2.53×10^−7^ and 0.00016, respectively). 8 genes from the genomic region Sp947_0842 to Sp947_0855, as well as Sp947_00631 and Sp947_02096, were significantly upregulated in the strains that were cleared from the lungs by 24 h post-infection, 9-47-Ear and 4559M (for 9-47-Ear vs 9-47M, 9-47-Ear vs 4559-Blood and 9-47M vs 4559M comparisons). Among these genes was a sialidase, Sp947_00844 (9-47-Ear vs 9-47M, FC = 313, *p* = 3.08×10^−10^; 9-47-Ear vs 4559-Blood, FC = 2.53, *p* = 1.14×10^−8^; 9-47M vs 4559M, FC = 0.01, *p* = 4.59×10^−8^). Alpha-glycerophosphate oxidase *glpO*, Sp947_02129, was also found to be upregulated in 9-47-Ear and 4559M (9-47-Ear vs 9-47M, FC = 5.86, *p* = 1.19×10^−37^; 9-47-Ear vs 4559-Blood, FC = 1.89, *p* = 2.26×10^−7^; 9-47M vs 4559M, FC = 0.25, *p* = 3.55×10^−23^).

### Extensive RafR-Specific Rewiring of Host Transcriptional Responses to Infection

The measured murine transcriptional response represents the aggregate gene expression of all (host) cells present during pneumococcal infection in the lung. These include epithelial cells, endothelial cells of lung vasculature, smooth muscle cells, fibroblasts, activated and non-activated immune cells. The host transcriptional response was specific to the infecting pneumococcal strain (**Figure 3a**). Specifically, there was a diverging host response to the ear isolate (9-47-Ear, dark orange) and blood isolate (4559-Blood, dark purple). Interestingly, *rafR* swap in blood isolate background (4559M, light purple) mimics the lung response to the wild type ear isolate (9-47-Ear, dark orange); the two strains harbor the D249 *rafR* allele. Surprisingly, the *rafR* swap in the 9-47-Ear background (9-47M, light orange) which harbours the G249 allele, does not drive the host response to mimic those of the wild type 4559-Blood strain (dark purple) that also has the G249 allele, but rather towards a new, third position of genome-wide expression.

**Figure 3.**
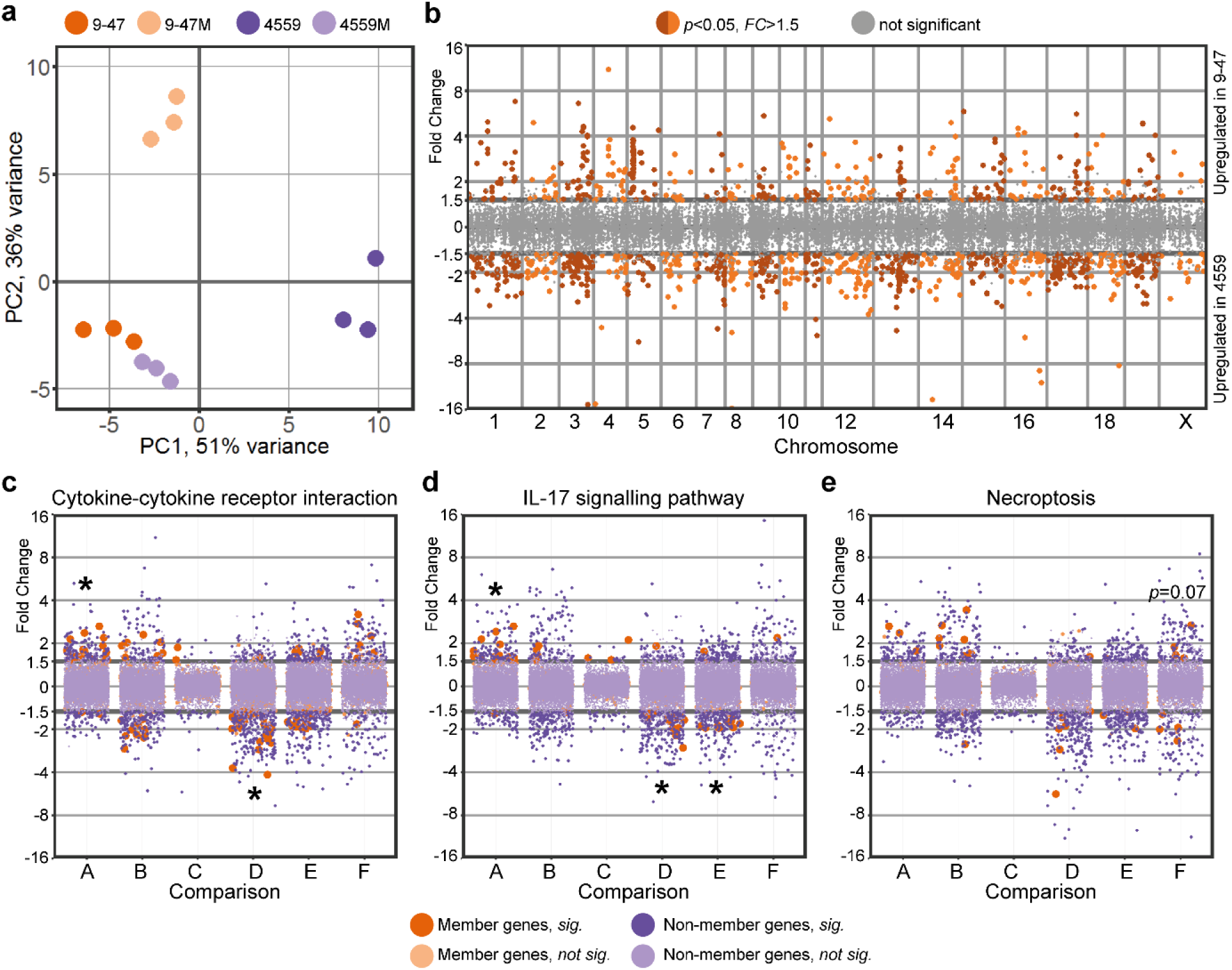
Single nucleotide polymorphism in pneumococcal *rafR* drives diverging host response. (**a)**. PCA plot illustrates murine lung response to the pneumococcal strains. Interestingly, host transcriptional response to *rafR* swap in blood isolate (4559M, light purple) is similar to the murine response to the original ear strain (9-47-Ear, dark orange). (**b)**. Differential gene expression of transcriptional response to pneumococcal ear and blood isolates shows a widespread transcriptional rewiring. Specifically, 433 genes are activated in response to infection by ear isolate (9-47-Ear) while 787 genes are activated (*FC*>1.5, *p*<0.05) by blood isolate (4559-Blood). Specific gene ontology terms are enriched in differentially expressed host genes in response to pneumococcal infection: cytokine-cytokine receptor interaction (**c**), interleukin-17 signaling pathway (**d**) and necroptosis (**e**). A: comparison between 9-47-Ear to 9-47M; B: 9-47-Ear to 4559-Blood; C: 9-47-Ear to 4559M; D: 9-47M to 4559-Blood; E: 9-47M to 4559M and F: 4559-Blood to 4559M. * denotes statistically significant functional enrichment for the indicated strain-strain comparison.

Genome-wide plotting of the murine transcriptional response to *S. pneumoniae* strain 9-47-Ear (ear isolate) and to strain 4559-Blood (blood isolate) shows an extensive rewiring of gene expression across the murine chromosomes, and the response is specific to the infecting strain (**Figure 3b**). Specifically, 433 murine genes are activated upon infection by the ear isolate 9-47-Ear (FC >1.5, *p* <0.05), while 787 genes are activated by infection with the blood isolate 4559-Blood (FC >1.5, *p* <0.05). Of the 9-47-Ear upregulated murine genes, only 37% were protein-coding genes, with the majority encoding pseudogenes and small RNA features. On the other hand, of the 787 4559-upregulated murine genes, 80% were protein-coding genes, while the rest encoded small RNA features. The 4559-upregulated genes include genes encoding proteins involved in multiple pathways such as general metabolism, peroxisome proliferator-activated receptor (PPAR) signaling, steroid hormone biosynthesis and cAMP signaling. Although both pneumococcal strains belong to the same capsular serotype and multi-locus sequence type (Amin et al., 2015), our data strongly suggest wildly diverging isolate-specific host responses during early infection.

In addition, *rafR* swap in the ear isolate background (9-47M) expressing the G249 *rafR* allele activates 271 murine genes (FC >1.5, *p* <0.05), while it represses 479 genes. The G249 *rafR*-activated genes include those involved in the Wnt signalling pathway (*Fzd2, Lgr6, Rspo1, Sost, Sox17, Wnt3a, Wnt7a*) and general calcium signalling pathway (*Adra1a, Adra1b, Adrb3, Cckar, Grin2c, P2rx6, Tacr1, Tacr2*). Conversely, 52% of the G249 *rafR*-repressed genes in lungs infected with 9-47M encode RNA features and 18 chemokines, chemokine ligands, interferons and interleukins. On the other hand, *rafR* swap in the blood isolate background (4559M) expressing the D249 *rafR* allele activates 328 murine genes (FC >1.5, *p* <0.05), and represses 472 genes. 73% of the D249 *rafR*-activated murine genes encode RNA features and 33 encode histone proteins. The activation of these histone proteins suggests a massive reorganization of gene regulation with numerous potential downstream impacts. In contrast, D249 *rafR*-repressed genes include genes encoding calmodulins (*Calm4, Calm13* and *Camk2a*) and phospholipases A2 (*Pla2g4b, Pla2g4d* and *Pla2g4f*).

Moreover, there are only 132 differentially expressed host genes (FC >1.5, *p* <0.05), in response to wild type 9-47-Ear compared to the response to strain 4559M (both having the D249 *rafR* allele), with 38 genes (FC >1.5, *p* <0.05), upregulated in strain 9-47-Ear and 94 genes in strain 4559M. Fascinatingly, the D249 *rafR* allele (strains 9-47-Ear and 4559M) is associated with a significant upregulation of RNA features, including antisense, intronic, long intergenic non-coding RNAs (lincRNAs) and micro RNAs (miRNAs). The resulting abundance of RNA species in murine cells upon pneumococcal infection has the potential for even more widespread transcriptional rewiring and fine-tuning of gene products later in the infection.

A quantified functional enrichment showed that certain gene functions are enriched in the murine response to pneumococcal strains. In particular, cytokine-cytokine receptor interaction is enriched in differentially expressed host genes because of *rafR* swap in the ear isolate background (**Figure 3c**, *comparison A*, 9-47-Ear vs. 9-47M, *p* = 9.5×10^−4^). Concurrently, the function is enriched in differentially expressed genes between mice infected with strain 9-47M and those infected with strain 4559-Blood (*comparison D, p* = 1.8 ×10^−4^). Since both strains harbor the G249 *rafR* allele, the differentially expressed genes encoding for cytokines and cytokine receptors are most likely attributable to unrelated genetic differences between the clinical isolates. Interestingly, this function is not enriched in differentially expressed genes between the *rafR* swap in the blood isolate background (4559M) and the wild type ear isolate (9-47-Ear, *comparison C*), both of which have the D249 *rafR* allele. Genes encoding chemokine ligands (*Cxcl2, Cxcl3, Cxcl10* and *Ccl20*), interleukin 17F (*Il17f*), interferon beta (*Ifnb1*) and a receptor of TNF (*Tnfrsf18*) are the common differentially expressed genes in lungs of mice infected with 9-47-Ear, 9-47M and 4559-Blood, with ascending expression from responses to 9-47M, 9-47-Ear and 4559-Blood. Other genes encoding chemokine ligands (*Ccl3, Ccl4, Ccl17, Ccl24, Cxcl5, Cxcl11* and *Xcl1*), interleukins (*Il1rn* and *Il13ra2*) and interferon gamma (*Ifng*) are more highly expressed in the ear isolate-infected lung (9-47-Ear) than in lungs infected by the *rafR* swap ear isolate (9-47M). Finally, genes encoding chemokine receptors (*Ccr1* and *Ccr6*), interleukin receptors (*Il1r2, Il10ra, Il17a, Il18rap, Il20ra, Il20rb, Il22* and *Il23r*), and interleukins (*Il1f5, Il1f6, Il1f8* and *Il6*) are more highly expressed in lungs infected by the blood isolate (4559-Blood) compared to the *rafR* swap in the ear isolate background (9-47M).

Interleukin 17, as part of the cytokine response, activates multitudes of downstream targets in defense against infectious agents (Onishi and Gaffen, 2010), and thus plays a central role in host response against pneumococcal infection. Here, we observe the same pattern of diverging activation among murine response to the pneumococcal strains (**Figure 3d**), with IL-17 associated genes being enriched in differentially expressed genes among the host transcriptional response to the *rafR* swap in the ear isolate background (*comparison A*, 9-47-Ear vs. 9-47M, *p* = 9.5×10^−4^). These genes are also enriched amongst the host response to pneumococcal strains with the G249 *rafR* allele (*comparison D*, 9-47M vs. 4559-Blood, *p* = 7.5×10^−4^) and to the *rafR* swap cognates (*comparison E*, 9-47M vs. 4559M, *p* = 0.022). However, there was no enrichment of IL-17 associated genes amongst the host response to pneumococcal strains with D249 *rafR* allele (*comparison C*, 9-47-Ear vs. 4559M). Common differentially expressed genes of this function include genes encoding interleukin 17F (*Il17f*) and chemokine ligands (*Cxcl2, Cxcl3* and *Ccl20*), with ascending expression level of response to 9-47M, 9-47-Ear and 4559-Blood. Specifically, the products of these genes regulate the recruitment of neutrophils and activate immune responses to extracellular pathogens.

In addition to the above, necroptosis, a programmed cell death, is almost significantly enriched (*p* = 0.07) in differentially expressed murine genes because of the *rafR* swap in the blood isolate background (**Figure 3e**, *comparison F*, 4559-Blood vs. 4559M). Genes encoding for histone cluster 2 (*Hist2h2ac, Hisr2h2aa1, Hist2h2aa2* and *H2afx*) are more highly expressed in murine lungs infected with the blood isolate (4559-Blood), while those encoding phospholipases A2 (*Pla2g4b, Pla2g4f* and *Pla2g4d*) and a subunit of calcium/calmodulin-dependent protein kinase II (*Camk2a*) are more highly expressed in the transcriptional response to the *rafR* swap in the blood isolate background (4559M).

### Validation of Host/Pathogen Transcriptomics

To validate the findings from the Dual RNA-seq, quantitative real time RT-PCR was performed on the RNA samples from the lungs 6 h post-infection. 19 pneumococcal and 18 murine genes were chosen for this validation, with the primers used listed in **Table 2**. Log_2_ FCs were compared between the Dual RNA-seq and the qRT-PCR datasets for 9-47-Ear vs 9-47M, 9-47-Ear vs 4559-Blood, 9-47M vs 4559M and 4559-Blood vs 4559M, for each gene, totalling 76 pneumococcal and 72 murine comparisons. A high degree of correlation was observed for both pneumococcal (*R*^*2*^ > 0.81, Pearson) and murine genes (*R*^*2*^ > 0.73, Pearson) **(Figure 4)**.

**Table 2.**
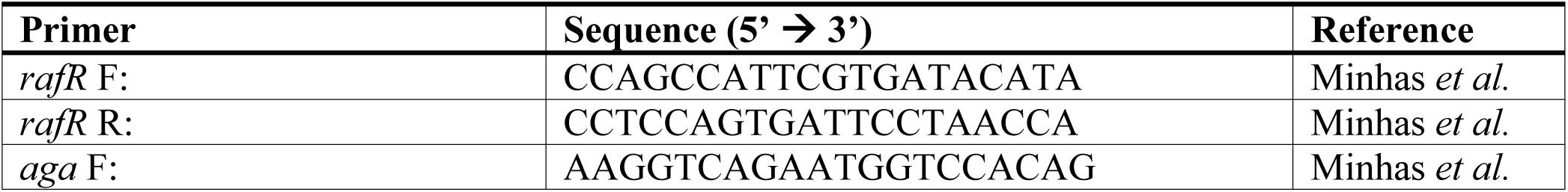
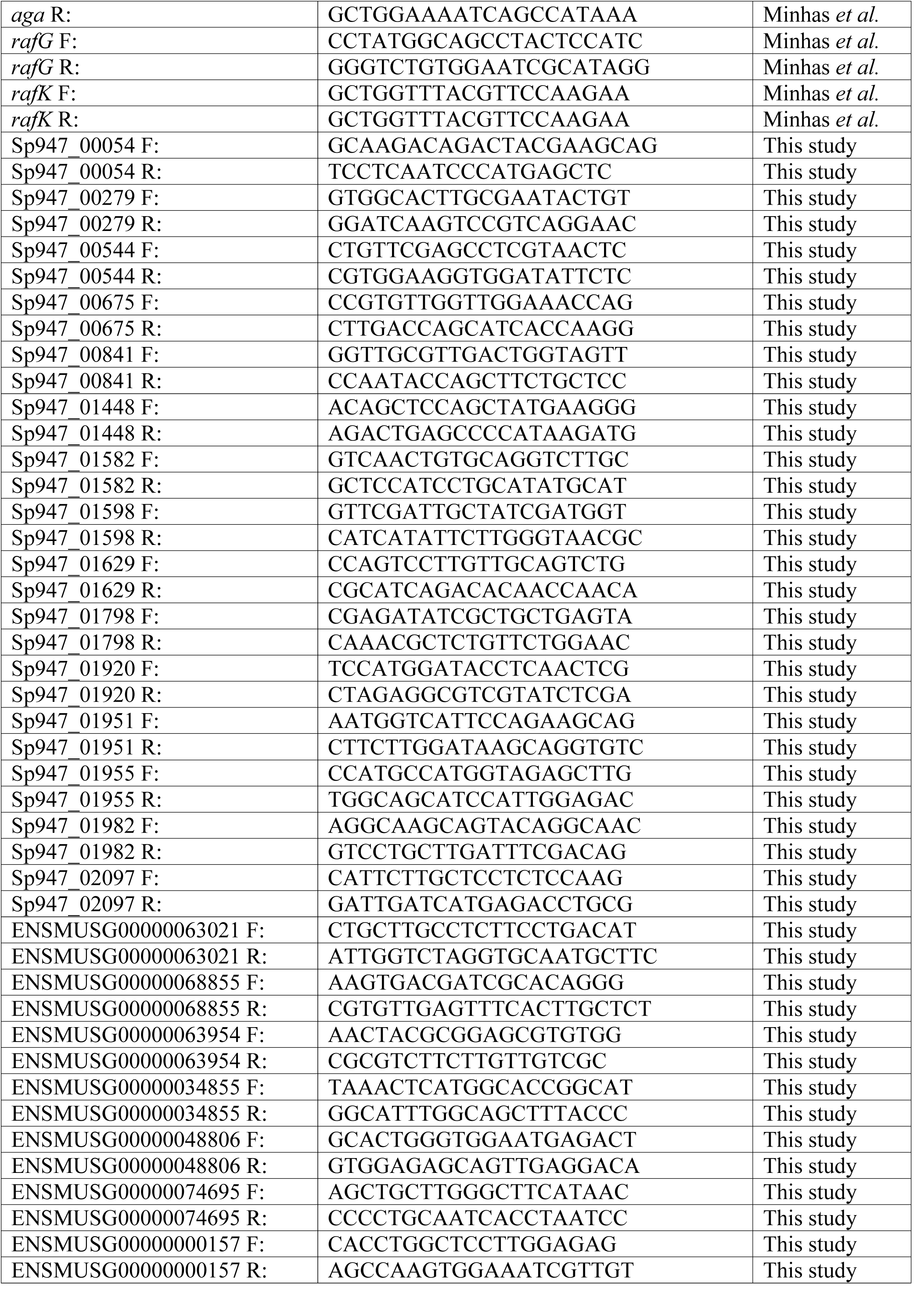
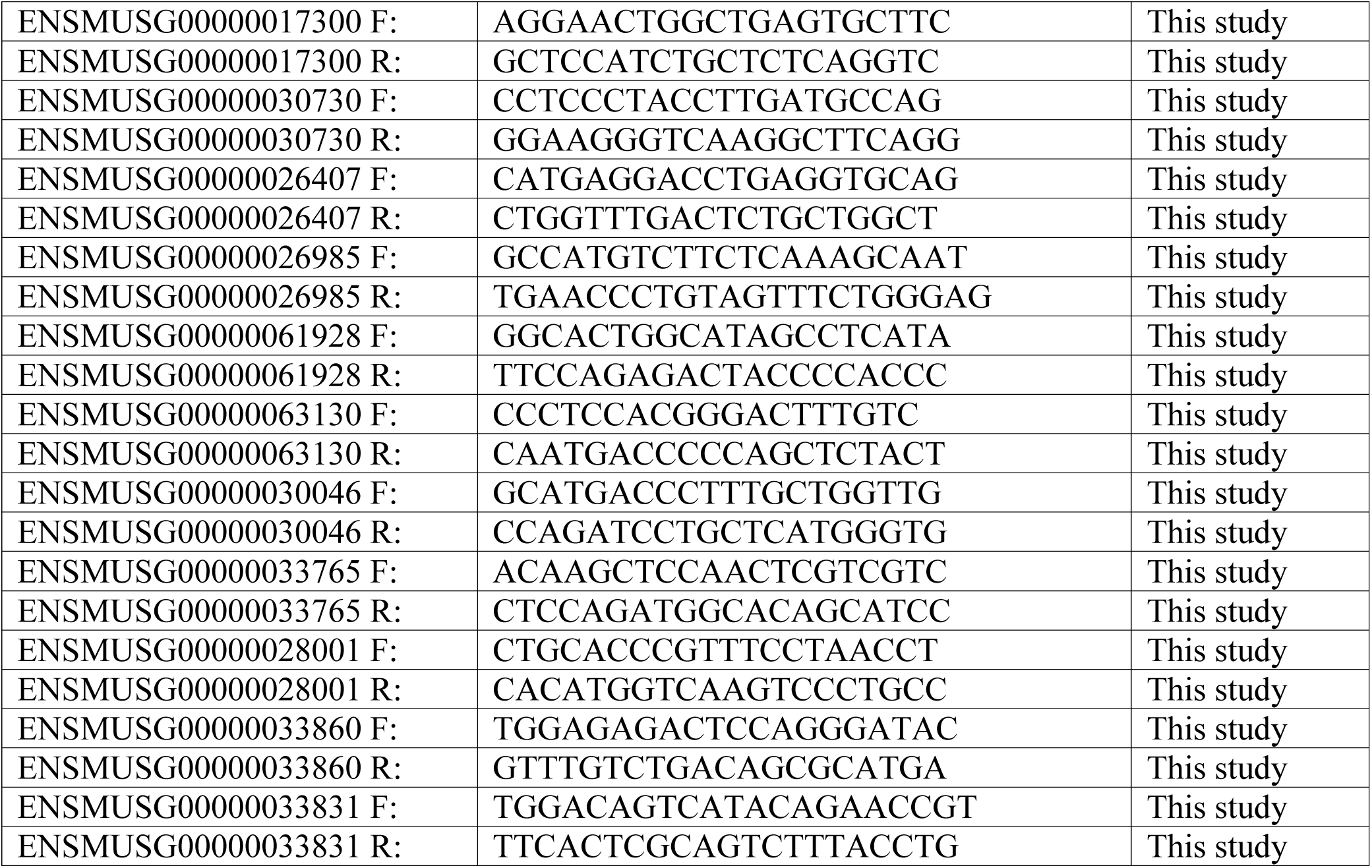
Oligonucleotide primers used in this study.

**Figure 4.**
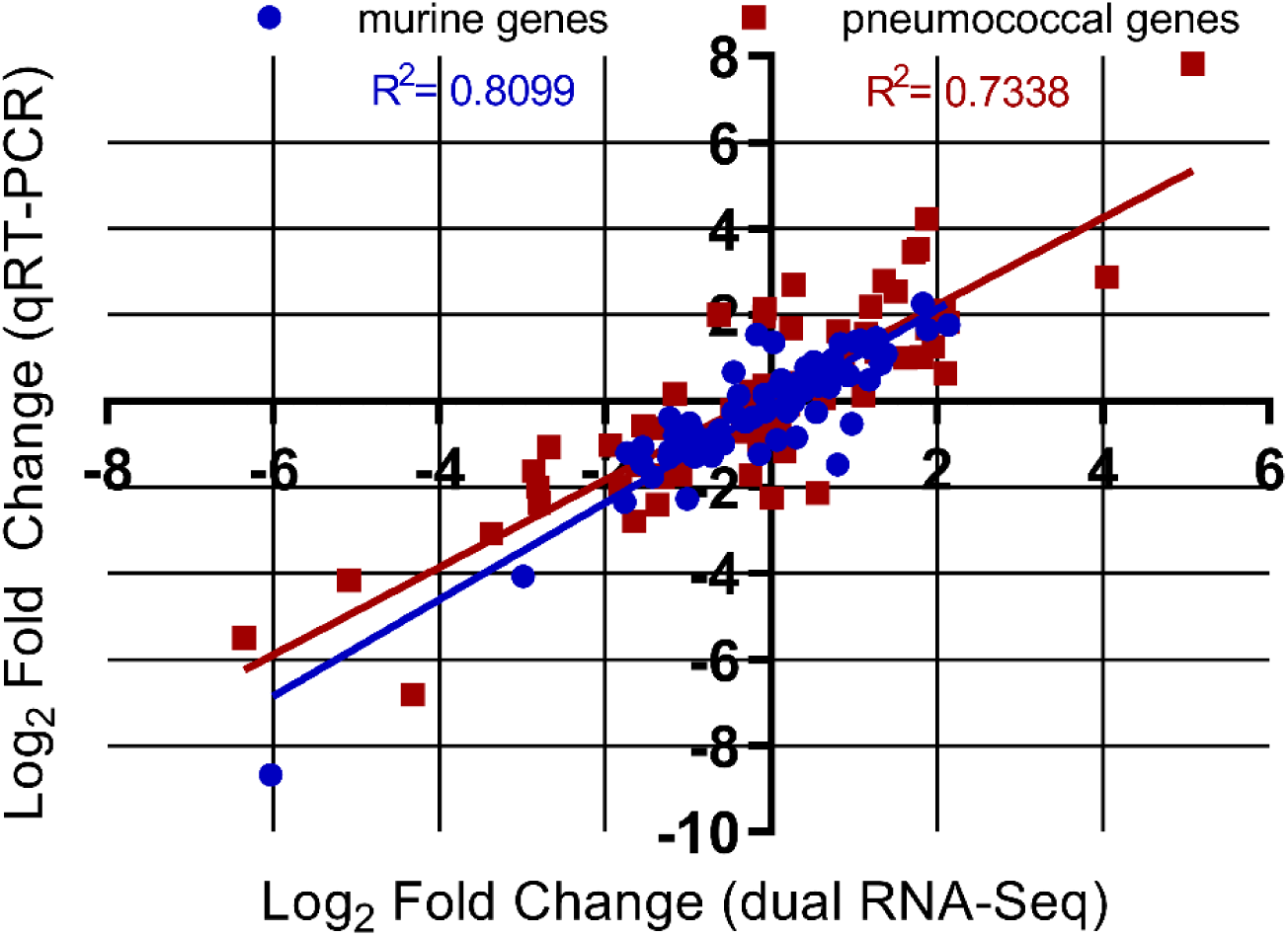
Gene expression values from the dual RNA-seq were confirmed by qRT-PCR, using the same isolated RNA used for the dual RNA-seq. 18 murine and 19 pneumococcal genes were chosen as validation targets. Log_2_ fold changes were plotted from qRT-PCR against dual RNA-seq log fold changes for 9-47-Ear vs 4559-Blood, 9-47-Ear vs 9-47M, 9-47M vs 4559M and 4559-Blood vs 4559M comparisons. A total of 72 murine and 76 pneumococcal comparisons were plotted, with a high degree of correlation observed for both species (*R*^*2*^ > 0.73, Pearson).

### Immune Cell Subsets Present in Infected Lung Tissue

The host RNA-seq data represent the pooled transcriptional responses of all cell types present in the lungs at the time of RNA extraction. Thus, at least some of the transcriptomic differences may be attributable to alterations in the relative abundance of given cell types, for example by differential recruitment of immune cell subsets to the site of infection. Accordingly, flow cytometry was used to quantify immune cell subsets present in lung tissue 6 h after infection with either 9-47-Ear, 4559-Blood, 9-47M or 4559M. The surface marker staining panel used (**Table 2**) allowed the identification and enumeration of natural killer (NK) cells, neutrophils, eosinophils, inflammatory monocytes (iMono), resident monocytes (rMono), alveolar macrophages (AMФ), interstitial macrophages (iMФ), CD11b-negative dendritic cells (CD11b-DC), CD11b-positive dendritic cells (CD11b+ DC), T cells and B cells (Yu et al., 2016). Of these, neutrophils, by far the most abundant cell type, were present in significantly higher numbers in murine lungs infected with 9-47-Ear (vs 4559-Blood, *p* < 0.01; vs 9-47M, *p* <0.05) and 4559M (vs 4559-Blood, *p* < 0.05) (**Figure 5**), both of which have the D249 *rafR* allele. NK cells were also found to be significantly higher in lungs infected with 9-47-Ear (vs 4559-Blood, *p* < 0.01 and vs 9-47M, *p* < 0.05), while eosinophils were raised in 4559M infected lungs (vs 4559-Blood, *p* < 0.05) (**Figure 5)**.

**Fig.5.**
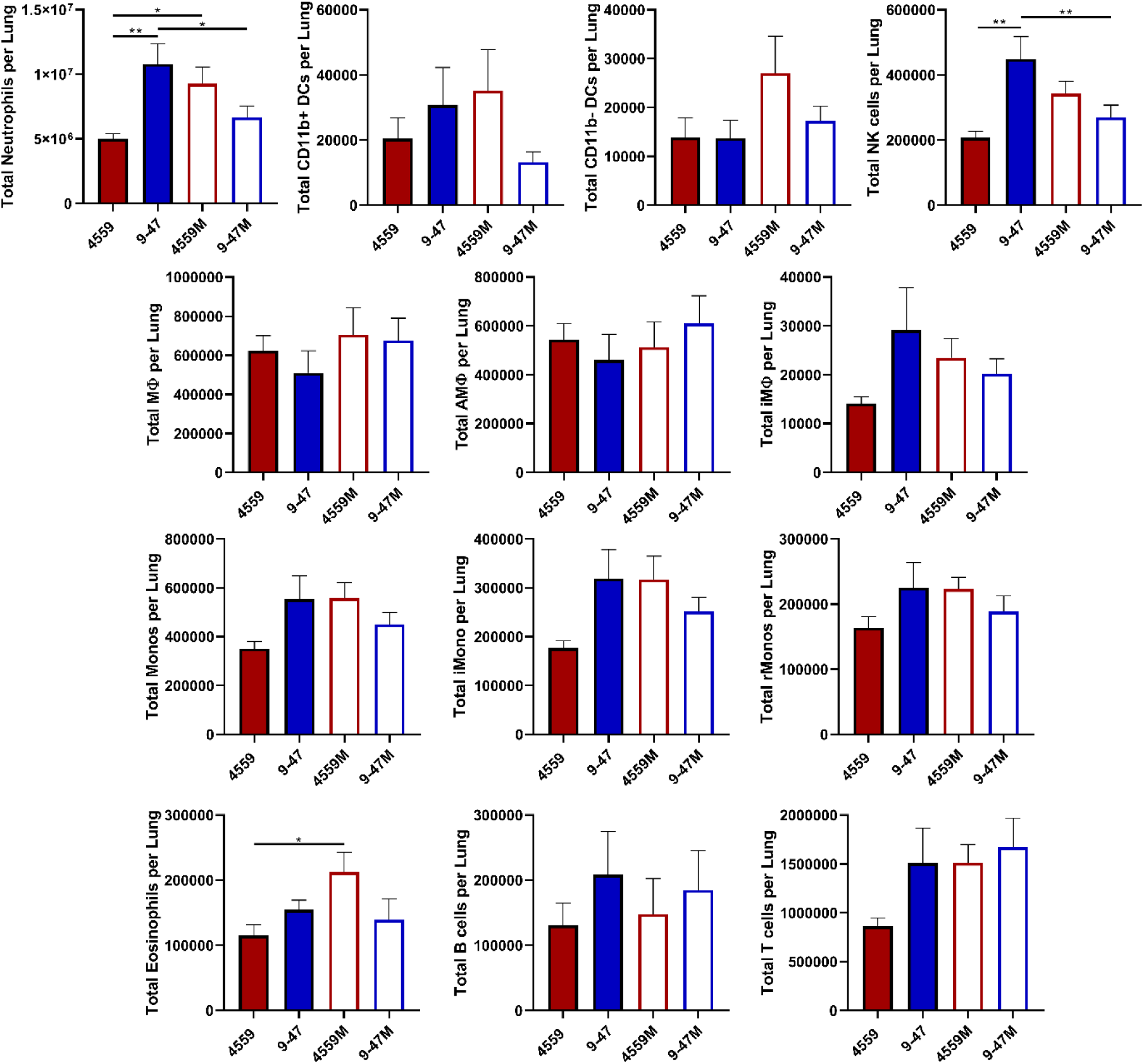
Quantification of immune cell subsets in murine lungs 6 h post infection. Groups of 8 mice per strain were challenged with either 9-47-Ear, 4559-Blood, 9-47M or 4559M. Single cell lung suspensions were prepared and stained with antibodies against various surface markers (**Table 1**) and analyzed by flow cytometry. Populations enumerated include: natural killer (NK) cells, neutrophils, eosinophils, inflammatory monocytes (iMono), resident monocytes (rMono), alveolar macrophages (AMФ), interstitial macrophages (iMФ), CD11b-negative dendritic cells (CD11b-DC), CD11b-positive dendritic cells (CD11b+ DC), T cells and B cells. Graphs shown represent pooled data from two independent experiments. All quantitative data are presented as mean ± S.E.M (*n* = 16 for each group), analyzed by one-way ANOVA (\**P* < 0.05; \*\**P* < 0.01).

### Impact of Neutrophil Depletion and IL-17A Neutralization on Pneumococcal Persistence in Murine Lungs

As shown above, neutrophils were more abundant in murine lungs infected with 9-47-Ear and 4559M, the strains containing the D249 *rafR* allele that are cleared from the lungs by 24h. This suggests that the recruitment and presence of neutrophils is crucial for bacterial clearance from the lung and differential neutrophil recruitment might be the underlying mechanism for the observed RafR-dependent tropism. To test this, we investigated the importance of neutrophils for persistence of pneumococci in the lungs in the murine IN challenge model. Injection of anti-mouse Ly6G antibody was used to deplete neutrophils in 32 mice, alongside an isotype control group treated with rat IgG2a. Neutrophil depletion was confirmed in the blood prior to challenge, with a 76.35% decrease in neutrophils seen in the anti-mouse ly6G treated mice, relative to the isotype control treated group (*p* < 0.0001) **(Figure 6A)**. Mice were then challenged with 10^8^ CFU of each strain, for both treatment groups. Bacterial loads were quantified in the nasopharynx and lungs 24 h post-challenge. No significant differences in bacterial numbers in the nasopharynx were seen between strains within each treatment group **(Figure 6B)**. Also, for both treatments, the numbers of bacteria in the lungs infected with 4559-Blood and 9-47M were significantly higher than 9-47-Ear and 4559 **(Figure 6C)**, which is consistent with our previous findings (Minhas et al., 2019). However, the anti-Ly6G-treated groups showed significantly higher lung bacterial loads compared to their respective isotype controls: 4559-Blood anti-Ly6G vs 4559-Blood control (*p* < 0.001), 947-Blood anti-Ly6G vs 947-Blood control (*p* < 0.01), 4559M anti-Ly6G vs 4559M control (*p* < 0.01) and 947M anti-Ly6G vs 947M control (*p* < 0.05) **(Figure 6C)**. Importantly, the lung bacterial loads of anti-Ly6G-treated 9-47-Ear and 4559M groups were not significantly different to the isotype control-treated 4559-Blood group **(Figure 6B)**. Thus, restriction of neutrophil infiltration into the lungs by depleting circulating neutrophils in mice challenged with the strains expressing the D249 *rafR* allele resulted in enhanced lung bacterial loads at 24 h similar to that seen in untreated mice challenged with the strains expressing the G249 *rafR* allele.

**Figure 6.**
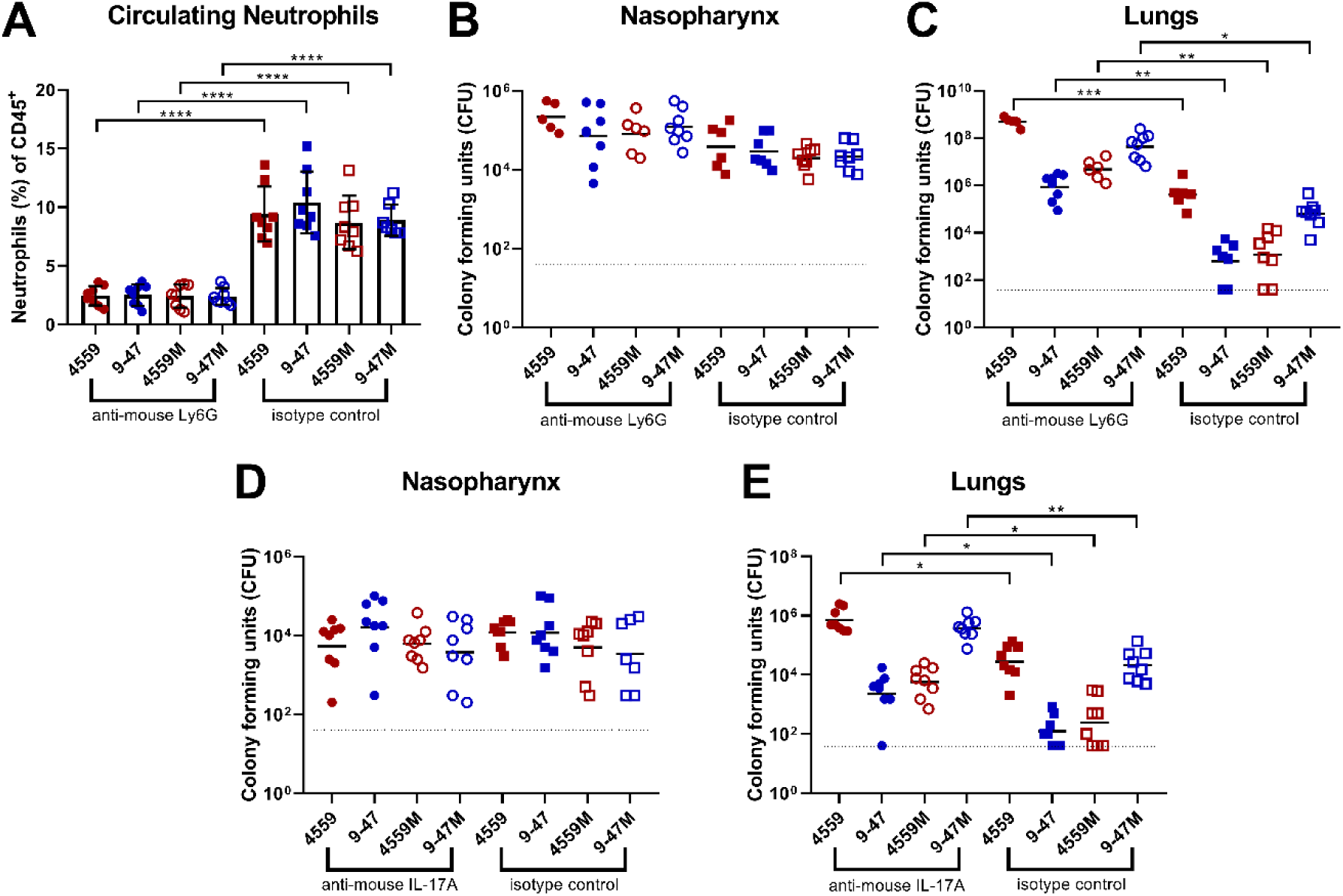
Impact of neutrophil depletion or anti-IL-17A on pneumococcal virulence. Groups of 8 mice were treated with either 350 μg of rat anti-mouse Ly6G or rat IgG2a isotype control, one and two days prior to pneumococcal challenge (B & C), or with IL-17A or mouse IgG1 isotype control (D & E), one day before, 2 h before and 6 h after intranasal challenge (see materials and methods). (A) Percentage of circulating neutrophils relative to live (CD45^+^) cells were calculated. Differences in circulating neutrophils between groups are indicated by asterisks: ****, *P* < 0.0001, by one-way ANOVA. (B, C, D & E) 24 h post-infection, numbers of pneumococci in the nasopharynx and lungs were quantitated (see Materials and Methods). NB: n is <8 for some groups because didn’t survive the challenge procedure, or until the time of harvest. Viable counts (total CFU per tissue) are shown for each mouse at each site; horizontal bars indicate the geometric mean (GM) CFU for each group; the broken line indicates the threshold for detection. Differences in GM bacterial loads between groups are indicated by asterisks: *, *P* < 0.05, **, *P* < 0.01, ***, *P* < 0.001, by unpaired *t*-test.

Given the known involvement of IL-17 in neutrophil recruitment into the lungs after infection (Lindén et al., 2005; McCarthy et al., 2014; Ritchie et al., 2018; Stoppelenburg et al., 2013), we also investigated the in vivo significance of the *rafR*-mediated differential expression of IL-17-associated genes between the various *S. pneumoniae* strains described above. Groups of mice were injected with anti-mouse IL-17A antibody, or a control murine IgG1 antibody, before and after pneumococcal challenge. Bacterial loads were quantified in the nasopharynx and lungs 24 h post-challenge. Again, no significant differences between strains in bacterial numbers in the nasopharynx were seen within each treatment group **(Figure 6D)**. However, similar to the results obtained using anti-Ly6G, groups treated with anti-IL-17A showed significantly higher bacterial numbers in the lungs compared to their respective isotype controls; 4559-Blood anti-IL-17A vs 4559-Blood control (*p* < 0.05); 947-Blood anti-IL-17A vs 947-Blood control (*p* < 0.05); 4559M anti-IL-17A vs 4559M control (*p* < 0.05); and 947M anti-IL-17A vs 947M control (*p* < 0.01) **(Figure 6E)**. Nevertheless, the impact of anti-IL-17 treatment on lung bacterial loads was not quite as dramatic as that of anti-Ly6G, as the number of bacteria in the lungs of anti-IL-17A-treated 9-47-Ear and 4559M groups remained significantly lower relative to the isotype control treated 4559-Blood group (both *p* < 0.05) **(Figure 6E)**. Together, these results show that pneumococcal strains carrying the D249 *rafR* allele cause a rapid influx of neutrophils, partly controlled by IL-17 expression in the host, leading to clearance from the lung, while the G249 *rafR* strains manage to remain ‘stealthy’ and hence can persist.

## Conclusions

In this study, we have used a dual RNA-seq approach, validated by qRT-PCR, to elucidate the complex interspecies interactions between murine lung cells and infecting *S. pneumoniae* blood and ear isolates that are closely related (same capsular serotype and ST type), but exhibit distinct virulence phenotypes in accordance with their original clinical isolation site. These differences are largely, but not completely, driven by a D249G SNP in the raffinose pathway transcriptional regulator gene *rafR*, which extensively impacts the bacterial transcriptome in the lung environment. The SNP affects expression of genes encoding multiple transmembrane transporters, including those for various sugars, and fine-tunes pneumococcal carbohydrate metabolism. This indicates that the differential expression of sugar catabolism pathways provides specific advantages in distinct host niches, implying differential niche-specific availability of one carbohydrate source versus another. Free sugars are in low abundance in the upper respiratory tract, but *S. pneumoniae* expresses a range of surface-associated exoglycosidases enabling it to scavenge constituent sugars (including galactose, *N*-acetylglucosamine, sialic acid and mannose) from complex host glycans present in respiratory secretions and on the epithelial surface (King et al., 2006; Shelburne et al., 2008; Buckwalter and King, 2012; Paixão et al., 2015; Robb et al., 2017). On the other hand, glucose is readily available in the blood and also in inflamed tissues, implying a marked alteration in the availability of this preferred carbohydrate source as invasive disease progresses (Philips et al., 2003). All these variations, and the downstream consequences thereof, are ultimately sensed by host cells, including epithelial and immune cells, resulting in the observed divergence of host response to the various strains, particularly with respect to expression of genes encoding cytokine and chemokine ligands and receptors, as well as those associated with programmed cell death.

Examination of the nature of the host response has provided important clues regarding the mechanism whereby the *rafR* SNP impacts virulence phenotype. By way of example, the dual RNA-seq data showed that expression of IL-17 related genes was enriched in mice infected with 9-47-Ear and 4559M, the strains that express the D249 *rafR* allele and which are cleared from the lungs by 24 h post-challenge. It is well known that IL-17 drives neutrophil recruitment into the lungs after infection (Lindén et al., 2005; McCarthy et al., 2014; Ritchie et al., 2018; Stoppelenburg et al., 2013). Additionally, neutrophil extravasation genes were shown to be upregulated in murine lungs 48 h post pneumococcal challenge (Ritchie and Evans, 2019). Indeed, we have shown here that neutrophils were present in the lungs 6 h post challenge at significantly higher numbers in mice infected with 9-47-Ear and 4559M compared with the strains expressing the G249 *rafR* allele **(Figure 5)**, as predicted by the dual RNA-seq data. Moreover, we went on to show that neutrophil depletion by treatment with anti-Ly6G increased bacterial numbers in the lungs of mice, relative to the isotype controls. Strikingly, pneumococcal numbers in the lungs of anti-Ly6G treated mice infected with 9-47-Ear and 4559M were not significantly different to that for isotype control-treated 4559-Blood-infected mice **(Figure 6C)**. In vivo neutralization of IL-17A also resulted in an increase in bacterial loads in the lungs of mice, relative to the isotype controls, for all challenge strains **(Figure 6E)**, although not to the same extent as seen with for neutrophil-depleted mice **(Figure 6C)**. The difference between the impact of IL-17A neutralization vs neutrophil depletion is likely due to the action of alternative neutrophil recruitment pathways (Craig et al., 2009; Peñaloza et al., 2015). Our findings demonstrate that the *rafR* SNP examined in this study has a wide spread effect on both the bacterial and host transcriptomes, with the strains expressing the G249 allele triggering a strong pro-inflammatory IL-17 response in the lungs post-infection. This response leads to an influx of neutrophils to the lungs, resulting in the clearance of bacteria. Conversely, expression of the D249 *rafR* allele results in a more subdued IL-17 host response, allowing for bacterial persistence in the lungs. Thus, our findings clearly indicate that modulation of neutrophil recruitment during the early stage of infection plays a key role in the capacity of a given *S. pneumoniae* strain to persist in the lungs, and the nature of disease ultimately caused by it.

Our previous studies have shown that in spite of early clearance from the lung, strains 9-47 Ear and 4559M, expressing the G249 *rafR* allele, have an enhanced capacity to spread to and/or proliferate in the ear and brain compartments (Amin et al., 2015; Minhas et al., 2019). It is not known whether differential carbohydrate metabolism better adapts these strains to available carbohydrate sources in these niches, or whether altered host pro-inflammatory responses contribute to ascension of the Eustachian tube or penetration of the blood-brain barrier. Unfortunately, the total numbers of pneumococci present in these niches are too low for pathogen-host transcriptomic analyses using available technologies.

Intra-species variation in virulence phenotype is a common feature of pathogenic microorganisms, which by nature are genetically diverse. *S. pneumoniae* is an exemplar of such diversity comprising at least 98 capsular serotypes superimposed on over 12,000 MLST types, and with a core genome that accounts for only 70% of genes (Weiser et al., 2018). Nevertheless, stark differences in pathogenic profile can result from the smallest of genetic differences between strains, as exemplified by the profound impact of a single SNP on both bacterial and host transcriptomes reported in this study.

## ACKNOWLEDGEMENTS

We thank V Benes (GeneCore, EMBL, Heidelberg) for his continuing support in library preparation and sequencing. We would like to acknowledge the Center for Information Technology of the University of Groningen for their support and for providing access to the Peregrine high-performance computing cluster. We would also like to thank Timona Tyllis and Todd Norton for their assistance in acquiring the flow cytometry samples, as well Alexandra Tikhomirova for her assistance with the murine experiments. This work was supported by the Swiss National Science Foundation (SNSF) (project grant 31003A_172861) to J.W.V., a National Health and Medical Research Council (NHMRC) Program Grant 1071659 to J.C.P. and a University of Adelaide Beacon Fellowship to C.T. The funders had no role in study design, data collection and interpretation, or the decision to submit the work for publication.

## AUTHOR CONTRIBUTIONS

Conceptualization: V.M., J.C.P., J.W.V. and C.T.; Methodology: V.M., R.A., J.W.V. and C.T.; Formal analysis: V.M., R.A., L.J.M., I.C., S.R.M., J.C.P., J.W.V. and C.T.; Investigation: V.M., R.A., H.W., S.C.D., K.T.M., and C.T.; Writing – original draft: V.M., R.A., J.C.P., J.W.V. and C.T.; Writing – review and editing: all authors; Supervision: J.C.P., J.W.V., and C.T.; Funding acquisition: J.C.P., J.W.V., and C.T.

## MATERIALS AND METHODS

### Bacterial Strains and Growth Conditions

*S. pneumoniae* strains used in this study are listed in the (**Table 1**). Cells were routinely grown in serum broth (SB) as required. Bacteria were plated on Columbia agar supplemented with 5% (vol/vol) horse blood (BA) and incubated at 37°C in 5% CO_2_ overnight.

### Intranasal Challenge of Mice and Extraction of RNA

Animal experiments were approved by the University of Adelaide Animal Ethics Committee. Groups of 12 outbred 6-week-old female Swiss (CD-1) mice (48 in total), were anesthetized by intraperitoneal injection of pentobarbital sodium (Nembutal) and challenged intranasally (IN) with 50 µl of bacterial suspension containing approximately 1 × 10^8^ CFU in SB of 4559-Blood, 9-47-Ear, 4559M or 9-47M. The challenge dose was confirmed retrospectively by serial dilution and plating on BA. Mice were euthanized by CO_2_ asphyxiation at 6 h and lungs placed in 1 ml TRIzol (Thermo Fisher). RNA was then extracted using acid-phenol-chloroform-isoamyl alcohol (125:21:1; pH 4.5; Ambion) and purified using the RNeasy minikit (Qiagen). For subsequent dual RNA-seq analyses, there were three replicates per strain, with each replicate derived from the lungs of four mice.

### RNA Library Preparation and Sequencing

RNA quality was checked using chip-based capillary electrophoresis. Samples were then simultaneously depleted from murine and pneumococcal ribosomal RNAs by dual rRNA-depletion as previously described (Aprianto et al., 2016). Stranded cDNA library preparation was performed according to the prescribed protocol (Illumina, US). Sequencing was performed for twelve samples in one lane of Illumina NextSeq 500, High Output Flowcell in 85 single end mode. Libraries were demultiplexed and analyzed further. Raw libraries are accessible at https://www.ncbi.nlm.nih.gov/geo/ with the accession number GSE123982.

### Sequence Data Analysis

Quality of raw libraries was checked (Andrews and Babraham Bioinformatics, 2010) (FastQC v0.11.8, Babraham Bioinformatics, UK). In order to improve the quality of alignment, we trimmed the reads (Bolger et al., 2014) using the following criteria: (i) removal of adapter sequence, if any, based on TruSeq3-SE library, (ii) removal of low quality leading and trailing nucleotides, (iii) a five-nucleotide sliding window was created for surviving reads, in which the average quality score must be above 20 and (iv) minimum remaining length must be above 50 (Trimmomatic v0.38). The quality of trimmed reads were confirmed using FastQC (Andrews and Babraham Bioinformatics, 2010).

As reference genomes, we created chimeric genomes by concatenating the in-house generated *S. pneumoniae* circular genome into the genome of Mus musculus (ENSEMBL, release 94, downloaded 9 October 2018). The corresponding annotation file was downloaded at the same time. The chimeric genome containing genome of strain 9-47-Ear was used as reference to align libraries from lung infected by strain 9-47-Ear (and its corresponding swap mutant) while the 4559-Blood chimeric genome was used to align 4559-Blood libraries. Notably, genome of *S. pneumoniae* isolate 4559-Blood has a plasmid. Alignment was performed by RNA-STAR (v2.6.0a) (Dobin et al., 2013) with the following options: (i) alignIntronMax 1 and (ii) sjdbOverhang 84. The aligned reads were the summarized (featureCount v1.6.3) according to the chimeric annotation file in stranded, multimapping (-M), fractionized (--fraction) and overlapping (-O) modes (Liao et al., 2014). In order to compare gene expression between strains from ear and blood isolate backgrounds, we prepared a common pneumococcal annotation file using Mauve v20150226 (Darling et al., 2004). ommon genes between 9-47-Ear and 4559-Blood were defined as having common coverage at least 90% and identity at least 90%. This single-pass alignment was selected onto chimeric genome was selected to minimize false discovery rate. However, due to this approach, we have to adjust the summarizing process, taking into account the overlapping nature of bacterial genes and its organization into operon structures.

We then analyzed host and pathogen libraries separately in R (R v3.5.2). Since reads coming from pneumococcal genes encoding bacterial rRNA dominate the pathogen libraries (average 64.5%, range between 61.4 to 67.3%), we excluded these pneumococcal ribosomal RNA reads from downstream analysis, but we did not do the same exclusion to reads coming murine ribosomal RNA genes due to effective rRNA depletion. Differential gene analysis was performed by DESeq2 v1.22.1 (Love et al., 2014) and genome-wide fold change was calculated within host and pathogen libraries for every two possible comparisons: strains 9-47-Ear to 9-47M, strains 9-47-Ear to 4559-Blood, strains 9-47-Ear to 4559M, strains 9-47M to 4559-Blood, strains 9-47M to 4559M and strains 4559-Blood to 4559M. Value of fold change was set to zero if the corresponding adjusted p-value (*padj*) is reported to be NA.

### Quantitative Real Time RT-PCR

Differences in levels of gene expression observed in the dual RNA-seq data were validated by one-step relative quantitative real-time RT-PCR (qRT-PCR) in a Roche LC480 real-time cycler essentially as previously described (Mahdi et al., 2008). The same RNA that was used for the dual RNA-seq was used in the RT-PCR validation. 19 pneumococcal genes and 18 murine genes were chosen for the validation. The specific primers used for the various genes are listed in **Table 2** and were used at a final concentration of 200 nM per reaction. As an internal control, primers specific for *gyrA* were employed. Amplification data were analysed using the comparative critical threshold (2^−^CT) method (Livak and Schmittgen, 2001).

### Flow Cytometry Analysis of Infected Murine Lungs

Groups of 8 outbred 6-week-old female Swiss (CD-1) mice (32 in total) were anesthetized and challenged with the bacterial suspension as outlined above in the total RNA extraction method. Mice were euthanized by CO_2_ asphyxiation at 6 h, then lungs were finely macerated in 1 mL prewarmed digestion medium (DMEM + 5% FCS, 10 mM HEPES, 2.5 mM CaCl_2_, 0.2 U mL^−1^ penicillin/gentamicin, 1 mg mL^−1^ collagenase IA, 30 U mL^−1^ DNase) and incubated at 37°C for 1 h with mixing every 20 min. Single cells were then prepared for acquisition on a BD LSRFortessa X20 flow cytometer as previously described (David et al., 2019). The single cell suspensions were stained using antibodies against surface markers listed in **Table 1**, allowing the enumeration of a number of immune cell subsets, as previously described (Yu et al., 2016).

### Neutrophil Depletion and IL-17A Blockade and Bacterial Load Quantification

Groups of 8 outbred 6-week-old female Swiss (CD-1) mice (64 in total) were intraperitoneally administered with either 350 μg of rat anti-mouse Ly6G or rat IgG2a isotype control antibodies, one and two days prior to pneumococcal challenge, or 200ug of either monoclonal anti-mouse IL-17A or mouse IgG1 isotype control antibodies one day prior to, 2 h before and 6 h after pneumococcal challenge. Mice were also cheek bled on day of challenge for confirmation of depletion of Ly6G-positive cells via flow cytometry, as previously described (Faget et al., 2018). Mice were then anesthetized and challenged with the bacterial suspension as outlined above in the total RNA extraction method, for each treatment group. Mice were euthanized by CO_2_ asphyxiation at 24 h, then nasopharynx and lung tissue samples were harvested and pneumococci enumerated in tissue homogenates as described previously via serial dilution and plating on BA containing gentamicin (Trappetti et al., 2011).

## QUANTIFICATION AND STATISTICAL ANALYSIS

For the RNA-seq data, enrichment tests to assess enrichment were performed by the built-in function, *fisher.test().* Corresponding *p*-values of the enrichment test were adjusted by Bonferroni correction. Resultant figures encompass data derived from three replicates per group, with each replicate derived from lungs of four mice. All other data are presented as mean ± standard error of mean (SEM) or geometric mean, and were analyzed by two-tailed unpaired Student’s *t*-test, one way ANOVA or Pearson correlation coefficient, using Prism v8.0d (GraphPad). Statistical significance was defined as *P* < 0.05. Data presented in figures are representative of at least two independent *in vivo* experiments, or at least 3 independent *in vitro* experiments.

## DATA AVAILABILITY

The transcriptomic datasets are available in the GEO repository, accession number GSE123982.

## REFERENCES

Amin, Z., Harvey, R.M., Wang, H., Hughes, C.E., Paton, A.W., Paton, J.C., and Trappetti, C. (2015). Isolation site influences virulence phenotype of serotype 14 Streptococcus pneumoniae strains belonging to multilocus sequence type 15. Infection and Immunity 83, 4781–4790.

Andrews, S., and Babraham Bioinformatics (2010). FastQC: A quality control tool for high throughput sequence data. Manual.

Aprianto, R., Slager, J., Holsappel, S., and Veening, J.-W. (2016). Time-resolved dual RNA-seq reveals extensive rewiring of lung epithelial and pneumococcal transcriptomes during early infection. Genome Biol. 17, 198.

Bidossi, A., Mulas, L., Decorosi, F., Colomba, L., Ricci, S., Pozzi, G., Deutscher, J., Viti, C., and Oggioni, M.R. (2012). A functional genomics approach to establish the complement of carbohydrate transporters in *Streptococcus pneumoniae*. PLoS ONE 7, e33320.

Bolger, A.M., Lohse, M., and Usadel, B. (2014). Trimmomatic: a flexible trimmer for Illumina sequence data. Bioinformatics 30, 2114–2120.

Buckwalter, C.M., and King, S.J. (2012). Pneumococcal carbohydrate transport: food for thought. Trends Microbiol 20, 517–522.

Craig, A., Mai, J., Cai, S., and Jeyaseelan, S. (2009). Neutrophil recruitment to the lungs during bacterial pneumonia. Infect. Immun. 77, 568–575.

Darling, A.C.E., Mau, B., Blattner, F.R., and Perna, N.T. (2004). Mauve: multiple alignment of conserved genomic sequence with rearrangements. Genome Res. 14, 1394–1403.

David, S.C., Laan, Z., Minhas, V., Chen, A.Y., Davies, J., Hirst, T.R., McColl, S.R., Alsharifi, M., and Paton, J.C. (2019). Enhanced safety and immunogenicity of a pneumococcal surface antigen A mutant whole-cell inactivated pneumococcal vaccine. Immunol. Cell Biol. 97, 726–739.

Dobin, A., Davis, C.A., Schlesinger, F., Drenkow, J., Zaleski, C., Jha, S., Batut, P., Chaisson, M., and Gingeras, T.R. (2013). STAR: ultrafast universal RNA-seq aligner. Bioinformatics 29, 15–21.

Enright, M.C., and Spratt, B.G. (1998). A multilocus sequence typing scheme for *Streptococcus pneumoniae*: identification of clones associated with serious invasive disease. Microbiology 144, 3049–3060.

Faget, J., Boivin, G., Ancey, P.-B., Gkasti, A., Mussard, J., Engblom, C., Pfirschke, C., Vazquez, J., Bendriss-Vermare, N., Caux, C., et al. (2018). Efficient and specific Ly6G+ cell depletion: A change in the current practices toward more relevant functional analyses of neutrophils. BioRxiv 498881.

Kadioglu, A., Weiser, J.N., Paton, J.C., and Andrew, P.W. (2008). The role of *Streptococcus pneumoniae* virulence factors in host respiratory colonization and disease. Nature Reviews Microbiology 6, 288–301.

King, S.J., Hippe, K.R., and Weiser, J.N. (2006). Deglycosylation of human glycoconjugates by the sequential activities of exoglycosidases expressed by *Streptococcus pneumoniae*. Mol. Microbiol. 59, 961–974.

Liao, Y., Smyth, G.K., and Shi, W. (2014). featureCounts: an efficient general purpose program for assigning sequence reads to genomic features. Bioinformatics 30, 923–930.

Lindén, A., Laan, M., and Anderson, G.P. (2005). Neutrophils, interleukin-17A and lung disease. Eur. Resp. J. 25, 159–172.

Livak, K.J., and Schmittgen, T.D. (2001). Analysis of relative gene expression data using real-time quantitative PCR and the 2(-Delta Delta C(T)) Method. Methods 25, 402–408.

Love, M.I., Huber, W., and Anders, S. (2014). Moderated estimation of fold change and dispersion for RNA-seq data with DESeq2. Genome Biology 15, 550.

Mahdi, L.K., Ogunniyi, A.D., LeMessurier, K.S., and Paton, J.C. (2008). Pneumococcal virulence gene expression and host cytokine profiles during pathogenesis of invasive disease. Infect. Immun. 76, 646–657.

McCarthy, M.K., Zhu, L., Procario, M.C., and Weinberg, J.B. (2014). IL-17 contributes to neutrophil recruitment but not to control of viral replication during acute mouse adenovirus type 1 respiratory infection. Virology 456–457, 259–267.

Minhas, V., Harvey, R.M., McAllister, L.J., Seemann, T., Syme, A.E., Baines, S.L., Paton, J.C., and Trappetti, C. (2019). Capacity to utilize raffinose dictates pneumococcal disease phenotype. mBio 10, e02596–18.

Onishi, R.M., and Gaffen, S.L. (2010). Interleukin-17 and its target genes: mechanisms of interleukin-17 function in disease. Immunology 129, 311–321.

Paixao, L., Oliveira, J., Verissimo, A., Vinga, S., Lourenco, E.C., Ventura, M.R., Kjos, M., Veening, J.W., Fernandes, V.E., Andrew, P.W., Yesilkaya, H., and Neves, A.R. (2015). Host glycan sugar-specific pathways in *Streptococcus pneumoniae*: galactose as a key sugar in colonisation and infection. PLoS One 10, e0121042.

Peñaloza, H.F., Nieto, P.A., Muñoz-Durango, N., Salazar-Echegarai, F.J., Torres, J., Parga, M.J., Alvarez-Lobos, M., Riedel, C.A., Kalergis, A.M., and Bueno, S.M. (2015). Interleukin-10 plays a key role in the modulation of neutrophils recruitment and lung inflammation during infection by *Streptococcus pneumoniae*. Immunology 146, 100–112.

Philips, B.J., Meguer, J.X., Redman, J., and Baker, E.H. (2003). Factors determining the appearance of glucose in upper and lower respiratory tract secretions. Intensive Care Med. 29, 2204–2210.

Ritchie, N.D., and Evans, T.J. (2019). Dual RNA-seq in *Streptococcus pneumoniae* infection reveals compartmentalized neutrophil responses in lung and pleural space. MSystems 4, e00216–19.

Ritchie, N.D., Ritchie, R., Bayes, H.K., Mitchell, T.J., and Evans, T.J. (2018). IL-17 can be protective or deleterious in murine pneumococcal pneumonia. PLOS Pathogens 14, e1007099.

Robb, M., Hobbs, J.K., Woodiga, S.A., Shapiro-Ward, S., Suits, M.D., McGregor, N., Brumer, H., Yesilkaya, H., King, S.J., and Boraston, A.B. (2017). Molecular characterization of *N*-glycan degradation and transport in *Streptococcus pneumoniae* and its contribution to virulence. PLoS Pathog 13, e1006090.

Shelburne, S.A., Davenport, M.T., Keith, D.B., and Musser, J.M. (2008). The role of complex carbohydrate catabolism in the pathogenesis of invasive streptococci. Trends Microbiol. 16, 318–325.

Stoppelenburg, A.J., Salimi, V., Hennus, M., Plantinga, M., Veld, R.H. in’t, Walk, J., Meerding, J., Coenjaerts, F., Bont, L., and Boes, M. (2013). Local IL-17A potentiates early neutrophil recruitment to the respiratory tract during severe RSV infection. PLOS ONE 8, e78461.

van Tonder, A.J., Gladstone, R.A., Lo, S.W., Nahm, M.H., du Plessis, M., Cornick, J., Kwambana-Adams, B., Madhi, S.A., Hawkins, P.A., Benisty, R., et al. (2019). Putative novel *cps* loci in a large global collection of pneumococci. Microb Genom 5, doi: 10.1099/mgen.0.000274.

Trappetti, C., Ogunniyi, A.D., Oggioni, M.R., and Paton, J.C. (2011). Extracellular matrix formation enhances the ability of *Streptococcus pneumoniae* to cause invasive sisease. PLoS One 6, e19844.

Trappetti, C., Maten, E. van der, Amin, Z., Potter, A.J., Chen, A.Y., Mourik, P.M. van, Lawrence, A.J., Paton, A.W., and Paton, J.C. (2013). Site of isolation determines biofilm formation and virulence phenotypes of *Streptococcus pneumoniae* serotype 3 clinical isolates. Infect. Immun. 81, 505–513.

Weiser, J.N., Ferreira, D.M., and Paton, J.C. (2018). *Streptococcus pneumoniae*: transmission, colonization and invasion. Nature Reviews Microbiology 16, 355–367.

Westermann, A.J., Barquist, L., and Vogel, J. (2017). Resolving host-pathogen interactions by dual RNA-seq. PLoS Pathog. 13, e1006033.

Wolf, T., Kämmer, P., Brunke, S., and Linde, J. (2018). Two’s company: studying interspecies relationships with dual RNA-seq. Curr Opin Microbiol 42, 7–12.

Yu, Y.-R.A., O’Koren, E.G., Hotten, D.F., Kan, M.J., Kopin, D., Nelson, E.R., Que, L., and Gunn, M.D. (2016). A Protocol for the comprehensive flow cytometric analysis of immune cells in normal and inflamed murine non-lymphoid tissues. PLoS One 11, e0150606.

