## Supplementary Tables S1-S12 for "*In Vivo* Dual RNA-Seq Analysis Reveals the Basis for Differential Tissue Tropism of Clinical Isolates of *Streptococcus pneumoniae*"

**Supplementary Information**

**Supplementary Table 1.** Pneumococcal differential gene expression in the lungs 6 h post-infection, 9-47-Ear vs 9-47M. Genes with fold change (FC) greater than 2 and *p* < 0.05 are shown. FC values highlighted in blue = upregulated in 9-47-Ear, while values highlighted in red = upregulated in 9-47M.

| **Locus tag in 9-47-Ear** | **Product** | **padj** | **FC** |
|---|---|---|---|
| Sp947_chr_00844 | Sialidase B | 3.08E-10 | 313.9807 |
| Sp947_chr_02077 | hypothetical protein | 4.46E-10 | 306.9412 |
| Sp947_chr_00842 | Sodium/glucose cotransporter | 2.22E-09 | 243.4822 |
| Sp947_chr_00841 | N-acetylneuraminate lyase | 4.53E-09 | 227.7963 |
| Sp947_chr_00845 | scyllo-inositol 2-dehydrogenase (NAD(+)) | 4.36E-09 | 221.051 |
| Sp947_chr_00848 | hypothetical protein | 1.19E-08 | 202.7867 |
| Sp947_chr_00853 | V-type sodium ATPase catalytic subunit A | 1.29E-06 | 100.5411 |
| Sp947_chr_00846 | Beta-glucoside kinase | 3.42E-06 | 98.18951 |
| Sp947_chr_00855 | V-type sodium ATPase subunit D | 8.34E-06 | 85.94879 |
| Sp947_chr_00851 | V-type sodium ATPase subunit C | 2.50E-05 | 72.46612 |
| Sp947_chr_00843 | hypothetical protein | 2.17E-05 | 65.97758 |
| Sp947_chr_00839 | HTH-type transcriptional regulator RpiR | 3.09E-05 | 61.28171 |
| Sp947_chr_00854 | V-type sodium ATPase subunit B | 1.32E-06 | 50.86992 |
| Sp947_chr_00120 | hypothetical protein | 3.00E-04 | 45.77325 |
| Sp947_chr_02078 | hypothetical protein | 1.52E-04 | 45.42925 |
| Sp947_chr_00852 | V-type sodium ATPase subunit G | 5.34E-04 | 44.97311 |
| Sp947_chr_01964 | 3-keto-L-gulonate-6-phosphate decarboxylase UlaD | 1.36E-03 | 44.2113 |
| Sp947_chr_00850 | V-type proton ATPase subunit E | 9.39E-04 | 39.74672 |
| Sp947_chr_00840 | Putative N-acetylmannosamine-6-phosphate 2-epimerase | 4.32E-04 | 37.87573 |
| Sp947_chr_01963 | L-ribulose-5-phosphate 3-epimerase UlaE | 2.11E-03 | 37.77052 |
| Sp947_chr_01962 | L-ribulose-5-phosphate 4-epimerase AraD | 2.75E-03 | 35.23993 |
| Sp947_chr_00849 | V-type sodium ATPase subunit K | 2.25E-03 | 34.7923 |
| Sp947_chr_01960 | putative L-ascorbate-6-phosphate lactonase UlaG | 1.16E-03 | 34.0495 |
| Sp947_chr_00847 | hypothetical protein | 3.85E-03 | 30.78829 |
| Sp947_chr_00118 | hypothetical protein | 2.13E-03 | 30.10216 |
| Sp947_chr_00631 | hypothetical protein | 1.22E-04 | 28.36211 |
| Sp947_chr_01965 | Ascorbate-specific PTS system EIIA component | 8.94E-03 | 28.12967 |
| Sp947_chr_00632 | ABC transporter ATP-binding protein YxdL | 7.13E-03 | 26.82578 |
| Sp947_chr_00117 | hypothetical protein | 9.36E-03 | 21.85911 |
| Sp947_chr_00116 | hypothetical protein | 1.27E-02 | 20.39713 |
| Sp947_chr_02096 | hypothetical protein | 2.92E-07 | 18.61494 |
| Sp947_chr_01966 | Ascorbate-specific PTS system EIIB component | 5.76E-02 | 18.33251 |

| | | | |
|---|---|---|---|
| Sp947_chr_01967 | Ascorbate-specific PTS system EIIC component | 1.17E-02 | 14.29112 |
| Sp947_chr_00630 | hypothetical protein | 4.67E-02 | 14.26137 |
| Sp947_chr_01105 | Transcription antiterminator LicT | 5.76E-02 | 12.54562 |
| Sp947_chr_00122 | hypothetical protein | 5.11E-02 | 12.44502 |
| Sp947_chr_00111 | Lactococcin-G-processing and transport ATP-binding protein LagD | 8.11E-02 | 12.07699 |
| Sp947_chr_01629 | PTS system trehalose-specific EIIBC component | 3.15E-18 | 11.47534 |
| Sp947_chr_00637 | hypothetical protein | 8.81E-02 | 11.10599 |
| Sp947_chr_00119 | hypothetical protein | 2.96E-02 | 10.07432 |
| Sp947_chr_01187 | hypothetical protein | 5.24E-02 | 8.833295 |
| Sp947_chr_01975 | Ubiquinone biosynthesis O-methyltransferase | 5.86E-02 | 8.275917 |
| Sp947_chr_00702 | Maltose 6'-phosphate phosphatase | 2.91E-01 | 8.131911 |
| Sp947_chr_00121 | hypothetical protein | 8.44E-02 | 8.011629 |
| Sp947_chr_01982 | ComG operon protein 1 | 1.19E-04 | 7.374729 |
| Sp947_chr_00133 | hypothetical protein | 6.34E-04 | 7.228914 |
| Sp947_chr_02066 | hypothetical protein | 2.74E-01 | 7.180618 |
| Sp947_chr_01949 | hypothetical protein | 2.49E-01 | 7.107773 |
| Sp947_chr_02151 | ComF operon protein 1 | 2.86E-01 | 6.998242 |
| Sp947_chr_01923 | tRNA-Pro(tgg) | 3.04E-01 | 6.979745 |
| Sp947_chr_01980 | hypothetical protein | 3.75E-02 | 6.946905 |
| Sp947_chr_02094 | Ornithine carbamoyltransferase%2C catabolic | 1.29E-13 | 6.850913 |
| Sp947_chr_00240 | Glucitol operon repressor | 2.50E-01 | 6.725672 |
| Sp947_chr_00922 | ComE operon protein 1 | 2.64E-01 | 6.597975 |
| Sp947_chr_01978 | hypothetical protein | 1.39E-01 | 6.480978 |
| Sp947_chr_01277 | hypothetical protein | 1.43E-01 | 6.459624 |
| Sp947_chr_00112 | hypothetical protein | 2.91E-01 | 6.188522 |
| Sp947_chr_02128 | putative glycerol uptake facilitator protein | 4.20E-14 | 6.053371 |
| Sp947_chr_01120 | hypothetical protein | 3.51E-01 | 6.024365 |
| Sp947_chr_01977 | hypothetical protein | 6.78E-02 | 5.999717 |
| Sp947_chr_00794 | tRNA-Gln(ttg) | 3.91E-01 | 5.87385 |
| Sp947_chr_01979 | hypothetical protein | 2.88E-02 | 5.870884 |
| Sp947_chr_02129 | Alpha-glycerophosphate oxidase | 1.19E-37 | 5.867422 |
| Sp947_chr_02097 | Putative dipeptidase | 1.26E-04 | 5.831657 |
| Sp947_chr_00134 | hypothetical protein | 1.28E-02 | 5.621892 |
| Sp947_chr_00659 | Aminopyrimidine aminohydrolase | 3.95E-01 | 5.584997 |
| Sp947_chr_02130 | Glycerol kinase | 1.19E-27 | 5.564175 |
| Sp947_chr_00147 | UDP-glucose 6-dehydrogenase | 6.94E-02 | 5.536174 |
| Sp947_chr_02065 | hypothetical protein | 2.63E-01 | 5.252184 |
| Sp947_chr_01700 | Inner membrane ABC transporter permease protein YcjP | 2.73E-01 | 5.193315 |
| Sp947_chr_02072 | 1-deoxy-D-xylulose-5-phosphate synthase | 2.52E-02 | 5.114776 |
| Sp947_chr_01044 | 3-isopropylmalate dehydratase small subunit | 4.14E-01 | 4.950902 |
| Sp947_chr_02093 | Arginine deiminase | 6.29E-05 | 4.916879 |
| Sp947_chr_00981 | Queuosine precursor transporter QueT | 2.91E-01 | 4.902071 |
| Sp947_chr_00135 | tRNA threonylcarbamoyladenosine biosynthesis protein TsaB | 2.15E-01 | 4.652439 |
| Sp947_chr_02029 | Phosphate-binding protein PstS 2 | 1.20E-06 | 4.568416 |
| Sp947_chr_01119 | hypothetical protein | 3.89E-01 | 4.33514 |

| | | | |
|---|---|---|---|
| Sp947_chr_00105 | hypothetical protein | 2.35E-06 | 4.320641 |
| Sp947_chr_01601 | hypothetical protein | 1.99E-01 | 4.317593 |
| Sp947_chr_01585 | hypothetical protein | 9.40E-06 | 4.244689 |
| Sp947_chr_02095 | Carbamate kinase 1 | 2.88E-02 | 4.210313 |
| Sp947_chr_01582 | Beta-glucoside kinase | 8.39E-11 | 4.021705 |
| Sp947_chr_02074 | Ascorbate-specific PTS system EIIC component | 4.93E-02 | 3.969839 |
| Sp947_chr_00983 | asd | 1.88E-02 | 3.920181 |
| Sp947_chr_02073 | Transketolase | 1.04E-01 | 3.916258 |
| Sp947_chr_00390 | Glycine | 3.47E-01 | 3.883027 |
| Sp947_chr_01611 | hypothetical protein | 4.51E-01 | 3.86601 |
| Sp947_chr_00146 | 2'-N-acetylparomamine deacetylase | 1.44E-01 | 3.836192 |
| Sp947_chr_00145 | hypothetical protein | 2.43E-02 | 3.78253 |
| Sp947_chr_00149 | hypothetical protein | 1.96E-43 | 3.78179 |
| Sp947_chr_01772 | Iron-uptake system permease protein FeuB | 1.10E-02 | 3.780349 |
| Sp947_chr_00109 | Argininosuccinate synthase | 1.38E-01 | 3.732843 |
| Sp947_chr_01631 | HTH-type transcriptional regulator DegA | 2.19E-02 | 3.731615 |
| Sp947_chr_01262 | Putative endo-beta-N-acetylglucosaminidase | 8.15E-02 | 3.596429 |
| Sp947_chr_01107 | PTS system lactose-specific EIICB component | 1.35E-02 | 3.562908 |
| Sp947_chr_01035 | hypothetical protein | 9.08E-02 | 3.496732 |
| Sp947_chr_01250 | putative ABC transporter ATP-binding protein YheS | 6.78E-02 | 3.438094 |
| Sp947_chr_00108 | Arginine-binding extracellular protein ArtP | 1.51E-01 | 3.41617 |
| Sp947_chr_01108 | 6-phospho-beta-galactosidase | 7.85E-02 | 3.360241 |
| Sp947_chr_00649 | ABC transporter ATP-binding protein NatA | 1.01E-01 | 3.333401 |
| Sp947_chr_01835 | Single-stranded DNA-binding protein | 7.89E-05 | 3.281171 |
| Sp947_chr_00055 | hypothetical protein | 2.19E-04 | 3.207721 |
| Sp947_chr_00144 | hypothetical protein | 2.73E-01 | 3.138686 |
| Sp947_chr_00641 | Fluoroquinolones export ATP-binding protein | 5.76E-02 | 3.109696 |
| Sp947_chr_01747 | hypothetical protein | 4.93E-01 | 3.109528 |
| Sp947_chr_01587 | L-arabinose transport system permease protein AraQ | 5.18E-09 | 3.081995 |
| Sp947_chr_01976 | hypothetical protein | 1.08E-01 | 3.078674 |
| Sp947_chr_01155 | Glucose-1-phosphate adenylyltransferase | 1.41E-05 | 3.071928 |
| Sp947_chr_01228 | Tyrosine-protein phosphatase | 3.76E-02 | 3.048501 |
| Sp947_chr_01154 | Glycogen biosynthesis protein GlgD | 3.87E-05 | 2.954565 |
| Sp947_chr_01100 | hypothetical protein | 4.09E-01 | 2.889683 |
| Sp947_chr_01993 | hypothetical protein | 2.76E-03 | 2.87091 |
| Sp947_chr_01263 | Autolysin | 2.73E-01 | 2.870719 |
| Sp947_chr_01613 | hypothetical protein | 3.54E-01 | 2.855586 |
| Sp947_chr_01588 | Lactose transport system permease protein LacF | 8.93E-08 | 2.852978 |
| Sp947_chr_00608 | putative response regulatory protein | 1.10E-09 | 2.84383 |
| Sp947_chr_01600 | Sialidase A | 1.33E-17 | 2.819847 |
| Sp947_chr_00173 | 6%2C7-dimethyl-8-ribityllumazine synthase | 4.42E-01 | 2.814878 |

| | | | |
|---|---|---|---|
| Sp947_chr_00069 | Putative tagatose-6-phosphate ketose/aldose isomerase | 5.19E-08 | 2.76399 |
| Sp947_chr_01584 | hypothetical protein | 1.66E-03 | 2.762851 |
| Sp947_chr_00521 | hypothetical protein | 3.78E-01 | 2.762754 |
| Sp947_chr_00250 | hypothetical protein | 4.70E-01 | 2.727314 |
| Sp947_chr_00721 | hypothetical protein | 3.97E-01 | 2.68465 |
| Sp947_chr_00987 | 2-hydroxymuconate tautomerase | 4.81E-01 | 2.666089 |
| Sp947_chr_01388 | hypothetical protein | 1.16E-01 | 2.622464 |
| Sp947_chr_00923 | ComE operon protein 3 | 2.15E-01 | 2.594794 |
| Sp947_chr_01013 | Serine recombinase PinR | 1.76E-01 | 2.580421 |
| Sp947_chr_01586 | Toxin-antitoxin biofilm protein TabA | 1.04E-02 | 2.560526 |
| Sp947_chr_00023 | competence protein ComW, recombinase RecX | 3.57E-01 | 2.551904 |
| Sp947_chr_01453 | hypothetical protein | 2.91E-01 | 2.548217 |
| Sp947_chr_00480 | hypothetical protein | 2.60E-01 | 2.514127 |
| Sp947_chr_00067 | PTS system mannose-specific EIID component | 4.61E-05 | 2.500185 |
| Sp947_chr_02076 | Mannitol-specific cryptic phosphotransferase enzyme IIA component | 4.56E-01 | 2.444106 |
| Sp947_chr_01589 | Putative ABC transporter substrate-binding protein YesO | 4.02E-17 | 2.425022 |
| Sp947_chr_00052 | Amidophosphoribosyltransferase | 2.35E-06 | 2.408122 |
| Sp947_chr_00457 | hypothetical protein | 3.54E-01 | 2.390357 |
| Sp947_chr_01583 | N-acetylneuraminate lyase | 4.63E-06 | 2.376665 |
| Sp947_chr_00925 | hypothetical protein | 4.93E-02 | 2.375513 |
| Sp947_chr_01278 | 50S ribosomal protein L7/L12 | 2.41E-07 | 2.35971 |
| Sp947_chr_00053 | Phosphoribosylformylglycinamidine cyclo-ligase | 1.21E-05 | 2.353489 |
| Sp947_chr_02127 | hypothetical protein | 4.95E-02 | 2.308295 |
| Sp947_chr_02131 | hypothetical protein | 3.40E-01 | 2.301176 |
| Sp947_chr_01756 | Acyl-coenzyme A thioesterase PaaI | 3.75E-01 | 2.288928 |
| Sp947_chr_01404 | hypothetical protein | 2.91E-01 | 2.276784 |
| Sp947_chr_01775 | putative ABC transporter solute-binding protein YclQ | 1.34E-01 | 2.268465 |
| Sp947_chr_00005 | Peptidyl-tRNA hydrolase | 3.69E-01 | 2.265023 |
| Sp947_chr_01153 | Glycogen synthase | 5.10E-03 | 2.260835 |
| Sp947_chr_01101 | Galactose-6-phosphate isomerase subunit LacA | 2.16E-01 | 2.243408 |
| Sp947_chr_00476 | Protein GrpE | 9.93E-04 | 2.233808 |
| Sp947_chr_00638 | hypothetical protein | 3.54E-01 | 2.218743 |
| Sp947_chr_02032 | Phosphate import ATP-binding protein PstB 3 | 2.74E-02 | 2.215497 |
| Sp947_chr_01224 | hypothetical protein | 3.49E-01 | 2.207203 |
| Sp947_chr_01810 | tRNA-Ser(tga) | 4.13E-01 | 2.19757 |
| Sp947_chr_00762 | Purine nucleoside phosphorylase 1 | 1.20E-01 | 2.193752 |
| Sp947_chr_01803 | HTH-type transcriptional regulator CdhR | 3.03E-01 | 2.193316 |
| Sp947_chr_02007 | tRNA-Gly(tcc) | 4.74E-01 | 2.1921 |
| Sp947_chr_00827 | Type-1 restriction enzyme EcoKI specificity protein | 2.00E-01 | 2.190873 |
| Sp947_chr_01630 | Sucrose-6-phosphate hydrolase | 3.05E-02 | 2.18504 |
| Sp947_chr_00106 | hypothetical protein | 3.27E-03 | 2.177719 |

| | | | |
|---|---|---|---|
| Sp947_chr_00066 | PTS system sorbose-specific EIIC component | 3.08E-03 | 2.174249 |
| Sp947_chr_02030 | Phosphate transport system permease protein PstC 1 | 8.44E-02 | 2.167527 |
| Sp947_chr_00150 | Nisin biosynthesis protein NisB | 7.57E-23 | 2.163768 |
| Sp947_chr_01099 | hypothetical protein | 3.93E-01 | 2.157605 |
| Sp947_chr_01479 | DegV domain-containing protein | 7.47E-03 | 2.15692 |
| Sp947_chr_02062 | hypothetical protein | 2.49E-01 | 2.144907 |
| Sp947_chr_00607 | Peptide methionine sulfoxide reductase MsrA/MsrB | 1.57E-06 | 2.132922 |
| Sp947_chr_00017 | tRNA-Ala(tgc) | 4.70E-01 | 2.123204 |
| Sp947_chr_01824 | tRNA-Ala(tgc) | 4.70E-01 | 2.123204 |
| Sp947_chr_01934 | tRNA-Ala(tgc) | 4.70E-01 | 2.123204 |
| Sp947_chr_02011 | tRNA-Ala(tgc) | 4.70E-01 | 2.123204 |
| Sp947_chr_00140 | hypothetical protein | 4.86E-01 | 2.122109 |
| Sp947_chr_01612 | Molybdenum import ATP-binding protein ModC | 3.11E-01 | 2.120579 |
| Sp947_chr_01773 | Ferric enterobactin transport system permease protein FepD | 2.48E-01 | 2.118436 |
| Sp947_chr_00477 | Chaperone protein DnaK | 3.11E-11 | 2.118333 |
| Sp947_chr_00645 | hypothetical protein | 1.33E-01 | 2.110241 |
| Sp947_chr_00056 | Bifunctional purine biosynthesis protein PurH | 2.62E-06 | 2.095397 |
| Sp947_chr_00496 | Bacteriocin lactacin-F subunit LafA | 4.09E-01 | 2.074411 |
| Sp947_chr_01478 | 4-hydroxy-tetrahydrodipicolinate reductase | 1.97E-02 | 2.068728 |
| Sp947_chr_02033 | Phosphate-specific transport system accessory protein PhoU | 3.11E-02 | 2.04991 |
| Sp947_chr_00174 | Riboflavin biosynthesis protein RibBA | 2.98E-01 | 2.047245 |
| Sp947_chr_01231 | hypothetical protein | 2.17E-01 | 2.042865 |
| Sp947_chr_00152 | hypothetical protein | 1.87E-20 | 2.035498 |
| Sp947_chr_02063 | hypothetical protein | 4.93E-01 | 2.032609 |
| Sp947_chr_00054 | Phosphoribosylglycinamide formyltransferase | 2.39E-02 | 2.031196 |
| Sp947_chr_01614 | hypothetical protein | 3.71E-01 | 2.028597 |
| Sp947_chr_02031 | Phosphate transport system permease protein PstA 2 | 7.01E-02 | 2.026564 |
| Sp947_chr_00475 | Heat-inducible transcription repressor HrcA | 7.89E-05 | 2.006056 |
| Sp947_chr_02088 | Beta-glucoside kinase | 5.00E-01 | 2.004401 |
| Sp947_chr_01317 | HTH-type transcriptional regulatory protein GabR | 3.80E-01 | 2.004158 |
| Sp947_chr_02092 | hypothetical protein | 4.45E-01 | 2.004096 |
| Sp947_chr_00279 | Alcohol dehydrogenase 1 | 4.32E-03 | 2.100505 |
| Sp947_chr_01847 | putative transcriptional regulatory protein | 2.22E-05 | 2.113858 |
| Sp947_chr_01234 | UTP--glucose-1-phosphate uridylyltransferase | 6.67E-04 | 2.154403 |
| Sp947_chr_01238 | Ribitol-5-phosphate cytidylyltransferase | 1.32E-04 | 2.183908 |
| Sp947_chr_00859 | Arginine regulator | 4.52E-03 | 2.209799 |
| Sp947_chr_01055 | Glucose-6-phosphate 1-dehydrogenase | 2.31E-14 | 2.212385 |
| Sp947_chr_01293 | Regulatory protein MsrR | 2.28E-03 | 2.222293 |
| Sp947_chr_01640 | Ribosomal RNA small subunit methyltransferase B | 9.93E-04 | 2.225681 |

| | | | |
|---|---|---|---|
| Sp947_chr_00191 | Folylpolyglutamate synthase | 3.11E-02 | 2.250767 |
| Sp947_chr_01653 | hypothetical protein | 1.43E-02 | 2.295077 |
| Sp947_chr_01300 | 3-dehydroquinate synthase | 2.71E-02 | 2.309334 |
| Sp947_chr_00490 | Lactococcin-G-processing and transport ATP-binding protein LagD | 2.35E-06 | 2.364015 |
| Sp947_chr_01419 | Arginine-binding extracellular protein ArtP | 4.40E-07 | 2.398254 |
| Sp947_chr_01447 | Multidrug resistance ABC transporter ATP-binding/permease protein BmrA | 1.06E-02 | 2.404999 |
| Sp947_chr_00613 | 2-succinyl-6-hydroxy-2%2C4-cyclohexadiene-1-carboxylate synthase | 4.54E-02 | 2.488993 |
| Sp947_chr_02091 | hypothetical protein | 9.36E-03 | 2.491938 |
| Sp947_chr_01850 | hypothetical protein | 4.99E-05 | 2.506163 |
| Sp947_chr_01236 | hypothetical protein | 4.88E-06 | 2.557541 |
| Sp947_chr_01851 | hypothetical protein | 2.43E-03 | 2.891242 |
| Sp947_chr_01237 | Ribulose-5-phosphate reductase | 1.55E-09 | 2.935155 |
| Sp947_chr_01235 | hypothetical protein | 4.67E-09 | 3.067205 |
| Sp947_chr_01848 | Pneumolysin | 7.93E-20 | 3.074683 |
| Sp947_chr_01445 | Multidrug resistance ABC transporter ATP-binding and permease protein | 9.93E-04 | 3.429428 |
| Sp947_chr_01421 | L-cystine transport system permease protein YecS | 4.60E-05 | 3.567119 |
| Sp947_chr_01952 | Lichenan permease IIC component | 3.40E-03 | 3.631755 |
| Sp947_chr_01849 | hypothetical protein | 8.04E-09 | 3.899592 |
| Sp947_chr_01951 | 6-phospho-beta-glucosidase GmuD | 6.09E-04 | 4.38906 |
| Sp947_chr_00675 | Pyruvate oxidase | 2.48E-78 | 5.288241 |
| Sp947_chr_00722 | Biotin transporter BioY2 | 9.36E-03 | 7.135698 |
| Sp947_chr_01592 | scyllo-inositol 2-dehydrogenase (NAD(+)) | 7.65E-10 | 16.61538 |
| Sp947_chr_01596 | Inner membrane ABC transporter permease protein YcjO | 1.70E-07 | 17.3257 |
| Sp947_chr_01598 | Evolved beta-galactosidase subunit beta | 7.36E-05 | 17.8072 |
| Sp947_chr_01595 | Inner membrane ABC transporter permease protein YcjP | 1.12E-06 | 19.42745 |
| Sp947_chr_01599 | hypothetical protein | 2.80E-03 | 23.35696 |
| Sp947_chr_01597 | Putative ABC transporter substrate-binding protein YesO | 1.24E-13 | 40.51841 |

**Supplementary Table 2.** Pneumococcal differential gene expression in the lungs 6 h post-infection, 9-47-Ear vs 4559-Blood. Genes with fold change (FC) greater than 2 and $p < 0.05$ are shown. FC values highlighted in blue = upregulated in 9-47-Ear, while values highlighted in red = upregulated in 4559-Blood.

| **Locus tag in 9-47-Ear** | **Product** | **padj** | **FC** |
|---|---|---|---|
| Sp947_chr_01964 | 3-keto-L-gulonate-6-phosphate decarboxylase UlaD | 9.92E-03 | 24.34795 |

| | | | |
|---|---|---|---|
| Sp947_chr_00305 | Unsaturated chondroitin disaccharide hydrolase | 7.83E-03 | 23.54004 |
| Sp947_chr_01629 | PTS system trehalose-specific EIIBC component | 2.80E-19 | 18.65254 |
| Sp947_chr_01663 | hypothetical protein | 4.24E-02 | 17.40567 |
| Sp947_chr_00108 | Arginine-binding extracellular protein ArtP | 9.63E-03 | 15.89418 |
| Sp947_chr_00309 | hypothetical protein | 4.66E-02 | 15.73089 |
| Sp947_chr_01967 | Ascorbate-specific PTS system EIIC component | 1.09E-02 | 15.09351 |
| Sp947_chr_00109 | Argininosuccinate synthase | 1.64E-02 | 13.49916 |
| Sp947_chr_01318 | Glutamine-binding periplasmic protein | 2.64E-07 | 7.648698 |
| Sp947_chr_01667 | Accessory Sec system protein Asp3 | 4.58E-02 | 7.043576 |
| Sp947_chr_01630 | Sucrose-6-phosphate hydrolase | 1.00E-06 | 6.702188 |
| Sp947_chr_02097 | Putative dipeptidase | 5.39E-05 | 6.495677 |
| Sp947_chr_01672 | General stress protein A | 3.63E-19 | 5.924728 |
| Sp947_chr_01670 | Accessory Sec system protein translocase subunit SecY2 | 3.19E-05 | 5.757463 |
| Sp947_chr_01631 | HTH-type transcriptional regulator DegA | 3.19E-03 | 5.461813 |
| Sp947_chr_00308 | PTS system mannose-specific EIID component | 3.92E-02 | 4.864117 |
| Sp947_chr_00854 | V-type sodium ATPase subunit B | 2.52E-06 | 4.860106 |
| Sp947_chr_01671 | Beta-1%2C6-galactofuranosyltransferase WbbI | 2.36E-05 | 4.837304 |
| Sp947_chr_02096 | hypothetical protein | 2.73E-05 | 4.789669 |
| Sp947_chr_01560 | putative thiol peroxidase | 6.59E-14 | 4.443783 |
| Sp947_chr_01666 | Protein translocase subunit SecA | 2.98E-07 | 4.414981 |
| Sp947_chr_01665 | Glycosyltransferase Gtf1 | 2.51E-05 | 4.413495 |
| Sp947_chr_01920 | putative ABC transporter ATP-binding protein YxlF | 1.07E-07 | 4.401048 |
| Sp947_chr_00855 | V-type sodium ATPase subunit D | 4.26E-03 | 4.394869 |
| Sp947_chr_01081 | L-methionine gamma-lyase | 3.24E-04 | 4.297872 |
| Sp947_chr_01447 | Multidrug resistance ABC transporter ATP-binding/permease protein BmrA | 4.48E-03 | 4.165307 |
| Sp947_chr_01444 | Cystathionine gamma-synthase/O-acetylhomoserine (thiol)-lyase | 6.76E-04 | 4.10799 |
| Sp947_chr_01772 | Iron-uptake system permease protein FeuB | 6.06E-03 | 4.051281 |
| Sp947_chr_01673 | General stress protein A | 1.82E-06 | 4.024977 |
| Sp947_chr_01669 | hypothetical protein | 2.01E-04 | 4.019284 |
| Sp947_chr_01443 | Cystathionine beta-lyase PatB | 6.28E-04 | 4.001289 |
| Sp947_chr_01664 | Glycosyltransferase-stabilizing protein Gtf2 | 6.80E-05 | 3.991404 |
| Sp947_chr_00104 | putative cell wall hydrolase LytN | 6.33E-10 | 3.97022 |
| Sp947_chr_01448 | Oligopeptide-binding protein SarA | 8.69E-35 | 3.885765 |
| Sp947_chr_02094 | Ornithine carbamoyltransferase%2C catabolic | 2.27E-09 | 3.843469 |
| Sp947_chr_02093 | Arginine deiminase | 1.20E-03 | 3.624133 |
| Sp947_chr_01668 | Accessory Sec system protein Asp2 | 9.01E-04 | 3.57589 |
| Sp947_chr_01993 | hypothetical protein | 2.75E-04 | 3.5502 |
| Sp947_chr_00262 | Alkanal monooxygenase alpha chain | 4.20E-02 | 3.513196 |
| Sp947_chr_00631 | hypothetical protein | 6.86E-03 | 3.468201 |
| Sp947_chr_01628 | Putative fructokinase | 3.76E-02 | 3.396119 |
| Sp947_chr_01674 | putative glycosyltransferase EpsJ | 3.86E-03 | 3.263426 |
| Sp947_chr_02070 | hypothetical protein | 3.40E-02 | 3.249364 |
| Sp947_chr_00169 | hypothetical protein | 1.81E-12 | 3.175252 |

| | | | |
|---|---|---|---|
| Sp947_chr_00105 | hypothetical protein | 9.82E-05 | 3.113409 |
| Sp947_chr_00544 | 5%2C10-methylenetetrahydrofolate reductase | 6.48E-07 | 3.063051 |
| Sp947_chr_02127 | hypothetical protein | 5.20E-03 | 3.051198 |
| Sp947_chr_01919 | hypothetical protein | 6.43E-04 | 3.008033 |
| Sp947_chr_00148 | hypothetical protein | 3.11E-03 | 2.944036 |
| Sp947_chr_00846 | Beta-glucoside kinase | 2.93E-02 | 2.853877 |
| Sp947_chr_01724 | Tryptophan synthase beta chain | 8.91E-04 | 2.848494 |
| Sp947_chr_01917 | Transcriptional regulatory protein DesR | 1.30E-04 | 2.799649 |
| Sp947_chr_01157 | NADP-dependent glyceraldehyde-3-phosphate dehydrogenase | 5.07E-05 | 2.764121 |
| Sp947_chr_00161 | Methionine import ATP-binding protein MetN | 6.86E-03 | 2.745542 |
| Sp947_chr_01732 | T-box | 4.38E-02 | 2.72911 |
| Sp947_chr_01840 | hypothetical protein | 2.90E-03 | 2.691428 |
| Sp947_chr_00425 | Acetolactate synthase large subunit | 1.12E-15 | 2.650045 |
| Sp947_chr_00848 | hypothetical protein | 5.95E-04 | 2.637928 |
| Sp947_chr_00845 | scyllo-inositol 2-dehydrogenase (NAD(+)) | 4.49E-07 | 2.594575 |
| Sp947_chr_01918 | Sensor histidine kinase DesK | 1.20E-05 | 2.551678 |
| Sp947_chr_00844 | Sialidase B | 1.14E-08 | 2.530319 |
| Sp947_chr_00692 | Leucine-%2C isoleucine-%2C valine-%2C threonine-%2C and alanine-binding protein | 2.08E-05 | 2.499057 |
| Sp947_chr_02160 | Secreted 45 kDa protein | 1.55E-13 | 2.495737 |
| Sp947_chr_00476 | Protein GrpE | 2.01E-04 | 2.420533 |
| Sp947_chr_00787 | Branched-chain-amino-acid aminotransferase | 1.35E-07 | 2.395717 |
| Sp947_chr_01424 | 6-phosphogluconolactonase | 5.77E-04 | 2.345615 |
| Sp947_chr_01660 | Serine/threonine transporter SstT | 2.89E-04 | 2.303845 |
| Sp947_chr_00232 | 2%2C3-bisphosphoglycerate-dependent phosphoglycerate mutase | 3.88E-02 | 2.243558 |
| Sp947_chr_00842 | Sodium/glucose cotransporter | 2.08E-05 | 2.238557 |
| Sp947_chr_01276 | NADP-specific glutamate dehydrogenase | 9.86E-19 | 2.229232 |
| Sp947_chr_01391 | Pyridoxal 5'-phosphate synthase subunit PdxS | 1.61E-04 | 2.172842 |
| Sp947_chr_00696 | High-affinity branched-chain amino acid transport ATP-binding protein LivF | 4.48E-03 | 2.171266 |
| Sp947_chr_01661 | hypothetical protein | 2.04E-02 | 2.153858 |
| Sp947_chr_00853 | V-type sodium ATPase catalytic subunit A | 9.55E-03 | 2.130379 |
| Sp947_chr_00543 | 5-methyltetrahydropteroyltriglutamate--homocysteine methyltransferase | 2.38E-05 | 2.118701 |
| Sp947_chr_00695 | Lipopolysaccharide export system ATP-binding protein LptB | 2.08E-02 | 2.098421 |
| Sp947_chr_00153 | hypothetical protein | 2.00E-19 | 2.013755 |
| Sp947_chr_01394 | NADH-dependent flavin reductase subunit 2 | 3.48E-02 | 2.043274 |
| Sp947_chr_00205 | 30S ribosomal protein S19 | 2.68E-08 | 2.057637 |
| Sp947_chr_00489 | Lactococcin A secretion protein LcnD | 5.57E-07 | 2.137788 |
| Sp947_chr_00057 | Phosphoribosylamine--glycine ligase | 2.04E-02 | 2.148252 |
| Sp947_chr_00155 | hypothetical protein | 1.27E-26 | 2.163473 |
| Sp947_chr_00151 | Nisin biosynthesis protein NisC | 3.70E-31 | 2.171804 |
| Sp947_chr_00498 | hypothetical protein | 3.74E-02 | 2.222467 |
| Sp947_chr_02091 | hypothetical protein | 2.88E-02 | 2.230061 |
| Sp947_chr_01955 | Aldehyde-alcohol dehydrogenase | 3.36E-10 | 2.239625 |
| Sp947_chr_00490 | Lactococcin-G-processing and transport ATP-binding protein LagD | 1.15E-05 | 2.2462 |

| | | | |
|---|---|---|---|
| Sp947_chr_02054 | Maltose transport system permease protein MalG | 2.55E-03 | 2.250996 |
| Sp947_chr_00152 | hypothetical protein | 3.16E-35 | 2.253763 |
| Sp947_chr_02053 | Maltose transport system permease protein MalF | 6.34E-05 | 2.292191 |
| Sp947_chr_00157 | L-cystine-binding protein TcyA | 2.04E-02 | 2.305243 |
| Sp947_chr_00491 | hypothetical protein | 2.93E-08 | 2.319449 |
| Sp947_chr_00150 | Nisin biosynthesis protein NisB | 2.65E-31 | 2.324591 |
| Sp947_chr_00499 | hypothetical protein | 2.99E-04 | 2.378979 |
| Sp947_chr_00154 | hypothetical protein | 4.99E-25 | 2.443422 |
| Sp947_chr_00281 | Guanine/hypoxanthine permease PbuO | 5.57E-07 | 2.494449 |
| Sp947_chr_00041 | hypothetical protein | 3.18E-02 | 2.52396 |
| Sp947_chr_01178 | NAD kinase | 5.94E-03 | 2.564431 |
| Sp947_chr_00050 | Phosphoribosylaminoimidazole-succinocarboxamide synthase | 1.84E-02 | 2.58736 |
| Sp947_chr_00263 | Pullulanase | 6.02E-06 | 2.823468 |
| Sp947_chr_00440 | Formate acetyltransferase | 2.27E-52 | 2.85616 |
| Sp947_chr_00279 | Alcohol dehydrogenase 1 | 5.38E-07 | 3.088478 |
| Sp947_chr_01753 | Uric acid permease PucK | 2.63E-06 | 3.150544 |
| Sp947_chr_00156 | hypothetical protein | 1.25E-03 | 3.349079 |
| Sp947_chr_01798 | Sucrose phosphorylase | 3.67E-02 | 3.943655 |
| Sp947_chr_00282 | hypothetical protein | 5.77E-04 | 5.106822 |
| Sp947_chr_01802 | Bifunctional alpha-galactosidase/sucrose kinase AgaSK | 2.44E-03 | 5.116292 |
| Sp947_chr_00676 | hypothetical protein | 1.18E-02 | 6.452692 |
| Sp947_chr_01801 | Multiple sugar-binding protein | 4.56E-03 | 6.788077 |
| Sp947_chr_00675 | Pyruvate oxidase | 7.84E-110 | 6.915359 |
| Sp947_chr_01592 | scyllo-inositol 2-dehydrogenase (NAD(+)) | 8.87E-15 | 29.54277 |
| Sp947_chr_01596 | Inner membrane ABC transporter permease protein YcjO | 2.70E-13 | 42.27417 |
| Sp947_chr_01598 | Evolved beta-galactosidase subunit beta | 2.11E-08 | 45.04628 |
| Sp947_chr_01595 | Inner membrane ABC transporter permease protein YcjP | 9.79E-12 | 49.58462 |
| Sp947_chr_01599 | hypothetical protein | 3.26E-06 | 80.56657 |
| Sp947_chr_01597 | Putative ABC transporter substrate-binding protein YesO | 1.55E-21 | 99.69291 |

**Supplementary Table 3.** Pneumococcal differential gene expression in the lungs 6 h post-infection, 9-47-Ear vs 4559M. Genes with fold change (FC) greater than 2 and $p < 0.05$ are shown. FC values highlighted in blue = upregulated in 9-47-Ear, while values highlighted in red = upregulated in 4559M.

| **Locus tag in 9-47-Ear** | **Product** | **padj** | **FC** |
|---|---|---|---|
| Sp947_chr_01081 | L-methionine gamma-lyase | 2.01E-06 | 6.049978 |
| Sp947_chr_01629 | PTS system trehalose-specific EIIBC component | 4.32E-17 | 5.71303 |
| Sp947_chr_01673 | General stress protein A | 2.68E-09 | 4.856349 |
| Sp947_chr_01318 | Glutamine-binding periplasmic protein | 3.42E-07 | 4.661576 |
| Sp947_chr_01448 | Oligopeptide-binding protein SarA | 1.19E-45 | 4.251728 |

| | | | |
|---|---|---|---|
| Sp947_chr_01671 | Beta-1%2C6-galactofuranosyltransferase WbbI | 5.13E-06 | 4.200336 |
| Sp947_chr_02096 | hypothetical protein | 1.64E-05 | 4.131014 |
| Sp947_chr_02097 | Putative dipeptidase | 2.74E-04 | 3.935156 |
| Sp947_chr_00544 | 5%2C10-methylenetetrahydrofolate reductase | 4.34E-11 | 3.926284 |
| Sp947_chr_01630 | Sucrose-6-phosphate hydrolase | 1.32E-05 | 3.842564 |
| Sp947_chr_00105 | hypothetical protein | 9.16E-07 | 3.720141 |
| Sp947_chr_01674 | putative glycosyltransferase EpsJ | 2.97E-04 | 3.645955 |
| Sp947_chr_02093 | Arginine deiminase | 6.31E-04 | 3.638204 |
| Sp947_chr_02094 | Ornithine carbamoyltransferase%2C catabolic | 2.11E-10 | 3.595965 |
| Sp947_chr_01669 | hypothetical protein | 9.12E-05 | 3.520343 |
| Sp947_chr_00145 | hypothetical protein | 1.10E-02 | 3.437721 |
| Sp947_chr_01672 | General stress protein A | 3.78E-14 | 3.337295 |
| Sp947_chr_01444 | Cystathionine gamma-synthase/O-acetylhomoserine (thiol)-lyase | 6.68E-04 | 3.323343 |
| Sp947_chr_01628 | Putative fructokinase | 1.81E-02 | 3.312402 |
| Sp947_chr_01993 | hypothetical protein | 1.09E-04 | 3.292474 |
| Sp947_chr_00104 | putative cell wall hydrolase LytN | 9.04E-10 | 3.274894 |
| Sp947_chr_01668 | Accessory Sec system protein Asp2 | 3.58E-04 | 3.271471 |
| Sp947_chr_02070 | hypothetical protein | 1.33E-02 | 3.268113 |
| Sp947_chr_01631 | HTH-type transcriptional regulator DegA | 1.62E-02 | 3.241793 |
| Sp947_chr_01666 | Protein translocase subunit SecA | 1.86E-06 | 3.194154 |
| Sp947_chr_01560 | putative thiol peroxidase | 1.87E-11 | 3.156039 |
| Sp947_chr_00148 | hypothetical protein | 3.69E-04 | 3.143065 |
| Sp947_chr_00843 | hypothetical protein | 6.34E-04 | 3.140506 |
| Sp947_chr_01446 | Multidrug resistance ABC transporter ATP-binding and permease protein | 2.44E-02 | 3.095084 |
| Sp947_chr_00631 | hypothetical protein | 5.70E-03 | 3.078157 |
| Sp947_chr_01670 | Accessory Sec system protein translocase subunit SecY2 | 6.16E-04 | 3.060934 |
| Sp947_chr_01447 | Multidrug resistance ABC transporter ATP-binding/permease protein BmrA | 6.68E-03 | 3.030447 |
| Sp947_chr_01665 | Glycosyltransferase Gtf1 | 2.91E-04 | 2.877465 |
| Sp947_chr_00846 | Beta-glucoside kinase | 1.36E-02 | 2.854048 |
| Sp947_chr_01664 | Glycosyltransferase-stabilizing protein Gtf2 | 4.96E-04 | 2.799165 |
| Sp947_chr_01660 | Serine/threonine transporter SstT | 1.28E-06 | 2.757889 |
| Sp947_chr_00855 | V-type sodium ATPase subunit D | 4.10E-02 | 2.573402 |
| Sp947_chr_01772 | Iron-uptake system permease protein FeuB | 3.65E-02 | 2.566198 |
| Sp947_chr_01443 | Cystathionine beta-lyase PatB | 1.32E-02 | 2.466567 |
| Sp947_chr_00695 | Lipopolysaccharide export system ATP-binding protein LptB | 1.70E-03 | 2.4024 |
| Sp947_chr_00854 | V-type sodium ATPase subunit B | 2.76E-03 | 2.359804 |
| Sp947_chr_01157 | NADP-dependent glyceraldehyde-3-phosphate dehydrogenase | 1.59E-04 | 2.339795 |
| Sp947_chr_00169 | hypothetical protein | 4.71E-09 | 2.339571 |
| Sp947_chr_00692 | Leucine-%2C isoleucine-%2C valine-%2C threonine-%2C and alanine-binding protein | 2.14E-05 | 2.312731 |
| Sp947_chr_00543 | 5-methyltetrahydropteroyltriglutamate--homocysteine methyltransferase | 2.80E-07 | 2.29823 |
| Sp947_chr_01661 | hypothetical protein | 5.48E-03 | 2.245706 |
| Sp947_chr_00701 | PTS system glucose-specific EIICBA component | 2.24E-04 | 2.216955 |
| Sp947_chr_01985 | N-acetylglucosamine-6-phosphate deacetylase | 1.28E-06 | 2.199836 |

| | | | |
|---|---|---|---|
| Sp947_chr_00845 | scyllo-inositol 2-dehydrogenase (NAD(+)) | 6.36E-06 | 2.193442 |
| Sp947_chr_00967 | putative amino acid permease YhdG | 6.47E-03 | 2.175611 |
| Sp947_chr_00853 | V-type sodium ATPase catalytic subunit A | 2.93E-03 | 2.171867 |
| Sp947_chr_00747 | hypothetical protein | 1.16E-02 | 2.164988 |
| Sp947_chr_01984 | Alcohol dehydrogenase | 6.59E-05 | 2.143827 |
| Sp947_chr_02160 | Secreted 45 kDa protein | 5.54E-11 | 2.11217 |
| Sp947_chr_00696 | High-affinity branched-chain amino acid transport ATP-binding protein LivF | 2.42E-03 | 2.088927 |
| Sp947_chr_00844 | Sialidase B | 1.59E-06 | 2.076986 |
| Sp947_chr_01840 | hypothetical protein | 1.72E-02 | 2.063451 |
| Sp947_chr_00842 | Sodium/glucose cotransporter | 4.98E-05 | 2.058073 |
| Sp947_chr_00787 | Branched-chain-amino-acid aminotransferase | 2.56E-06 | 2.038246 |
| Sp947_chr_02071 | Dihydroxy-acid dehydratase | 1.13E-06 | 2.036651 |
| Sp947_chr_00841 | N-acetylneuraminate lyase | 2.76E-03 | 2.028736 |
| Sp947_chr_00219 | 50S ribosomal protein L30 | 4.02E-06 | 2.006281 |
| Sp947_chr_01505 | putative MFS-type transporter YhjX | 1.81E-02 | 2.023671 |
| Sp947_chr_00337 | Putative O-antigen transporter | 1.76E-08 | 2.108266 |
| Sp947_chr_00285 | GTP cyclohydrolase 1 | 7.38E-03 | 2.146718 |
| Sp947_chr_01309 | Spermidine/putrescine-binding periplasmic protein | 9.70E-03 | 2.183815 |
| Sp947_chr_00051 | Phosphoribosylformylglycinamidine synthase | 2.50E-18 | 2.248477 |
| Sp947_chr_00499 | hypothetical protein | 3.77E-04 | 2.270129 |
| Sp947_chr_00489 | Lactococcin A secretion protein LcnD | 4.44E-09 | 2.301998 |
| Sp947_chr_00057 | Phosphoribosylamine--glycine ligase | 3.82E-03 | 2.336422 |
| Sp947_chr_01178 | NAD kinase | 7.88E-03 | 2.39753 |
| Sp947_chr_00440 | Formate acetyltransferase | 4.09E-39 | 2.453134 |
| Sp947_chr_00491 | hypothetical protein | 2.44E-11 | 2.578315 |
| Sp947_chr_01048 | GMP reductase | 1.47E-04 | 2.657956 |
| Sp947_chr_00060 | hypothetical protein | 2.11E-02 | 2.684809 |
| Sp947_chr_00490 | Lactococcin-G-processing and transport ATP-binding protein LagD | 8.32E-10 | 2.783617 |
| Sp947_chr_00281 | Guanine/hypoxanthine permease PbuO | 1.01E-11 | 3.141839 |
| Sp947_chr_00050 | Phosphoribosylaminoimidazole-succinocarboxamide synthase | 8.38E-04 | 3.197571 |
| Sp947_chr_01752 | Xanthine phosphoribosyltransferase | 5.58E-03 | 3.533145 |
| Sp947_chr_01753 | Uric acid permease PucK | 1.84E-11 | 4.438869 |
| Sp947_chr_00282 | hypothetical protein | 5.34E-05 | 5.973551 |
| Sp947_chr_00675 | Pyruvate oxidase | 2.07E-119 | 7.278273 |
| Sp947_chr_00676 | hypothetical protein | 3.25E-03 | 7.516116 |
| Sp947_chr_01592 | scyllo-inositol 2-dehydrogenase (NAD(+)) | 1.85E-18 | 43.48026 |
| Sp947_chr_01598 | Evolved beta-galactosidase subunit beta | 4.44E-09 | 49.62974 |
| Sp947_chr_01596 | Inner membrane ABC transporter permease protein YcjO | 1.62E-14 | 50.00969 |
| Sp947_chr_01599 | hypothetical protein | 1.54E-05 | 59.75261 |
| Sp947_chr_01595 | Inner membrane ABC transporter permease protein YcjP | 4.01E-13 | 60.7687 |
| Sp947_chr_01597 | Putative ABC transporter substrate-binding protein YesO | 7.96E-23 | 115.2393 |

**Supplementary Table 4.** Pneumococcal differential gene expression in the lungs 6 h post-infection, 9-47M vs 4559-Blood. Genes with fold change (FC) greater than 2 and $p$ < 0.05 are shown. FC values highlighted in blue = upregulated in 9-47M, while values highlighted in red = upregulated in 4559-Blood.

| Locus tag in 9-47-Ear | Product | padj | FC |
|---|---|---|---|
| Sp947_chr_00305 | Unsaturated chondroitin disaccharide hydrolase | 5.14E-05 | 68.93754 |
| Sp947_chr_00306 | PTS system sorbose-specific EIIB component | 3.51E-04 | 49.45218 |
| Sp947_chr_00300 | KHG/KDPG aldolase | 3.91E-03 | 24.0683 |
| Sp947_chr_01663 | hypothetical protein | 4.89E-02 | 14.9774 |
| Sp947_chr_00307 | N-acetylgalactosamine permease IIC component 1 | 1.38E-02 | 13.15252 |
| Sp947_chr_01954 | PTS system oligo-beta-mannoside-specific EIIA component | 1.43E-02 | 12.11948 |
| Sp947_chr_01722 | hypothetical protein | 4.94E-02 | 11.94383 |
| Sp947_chr_00301 | 2-dehydro-3-deoxygluconokinase | 6.47E-04 | 11.29777 |
| Sp947_chr_01447 | Multidrug resistance ABC transporter ATP-binding/permease protein BmrA | 7.75E-08 | 10.01756 |
| Sp947_chr_00308 | PTS system mannose-specific EIID component | 2.03E-03 | 7.771254 |
| Sp947_chr_00262 | Alkanal monooxygenase alpha chain | 1.33E-04 | 6.912513 |
| Sp947_chr_00193 | Cardiolipin synthase | 4.62E-03 | 6.683222 |
| Sp947_chr_01672 | General stress protein A | 1.61E-20 | 6.353083 |
| Sp947_chr_01448 | Oligopeptide-binding protein SarA | 2.32E-67 | 6.342907 |
| Sp947_chr_01318 | Glutamine-binding periplasmic protein | 8.47E-06 | 6.056553 |
| Sp947_chr_01670 | Accessory Sec system protein translocase subunit SecY2 | 3.95E-05 | 5.615085 |
| Sp947_chr_01951 | 6-phospho-beta-glucosidase GmuD | 2.41E-04 | 5.149937 |
| Sp947_chr_01671 | Beta-1%2C6-galactofuranosyltransferase WbbI | 2.30E-05 | 4.811812 |
| Sp947_chr_01674 | putative glycosyltransferase EpsJ | 2.18E-05 | 4.795206 |
| Sp947_chr_01669 | hypothetical protein | 1.60E-05 | 4.73273 |
| Sp947_chr_01444 | Cystathionine gamma-synthase/O-acetylhomoserine (thiol)-lyase | 1.22E-04 | 4.673266 |
| Sp947_chr_01665 | Glycosyltransferase Gtf1 | 1.18E-05 | 4.600689 |
| Sp947_chr_01849 | hypothetical protein | 1.09E-09 | 4.492743 |
| Sp947_chr_01666 | Protein translocase subunit SecA | 2.78E-07 | 4.406684 |
| Sp947_chr_01952 | Lichenan permease IIC component | 9.92E-04 | 4.298464 |
| Sp947_chr_01668 | Accessory Sec system protein Asp2 | 1.11E-04 | 4.155341 |
| Sp947_chr_01673 | General stress protein A | 1.46E-06 | 4.054469 |
| Sp947_chr_01445 | Multidrug resistance ABC transporter ATP-binding and permease protein | 2.20E-04 | 4.038759 |
| Sp947_chr_00104 | putative cell wall hydrolase LytN | 6.98E-10 | 3.983611 |
| Sp947_chr_01664 | Glycosyltransferase-stabilizing protein Gtf2 | 8.77E-05 | 3.910847 |
| Sp947_chr_01157 | NADP-dependent glyceraldehyde-3-phosphate dehydrogenase | 1.24E-08 | 3.804836 |
| Sp947_chr_01920 | putative ABC transporter ATP-binding protein YxlF | 3.68E-06 | 3.759168 |
| Sp947_chr_01995 | putative FMN/FAD exporter YeeO | 4.45E-02 | 3.737599 |
| Sp947_chr_01446 | Multidrug resistance ABC transporter ATP-binding and permease protein | 2.66E-03 | 3.726174 |
| Sp947_chr_01443 | Cystathionine beta-lyase PatB | 2.07E-03 | 3.550934 |

| | | | |
|---|---|---|---|
| Sp947_chr_00544 | 5%2C10-methylenetetrahydrofolate reductase | 7.79E-09 | 3.549207 |
| Sp947_chr_00161 | Methionine import ATP-binding protein MetN | 2.08E-04 | 3.538504 |
| Sp947_chr_01837 | Phenylalanine--tRNA ligase beta subunit | 3.15E-02 | 3.43443 |
| Sp947_chr_01848 | Pneumolysin | 9.47E-22 | 3.364311 |
| Sp947_chr_01420 | Glutamine transport ATP-binding protein GlnQ | 7.35E-08 | 3.349461 |
| Sp947_chr_00162 | Methionine import system permease protein MetP | 3.23E-03 | 3.322192 |
| Sp947_chr_01081 | L-methionine gamma-lyase | 8.60E-03 | 3.120338 |
| Sp947_chr_01260 | Putative phosphatase | 3.26E-02 | 3.115464 |
| Sp947_chr_01630 | Sucrose-6-phosphate hydrolase | 1.34E-02 | 3.067307 |
| Sp947_chr_02070 | hypothetical protein | 4.65E-02 | 2.98896 |
| Sp947_chr_00181 | putative murein peptide carboxypeptidase | 4.94E-02 | 2.972231 |
| Sp947_chr_01419 | Arginine-binding extracellular protein ArtP | 1.35E-09 | 2.930964 |
| Sp947_chr_00482 | ABC-type transporter ATP-binding protein EcsA | 2.93E-02 | 2.922061 |
| Sp947_chr_01055 | Glucose-6-phosphate 1-dehydrogenase | 9.29E-23 | 2.874562 |
| Sp947_chr_01918 | Sensor histidine kinase DesK | 6.21E-07 | 2.828767 |
| Sp947_chr_01851 | hypothetical protein | 2.91E-03 | 2.786765 |
| Sp947_chr_01339 | Glucosamine-6-phosphate deaminase | 9.28E-07 | 2.746131 |
| Sp947_chr_01850 | hypothetical protein | 1.10E-05 | 2.743522 |
| Sp947_chr_01732 | T-box | 3.58E-02 | 2.720698 |
| Sp947_chr_00543 | 5-methyltetrahydropteroyltriglutamate--homocysteine methyltransferase | 2.51E-09 | 2.709082 |
| Sp947_chr_01847 | putative transcriptional regulatory protein | 6.58E-08 | 2.646376 |
| Sp947_chr_01238 | Ribitol-5-phosphate cytidylyltransferase | 4.11E-06 | 2.589054 |
| Sp947_chr_01056 | Arginine transport ATP-binding protein ArtM | 2.77E-13 | 2.557431 |
| Sp947_chr_01421 | L-cystine transport system permease protein YecS | 2.65E-03 | 2.512296 |
| Sp947_chr_01282 | putative ABC transporter ATP-binding protein | 2.62E-03 | 2.508221 |
| Sp947_chr_01642 | Primosomal protein N' | 3.98E-04 | 2.500435 |
| Sp947_chr_00696 | High-affinity branched-chain amino acid transport ATP-binding protein LivF | 3.30E-04 | 2.495921 |
| Sp947_chr_01919 | hypothetical protein | 7.59E-03 | 2.477463 |
| Sp947_chr_01917 | Transcriptional regulatory protein DesR | 1.19E-03 | 2.460707 |
| Sp947_chr_01724 | Tryptophan synthase beta chain | 6.73E-03 | 2.43327 |
| Sp947_chr_00434 | hypothetical protein | 1.20E-02 | 2.423222 |
| Sp947_chr_00191 | Folylpolyglutamate synthase | 1.37E-02 | 2.406573 |
| Sp947_chr_00787 | Branched-chain-amino-acid aminotransferase | 1.28E-07 | 2.396524 |
| Sp947_chr_01660 | Serine/threonine transporter SstT | 1.56E-04 | 2.365193 |
| Sp947_chr_00817 | PTS system fructose-specific EIIABC component | 3.52E-05 | 2.357843 |
| Sp947_chr_02040 | N-acetyldiaminopimelate deacetylase | 2.44E-03 | 2.31786 |
| Sp947_chr_00425 | Acetolactate synthase large subunit | 9.17E-11 | 2.279343 |
| Sp947_chr_00169 | hypothetical protein | 4.14E-06 | 2.275144 |
| Sp947_chr_00944 | putative S-adenosyl-L-methionine-dependent methyltransferase TehB | 6.05E-03 | 2.260243 |
| Sp947_chr_01560 | putative thiol peroxidase | 4.54E-04 | 2.24987 |
| Sp947_chr_01215 | putative peptidoglycan glycosyltransferase FtsW | 4.18E-05 | 2.245616 |
| Sp947_chr_02018 | putative ABC transporter ATP-binding protein | 2.54E-04 | 2.232901 |
| Sp947_chr_00985 | 4-hydroxy-tetrahydrodipicolinate synthase | 9.09E-04 | 2.211691 |
| Sp947_chr_01283 | putative ABC transporter ATP-binding protein | 9.29E-03 | 2.209084 |

| | | | |
|---|---|---|---|
| Sp947_chr_02056 | HTH-type transcriptional regulator MalR | 2.43E-02 | 2.208454 |
| Sp947_chr_00859 | Arginine regulator | 4.43E-03 | 2.173132 |
| Sp947_chr_01470 | hypothetical protein | 1.28E-07 | 2.172156 |
| Sp947_chr_00967 | putative amino acid permease YhdG | 8.74E-03 | 2.141793 |
| Sp947_chr_00695 | Lipopolysaccharide export system ATP-binding protein LptB | 1.97E-02 | 2.066565 |
| Sp947_chr_01653 | hypothetical protein | 2.93E-02 | 2.065838 |
| Sp947_chr_01057 | Glutamine transport system permease protein GlnP | 2.03E-12 | 2.062275 |
| Sp947_chr_01661 | hypothetical protein | 2.78E-02 | 2.054099 |
| Sp947_chr_01640 | Ribosomal RNA small subunit methyltransferase B | 2.91E-03 | 2.053249 |
| Sp947_chr_01281 | 5-methylthioadenosine/S-adenosylhomocysteine deaminase | 7.78E-03 | 2.046941 |
| Sp947_chr_00789 | hypothetical protein | 6.20E-04 | 2.033781 |
| Sp947_chr_01235 | hypothetical protein | 1.92E-04 | 2.031984 |
| Sp947_chr_00258 | Putative zinc metalloprotease | 6.05E-03 | 2.019156 |
| Sp947_chr_02160 | Secreted 45 kDa protein | 1.02E-07 | 2.017272 |
| Sp947_chr_00790 | hypothetical protein | 8.94E-06 | 2.016723 |
| Sp947_chr_01391 | Pyridoxal 5'-phosphate synthase subunit PdxS | 9.79E-04 | 2.001587 |
| Sp947_chr_00210 | 30S ribosomal protein S17 | 4.67E-07 | 2.011481 |
| Sp947_chr_01154 | Glycogen biosynthesis protein GlgD | 1.55E-02 | 2.038321 |
| Sp947_chr_01279 | 50S ribosomal protein L10 | 1.09E-07 | 2.043546 |
| Sp947_chr_01477 | CCA-adding enzyme | 4.41E-02 | 2.051156 |
| Sp947_chr_01432 | ATP synthase subunit c | 6.93E-03 | 2.052648 |
| Sp947_chr_00225 | 30S ribosomal protein S13 | 1.54E-08 | 2.076816 |
| Sp947_chr_00498 | hypothetical protein | 4.94E-02 | 2.116476 |
| Sp947_chr_00928 | 50S ribosomal protein L35 | 2.34E-03 | 2.124092 |
| Sp947_chr_00608 | putative response regulatory protein | 3.31E-05 | 2.136323 |
| Sp947_chr_00228 | 50S ribosomal protein L17 | 3.58E-08 | 2.137006 |
| Sp947_chr_01478 | 4-hydroxy-tetrahydrodipicolinate reductase | 6.93E-03 | 2.173119 |
| Sp947_chr_00501 | hypothetical protein | 5.79E-03 | 2.174678 |
| Sp947_chr_01479 | DegV domain-containing protein | 3.33E-03 | 2.221312 |
| Sp947_chr_01591 | Putative N-acetylmannosamine-6-phosphate 2-epimerase | 7.35E-08 | 2.23104 |
| Sp947_chr_00059 | N5-carboxyaminoimidazole ribonucleotide synthase | 6.69E-04 | 2.236367 |
| Sp947_chr_01589 | Putative ABC transporter substrate-binding protein YesO | 9.43E-15 | 2.28675 |
| Sp947_chr_01584 | hypothetical protein | 1.07E-02 | 2.333389 |
| Sp947_chr_01590 | PTS system glucose-specific EIICBA component | 5.93E-17 | 2.346142 |
| Sp947_chr_00057 | Phosphoribosylamine--glycine ligase | 6.10E-03 | 2.431764 |
| Sp947_chr_01596 | Inner membrane ABC transporter permease protein YcjO | 4.53E-05 | 2.439969 |
| Sp947_chr_01597 | Putative ABC transporter substrate-binding protein YesO | 1.57E-11 | 2.460435 |
| Sp947_chr_00577 | Branched-chain amino acid transport system 2 carrier protein | 2.12E-06 | 2.461054 |
| Sp947_chr_00499 | hypothetical protein | 2.11E-04 | 2.4774 |
| Sp947_chr_01600 | Sialidase A | 1.40E-13 | 2.501157 |
| Sp947_chr_01598 | Evolved beta-galactosidase subunit beta | 1.08E-03 | 2.529667 |
| Sp947_chr_00925 | hypothetical protein | 1.77E-02 | 2.540488 |

| | | | |
|---|---|---|---|
| Sp947_chr_01595 | Inner membrane ABC transporter permease protein YcjP | 4.13E-05 | 2.552297 |
| Sp947_chr_00282 | hypothetical protein | 3.15E-02 | 2.616579 |
| Sp947_chr_01586 | Toxin-antitoxin biofilm protein TabA | 2.91E-03 | 2.76266 |
| Sp947_chr_01101 | Galactose-6-phosphate isomerase subunit LacA | 4.83E-02 | 2.778676 |
| Sp947_chr_00493 | hypothetical protein | 1.56E-04 | 2.785461 |
| Sp947_chr_02029 | Phosphate-binding protein PstS 2 | 2.15E-03 | 2.865929 |
| Sp947_chr_01278 | 50S ribosomal protein L7/L12 | 3.87E-12 | 2.996972 |
| Sp947_chr_00488 | hypothetical protein | 4.04E-02 | 3.028346 |
| Sp947_chr_01585 | hypothetical protein | 1.43E-03 | 3.034917 |
| Sp947_chr_01588 | Lactose transport system permease protein LacF | 8.20E-09 | 3.043549 |
| Sp947_chr_01587 | L-arabinose transport system permease protein AraQ | 4.60E-09 | 3.098046 |
| Sp947_chr_02129 | Alpha-glycerophosphate oxidase | 2.28E-14 | 3.098558 |
| Sp947_chr_01753 | Uric acid permease PucK | 6.97E-06 | 3.127739 |
| Sp947_chr_01583 | N-acetylneuraminate lyase | 8.65E-11 | 3.160283 |
| Sp947_chr_02130 | Glycerol kinase | 6.99E-12 | 3.188476 |
| Sp947_chr_02128 | putative glycerol uptake facilitator protein | 5.65E-06 | 3.28419 |
| Sp947_chr_00056 | Bifunctional purine biosynthesis protein PurH | 4.99E-17 | 3.314739 |
| Sp947_chr_01233 | DNA processing protein DprA | 1.59E-02 | 3.332503 |
| Sp947_chr_00051 | Phosphoribosylformylglycinamidine synthase | 6.17E-29 | 3.43114 |
| Sp947_chr_01599 | hypothetical protein | 6.34E-04 | 3.44936 |
| Sp947_chr_00156 | hypothetical protein | 1.20E-03 | 3.475747 |
| Sp947_chr_00153 | hypothetical protein | 1.72E-53 | 3.578283 |
| Sp947_chr_00155 | hypothetical protein | 3.41E-64 | 3.592961 |
| Sp947_chr_00053 | Phosphoribosylformylglycinamidine cyclo-ligase | 3.79E-13 | 3.631123 |
| Sp947_chr_01107 | PTS system lactose-specific EIICB component | 7.51E-03 | 3.644307 |
| Sp947_chr_00154 | hypothetical protein | 7.99E-47 | 3.765825 |
| Sp947_chr_01835 | Single-stranded DNA-binding protein | 4.27E-06 | 3.783469 |
| Sp947_chr_00050 | Phosphoribosylaminoimidazole-succinocarboxamide synthase | 1.15E-03 | 3.934432 |
| Sp947_chr_00151 | Nisin biosynthesis protein NisC | 1.55E-88 | 4.039514 |
| Sp947_chr_00054 | Phosphoribosylglycinamide formyltransferase | 1.05E-08 | 4.05498 |
| Sp947_chr_01976 | hypothetical protein | 1.43E-02 | 4.070542 |
| Sp947_chr_00491 | hypothetical protein | 1.08E-16 | 4.148617 |
| Sp947_chr_00157 | L-cystine-binding protein TcyA | 1.27E-04 | 4.197553 |
| Sp947_chr_00052 | Amidophosphoribosyltransferase | 2.70E-18 | 4.311536 |
| Sp947_chr_01752 | Xanthine phosphoribosyltransferase | 8.75E-03 | 4.371717 |
| Sp947_chr_01035 | hypothetical protein | 1.51E-02 | 4.523322 |
| Sp947_chr_00152 | hypothetical protein | 1.47E-102 | 4.587529 |
| Sp947_chr_00496 | Bacteriocin lactacin-F subunit LafA | 3.41E-03 | 4.761852 |
| Sp947_chr_01582 | Beta-glucoside kinase | 5.11E-14 | 4.796863 |
| Sp947_chr_01979 | hypothetical protein | 4.17E-02 | 4.975354 |
| Sp947_chr_00150 | Nisin biosynthesis protein NisB | 2.28E-105 | 5.029877 |
| Sp947_chr_00149 | hypothetical protein | 9.98E-80 | 5.84338 |
| Sp947_chr_00055 | hypothetical protein | 2.54E-10 | 5.908767 |
| Sp947_chr_00923 | ComE operon protein 3 | 6.38E-04 | 5.932836 |

| | | | |
|---|---|---|---|
| Sp947_chr_01974 | Ubiquinone/menaquinone biosynthesis C-methyltransferase UbiE | 6.02E-03 | 6.304239 |
| Sp947_chr_01982 | ComG operon protein 1 | 1.51E-04 | 7.042161 |
| Sp947_chr_00133 | hypothetical protein | 5.43E-04 | 7.078549 |
| Sp947_chr_00631 | hypothetical protein | 3.43E-02 | 8.177758 |
| Sp947_chr_01601 | hypothetical protein | 1.07E-02 | 8.512209 |
| Sp947_chr_00134 | hypothetical protein | 4.34E-04 | 8.597105 |
| Sp947_chr_01187 | hypothetical protein | 3.55E-02 | 8.767671 |
| Sp947_chr_00119 | hypothetical protein | 2.86E-02 | 9.128069 |
| Sp947_chr_01977 | hypothetical protein | 3.77E-03 | 10.4118 |
| Sp947_chr_00854 | V-type sodium ATPase subunit B | 1.03E-02 | 10.46683 |
| Sp947_chr_01975 | Ubiquinone biosynthesis O-methyltransferase | 1.60E-02 | 10.52488 |
| Sp947_chr_01949 | hypothetical protein | 4.65E-02 | 12.94222 |
| Sp947_chr_00632 | ABC transporter ATP-binding protein YxdL | 2.09E-02 | 17.25928 |
| Sp947_chr_00111 | Lactococcin-G-processing and transport ATP-binding protein LagD | 1.55E-02 | 18.33763 |
| Sp947_chr_00855 | V-type sodium ATPase subunit D | 7.50E-03 | 19.55662 |
| Sp947_chr_00850 | V-type proton ATPase subunit E | 9.84E-03 | 19.86727 |
| Sp947_chr_00840 | Putative N-acetylmannosamine-6-phosphate 2-epimerase | 2.06E-03 | 24.75437 |
| Sp947_chr_00120 | hypothetical protein | 2.86E-03 | 25.71408 |
| Sp947_chr_00116 | hypothetical protein | 2.91E-03 | 26.96818 |
| Sp947_chr_00851 | V-type sodium ATPase subunit C | 1.83E-03 | 28.20511 |
| Sp947_chr_00843 | hypothetical protein | 6.39E-04 | 31.47893 |
| Sp947_chr_00117 | hypothetical protein | 1.32E-03 | 31.85973 |
| Sp947_chr_00846 | Beta-glucoside kinase | 6.62E-04 | 34.40566 |
| Sp947_chr_00118 | hypothetical protein | 3.54E-04 | 40.73953 |
| Sp947_chr_02078 | hypothetical protein | 1.92E-04 | 40.77519 |
| Sp947_chr_00839 | HTH-type transcriptional regulator RpiR | 1.25E-04 | 44.12853 |
| Sp947_chr_00853 | V-type sodium ATPase catalytic subunit A | 7.65E-05 | 47.19399 |
| Sp947_chr_00848 | hypothetical protein | 6.15E-06 | 76.87348 |
| Sp947_chr_00845 | scyllo-inositol 2-dehydrogenase (NAD(+)) | 2.21E-06 | 85.19736 |
| Sp947_chr_00842 | Sodium/glucose cotransporter | 4.67E-07 | 108.7675 |
| Sp947_chr_00841 | N-acetylneuraminate lyase | 4.39E-07 | 115.1438 |
| Sp947_chr_00844 | Sialidase B | 1.77E-07 | 124.0874 |
| Sp947_chr_02077 | hypothetical protein | 8.20E-09 | 199.0553 |

**Supplementary Table 5.** Pneumococcal differential gene expression in the lungs 6 h post-infection, 9-47M vs 4559M. Genes with fold change (FC) greater than 2 and $p < 0.05$ are shown. FC values highlighted in blue = upregulated in 9-47M, while values highlighted in red = upregulated in 4559M.

| **Locus tag in 9-47-Ear** | **Product** | **padj** | **FC** |
|---|---|---|---|
| Sp947_chr_00301 | 2-dehydro-3-deoxygluconokinase | 1.17E-04 | 9.670733 |
| Sp947_chr_01712 | Lichenan permease IIC component | 3.56E-02 | 8.609312 |
| Sp947_chr_01445 | Multidrug resistance ABC transporter ATP-binding and permease protein | 4.16E-08 | 8.483181 |
| Sp947_chr_00306 | PTS system sorbose-specific EIIB component | 8.86E-03 | 7.513945 |
| Sp947_chr_01447 | Multidrug resistance ABC transporter ATP-binding/permease protein BmrA | 7.53E-09 | 7.288221 |
| Sp947_chr_01448 | Oligopeptide-binding protein SarA | 3.21E-85 | 6.940285 |

| | | | |
|---|---|---|---|
| Sp947_chr_00302 | hypothetical protein | 1.64E-02 | 6.743771 |
| Sp947_chr_01446 | Multidrug resistance ABC transporter ATP-binding and permease protein | 2.12E-05 | 6.075821 |
| Sp947_chr_00307 | N-acetylgalactosamine permease IIC component 1 | 2.53E-02 | 5.42795 |
| Sp947_chr_01674 | putative glycosyltransferase EpsJ | 5.67E-07 | 5.357286 |
| Sp947_chr_00308 | PTS system mannose-specific EIID component | 2.24E-03 | 4.992152 |
| Sp947_chr_00305 | Unsaturated chondroitin disaccharide hydrolase | 9.64E-03 | 4.929947 |
| Sp947_chr_01673 | General stress protein A | 3.78E-09 | 4.891933 |
| Sp947_chr_00664 | Putative HMP/thiamine import ATP-binding protein YkoD | 4.57E-02 | 4.859076 |
| Sp947_chr_00544 | 5%2C10-methylenetetrahydrofolate reductase | 1.52E-13 | 4.549448 |
| Sp947_chr_01081 | L-methionine gamma-lyase | 2.58E-04 | 4.3924 |
| Sp947_chr_01802 | Bifunctional alpha-galactosidase/sucrose kinase AgaSK | 9.20E-03 | 4.294223 |
| Sp947_chr_01671 | Beta-1%2C6-galactofuranosyltransferase WbbI | 7.43E-06 | 4.178201 |
| Sp947_chr_01669 | hypothetical protein | 6.03E-06 | 4.145224 |
| Sp947_chr_01668 | Accessory Sec system protein Asp2 | 4.18E-05 | 3.801593 |
| Sp947_chr_01444 | Cystathionine gamma-synthase/O-acetylhomoserine (thiol)-lyase | 1.32E-04 | 3.780648 |
| Sp947_chr_01749 | Exodeoxyribonuclease | 1.34E-03 | 3.759186 |
| Sp947_chr_01318 | Glutamine-binding periplasmic protein | 4.34E-05 | 3.691227 |
| Sp947_chr_00722 | Biotin transporter BioY2 | 3.56E-02 | 3.673481 |
| Sp947_chr_01672 | General stress protein A | 1.09E-15 | 3.578579 |
| Sp947_chr_01850 | hypothetical protein | 1.54E-08 | 3.371368 |
| Sp947_chr_00104 | putative cell wall hydrolase LytN | 1.48E-09 | 3.28594 |
| Sp947_chr_01711 | Arylsulfatase | 3.74E-02 | 3.285935 |
| Sp947_chr_01951 | 6-phospho-beta-glucosidase GmuD | 2.12E-03 | 3.238088 |
| Sp947_chr_01952 | Lichenan permease IIC component | 3.22E-03 | 3.220857 |
| Sp947_chr_01157 | NADP-dependent glyceraldehyde-3-phosphate dehydrogenase | 3.10E-08 | 3.220747 |
| Sp947_chr_01666 | Protein translocase subunit SecA | 2.62E-06 | 3.188151 |
| Sp947_chr_01849 | hypothetical protein | 3.61E-07 | 3.031185 |
| Sp947_chr_02070 | hypothetical protein | 2.76E-02 | 3.006206 |
| Sp947_chr_01665 | Glycosyltransferase Gtf1 | 1.78E-04 | 2.99951 |
| Sp947_chr_01670 | Accessory Sec system protein translocase subunit SecY2 | 1.04E-03 | 2.985239 |
| Sp947_chr_00543 | 5-methyltetrahydropteroyltriglutamate--homocysteine methyltransferase | 3.12E-12 | 2.938637 |
| Sp947_chr_01731 | hypothetical protein | 1.04E-02 | 2.933719 |
| Sp947_chr_01851 | hypothetical protein | 8.59E-04 | 2.875592 |
| Sp947_chr_01339 | Glucosamine-6-phosphate deaminase | 2.98E-08 | 2.866661 |
| Sp947_chr_01660 | Serine/threonine transporter SstT | 7.70E-07 | 2.831328 |
| Sp947_chr_01664 | Glycosyltransferase-stabilizing protein Gtf2 | 8.36E-04 | 2.742671 |
| Sp947_chr_01848 | Pneumolysin | 5.90E-18 | 2.731212 |
| Sp947_chr_00967 | putative amino acid permease YhdG | 2.01E-04 | 2.694598 |
| Sp947_chr_01281 | 5-methylthioadenosine/S-adenosylhomocysteine deaminase | 1.08E-04 | 2.610116 |
| Sp947_chr_01238 | Ribitol-5-phosphate cytidylyltransferase | 5.25E-07 | 2.603372 |
| Sp947_chr_02040 | N-acetyldiaminopimelate deacetylase | 3.36E-04 | 2.491103 |
| Sp947_chr_01847 | putative transcriptional regulatory protein | 4.75E-08 | 2.46311 |

| | | | |
|---|---|---|---|
| Sp947_chr_01055 | Glucose-6-phosphate 1-dehydrogenase | 8.83E-19 | 2.41423 |
| Sp947_chr_00279 | Alcohol dehydrogenase 1 | 1.79E-04 | 2.402247 |
| Sp947_chr_00696 | High-affinity branched-chain amino acid transport ATP-binding protein LivF | 1.78E-04 | 2.40127 |
| Sp947_chr_00161 | Methionine import ATP-binding protein MetN | 5.07E-03 | 2.36814 |
| Sp947_chr_00695 | Lipopolysaccharide export system ATP-binding protein LptB | 2.38E-03 | 2.365929 |
| Sp947_chr_00162 | Methionine import system permease protein MetP | 2.47E-02 | 2.354175 |
| Sp947_chr_01237 | Ribulose-5-phosphate reductase | 2.21E-07 | 2.347699 |
| Sp947_chr_00694 | High-affinity branched-chain amino acid transport system permease protein LivH | 6.27E-04 | 2.285144 |
| Sp947_chr_02071 | Dihydroxy-acid dehydratase | 9.70E-09 | 2.2807 |
| Sp947_chr_01640 | Ribosomal RNA small subunit methyltransferase B | 2.39E-04 | 2.277034 |
| Sp947_chr_00985 | 4-hydroxy-tetrahydrodipicolinate synthase | 3.06E-04 | 2.222209 |
| Sp947_chr_01421 | L-cystine transport system permease protein YecS | 6.47E-03 | 2.19205 |
| Sp947_chr_01443 | Cystathionine beta-lyase PatB | 4.50E-02 | 2.188948 |
| Sp947_chr_01056 | Arginine transport ATP-binding protein ArtM | 1.14E-10 | 2.147633 |
| Sp947_chr_01661 | hypothetical protein | 1.13E-02 | 2.141693 |
| Sp947_chr_01287 | Adapter protein MecA | 6.92E-03 | 2.108023 |
| Sp947_chr_01420 | Glutamine transport ATP-binding protein GlnQ | 3.04E-04 | 2.064193 |
| Sp947_chr_00191 | Folylpolyglutamate synthase | 3.39E-02 | 2.058539 |
| Sp947_chr_01057 | Glutamine transport system permease protein GlnP | 1.08E-13 | 2.054665 |
| Sp947_chr_00789 | hypothetical protein | 2.13E-04 | 2.044356 |
| Sp947_chr_00787 | Branched-chain-amino-acid aminotransferase | 3.56E-06 | 2.038933 |
| Sp947_chr_00403 | Malonyl CoA-acyl carrier protein transacylase | 1.35E-02 | 2.029537 |
| Sp947_chr_01985 | N-acetylglucosamine-6-phosphate deacetylase | 4.34E-05 | 2.007822 |
| Sp947_chr_01955 | Aldehyde-alcohol dehydrogenase | 2.78E-07 | 2.00535 |
| Sp947_chr_00283 | Dihydropteroate synthase | 1.34E-02 | 2.008932 |
| Sp947_chr_01500 | Trehalose import ATP-binding protein SugC | 3.16E-06 | 2.026012 |
| Sp947_chr_00609 | Sensor histidine kinase YehU | 1.74E-06 | 2.070439 |
| Sp947_chr_00083 | Histidine protein kinase SaeS | 5.36E-03 | 2.073337 |
| Sp947_chr_01155 | Glucose-1-phosphate adenylyltransferase | 8.09E-03 | 2.093897 |
| Sp947_chr_01279 | 50S ribosomal protein L10 | 1.05E-08 | 2.096326 |
| Sp947_chr_01154 | Glycogen biosynthesis protein GlgD | 8.29E-03 | 2.098713 |
| Sp947_chr_01590 | PTS system glucose-specific EIICBA component | 5.60E-14 | 2.138227 |
| Sp947_chr_00500 | hypothetical protein | 1.66E-06 | 2.139178 |
| Sp947_chr_00502 | hypothetical protein | 7.03E-05 | 2.147247 |
| Sp947_chr_01589 | Putative ABC transporter substrate-binding protein YesO | 3.23E-13 | 2.149307 |
| Sp947_chr_00066 | PTS system sorbose-specific EIIC component | 1.90E-03 | 2.153142 |
| Sp947_chr_00065 | PTS system sorbose-specific EIIB component | 2.53E-02 | 2.169443 |
| Sp947_chr_00069 | Putative tagatose-6-phosphate ketose/aldose isomerase | 4.34E-05 | 2.18834 |
| Sp947_chr_00925 | hypothetical protein | 4.99E-02 | 2.232677 |
| Sp947_chr_00067 | PTS system mannose-specific EIID component | 2.72E-04 | 2.245442 |
| Sp947_chr_01591 | Putative N-acetylmannosamine-6-phosphate 2-epimerase | 1.55E-08 | 2.259193 |

| | | | |
|---|---|---|---|
| Sp947_chr_00284 | Folylpolyglutamate synthase | 1.47E-04 | 2.262639 |
| Sp947_chr_00582 | 50S ribosomal protein L1 | 1.92E-11 | 2.266385 |
| Sp947_chr_01478 | 4-hydroxy-tetrahydrodipicolinate reductase | 1.95E-03 | 2.316861 |
| Sp947_chr_00228 | 50S ribosomal protein L17 | 7.30E-11 | 2.339842 |
| Sp947_chr_00608 | putative response regulatory protein | 7.07E-07 | 2.352334 |
| Sp947_chr_00499 | hypothetical protein | 3.48E-04 | 2.364047 |
| Sp947_chr_00930 | Lactoylglutathione lyase | 3.61E-02 | 2.382768 |
| Sp947_chr_02030 | Phosphate transport system permease protein PstC 1 | 1.96E-02 | 2.434936 |
| Sp947_chr_01479 | DegV domain-containing protein | 4.56E-04 | 2.439687 |
| Sp947_chr_01600 | Sialidase A | 1.52E-13 | 2.448881 |
| Sp947_chr_00281 | Guanine/hypoxanthine permease PbuO | 1.26E-07 | 2.458395 |
| Sp947_chr_00059 | N5-carboxyaminoimidazole ribonucleotide synthase | 6.17E-05 | 2.458645 |
| Sp947_chr_01584 | hypothetical protein | 3.37E-03 | 2.507546 |
| Sp947_chr_01599 | hypothetical protein | 1.54E-02 | 2.558236 |
| Sp947_chr_01592 | scyllo-inositol 2-dehydrogenase (NAD(+)) | 4.91E-07 | 2.616868 |
| Sp947_chr_00285 | GTP cyclohydrolase 1 | 1.08E-03 | 2.622738 |
| Sp947_chr_00577 | Branched-chain amino acid transport system 2 carrier protein | 1.03E-07 | 2.63016 |
| Sp947_chr_00057 | Phosphoribosylamine--glycine ligase | 1.46E-03 | 2.644768 |
| Sp947_chr_00493 | hypothetical protein | 2.13E-04 | 2.678999 |
| Sp947_chr_02032 | Phosphate import ATP-binding protein PstB 3 | 1.01E-03 | 2.722278 |
| Sp947_chr_01598 | Evolved beta-galactosidase subunit beta | 1.56E-04 | 2.78706 |
| Sp947_chr_00154 | hypothetical protein | 2.65E-28 | 2.805147 |
| Sp947_chr_01597 | Putative ABC transporter substrate-binding protein YesO | 1.60E-16 | 2.844123 |
| Sp947_chr_01586 | Toxin-antitoxin biofilm protein TabA | 1.39E-03 | 2.849225 |
| Sp947_chr_02031 | Phosphate transport system permease protein PstA 2 | 5.46E-04 | 2.861119 |
| Sp947_chr_01596 | Inner membrane ABC transporter permease protein YcjO | 2.53E-07 | 2.886446 |
| Sp947_chr_00153 | hypothetical protein | 1.82E-37 | 2.892108 |
| Sp947_chr_00155 | hypothetical protein | 1.12E-45 | 2.931856 |
| Sp947_chr_00151 | Nisin biosynthesis protein NisC | 7.90E-55 | 2.999037 |
| Sp947_chr_00282 | hypothetical protein | 6.70E-03 | 3.060665 |
| Sp947_chr_02033 | Phosphate-specific transport system accessory protein PhoU | 2.38E-05 | 3.077587 |
| Sp947_chr_01595 | Inner membrane ABC transporter permease protein YcjP | 1.02E-07 | 3.127981 |
| Sp947_chr_01588 | Lactose transport system permease protein LacF | 6.39E-10 | 3.167722 |
| Sp947_chr_01583 | N-acetylneuraminate lyase | 1.77E-11 | 3.181516 |
| Sp947_chr_00488 | hypothetical protein | 2.55E-02 | 3.198131 |
| Sp947_chr_01278 | 50S ribosomal protein L7/L12 | 2.57E-14 | 3.199195 |
| Sp947_chr_02062 | hypothetical protein | 1.23E-02 | 3.218262 |
| Sp947_chr_00157 | L-cystine-binding protein TcyA | 1.97E-03 | 3.281804 |
| Sp947_chr_02128 | putative glycerol uptake facilitator protein | 1.40E-06 | 3.376722 |
| Sp947_chr_00152 | hypothetical protein | 1.56E-66 | 3.408858 |
| Sp947_chr_01587 | L-arabinose transport system permease protein AraQ | 2.04E-11 | 3.436735 |
| Sp947_chr_01108 | 6-phospho-beta-galactosidase | 3.63E-02 | 3.572337 |
| Sp947_chr_00150 | Nisin biosynthesis protein NisB | 3.01E-67 | 3.640087 |

| | | | |
|---|---|---|---|
| Sp947_chr_00053 | Phosphoribosylformylglycinamidine cyclo-ligase | 7.11E-14 | 3.646901 |
| Sp947_chr_01835 | Single-stranded DNA-binding protein | 4.27E-06 | 3.681599 |
| Sp947_chr_02130 | Glycerol kinase | 5.10E-16 | 3.692078 |
| Sp947_chr_01585 | hypothetical protein | 2.56E-05 | 3.869586 |
| Sp947_chr_00056 | Bifunctional purine biosynthesis protein PurH | 3.59E-23 | 3.888367 |
| Sp947_chr_00496 | Bacteriocin lactacin-F subunit LafA | 1.15E-02 | 3.987266 |
| Sp947_chr_00054 | Phosphoribosylglycinamide formyltransferase | 8.51E-09 | 3.987798 |
| Sp947_chr_00983 | asd | 9.32E-03 | 3.997549 |
| Sp947_chr_02129 | Alpha-glycerophosphate oxidase | 3.55E-23 | 4.018257 |
| Sp947_chr_01035 | hypothetical protein | 2.41E-02 | 4.140521 |
| Sp947_chr_00051 | Phosphoribosylformylglycinamidine synthase | 2.28E-42 | 4.235246 |
| Sp947_chr_00052 | Amidophosphoribosyltransferase | 1.19E-18 | 4.268644 |
| Sp947_chr_02029 | Phosphate-binding protein PstS 2 | 1.30E-06 | 4.397577 |
| Sp947_chr_01753 | Uric acid permease PucK | 2.01E-10 | 4.406739 |
| Sp947_chr_01976 | hypothetical protein | 6.51E-03 | 4.475349 |
| Sp947_chr_02096 | hypothetical protein | 2.68E-02 | 4.506143 |
| Sp947_chr_00491 | hypothetical protein | 3.28E-20 | 4.611629 |
| Sp947_chr_01582 | Beta-glucoside kinase | 2.18E-14 | 4.75467 |
| Sp947_chr_00050 | Phosphoribosylaminoimidazole-succinocarboxamide synthase | 7.03E-05 | 4.86234 |
| Sp947_chr_00149 | hypothetical protein | 9.11E-65 | 4.877677 |
| Sp947_chr_00923 | ComE operon protein 3 | 1.39E-03 | 5.294791 |
| Sp947_chr_01752 | Xanthine phosphoribosyltransferase | 8.53E-04 | 5.746786 |
| Sp947_chr_00055 | hypothetical protein | 9.95E-11 | 5.925865 |
| Sp947_chr_01974 | Ubiquinone/menaquinone biosynthesis C-methyltransferase UbiE | 5.04E-03 | 6.407721 |
| Sp947_chr_01601 | hypothetical protein | 2.99E-02 | 6.738993 |
| Sp947_chr_00133 | hypothetical protein | 3.54E-04 | 7.246645 |
| Sp947_chr_01979 | hypothetical protein | 5.49E-03 | 7.304412 |
| Sp947_chr_01980 | hypothetical protein | 1.67E-02 | 7.537585 |
| Sp947_chr_01982 | ComG operon protein 1 | 1.75E-05 | 8.446919 |
| Sp947_chr_00134 | hypothetical protein | 4.13E-04 | 8.447782 |
| Sp947_chr_00631 | hypothetical protein | 2.15E-02 | 9.213989 |
| Sp947_chr_01977 | hypothetical protein | 5.49E-03 | 9.600591 |
| Sp947_chr_00119 | hypothetical protein | 2.24E-02 | 9.657972 |
| Sp947_chr_01975 | Ubiquinone biosynthesis O-methyltransferase | 1.24E-02 | 11.19827 |
| Sp947_chr_00849 | V-type sodium ATPase subunit K | 4.67E-02 | 12.68675 |
| Sp947_chr_00847 | hypothetical protein | 4.77E-02 | 12.75212 |
| Sp947_chr_00632 | ABC transporter ATP-binding protein YxdL | 3.91E-02 | 13.87838 |
| Sp947_chr_00115 | hypothetical protein | 4.98E-02 | 15.08898 |
| Sp947_chr_00122 | hypothetical protein | 7.13E-03 | 19.74573 |
| Sp947_chr_00852 | V-type sodium ATPase subunit G | 9.01E-03 | 20.06019 |
| Sp947_chr_00843 | hypothetical protein | 3.51E-03 | 21.00858 |
| Sp947_chr_00854 | V-type sodium ATPase subunit B | 2.61E-04 | 21.55684 |
| Sp947_chr_00111 | Lactococcin-G-processing and transport ATP-binding protein LagD | 7.18E-03 | 22.70188 |
| Sp947_chr_00850 | V-type proton ATPase subunit E | 1.94E-03 | 29.81791 |
| Sp947_chr_00116 | hypothetical protein | 1.53E-03 | 30.65687 |
| Sp947_chr_00840 | Putative N-acetylmannosamine-6-phosphate 2-epimerase | 6.27E-04 | 31.88306 |
| Sp947_chr_00855 | V-type sodium ATPase subunit D | 7.10E-04 | 33.3989 |
| Sp947_chr_00851 | V-type sodium ATPase subunit C | 7.66E-04 | 33.41091 |

| | | | |
|---|---|---|---|
| Sp947_chr_00839 | HTH-type transcriptional regulator RpiR | 4.17E-04 | 33.69369 |
| Sp947_chr_00846 | Beta-glucoside kinase | 6.02E-04 | 34.40359 |
| Sp947_chr_02078 | hypothetical protein | 2.89E-04 | 37.03186 |
| Sp947_chr_00117 | hypothetical protein | 6.19E-04 | 37.13564 |
| Sp947_chr_00120 | hypothetical protein | 3.83E-04 | 39.34665 |
| Sp947_chr_00853 | V-type sodium ATPase catalytic subunit A | 8.31E-05 | 46.29248 |
| Sp947_chr_00118 | hypothetical protein | 1.01E-04 | 52.43328 |
| Sp947_chr_00845 | scyllo-inositol 2-dehydrogenase (NAD(+)) | 6.99E-07 | 100.7781 |
| Sp947_chr_00848 | hypothetical protein | 5.25E-07 | 111.61 |
| Sp947_chr_00841 | N-acetylneuraminate lyase | 4.68E-07 | 112.2849 |
| Sp947_chr_00842 | Sodium/glucose cotransporter | 2.63E-07 | 118.3059 |
| Sp947_chr_00844 | Sialidase B | 4.59E-08 | 151.1713 |
| Sp947_chr_02077 | hypothetical protein | 6.25E-09 | 206.1168 |

**Supplementary Table 6.** Pneumococcal differential gene expression in the lungs 6 h post-infection, 4559-Blood vs 4559M. Genes with fold change (FC) greater than 2 and $p < 0.05$ are shown. FC values highlighted in blue = upregulated in 4559-Blood, while values highlighted in red = upregulated in 4559M.

| Locus tag in 9-47-Ear | Product | padj | FC |
|---|---|---|---|
| Sp947_chr_00279 | Alcohol dehydrogenase 1 | 2.23E-08 | 3.532144 |
| Sp947_chr_01955 | Aldehyde-alcohol dehydrogenase | 1.15E-20 | 3.234431 |
| Sp947_chr_00088 | putative multiple-sugar transport system permease YteP | 3.50E-02 | 2.176402 |
| Sp947_chr_02054 | Maltose transport system permease protein MalG | 9.80E-03 | 2.120116 |
| Sp947_chr_01917 | Transcriptional regulatory protein DesR | 1.14E-02 | 2.254388 |
| Sp947_chr_02033 | Phosphate-specific transport system accessory protein PhoU | 5.24E-03 | 2.313659 |
| Sp947_chr_01629 | PTS system trehalose-specific EIIBC component | 1.03E-02 | 3.264913 |
| Sp947_chr_01920 | putative ABC transporter ATP-binding protein YxlF | 1.75E-04 | 3.369874 |

**Supplementary Table 7.** Murine differential gene expression in the lungs 6 h post-infection, 9-47-Ear vs 9-47M. Genes with fold change (FC) greater than 1.5 and $p < 0.05$ are shown. FC values highlighted in blue = upregulated in 9-47-Ear, while values highlighted in red = upregulated in 9-47M.

| Gene Stable ID | Gene Name | padj | FC |
|---|---|---|---|
| ENSMUSG00000065254 | *Gm23973* | 1.17E-08 | 6.0256 |
| ENSMUSG00000088675 | *Rprl1* | 2.45E-08 | 5.382293 |
| ENSMUSG00000106222 | *Rprl1* | 2.45E-08 | 5.382293 |
| ENSMUSG00000096269 | *Rpl31-ps21* | 6.02E-06 | 5.232448 |
| ENSMUSG00000094668 | *Gm24871* | 1.75E-06 | 4.724852 |
| ENSMUSG00000064682 | *Gm25813* | 1.04E-09 | 4.405394 |
| ENSMUSG00000065870 | *Rnu3a* | 4.31E-15 | 4.246867 |
| ENSMUSG00000106147 | *Rnu3a* | 4.31E-15 | 4.246867 |
| ENSMUSG00000095676 | *Gm25099* | 2.12E-22 | 4.006482 |

| | | | |
|---|---|---|---|
| ENSMUSG00000104896 | *Rnu3b4* | 2.62E-19 | 3.731483 |
| ENSMUSG00000105115 | *Rnu3b2* | 1.58E-19 | 3.730893 |
| ENSMUSG00000098925 | *Rnu3b2* | 1.58E-19 | 3.730893 |
| ENSMUSG00000098641 | *Rnu3b4* | 3.16E-19 | 3.726678 |
| ENSMUSG00000098943 | *Rnu3b1* | 5.09E-19 | 3.713982 |
| ENSMUSG00000105025 | *Rnu3b1* | 5.09E-19 | 3.713982 |
| ENSMUSG00000099291 | *Rnu3b3* | 1.37E-18 | 3.710655 |
| ENSMUSG00000104856 | *Rnu3b3* | 1.37E-18 | 3.710655 |
| ENSMUSG00000065232 | *Gm22973* | 7.53E-12 | 3.699468 |
| ENSMUSG00000095738 | *Gm25313* | 7.10E-11 | 3.614973 |
| ENSMUSG00000064994 | *Gm22422* | 2.64E-07 | 3.494379 |
| ENSMUSG00000065176 | *Rnu12* | 2.59E-22 | 3.424881 |
| ENSMUSG00000077254 | *Gm26079* | 7.16E-09 | 3.417376 |
| ENSMUSG00000095892 | *Rnu5g* | 1.98E-10 | 3.377252 |
| ENSMUSG00000093956 | *Gm24497* | 1.37E-09 | 3.358284 |
| ENSMUSG00000094050 | *Gm23472* | 7.29E-10 | 3.353025 |
| ENSMUSG00000080465 | *Gm22486* | 2.03E-10 | 3.35003 |
| ENSMUSG00000064856 | *Gm23444* | 1.29E-09 | 3.346978 |
| ENSMUSG00000064899 | *Snord118* | 1.39E-05 | 3.336753 |
| ENSMUSG00000093843 | *Gm25939* | 5.28E-08 | 3.334159 |
| ENSMUSG00000094655 | *Gm25360* | 1.09E-16 | 3.3217 |
| ENSMUSG00000065820 | *Gm26316* | 1.36E-09 | 3.301702 |
| ENSMUSG00000065767 | *Gm23849* | 3.38E-09 | 3.230541 |
| ENSMUSG00000064702 | *Gm24950* | 4.61E-09 | 3.217205 |
| ENSMUSG00000065251 | *Gm23971* | 4.61E-09 | 3.217205 |
| ENSMUSG00000065944 | *Rnu2-10* | 3.79E-09 | 3.210903 |
| ENSMUSG00000080538 | *Gm25541* | 1.66E-11 | 3.172594 |
| ENSMUSG00000027261 | *Hao1* | 7.14E-05 | 3.03458 |
| ENSMUSG00000094377 | *Gm24407* | 3.83E-10 | 3.022053 |
| ENSMUSG00000032494 | *Tdgf1* | 4.29E-04 | 3.003122 |
| ENSMUSG00000065911 | *Gm24447* | 3.13E-11 | 2.969689 |
| ENSMUSG00000065824 | *Gm26315* | 3.28E-11 | 2.966459 |
| ENSMUSG00000089255 | *Snora78* | 5.52E-04 | 2.891999 |
| ENSMUSG00000096243 | *Gm24265* | 9.59E-08 | 2.856449 |
| ENSMUSG00000064945 | *Rny3* | 3.87E-08 | 2.845147 |
| ENSMUSG00000085348 | *Myhas* | 3.19E-07 | 2.838211 |
| ENSMUSG00000064694 | *Gm24146* | 1.10E-06 | 2.83174 |
| ENSMUSG00000065087 | *Snord22* | 2.41E-26 | 2.800736 |
| ENSMUSG00000115207 | *Gm49132* | 4.18E-02 | 2.775687 |
| ENSMUSG00000084744 | *Gm25291* | 1.83E-05 | 2.768169 |
| ENSMUSG00000091993 | *B930036N10Rik* | 2.04E-05 | 2.722677 |
| ENSMUSG00000084708 | *Gm22988* | 4.52E-11 | 2.714612 |
| ENSMUSG00000114163 | *Gm32296* | 2.07E-03 | 2.66448 |
| ENSMUSG00000010362 | *Rdm1* | 7.28E-09 | 2.661396 |
| ENSMUSG00000064453 | *Snord21* | 2.98E-04 | 2.630897 |
| ENSMUSG00000055170 | *Ifng* | 3.23E-05 | 2.609289 |
| ENSMUSG00000044092 | *C130050O18Rik* | 5.69E-17 | 2.609004 |
| ENSMUSG00000000386 | *Mx1* | 3.70E-72 | 2.588273 |
| ENSMUSG00000065118 | *Gm23297* | 5.39E-04 | 2.56856 |
| ENSMUSG00000064966 | *Snord15b* | 5.58E-08 | 2.566291 |
| ENSMUSG00000065701 | *Rny1* | 2.33E-06 | 2.562933 |
| ENSMUSG00000087819 | *Gm25117* | 1.86E-10 | 2.551007 |
| ENSMUSG00000111734 | *Gm29825* | 1.81E-02 | 2.544426 |

| | | | |
|---|---|---|---|
| ENSMUSG00000010342 | *Tex14* | 8.31E-10 | 2.529067 |
| ENSMUSG00000064901 | *Snora21* | 1.95E-03 | 2.517886 |
| ENSMUSG00000087775 | *Rprl2* | 4.05E-05 | 2.454835 |
| ENSMUSG00000065822 | *Snord15a* | 1.38E-04 | 2.451242 |
| ENSMUSG00000097770 | *Gm26776* | 9.57E-03 | 2.44156 |
| ENSMUSG00000020641 | *Rsad2* | 7.09E-26 | 2.429777 |
| ENSMUSG00000043681 | *Fam25c* | 1.39E-04 | 2.404388 |
| ENSMUSG00000003545 | *Fosb* | 5.64E-13 | 2.404057 |
| ENSMUSG00000025431 | *Crisp1* | 9.80E-05 | 2.397804 |
| ENSMUSG00000101133 | *Gm29050* | 8.16E-03 | 2.391731 |
| ENSMUSG00000088252 | *Snord13* | 6.50E-05 | 2.386827 |
| ENSMUSG00000077323 | *Rnu11* | 6.57E-07 | 2.38433 |
| ENSMUSG00000097131 | *D230017M19Rik* | 1.76E-08 | 2.381225 |
| ENSMUSG00000092837 | *Rpph1* | 7.69E-06 | 2.372261 |
| ENSMUSG00000071470 | *Ccnb1ip1* | 7.38E-06 | 2.360943 |
| ENSMUSG00000092819 | *Gm23639* | 7.41E-03 | 2.352266 |
| ENSMUSG00000048806 | *Ifnb1* | 7.71E-03 | 2.351491 |
| ENSMUSG00000117269 | *AC151275.1* | 2.17E-05 | 2.349181 |
| ENSMUSG00000087477 | *Gm13822* | 8.85E-05 | 2.342073 |
| ENSMUSG00000065649 | *Snora74a* | 1.06E-06 | 2.339372 |
| ENSMUSG00000105797 | *Gm5149* | 1.85E-04 | 2.330504 |
| ENSMUSG00000035692 | *Isg15* | 4.03E-49 | 2.315568 |
| ENSMUSG00000069310 | *Hist1h3c* | 2.16E-03 | 2.278831 |
| ENSMUSG00000090101 | *Snhg9* | 2.99E-03 | 2.269589 |
| ENSMUSG00000020216 | *Jsrp1* | 4.96E-03 | 2.263278 |
| ENSMUSG00000069265 | *Hist1h3a* | 8.88E-03 | 2.259961 |
| ENSMUSG00000104863 | *Gm49347* | 5.93E-05 | 2.258448 |
| ENSMUSG00000099517 | *Hist1h3g* | 1.60E-02 | 2.249359 |
| ENSMUSG00000057596 | *Trim30d* | 3.93E-09 | 2.247087 |
| ENSMUSG00000064604 | *Snora44* | 2.55E-04 | 2.227773 |
| ENSMUSG00000015437 | *Gzmb* | 1.49E-07 | 2.22751 |
| ENSMUSG00000046229 | *Scand1* | 7.61E-03 | 2.223718 |
| ENSMUSG00000077167 | *Gm24119* | 2.38E-03 | 2.220322 |
| ENSMUSG00000065126 | *Snord104* | 3.21E-09 | 2.21526 |
| ENSMUSG00000096887 | *Gm20594* | 7.60E-12 | 2.198159 |
| ENSMUSG00000080542 | *Gm22710* | 8.69E-04 | 2.195982 |
| ENSMUSG00000064999 | *Gm26035* | 2.79E-06 | 2.192941 |
| ENSMUSG00000069273 | *Hist1h3e* | 3.58E-06 | 2.184059 |
| ENSMUSG00000080365 | *Gm25776* | 1.43E-04 | 2.181141 |
| ENSMUSG00000031289 | *Il13ra2* | 4.89E-06 | 2.177819 |
| ENSMUSG00000016327 | *Atp1b4* | 1.73E-02 | 2.166814 |
| ENSMUSG00000064380 | *Gm26448* | 1.91E-09 | 2.16627 |
| ENSMUSG00000111821 | *Gm48545* | 9.69E-05 | 2.152735 |
| ENSMUSG00000097060 | *Gm26759* | 9.57E-10 | 2.145737 |
| ENSMUSG00000034855 | *Cxcl10* | 6.33E-14 | 2.12518 |
| ENSMUSG00000067455 | *Hist1h4j* | 7.47E-04 | 2.121665 |
| ENSMUSG00000045932 | *Ifit2* | 1.49E-41 | 2.119366 |
| ENSMUSG00000096349 | *Gm22513* | 1.67E-02 | 2.109175 |
| ENSMUSG00000064655 | *Gm25788* | 7.16E-03 | 2.098184 |
| ENSMUSG00000094344 | *Gm11942* | 7.06E-20 | 2.094663 |
| ENSMUSG00000023341 | *Mx2* | 4.66E-63 | 2.093454 |
| ENSMUSG00000060093 | *Hist1h4a* | 7.15E-07 | 2.093403 |
| ENSMUSG00000030670 | *Cyp2r1* | 5.70E-03 | 2.090606 |

| | | | |
|---|---|---|---|
| ENSMUSG00000044748 | *Defb1* | 3.78E-02 | 2.074292 |
| ENSMUSG00000107722 | *2900060B14Rik* | 1.19E-03 | 2.067021 |
| ENSMUSG00000031757 | *Mt4* | 1.78E-04 | 2.063615 |
| ENSMUSG00000088025 | *Rprl3* | 1.20E-03 | 2.055532 |
| ENSMUSG00000095590 | *Gm24305* | 4.50E-02 | 2.053062 |
| ENSMUSG00000081163 | *Defa-ps12* | 2.72E-02 | 2.051928 |
| ENSMUSG00000038357 | *Camp* | 9.67E-07 | 2.049958 |
| ENSMUSG00000097457 | *2310031A07Rik* | 4.24E-02 | 2.038639 |
| ENSMUSG00000064387 | *Snora73a* | 2.52E-11 | 2.036576 |
| ENSMUSG00000031343 | *Gabra3* | 1.84E-06 | 2.035071 |
| ENSMUSG00000115420 | *AL732506.1* | 1.46E-06 | 2.031062 |
| ENSMUSG00000088088 | *Rmrp* | 1.46E-06 | 2.031062 |
| ENSMUSG00000086336 | *Gm12979* | 4.46E-02 | 2.029161 |
| ENSMUSG00000042761 | *Mrap2* | 8.11E-03 | 2.023479 |
| ENSMUSG00000066443 | *Gm10163* | 1.60E-02 | 2.02177 |
| ENSMUSG00000110421 | *Gm9689* | 8.49E-03 | 2.017106 |
| ENSMUSG00000058385 | *Hist1h2bg* | 2.16E-05 | 2.015475 |
| ENSMUSG00000069274 | *Hist1h4f* | 6.77E-04 | 2.001288 |
| ENSMUSG00000083596 | *Rpl21-ps15* | 2.72E-02 | 1.997761 |
| ENSMUSG00000020638 | *Cmpk2* | 1.22E-42 | 1.997189 |
| ENSMUSG00000046748 | *Tmem45a2* | 1.42E-06 | 1.980568 |
| ENSMUSG00000114279 | *Hist1h2bm* | 3.13E-02 | 1.978823 |
| ENSMUSG00000088948 | *Gm23262* | 5.31E-05 | 1.970935 |
| ENSMUSG00000029019 | *Nppb* | 2.19E-04 | 1.958092 |
| ENSMUSG00000102189 | *Gm37194* | 8.10E-06 | 1.955291 |
| ENSMUSG00000059230 | *Defb4* | 2.95E-05 | 1.953494 |
| ENSMUSG00000106366 | *Mir3069* | 3.60E-02 | 1.950774 |
| ENSMUSG00000065259 | *Snora30* | 2.68E-02 | 1.943307 |
| ENSMUSG00000017300 | *Tnnc2* | 1.69E-15 | 1.943187 |
| ENSMUSG00000033213 | *AA467197* | 1.20E-05 | 1.94194 |
| ENSMUSG00000065353 | *Snora73b* | 3.61E-05 | 1.941924 |
| ENSMUSG00000062461 | *Gm5453* | 2.89E-10 | 1.940423 |
| ENSMUSG00000100551 | *Gm19503* | 1.62E-07 | 1.936219 |
| ENSMUSG00000060639 | *Hist1h4i* | 6.90E-04 | 1.935604 |
| ENSMUSG00000087968 | *Gm25395* | 8.91E-07 | 1.934721 |
| ENSMUSG00000106649 | *Gm49349* | 3.20E-06 | 1.929867 |
| ENSMUSG00000068397 | *Gm10240* | 2.63E-02 | 1.927078 |
| ENSMUSG00000100210 | *Hist1h3f* | 8.72E-04 | 1.921571 |
| ENSMUSG00000004814 | *Ccl24* | 4.06E-02 | 1.915198 |
| ENSMUSG00000063001 | *Gm9701* | 1.53E-02 | 1.914728 |
| ENSMUSG00000047501 | *Cldn4* | 5.80E-05 | 1.913371 |
| ENSMUSG00000091028 | *Gm10722* | 2.29E-02 | 1.902823 |
| ENSMUSG00000065686 | *Snora5c* | 2.53E-02 | 1.902247 |
| ENSMUSG00000114456 | *Hist1h2bh* | 4.42E-03 | 1.899751 |
| ENSMUSG00000030107 | *Usp18* | 4.19E-35 | 1.891192 |
| ENSMUSG00000108332 | *D530033B14Rik* | 5.32E-17 | 1.887082 |
| ENSMUSG00000065226 | *Gm25791* | 4.61E-06 | 1.885368 |
| ENSMUSG00000027233 | *Patl2* | 4.30E-02 | 1.87672 |
| ENSMUSG00000052837 | *Junb* | 3.04E-33 | 1.873304 |
| ENSMUSG00000045027 | *Prss22* | 2.77E-08 | 1.873299 |
| ENSMUSG00000095217 | *Hist1h2bn* | 6.37E-03 | 1.869101 |
| ENSMUSG00000065145 | *Vaultrc5* | 1.08E-03 | 1.86865 |
| ENSMUSG00000046049 | *Rp1l1* | 1.93E-03 | 1.864557 |

| | | | |
|---|---|---|---|
| ENSMUSG00000085532 | *B430319H21Rik* | 7.16E-05 | 1.864074 |
| ENSMUSG00000069266 | *Hist1h4b* | 1.11E-02 | 1.861964 |
| ENSMUSG00000074001 | *Klhl40* | 9.93E-04 | 1.861333 |
| ENSMUSG00000085806 | *Gm12023* | 1.61E-04 | 1.860825 |
| ENSMUSG00000026535 | *Ifi202b* | 5.49E-24 | 1.860159 |
| ENSMUSG00000041827 | *Oasl1* | 1.59E-25 | 1.858532 |
| ENSMUSG00000038295 | *Atg9b* | 2.43E-03 | 1.857709 |
| ENSMUSG00000094392 | *Gm3788* | 6.33E-05 | 1.85749 |
| ENSMUSG00000058773 | *Hist1h1b* | 1.03E-03 | 1.854963 |
| ENSMUSG00000075031 | *Hist1h2bb* | 1.13E-02 | 1.853781 |
| ENSMUSG00000089542 | *Gm25835* | 2.47E-06 | 1.852865 |
| ENSMUSG00000106392 | *Gm5870* | 4.66E-02 | 1.852755 |
| ENSMUSG00000083832 | *Gm13771* | 1.90E-05 | 1.846272 |
| ENSMUSG00000074433 | *Lce3e* | 1.17E-02 | 1.844793 |
| ENSMUSG00000090293 | *Gm17034* | 3.68E-02 | 1.844258 |
| ENSMUSG00000088008 | *Gm25492* | 1.09E-02 | 1.841263 |
| ENSMUSG00000082069 | *Gm16242* | 2.24E-03 | 1.83945 |
| ENSMUSG00000107480 | *Gm44165* | 8.52E-05 | 1.836863 |
| ENSMUSG00000064288 | *Hist1h4k* | 1.54E-03 | 1.836703 |
| ENSMUSG00000091383 | *Hist1h2al* | 1.56E-06 | 1.834135 |
| ENSMUSG00000070644 | *Etnk2* | 2.72E-02 | 1.83037 |
| ENSMUSG00000034459 | *Ifit1* | 9.35E-29 | 1.829028 |
| ENSMUSG00000108414 | *Snhg1* | 1.47E-18 | 1.82864 |
| ENSMUSG00000101191 | *Gm28809* | 1.83E-02 | 1.825134 |
| ENSMUSG00000095580 | *Rnu1b1* | 1.57E-03 | 1.824702 |
| ENSMUSG00000093834 | *Rnu1b2* | 1.57E-03 | 1.824702 |
| ENSMUSG00000087090 | *Nctc1* | 3.87E-14 | 1.821732 |
| ENSMUSG00000031231 | *Cox7b* | 1.15E-18 | 1.818275 |
| ENSMUSG00000064451 | *Snora23* | 4.39E-02 | 1.815006 |
| ENSMUSG00000026418 | *Tnni1* | 8.22E-03 | 1.811313 |
| ENSMUSG00000022586 | *Ly6i* | 6.13E-15 | 1.806331 |
| ENSMUSG00000102135 | *Gm37108* | 2.29E-04 | 1.803613 |
| ENSMUSG00000057465 | *Saa2* | 2.25E-09 | 1.803135 |
| ENSMUSG00000018930 | *Ccl4* | 5.57E-23 | 1.800731 |
| ENSMUSG00000104956 | *4930429D17Rik* | 1.82E-02 | 1.800304 |
| ENSMUSG00000049173 | *Myoz3* | 2.97E-04 | 1.793444 |
| ENSMUSG00000105361 | *AY036118* | 4.61E-03 | 1.788552 |
| ENSMUSG00000104862 | *Gm6520* | 5.48E-03 | 1.787612 |
| ENSMUSG00000088529 | *Gm26083* | 3.15E-03 | 1.7854 |
| ENSMUSG00000085241 | *Snhg3* | 8.45E-07 | 1.784438 |
| ENSMUSG00000068885 | *Lce3f* | 3.45E-03 | 1.781149 |
| ENSMUSG00000085609 | *1700016P03Rik* | 1.93E-02 | 1.781038 |
| ENSMUSG00000052305 | *Hbb-bs* | 1.24E-04 | 1.779109 |
| ENSMUSG00000069306 | *Hist1h4m* | 8.36E-03 | 1.776996 |
| ENSMUSG00000074403 | *Hist2h3b* | 4.10E-02 | 1.775714 |
| ENSMUSG00000038508 | *Gdf15* | 1.58E-02 | 1.774828 |
| ENSMUSG00000111713 | *Gm20234* | 3.65E-02 | 1.77292 |
| ENSMUSG00000105826 | *Gm42141* | 1.78E-02 | 1.772001 |
| ENSMUSG00000068855 | *Hist2h2ac* | 3.41E-03 | 1.763701 |
| ENSMUSG00000064923 | *Gm22042* | 2.24E-03 | 1.762643 |
| ENSMUSG00000033196 | *Myh2* | 1.57E-05 | 1.760939 |
| ENSMUSG00000095701 | *Gm24830* | 2.16E-03 | 1.760782 |
| ENSMUSG00000096838 | *Gm26232* | 2.16E-03 | 1.760782 |

| | | | |
|---|---|---|---|
| ENSMUSG00000065773 | *Rnu1b6* | 2.16E-03 | 1.760782 |
| ENSMUSG00000092702 | *Gm24514* | 2.48E-03 | 1.758247 |
| ENSMUSG00000031097 | *Tnni2* | 2.03E-10 | 1.757473 |
| ENSMUSG00000021768 | *Dusp13* | 1.61E-02 | 1.754751 |
| ENSMUSG00000068697 | *Myoz1* | 2.09E-06 | 1.754697 |
| ENSMUSG00000058427 | *Cxcl2* | 4.52E-11 | 1.754009 |
| ENSMUSG00000096010 | *Hist4h4* | 8.63E-04 | 1.751589 |
| ENSMUSG00000100969 | *1700030N03Rik* | 1.26E-04 | 1.75055 |
| ENSMUSG00000065037 | *Rn7sk* | 2.99E-04 | 1.749012 |
| ENSMUSG00000061482 | *Hist1h4d* | 9.27E-08 | 1.745652 |
| ENSMUSG00000081159 | *Gm6023* | 3.99E-02 | 1.745032 |
| ENSMUSG00000077714 | *Snord17* | 9.77E-04 | 1.74339 |
| ENSMUSG00000077563 | *Snora68* | 1.10E-03 | 1.741607 |
| ENSMUSG00000059434 | *Gckr* | 1.39E-05 | 1.737341 |
| ENSMUSG00000036731 | *Cysrt1* | 2.64E-02 | 1.736202 |
| ENSMUSG00000009070 | *Rsph14* | 1.51E-06 | 1.732995 |
| ENSMUSG00000038037 | *Socs1* | 3.91E-10 | 1.729735 |
| ENSMUSG00000115783 | *Bc1* | 1.91E-02 | 1.72912 |
| ENSMUSG00000095260 | *Gm25890* | 3.42E-03 | 1.728617 |
| ENSMUSG00000094812 | *Gm22614* | 3.42E-03 | 1.728617 |
| ENSMUSG00000079434 | *Neu2* | 2.66E-03 | 1.727284 |
| ENSMUSG00000066510 | *Ankdd1a* | 6.66E-07 | 1.722526 |
| ENSMUSG00000052565 | *Hist1h1d* | 6.03E-05 | 1.722286 |
| ENSMUSG00000026573 | *Xcl1* | 2.28E-02 | 1.721571 |
| ENSMUSG00000096084 | *Gm21850* | 5.20E-05 | 1.721306 |
| ENSMUSG00000076066 | *Mir223* | 1.46E-04 | 1.719947 |
| ENSMUSG00000060183 | *Cxcl11* | 1.19E-05 | 1.718861 |
| ENSMUSG00000056300 | *Zfp981* | 3.47E-08 | 1.717174 |
| ENSMUSG00000112041 | *9530020I12Rik* | 4.54E-02 | 1.716234 |
| ENSMUSG00000081642 | *Gm13532* | 2.45E-02 | 1.71339 |
| ENSMUSG00000080950 | *Gm7278* | 4.27E-03 | 1.71199 |
| ENSMUSG00000102630 | *Gm37289* | 3.32E-02 | 1.708678 |
| ENSMUSG00000073902 | *Gm1966* | 8.32E-11 | 1.70586 |
| ENSMUSG00000049734 | *Trex1* | 1.10E-13 | 1.705391 |
| ENSMUSG00000060981 | *Hist1h4h* | 1.23E-04 | 1.704353 |
| ENSMUSG00000032174 | *Icam5* | 9.90E-03 | 1.703716 |
| ENSMUSG00000088789 | *Scarna13* | 2.42E-04 | 1.703392 |
| ENSMUSG00000103308 | *Gm37800* | 1.54E-05 | 1.702856 |
| ENSMUSG00000062727 | *Hist1h2bk* | 1.47E-02 | 1.700599 |
| ENSMUSG00000088185 | *Scarna2* | 3.84E-05 | 1.700093 |
| ENSMUSG00000113198 | *Gm6988* | 1.11E-04 | 1.700093 |
| ENSMUSG00000059108 | *Ifitm6* | 1.55E-16 | 1.69831 |
| ENSMUSG00000065862 | *Gm24029* | 5.06E-03 | 1.697729 |
| ENSMUSG00000092805 | *Gm26461* | 8.64E-03 | 1.69729 |
| ENSMUSG00000070271 | *Gm13268* | 2.67E-02 | 1.696212 |
| ENSMUSG00000109052 | *Gm45012* | 1.88E-02 | 1.694509 |
| ENSMUSG00000028037 | *Ifi44* | 9.21E-19 | 1.693727 |
| ENSMUSG00000097413 | *A830052D11Rik* | 3.38E-02 | 1.693497 |
| ENSMUSG00000026358 | *Rgs1* | 1.60E-07 | 1.69318 |
| ENSMUSG00000054325 | *Lce3a* | 4.97E-03 | 1.691831 |
| ENSMUSG00000053560 | *Ier2* | 3.16E-19 | 1.69155 |
| ENSMUSG00000073940 | *Hbb-bt* | 7.55E-04 | 1.691523 |
| ENSMUSG00000051319 | *1500011K16Rik* | 1.19E-02 | 1.691061 |

| | | | |
|---|---|---|---|
| ENSMUSG00000064179 | *Tnnt1* | 3.07E-03 | 1.690918 |
| ENSMUSG00000074115 | *Saa1* | 9.38E-13 | 1.688411 |
| ENSMUSG00000069305 | *Hist1h4n* | 9.18E-03 | 1.688272 |
| ENSMUSG00000028341 | *Nr4a3* | 5.23E-35 | 1.683154 |
| ENSMUSG00000071637 | *Cebpd* | 3.66E-31 | 1.680871 |
| ENSMUSG00000065637 | *Gm26397* | 1.11E-04 | 1.68061 |
| ENSMUSG00000057160 | *Gm16372* | 1.59E-04 | 1.680286 |
| ENSMUSG00000105790 | *Gm24105* | 9.29E-04 | 1.678237 |
| ENSMUSG00000092674 | *Gm24105* | 8.40E-04 | 1.675874 |
| ENSMUSG00000060678 | *Hist1h4c* | 3.44E-05 | 1.674695 |
| ENSMUSG00000042918 | *Mamstr* | 4.07E-16 | 1.672552 |
| ENSMUSG00000109875 | *Gm45456* | 1.06E-02 | 1.6722 |
| ENSMUSG00000030399 | *Ckm* | 1.29E-07 | 1.666114 |
| ENSMUSG00000030672 | *Mylpf* | 3.88E-04 | 1.662639 |
| ENSMUSG00000005716 | *Pvalb* | 2.77E-02 | 1.662341 |
| ENSMUSG00000042031 | *Lce3b* | 7.71E-03 | 1.661896 |
| ENSMUSG00000109905 | *Gm7380* | 1.71E-03 | 1.660339 |
| ENSMUSG00000025172 | *Ankrd2* | 2.03E-04 | 1.659684 |
| ENSMUSG00000026826 | *Nr4a2* | 3.05E-06 | 1.659502 |
| ENSMUSG00000094338 | *Hist1h2bl* | 8.49E-03 | 1.656315 |
| ENSMUSG00000029379 | *Cxcl3* | 2.23E-17 | 1.656114 |
| ENSMUSG00000035852 | *Misp* | 4.46E-09 | 1.656018 |
| ENSMUSG00000086777 | *Far2os2* | 4.90E-02 | 1.655464 |
| ENSMUSG00000053469 | *Tg* | 1.99E-19 | 1.649527 |
| ENSMUSG00000104740 | *Mir6516* | 2.80E-02 | 1.648462 |
| ENSMUSG00000071858 | *Gm94* | 2.34E-02 | 1.647909 |
| ENSMUSG00000031972 | *Acta1* | 6.98E-10 | 1.647593 |
| ENSMUSG00000029635 | *Cdk8* | 1.63E-05 | 1.64378 |
| ENSMUSG00000081684 | *Rps2-ps13* | 1.03E-05 | 1.64243 |
| ENSMUSG00000067767 | *Clec4b2* | 1.61E-03 | 1.642307 |
| ENSMUSG00000045475 | *Lce3c* | 9.18E-03 | 1.642182 |
| ENSMUSG00000017204 | *Gsdma* | 1.46E-02 | 1.640548 |
| ENSMUSG00000029378 | *Areg* | 3.84E-05 | 1.638461 |
| ENSMUSG00000085249 | *Gm12847* | 1.43E-02 | 1.636783 |
| ENSMUSG00000097122 | *Gm26624* | 4.43E-08 | 1.636704 |
| ENSMUSG00000027913 | *Crct1* | 1.49E-03 | 1.63647 |
| ENSMUSG00000050463 | *Krt78* | 8.75E-06 | 1.635311 |
| ENSMUSG00000115518 | *Gm10791* | 1.41E-03 | 1.634944 |
| ENSMUSG00000086213 | *A330040F15Rik* | 1.95E-07 | 1.633556 |
| ENSMUSG00000004885 | *Crabp2* | 2.87E-02 | 1.63166 |
| ENSMUSG00000066491 | *Cox6c2* | 3.99E-07 | 1.630614 |
| ENSMUSG00000062826 | *Ces2f* | 8.39E-04 | 1.627736 |
| ENSMUSG00000046031 | *Calhm6* | 1.04E-03 | 1.627261 |
| ENSMUSG00000064382 | *Gm26447* | 7.12E-04 | 1.626644 |
| ENSMUSG00000035653 | *Lrfn5* | 3.15E-02 | 1.624893 |
| ENSMUSG00000063739 | *Gm4963* | 8.67E-05 | 1.623848 |
| ENSMUSG00000074896 | *Ifit3* | 8.41E-19 | 1.622777 |
| ENSMUSG00000062028 | *Irgc1* | 7.95E-04 | 1.621717 |
| ENSMUSG00000063820 | *Arl9* | 3.59E-02 | 1.621614 |
| ENSMUSG00000045377 | *Tmem88* | 4.72E-06 | 1.619556 |
| ENSMUSG00000114185 | *Gm47662* | 3.98E-02 | 1.619275 |
| ENSMUSG00000041872 | *Il17f* | 1.86E-04 | 1.619247 |
| ENSMUSG00000002500 | *Rpl3l* | 2.59E-03 | 1.616885 |

| | | | |
|---|---|---|---|
| ENSMUSG00000075042 | *4930431P03Rik* | 1.10E-02 | 1.615685 |
| ENSMUSG00000044165 | *Bcl2l15* | 2.68E-05 | 1.615489 |
| ENSMUSG00000017697 | *Ada* | 9.34E-11 | 1.615438 |
| ENSMUSG00000091957 | *Rps2-ps10* | 3.10E-05 | 1.615002 |
| ENSMUSG00000025498 | *Irf7* | 2.69E-23 | 1.614017 |
| ENSMUSG00000114277 | *Gm48583* | 7.46E-04 | 1.611868 |
| ENSMUSG00000104818 | *Gm43661* | 3.64E-03 | 1.609404 |
| ENSMUSG00000063651 | *Cnfn* | 5.49E-04 | 1.609357 |
| ENSMUSG00000081700 | *Atp5k-ps2* | 3.69E-03 | 1.607129 |
| ENSMUSG00000096205 | *Gm22068* | 9.60E-03 | 1.604372 |
| ENSMUSG00000098181 | *Rps12-ps24* | 4.51E-03 | 1.603992 |
| ENSMUSG00000056054 | *S100a8* | 3.40E-07 | 1.600618 |
| ENSMUSG00000028896 | *Rcc1* | 4.47E-11 | 1.599941 |
| ENSMUSG00000031780 | *Ccl17* | 5.39E-06 | 1.598957 |
| ENSMUSG00000109729 | *Gm45418* | 3.30E-05 | 1.597842 |
| ENSMUSG00000029862 | *Clcn1* | 1.22E-04 | 1.595774 |
| ENSMUSG00000097134 | *1110002J07Rik* | 8.36E-04 | 1.595135 |
| ENSMUSG00000094826 | *Gm23804* | 1.22E-03 | 1.594121 |
| ENSMUSG00000074149 | *Gm10634* | 2.88E-07 | 1.59407 |
| ENSMUSG00000029371 | *Cxcl5* | 5.61E-15 | 1.593155 |
| ENSMUSG00000111752 | *Gm38575* | 1.27E-10 | 1.593054 |
| ENSMUSG00000057580 | *Gm10012* | 1.95E-02 | 1.592497 |
| ENSMUSG00000114790 | *4921509O07Rik* | 4.54E-04 | 1.592138 |
| ENSMUSG00000027368 | *Dusp2* | 4.18E-09 | 1.591586 |
| ENSMUSG00000113178 | *Mylf-ps* | 1.12E-06 | 1.591195 |
| ENSMUSG00000048572 | *Tmem252* | 4.44E-31 | 1.589873 |
| ENSMUSG00000095969 | *Rnu1a1* | 1.30E-02 | 1.589137 |
| ENSMUSG00000022602 | *Arc* | 3.85E-02 | 1.589088 |
| ENSMUSG00000105145 | *Gm47299* | 1.40E-02 | 1.587799 |
| ENSMUSG00000105788 | *Gm47295* | 1.40E-02 | 1.587791 |
| ENSMUSG00000020061 | *Mybpc1* | 1.21E-11 | 1.587253 |
| ENSMUSG00000000157 | *Itgb2l* | 7.93E-11 | 1.586842 |
| ENSMUSG00000028680 | *Plk3* | 1.73E-13 | 1.586408 |
| ENSMUSG00000053553 | *3110082I17Rik* | 4.41E-10 | 1.586388 |
| ENSMUSG00000070167 | *Snora57* | 1.06E-03 | 1.585311 |
| ENSMUSG00000024430 | *Cabyr* | 2.76E-02 | 1.584461 |
| ENSMUSG00000087066 | *Gm15518* | 1.60E-08 | 1.584091 |
| ENSMUSG00000026166 | *Ccl20* | 3.82E-11 | 1.583687 |
| ENSMUSG00000014773 | *Dll1* | 1.32E-28 | 1.583278 |
| ENSMUSG00000090582 | *Gm17024* | 6.03E-07 | 1.582913 |
| ENSMUSG00000096659 | *Gm25679* | 1.19E-03 | 1.5829 |
| ENSMUSG00000093815 | *Gm26444* | 1.19E-03 | 1.5829 |
| ENSMUSG00000082838 | *Gm14519* | 5.79E-05 | 1.581682 |
| ENSMUSG00000062488 | *Ifit3b* | 7.88E-13 | 1.57948 |
| ENSMUSG00000070866 | *Zfp804a* | 2.93E-03 | 1.577636 |
| ENSMUSG00000053522 | *Lgals7* | 4.18E-02 | 1.57711 |
| ENSMUSG00000038670 | *Mybpc2* | 9.70E-09 | 1.576178 |
| ENSMUSG00000115355 | *4930445E18Rik* | 1.31E-03 | 1.574801 |
| ENSMUSG00000085434 | *Gm11725* | 3.76E-02 | 1.574487 |
| ENSMUSG00000037977 | *6430571L13Rik* | 1.13E-02 | 1.573903 |
| ENSMUSG00000089281 | *Scarna6* | 8.23E-03 | 1.573558 |
| ENSMUSG00000032327 | *Stra6* | 5.98E-03 | 1.572608 |
| ENSMUSG00000064816 | *Gm22357* | 1.73E-02 | 1.572211 |

| | | | |
|---|---|---|---|
| ENSMUSG00000008384 | *Sertad1* | 1.12E-13 | 1.571184 |
| ENSMUSG00000054582 | *Pabpc1l* | 2.99E-05 | 1.569168 |
| ENSMUSG00000026628 | *Atf3* | 3.62E-22 | 1.569059 |
| ENSMUSG00000057003 | *Myh4* | 1.65E-08 | 1.568031 |
| ENSMUSG00000113722 | *Snhg10* | 1.06E-03 | 1.566332 |
| ENSMUSG00000001014 | *Icam4* | 6.99E-03 | 1.565655 |
| ENSMUSG00000041954 | *Tnfrsf18* | 6.61E-03 | 1.563024 |
| ENSMUSG00000051439 | *Cd14* | 3.56E-27 | 1.562597 |
| ENSMUSG00000040328 | *Olfr56* | 4.75E-15 | 1.562502 |
| ENSMUSG00000096214 | *Gm22634* | 1.28E-03 | 1.562461 |
| ENSMUSG00000096206 | *Gm22317* | 1.28E-03 | 1.562461 |
| ENSMUSG00000033730 | *Egr3* | 2.40E-10 | 1.56174 |
| ENSMUSG00000116961 | *AC117662.4* | 2.23E-03 | 1.560197 |
| ENSMUSG00000090698 | *Apold1* | 1.43E-10 | 1.560015 |
| ENSMUSG00000109700 | *Gm21123* | 2.06E-03 | 1.559279 |
| ENSMUSG00000059741 | *Myl3* | 2.72E-06 | 1.558851 |
| ENSMUSG00000057280 | *Musk* | 2.22E-06 | 1.556841 |
| ENSMUSG00000053508 | *Gtsf2* | 6.80E-07 | 1.553985 |
| ENSMUSG00000030156 | *Cd69* | 5.45E-07 | 1.553384 |
| ENSMUSG00000089617 | *Scarna10* | 1.55E-02 | 1.551375 |
| ENSMUSG00000099587 | *Gm28967* | 1.55E-02 | 1.551375 |
| ENSMUSG00000030669 | *Calca* | 7.44E-05 | 1.551054 |
| ENSMUSG00000078616 | *Trim30c* | 4.08E-08 | 1.550782 |
| ENSMUSG00000060438 | *Rps10-ps1* | 1.39E-08 | 1.550238 |
| ENSMUSG00000024205 | *Rpl36-ps2* | 1.69E-04 | 1.549767 |
| ENSMUSG00000054169 | *Ceacam10* | 1.16E-05 | 1.548901 |
| ENSMUSG00000103313 | *Gm38357* | 3.20E-19 | 1.548815 |
| ENSMUSG00000110697 | *Gm31718* | 2.21E-17 | 1.548758 |
| ENSMUSG00000025726 | *Slc28a1* | 3.17E-02 | 1.545524 |
| ENSMUSG00000009633 | *G0s2* | 6.94E-06 | 1.54482 |
| ENSMUSG00000028435 | *Aqp3* | 1.82E-03 | 1.543177 |
| ENSMUSG00000030592 | *Ryr1* | 4.26E-06 | 1.541763 |
| ENSMUSG00000022126 | *Acod1* | 2.57E-26 | 1.541705 |
| ENSMUSG00000023034 | *Nr4a1* | 1.81E-17 | 1.541288 |
| ENSMUSG00000000982 | *Ccl3* | 6.13E-15 | 1.541237 |
| ENSMUSG00000078763 | *Slfn1* | 5.73E-17 | 1.541102 |
| ENSMUSG00000043953 | *Ccrl2* | 5.73E-17 | 1.540659 |
| ENSMUSG00000021250 | *Fos* | 1.14E-13 | 1.540618 |
| ENSMUSG00000113998 | *Gm40909* | 1.24E-05 | 1.539072 |
| ENSMUSG00000055775 | *Myh8* | 3.34E-07 | 1.538545 |
| ENSMUSG00000113778 | *Gm40847* | 4.05E-02 | 1.537382 |
| ENSMUSG00000003752 | *Itpkc* | 4.85E-21 | 1.537115 |
| ENSMUSG00000039236 | *Isg20* | 3.09E-19 | 1.536846 |
| ENSMUSG00000095891 | *Gm10717* | 4.11E-02 | 1.536618 |
| ENSMUSG00000039001 | *Rps21* | 9.90E-03 | 1.536396 |
| ENSMUSG00000082884 | *Gm13339* | 7.50E-04 | 1.536237 |
| ENSMUSG00000048003 | *Catsper4* | 8.39E-03 | 1.535787 |
| ENSMUSG00000078122 | *F630028O10Rik* | 4.20E-06 | 1.533738 |
| ENSMUSG00000026896 | *Ifih1* | 3.30E-37 | 1.533564 |
| ENSMUSG00000044162 | *Tnip3* | 1.14E-13 | 1.533448 |
| ENSMUSG00000060988 | *Galnt13* | 9.45E-06 | 1.532422 |
| ENSMUSG00000114980 | *4933432I03Rik* | 8.97E-11 | 1.532082 |
| ENSMUSG00000022026 | *Olfm4* | 2.93E-15 | 1.531757 |

| | | | |
|---|---|---|---|
| ENSMUSG00000051627 | *Hist1h1e* | 2.39E-07 | 1.530874 |
| ENSMUSG00000088835 | *Gm23547* | 1.43E-04 | 1.528212 |
| ENSMUSG00000024912 | *Fosl1* | 1.80E-06 | 1.524675 |
| ENSMUSG00000062345 | *Serpinb2* | 8.22E-08 | 1.524046 |
| ENSMUSG00000038418 | *Egr1* | 4.00E-21 | 1.523384 |
| ENSMUSG00000095280 | *Gm21738* | 2.33E-02 | 1.52126 |
| ENSMUSG00000026407 | *Cacna1s* | 4.38E-06 | 1.519984 |
| ENSMUSG00000074146 | *4930579C12Rik* | 7.58E-07 | 1.519858 |
| ENSMUSG00000099250 | *Rn7s2* | 1.74E-02 | 1.51752 |
| ENSMUSG00000099021 | *Rn7s1* | 1.74E-02 | 1.517472 |
| ENSMUSG00000059326 | *Csf2ra* | 1.63E-03 | 1.517321 |
| ENSMUSG00000089999 | *Gm6485* | 5.58E-09 | 1.517 |
| ENSMUSG00000043165 | *Lor* | 8.24E-05 | 1.516516 |
| ENSMUSG00000105195 | *Gm43584* | 5.14E-04 | 1.516166 |
| ENSMUSG00000089929 | *Bcl2a1b* | 2.06E-12 | 1.515751 |
| ENSMUSG00000081857 | *Gm13624* | 3.76E-02 | 1.515199 |
| ENSMUSG00000030787 | *Lyve1* | 1.11E-18 | 1.514832 |
| ENSMUSG00000116848 | *AC138306.1* | 1.12E-11 | 1.514698 |
| ENSMUSG00000104348 | *Gm37691* | 6.21E-04 | 1.514665 |
| ENSMUSG00000066861 | *Oas1g* | 1.81E-08 | 1.514221 |
| ENSMUSG00000020108 | *Ddit4* | 2.93E-14 | 1.513448 |
| ENSMUSG00000089762 | *Ier5l* | 3.07E-03 | 1.513212 |
| ENSMUSG00000095616 | *Gm26244* | 3.81E-02 | 1.513043 |
| ENSMUSG00000098178 | *Gm42418* | 7.46E-03 | 1.512104 |
| ENSMUSG00000081695 | *Rpsa-ps5* | 2.21E-02 | 1.510847 |
| ENSMUSG00000073402 | *Gm8909* | 3.59E-08 | 1.510563 |
| ENSMUSG00000086591 | *Dnah2os* | 1.17E-02 | 1.509414 |
| ENSMUSG00000105398 | *Gm47305* | 1.69E-02 | 1.508695 |
| ENSMUSG00000106279 | *Gm47287* | 1.69E-02 | 1.508691 |
| ENSMUSG00000087700 | *Gm15283* | 2.52E-02 | 1.508122 |
| ENSMUSG00000039070 | *Cpa4* | 3.68E-02 | 1.507901 |
| ENSMUSG00000075122 | *Cd80* | 2.72E-16 | 1.507703 |
| ENSMUSG00000069792 | *Wfdc17* | 2.78E-06 | 1.507287 |
| ENSMUSG00000069372 | *Ctxn3* | 3.67E-03 | 1.507089 |
| ENSMUSG00000087590 | *Epb41l4aos* | 1.11E-02 | 1.506513 |
| ENSMUSG00000026981 | *Il1rn* | 3.16E-19 | 1.505344 |
| ENSMUSG00000026452 | *Syt2* | 1.70E-09 | 1.505325 |
| ENSMUSG00000000204 | *Slfn4* | 4.49E-06 | 1.504687 |
| ENSMUSG00000046961 | *Gpr156* | 2.21E-03 | 1.504602 |
| ENSMUSG00000070692 | *Rps12-ps10* | 4.70E-02 | 1.504592 |
| ENSMUSG00000087881 | *Gm22442* | 1.39E-03 | 1.50459 |
| ENSMUSG00000086290 | *Snhg12* | 4.30E-06 | 1.504012 |
| ENSMUSG00000026121 | *Sema4c* | 6.49E-23 | 1.503955 |
| ENSMUSG00000087006 | *Gm13889* | 6.41E-14 | 1.501606 |
| ENSMUSG00000102496 | *Gm36989* | 8.23E-03 | 1.501532 |
| ENSMUSG00000079597 | *Gm5483* | 3.51E-09 | 1.500636 |
| ENSMUSG00000041117 | *Ccdc8* | 4.06E-05 | 1.500681 |
| ENSMUSG00000058571 | *Gpc6* | 1.32E-14 | 1.500895 |
| ENSMUSG00000036411 | *9530077C05Rik* | 6.19E-05 | 1.501083 |
| ENSMUSG00000052117 | *D630039A03Rik* | 2.45E-02 | 1.502702 |
| ENSMUSG00000104168 | *Gm38250* | 2.20E-02 | 1.503235 |
| ENSMUSG00000075410 | *Prcd* | 1.29E-02 | 1.506303 |
| ENSMUSG00000100005 | *B130024G19Rik* | 2.07E-17 | 1.507802 |

| | | | |
|---|---|---|---|
| ENSMUSG00000075592 | *Nynrin* | 6.90E-13 | 1.509107 |
| ENSMUSG00000038236 | *Hoxa7* | 4.80E-02 | 1.509405 |
| ENSMUSG00000091183 | *Gm5141* | 4.66E-03 | 1.509918 |
| ENSMUSG00000110206 | *Flt3l* | 1.03E-05 | 1.510135 |
| ENSMUSG00000078870 | *Gm14410* | 1.70E-12 | 1.510221 |
| ENSMUSG00000048402 | *Gli2* | 2.01E-16 | 1.510632 |
| ENSMUSG00000027318 | *Adam33* | 7.24E-05 | 1.510692 |
| ENSMUSG00000070867 | *Trabd2b* | 1.19E-23 | 1.510708 |
| ENSMUSG00000002346 | *Slc25a42* | 3.08E-09 | 1.51104 |
| ENSMUSG00000038119 | *Cdon* | 2.13E-13 | 1.511472 |
| ENSMUSG00000000901 | *Mmp11* | 5.59E-04 | 1.511928 |
| ENSMUSG00000030125 | *Lrrc23* | 5.32E-04 | 1.514186 |
| ENSMUSG00000021708 | *Rasgrf2* | 3.43E-12 | 1.514585 |
| ENSMUSG00000030494 | *Rhpn2* | 6.96E-09 | 1.515185 |
| ENSMUSG00000042788 | *Fam166b* | 4.27E-02 | 1.515671 |
| ENSMUSG00000024206 | *Rfx2* | 1.63E-11 | 1.516097 |
| ENSMUSG00000026494 | *Kif26b* | 4.56E-12 | 1.516931 |
| ENSMUSG00000017167 | *Cntnap1* | 8.30E-04 | 1.517416 |
| ENSMUSG00000022456 | *3-Sep* | 6.51E-04 | 1.518584 |
| ENSMUSG00000086158 | *Ccpg1os* | 2.54E-02 | 1.519402 |
| ENSMUSG00000060548 | *Tnfrsf19* | 1.99E-11 | 1.519979 |
| ENSMUSG00000031665 | *Sall1* | 1.66E-02 | 1.520783 |
| ENSMUSG00000020642 | *Rnf144a* | 4.31E-15 | 1.521445 |
| ENSMUSG00000085517 | *Gm12963* | 2.62E-08 | 1.522068 |
| ENSMUSG00000026785 | *Pkn3* | 1.39E-05 | 1.522559 |
| ENSMUSG00000041429 | *Nthl1* | 1.53E-02 | 1.522772 |
| ENSMUSG00000099564 | *Gm28729* | 3.26E-02 | 1.52279 |
| ENSMUSG00000050751 | *Pgbd5* | 3.26E-03 | 1.523024 |
| ENSMUSG00000024063 | *Lbh* | 1.62E-22 | 1.524459 |
| ENSMUSG00000051590 | *Map3k19* | 2.04E-03 | 1.524752 |
| ENSMUSG00000026109 | *Tmeff2* | 7.24E-05 | 1.524936 |
| ENSMUSG00000023266 | *Frs3* | 3.39E-02 | 1.525124 |
| ENSMUSG00000019945 | *Cabcoco1* | 1.24E-04 | 1.52674 |
| ENSMUSG00000026259 | *Ngef* | 2.29E-07 | 1.529308 |
| ENSMUSG00000034522 | *Zfp395* | 1.20E-09 | 1.531411 |
| ENSMUSG00000101801 | *1700020G17Rik* | 1.60E-08 | 1.531815 |
| ENSMUSG00000027656 | *Wisp2* | 1.30E-21 | 1.531853 |
| ENSMUSG00000022199 | *Slc22a17* | 3.25E-05 | 1.532662 |
| ENSMUSG00000045875 | *Adra1a* | 2.84E-04 | 1.533756 |
| ENSMUSG00000072601 | *Ear1* | 4.22E-05 | 1.534311 |
| ENSMUSG00000037492 | *Zmat4* | 1.17E-06 | 1.534582 |
| ENSMUSG00000050541 | *Adra1b* | 9.82E-06 | 1.535536 |
| ENSMUSG00000050014 | *Apol10b* | 1.20E-02 | 1.536867 |
| ENSMUSG00000059824 | *Dbp* | 3.21E-07 | 1.537822 |
| ENSMUSG00000026691 | *Fmo3* | 6.57E-08 | 1.538988 |
| ENSMUSG00000085024 | *C230035I16Rik* | 9.71E-03 | 1.539108 |
| ENSMUSG00000026347 | *Tmem163* | 2.07E-16 | 1.541624 |
| ENSMUSG00000097591 | *A330032B11Rik* | 4.27E-02 | 1.541666 |
| ENSMUSG00000075389 | *2810410L24Rik* | 2.04E-04 | 1.542084 |
| ENSMUSG00000022543 | *4930451G09Rik* | 1.75E-02 | 1.543206 |
| ENSMUSG00000106951 | *5930430L01Rik* | 7.93E-03 | 1.544424 |
| ENSMUSG00000022758 | *P2rx6* | 8.50E-04 | 1.545377 |
| ENSMUSG00000026888 | *Grb14* | 2.00E-26 | 1.54642 |

| | | | |
|---|---|---|---|
| ENSMUSG00000026824 | *Kcnj3* | 9.08E-11 | 1.547942 |
| ENSMUSG00000045555 | *Mettl24* | 1.86E-10 | 1.550112 |
| ENSMUSG00000022053 | *Ebf2* | 8.28E-10 | 1.550167 |
| ENSMUSG00000031489 | *Adrb3* | 2.08E-05 | 1.550335 |
| ENSMUSG00000074384 | *AI429214* | 2.69E-02 | 1.55054 |
| ENSMUSG00000054021 | *Sirt5* | 4.74E-06 | 1.551042 |
| ENSMUSG00000055945 | *Prr18* | 5.17E-03 | 1.551295 |
| ENSMUSG00000033595 | *Lgi3* | 1.84E-11 | 1.55237 |
| ENSMUSG00000103183 | *Gm37090* | 2.52E-03 | 1.5557 |
| ENSMUSG00000061601 | *Pclo* | 1.11E-04 | 1.555835 |
| ENSMUSG00000112714 | *Gm4798* | 2.86E-05 | 1.556475 |
| ENSMUSG00000020182 | *Ddc* | 2.94E-02 | 1.556886 |
| ENSMUSG00000030041 | *M1ap* | 4.17E-06 | 1.556913 |
| ENSMUSG00000066319 | *Rtp3* | 1.02E-03 | 1.559718 |
| ENSMUSG00000013089 | *Etv5* | 2.23E-20 | 1.561183 |
| ENSMUSG00000097805 | *Gm17473* | 3.87E-07 | 1.561625 |
| ENSMUSG00000017309 | *Cd300lg* | 4.88E-06 | 1.561748 |
| ENSMUSG00000085982 | *9530051G07Rik* | 4.48E-02 | 1.563023 |
| ENSMUSG00000029053 | *Prkcz* | 4.17E-13 | 1.563284 |
| ENSMUSG00000030607 | *Acan* | 9.06E-03 | 1.563675 |
| ENSMUSG00000046456 | *Tmem150b* | 1.36E-03 | 1.564973 |
| ENSMUSG00000093479 | *Gm20629* | 2.73E-09 | 1.565239 |
| ENSMUSG00000037610 | *Kcnmb2* | 7.66E-08 | 1.5685 |
| ENSMUSG00000086368 | *Gm13830* | 7.26E-03 | 1.569182 |
| ENSMUSG00000043088 | *Il17re* | 2.31E-04 | 1.569494 |
| ENSMUSG00000016262 | *Sertad4* | 6.83E-03 | 1.56952 |
| ENSMUSG00000020682 | *Mmp28* | 4.74E-10 | 1.570816 |
| ENSMUSG00000084939 | *Gm830* | 1.84E-02 | 1.572365 |
| ENSMUSG00000039763 | *Dnajc28* | 3.97E-02 | 1.575323 |
| ENSMUSG00000020734 | *Grin2c* | 3.03E-02 | 1.577359 |
| ENSMUSG00000026676 | *Ccdc3* | 6.28E-11 | 1.578234 |
| ENSMUSG00000028871 | *Rspo1* | 1.42E-02 | 1.579214 |
| ENSMUSG00000017007 | *Rbpjl* | 8.03E-08 | 1.581836 |
| ENSMUSG00000019831 | *Wasf1* | 2.22E-02 | 1.583533 |
| ENSMUSG00000042793 | *Lgr6* | 1.16E-16 | 1.585253 |
| ENSMUSG00000087424 | *5730405O15Rik* | 2.20E-02 | 1.587279 |
| ENSMUSG00000078794 | *Dact3* | 2.02E-05 | 1.590225 |
| ENSMUSG00000095079 | *Igha* | 8.37E-05 | 1.592326 |
| ENSMUSG00000050022 | *Amz1* | 5.27E-05 | 1.593171 |
| ENSMUSG00000036854 | *Hspb6* | 2.04E-03 | 1.597689 |
| ENSMUSG00000021719 | *Rgs7bp* | 2.95E-09 | 1.59776 |
| ENSMUSG00000025902 | *Sox17* | 1.02E-13 | 1.598591 |
| ENSMUSG00000074219 | *Gm10644* | 2.18E-02 | 1.598961 |
| ENSMUSG00000037846 | *Rtkn2* | 5.55E-07 | 1.598991 |
| ENSMUSG00000102082 | *Gm28675* | 4.81E-02 | 1.599203 |
| ENSMUSG00000086141 | *9030622O22Rik* | 1.25E-10 | 1.604845 |
| ENSMUSG00000089941 | *Gm16168* | 1.79E-03 | 1.606211 |
| ENSMUSG00000112468 | *Gm47720* | 2.62E-06 | 1.608526 |
| ENSMUSG00000068323 | *Slc4a5* | 6.46E-08 | 1.609607 |
| ENSMUSG00000116097 | *Gm36738* | 2.28E-02 | 1.611492 |
| ENSMUSG00000019312 | *Grb7* | 8.16E-04 | 1.611956 |
| ENSMUSG00000059659 | *Gm10069* | 3.68E-05 | 1.612558 |
| ENSMUSG00000031220 | *Awat2* | 1.41E-02 | 1.613621 |

| | | | |
|---|---|---|---|
| ENSMUSG00000090206 | *Tepp* | 4.13E-03 | 1.616264 |
| ENSMUSG00000087514 | *Gm16076* | 2.99E-02 | 1.62071 |
| ENSMUSG00000041351 | *Rap1gap* | 2.36E-09 | 1.621611 |
| ENSMUSG00000069378 | *Prdm6* | 1.85E-18 | 1.623444 |
| ENSMUSG00000036923 | *Stox1* | 5.18E-07 | 1.623817 |
| ENSMUSG00000050288 | *Fzd2* | 8.28E-04 | 1.624159 |
| ENSMUSG00000086607 | *4930511M06Rik* | 6.52E-03 | 1.624741 |
| ENSMUSG00000034685 | *Fam171a2* | 6.79E-04 | 1.62484 |
| ENSMUSG00000062542 | *Syt9* | 1.57E-02 | 1.627124 |
| ENSMUSG00000007279 | *Scube2* | 2.67E-18 | 1.629611 |
| ENSMUSG00000054057 | *A930004D18Rik* | 3.30E-02 | 1.629982 |
| ENSMUSG00000066438 | *Plekhd1* | 2.77E-07 | 1.631307 |
| ENSMUSG00000085682 | *Gm14267* | 1.18E-03 | 1.635757 |
| ENSMUSG00000074793 | *Hspa12b* | 6.74E-21 | 1.635809 |
| ENSMUSG00000030495 | *Slc7a10* | 3.89E-09 | 1.637942 |
| ENSMUSG00000033590 | *Myo5c* | 1.66E-15 | 1.641777 |
| ENSMUSG00000030562 | *Nox4* | 8.94E-09 | 1.642642 |
| ENSMUSG00000083817 | *Gm14400* | 1.77E-02 | 1.642954 |
| ENSMUSG00000099553 | *Gm29538* | 1.68E-03 | 1.64385 |
| ENSMUSG00000058420 | *Syt17* | 3.26E-06 | 1.643854 |
| ENSMUSG00000054459 | *Vsnl1* | 1.48E-12 | 1.644077 |
| ENSMUSG00000037973 | *Ccdc129* | 8.09E-09 | 1.644828 |
| ENSMUSG00000059146 | *Ntrk3* | 4.47E-22 | 1.645531 |
| ENSMUSG00000056155 | *Nanos3* | 4.06E-03 | 1.646041 |
| ENSMUSG00000106749 | *Gm43281* | 3.37E-06 | 1.64753 |
| ENSMUSG00000072599 | *Ear-ps2* | 8.09E-03 | 1.648472 |
| ENSMUSG00000069518 | *Gm10271* | 4.49E-02 | 1.649982 |
| ENSMUSG00000074771 | *Ankef1* | 3.41E-04 | 1.656591 |
| ENSMUSG00000041479 | *Syt15* | 1.37E-05 | 1.659181 |
| ENSMUSG00000049511 | *Htr1b* | 2.21E-04 | 1.662514 |
| ENSMUSG00000063430 | *Wscd2* | 1.63E-07 | 1.662763 |
| ENSMUSG00000086596 | *Susd5* | 3.88E-05 | 1.663486 |
| ENSMUSG00000011263 | *Exoc3l2* | 2.11E-02 | 1.671039 |
| ENSMUSG00000034584 | *Exph5* | 2.07E-17 | 1.671345 |
| ENSMUSG00000022840 | *Adcy5* | 1.15E-24 | 1.673312 |
| ENSMUSG00000028801 | *Stpg1* | 1.90E-04 | 1.675936 |
| ENSMUSG00000027827 | *Kcnab1* | 1.06E-12 | 1.685414 |
| ENSMUSG00000020614 | *Fam20a* | 7.24E-19 | 1.686889 |
| ENSMUSG00000021499 | *Catsper3* | 7.33E-03 | 1.687729 |
| ENSMUSG00000085171 | *D830026I12Rik* | 4.25E-02 | 1.689753 |
| ENSMUSG00000035435 | *Abca17* | 2.81E-10 | 1.692557 |
| ENSMUSG00000030043 | *Tacr1* | 7.60E-05 | 1.693777 |
| ENSMUSG00000041020 | *Map7d2* | 2.75E-02 | 1.695372 |
| ENSMUSG00000110380 | *Gm45332* | 6.76E-03 | 1.695675 |
| ENSMUSG00000105376 | *Gm36535* | 1.64E-02 | 1.696479 |
| ENSMUSG00000101605 | *Ace3* | 1.74E-05 | 1.696685 |
| ENSMUSG00000037709 | *Fam13a* | 2.63E-08 | 1.704891 |
| ENSMUSG00000109165 | *Gm45148* | 5.67E-03 | 1.705895 |
| ENSMUSG00000020169 | *Best3* | 1.46E-07 | 1.71572 |
| ENSMUSG00000048728 | *Zfp454* | 4.49E-02 | 1.717216 |
| ENSMUSG00000028755 | *Cda* | 1.99E-02 | 1.717754 |
| ENSMUSG00000085282 | *Gm15663* | 1.21E-02 | 1.718236 |
| ENSMUSG00000048096 | *Lmod1* | 9.53E-07 | 1.718553 |

| | | | |
|---|---|---|---|
| ENSMUSG00000103738 | *Gm37652* | 1.24E-03 | 1.722127 |
| ENSMUSG00000108580 | *Gm39094* | 1.47E-22 | 1.733461 |
| ENSMUSG00000027380 | *Acoxl* | 2.48E-08 | 1.733573 |
| ENSMUSG00000063535 | *Zfp773* | 3.78E-03 | 1.735884 |
| ENSMUSG00000027186 | *Elf5* | 3.75E-05 | 1.74138 |
| ENSMUSG00000029193 | *Cckar* | 1.05E-05 | 1.742803 |
| ENSMUSG00000108767 | *Gm29763* | 2.99E-02 | 1.742917 |
| ENSMUSG00000032357 | *Tinag* | 2.33E-06 | 1.748176 |
| ENSMUSG00000092515 | *C87198* | 4.36E-02 | 1.750226 |
| ENSMUSG00000070661 | *Rnf186* | 4.40E-02 | 1.752275 |
| ENSMUSG00000030380 | *Mzf1* | 1.63E-02 | 1.763149 |
| ENSMUSG00000042428 | *Mgat3* | 6.40E-12 | 1.763723 |
| ENSMUSG00000044708 | *Kcnj10* | 2.22E-02 | 1.767881 |
| ENSMUSG00000041423 | *Paqr6* | 2.72E-02 | 1.768799 |
| ENSMUSG00000089712 | *Gm15889* | 6.62E-03 | 1.770295 |
| ENSMUSG00000063730 | *Hsd3b2* | 2.08E-02 | 1.773821 |
| ENSMUSG00000101693 | *Gm19461* | 2.09E-02 | 1.777561 |
| ENSMUSG00000064294 | *Aox3* | 3.77E-15 | 1.778141 |
| ENSMUSG00000086313 | *Gm15940* | 2.10E-02 | 1.780429 |
| ENSMUSG00000044694 | *2010007H06Rik* | 1.51E-06 | 1.781054 |
| ENSMUSG00000108481 | *Gm33248* | 4.21E-13 | 1.784046 |
| ENSMUSG00000061397 | *Krt79* | 6.09E-04 | 1.786124 |
| ENSMUSG00000094786 | *Gm14403* | 1.13E-16 | 1.786856 |
| ENSMUSG00000079436 | *Kcnj13* | 4.22E-02 | 1.78992 |
| ENSMUSG00000038233 | *Fam198a* | 1.05E-06 | 1.790914 |
| ENSMUSG00000042256 | *Ptchd4* | 7.00E-05 | 1.791307 |
| ENSMUSG00000052026 | *Slc6a7* | 4.86E-05 | 1.792733 |
| ENSMUSG00000097124 | *A530020G20Rik* | 7.37E-06 | 1.792876 |
| ENSMUSG00000048988 | *Elfn1* | 1.27E-13 | 1.803612 |
| ENSMUSG00000057719 | *Sh3rf2* | 2.42E-19 | 1.804366 |
| ENSMUSG00000111761 | *9230112J17Rik* | 3.13E-04 | 1.810487 |
| ENSMUSG00000027886 | *1700013F07Rik* | 2.43E-02 | 1.810795 |
| ENSMUSG00000082292 | *Gm12250* | 3.69E-06 | 1.81246 |
| ENSMUSG00000049107 | *Ntf3* | 7.06E-20 | 1.821726 |
| ENSMUSG00000068373 | *D430041D05Rik* | 5.87E-17 | 1.830741 |
| ENSMUSG00000020566 | *Atp6v1c2* | 1.23E-25 | 1.832003 |
| ENSMUSG00000114028 | *9630002D21Rik* | 3.96E-03 | 1.832354 |
| ENSMUSG00000102590 | *Mannr* | 1.75E-04 | 1.836166 |
| ENSMUSG00000006567 | *Atp7b* | 2.18E-17 | 1.836476 |
| ENSMUSG00000076613 | *Ighg2b* | 2.30E-02 | 1.837406 |
| ENSMUSG00000102437 | *Gm38048* | 2.54E-08 | 1.843504 |
| ENSMUSG00000025577 | *Cbx2* | 1.42E-04 | 1.846281 |
| ENSMUSG00000037279 | *Ovol2* | 2.30E-03 | 1.852727 |
| ENSMUSG00000030093 | *Wnt7a* | 1.05E-06 | 1.855136 |
| ENSMUSG00000097453 | *Gm26894* | 1.59E-03 | 1.856702 |
| ENSMUSG00000087518 | *Gm13561* | 3.22E-03 | 1.859076 |
| ENSMUSG00000044633 | *B530045E10Rik* | 8.05E-19 | 1.860022 |
| ENSMUSG00000009394 | *Syn2* | 1.55E-17 | 1.865441 |
| ENSMUSG00000085069 | *Gm13111* | 1.65E-04 | 1.866473 |
| ENSMUSG00000079173 | *Zan* | 4.11E-04 | 1.868008 |
| ENSMUSG00000030098 | *Grip2* | 4.24E-11 | 1.871182 |
| ENSMUSG00000074575 | *Kcng1* | 4.71E-02 | 1.871584 |
| ENSMUSG00000116766 | *AC171205.1* | 3.48E-02 | 1.87324 |

| | | | |
|---|---|---|---|
| ENSMUSG00000027513 | *Pck1* | 3.53E-10 | 1.874273 |
| ENSMUSG00000056824 | *Zfp663* | 6.72E-03 | 1.880195 |
| ENSMUSG00000086843 | *E030013I19Rik* | 3.55E-05 | 1.886677 |
| ENSMUSG00000108218 | *Olfr1372-ps1* | 6.73E-09 | 1.893798 |
| ENSMUSG00000087413 | *Gm11266* | 3.96E-14 | 1.899133 |
| ENSMUSG00000093550 | *Higd1c* | 3.35E-02 | 1.906324 |
| ENSMUSG00000020335 | *Zfp354b* | 1.86E-02 | 1.906344 |
| ENSMUSG00000027460 | *Angpt4* | 7.61E-09 | 1.918001 |
| ENSMUSG00000036295 | *Lrrn3* | 4.53E-05 | 1.925128 |
| ENSMUSG00000090166 | *Ear10* | 1.40E-06 | 1.929094 |
| ENSMUSG00000099338 | *2810030D12Rik* | 6.44E-05 | 1.935947 |
| ENSMUSG00000032502 | *Stac* | 6.19E-06 | 1.936727 |
| ENSMUSG00000079039 | *Gm11037* | 2.50E-02 | 1.940577 |
| ENSMUSG00000105786 | *Gm18957* | 1.48E-02 | 1.942241 |
| ENSMUSG00000117322 | *AC154766.4* | 2.55E-02 | 1.953694 |
| ENSMUSG00000022504 | *Ciita* | 2.96E-28 | 1.959704 |
| ENSMUSG00000028919 | *Arhgef19* | 2.26E-03 | 1.968027 |
| ENSMUSG00000110891 | *Gm40639* | 3.60E-02 | 1.972536 |
| ENSMUSG00000104376 | *Gm37516* | 1.20E-02 | 1.974628 |
| ENSMUSG00000029601 | *Iqcd* | 8.65E-03 | 1.990538 |
| ENSMUSG00000026870 | *Cutal* | 1.79E-04 | 1.994395 |
| ENSMUSG00000110631 | *Gm42047* | 5.59E-03 | 2.007803 |
| ENSMUSG00000085882 | *2610507I01Rik* | 5.32E-05 | 2.01961 |
| ENSMUSG00000109770 | *Gm30085* | 3.97E-02 | 2.026925 |
| ENSMUSG00000086390 | *1810019D21Rik* | 1.01E-09 | 2.035809 |
| ENSMUSG00000080797 | *Gm15760* | 2.06E-02 | 2.049068 |
| ENSMUSG00000033849 | *B3galt2* | 1.86E-03 | 2.059198 |
| ENSMUSG00000031725 | *Ces1f* | 6.83E-03 | 2.092553 |
| ENSMUSG00000087500 | *Gm12426* | 5.91E-03 | 2.107058 |
| ENSMUSG00000092004 | *Gm17482* | 9.93E-07 | 2.112466 |
| ENSMUSG00000019909 | *Fam162b* | 5.07E-04 | 2.116376 |
| ENSMUSG00000115307 | *Gm49313* | 1.84E-04 | 2.139641 |
| ENSMUSG00000060275 | *Nrg2* | 7.68E-08 | 2.145009 |
| ENSMUSG00000025789 | *St8sia2* | 1.28E-15 | 2.169809 |
| ENSMUSG00000108390 | *Gm39038* | 1.64E-03 | 2.192189 |
| ENSMUSG00000026147 | *Col9a1* | 1.56E-02 | 2.194131 |
| ENSMUSG00000053961 | *Ang5* | 2.31E-03 | 2.198318 |
| ENSMUSG00000086155 | *9430041J12Rik* | 2.78E-02 | 2.203234 |
| ENSMUSG00000027796 | *Smad9* | 3.65E-14 | 2.220367 |
| ENSMUSG00000103160 | *C130012C08Rik* | 2.01E-03 | 2.234457 |
| ENSMUSG00000020081 | *Tacr2* | 1.82E-02 | 2.253652 |
| ENSMUSG00000069830 | *Nlrp1a* | 2.69E-07 | 2.281738 |
| ENSMUSG00000108617 | *Gm31749* | 6.04E-03 | 2.285686 |
| ENSMUSG00000009900 | *Wnt3a* | 5.16E-09 | 2.298134 |
| ENSMUSG00000027831 | *Veph1* | 9.72E-25 | 2.301924 |
| ENSMUSG00000061171 | *Slc38a11* | 3.50E-05 | 2.353128 |
| ENSMUSG00000001494 | *Sost* | 4.34E-04 | 2.412139 |
| ENSMUSG00000026304 | *Rab17* | 1.09E-03 | 2.608575 |
| ENSMUSG00000031551 | *Ido1* | 1.39E-05 | 3.239545 |
| ENSMUSG00000092591 | *Gm20429* | 2.09E-04 | 3.383042 |
| ENSMUSG00000071679 | *Rtl4* | 6.39E-21 | 4.582634 |
| ENSMUSG00000085154 | *C130046K22Rik* | 1.28E-13 | 27.59869 |

**Supplementary Table 8.** Murine differential gene expression in the lungs 6 h post-infection, 9-47-Ear vs 4559-Blood. Genes with fold change (FC) greater than 1.5 and $p < 0.05$ are shown. FC values highlighted in blue = upregulated in 9-47-Ear, while values highlighted in red = upregulated in 4559-Blood.

| Gene Stable ID | Gene Name | padj | FC |
|---|---|---|---|
| ENSMUSG00000096349 | *Gm22513* | 3.77E-18 | 10.97464 |
| ENSMUSG00000094668 | *Gm24871* | 1.29E-07 | 6.745915 |
| ENSMUSG00000064899 | *Snord118* | 1.08E-09 | 6.570577 |
| ENSMUSG00000097554 | *Gm26825* | 7.25E-04 | 5.791676 |
| ENSMUSG00000109641 | *Gm45556* | 1.47E-03 | 5.598475 |
| ENSMUSG00000105361 | *AY036118* | 3.15E-13 | 5.461151 |
| ENSMUSG00000080538 | *Gm25541* | 4.06E-18 | 5.174635 |
| ENSMUSG00000093843 | *Gm25939* | 7.29E-13 | 4.925819 |
| ENSMUSG00000095676 | *Gm25099* | 9.26E-27 | 4.906139 |
| ENSMUSG00000065870 | *Rnu3a* | 1.74E-16 | 4.884372 |
| ENSMUSG00000106147 | *Rnu3a* | 1.74E-16 | 4.884372 |
| ENSMUSG00000095738 | *Gm25313* | 8.71E-14 | 4.857934 |
| ENSMUSG00000095892 | *Rnu5g* | 4.27E-14 | 4.78872 |
| ENSMUSG00000104896 | *Rnu3b4* | 9.34E-24 | 4.622038 |
| ENSMUSG00000098641 | *Rnu3b4* | 1.21E-23 | 4.615922 |
| ENSMUSG00000099291 | *Rnu3b3* | 6.68E-23 | 4.590292 |
| ENSMUSG00000104856 | *Rnu3b3* | 6.68E-23 | 4.590292 |
| ENSMUSG00000105115 | *Rnu3b2* | 7.99E-24 | 4.578094 |
| ENSMUSG00000098925 | *Rnu3b2* | 7.99E-24 | 4.578094 |
| ENSMUSG00000098943 | *Rnu3b1* | 3.21E-23 | 4.559078 |
| ENSMUSG00000105025 | *Rnu3b1* | 3.21E-23 | 4.559078 |
| ENSMUSG00000085393 | *Gm11280* | 1.04E-03 | 4.538932 |
| ENSMUSG00000064945 | *Rny3* | 1.19E-15 | 4.476071 |
| ENSMUSG00000042385 | *Gzmk* | 5.37E-03 | 4.35977 |
| ENSMUSG00000094050 | *Gm23472* | 1.36E-13 | 4.320851 |
| ENSMUSG00000093956 | *Gm24497* | 4.43E-13 | 4.291089 |
| ENSMUSG00000064856 | *Gm23444* | 2.34E-13 | 4.288303 |
| ENSMUSG00000088675 | *Rprl1* | 2.41E-06 | 4.192998 |
| ENSMUSG00000106222 | *Rprl1* | 2.41E-06 | 4.192998 |
| ENSMUSG00000065176 | *Rnu12* | 1.99E-27 | 4.109879 |
| ENSMUSG00000065701 | *Rny1* | 7.64E-13 | 4.069417 |
| ENSMUSG00000065254 | *Gm23973* | 3.03E-06 | 4.059083 |
| ENSMUSG00000032494 | *Tdgf1* | 1.92E-05 | 4.053171 |
| ENSMUSG00000065820 | *Gm26316* | 9.85E-13 | 4.048075 |
| ENSMUSG00000064682 | *Gm25813* | 9.90E-09 | 4.046294 |
| ENSMUSG00000064702 | *Gm24950* | 3.05E-12 | 3.981116 |
| ENSMUSG00000065251 | *Gm23971* | 3.05E-12 | 3.981116 |
| ENSMUSG00000065944 | *Rnu2-10* | 2.22E-12 | 3.976161 |
| ENSMUSG00000065767 | *Gm23849* | 3.07E-12 | 3.962822 |
| ENSMUSG00000077323 | *Rnu11* | 6.04E-13 | 3.962105 |
| ENSMUSG00000094377 | *Gm24407* | 2.62E-14 | 3.808851 |
| ENSMUSG00000065232 | *Gm22973* | 6.93E-12 | 3.795942 |
| ENSMUSG00000067455 | *Hist1h4j* | 3.60E-08 | 3.778868 |
| ENSMUSG00000060093 | *Hist1h4a* | 2.31E-16 | 3.753049 |
| ENSMUSG00000096243 | *Gm24265* | 2.84E-11 | 3.648484 |

| | | | |
|---|---|---|---|
| ENSMUSG00000065845 | *Gm25189* | 2.58E-03 | 3.636942 |
| ENSMUSG00000064943 | *Gm23240* | 2.39E-03 | 3.62003 |
| ENSMUSG00000069300 | *Hist1h2bj* | 2.08E-05 | 3.616983 |
| ENSMUSG00000095616 | *Gm26244* | 1.81E-09 | 3.585974 |
| ENSMUSG00000088252 | *Snord13* | 3.16E-08 | 3.500728 |
| ENSMUSG00000064941 | *Gm23238* | 2.49E-12 | 3.477513 |
| ENSMUSG00000095969 | *Rnu1a1* | 4.33E-10 | 3.447212 |
| ENSMUSG00000096659 | *Gm25679* | 2.63E-16 | 3.411828 |
| ENSMUSG00000093815 | *Gm26444* | 2.63E-16 | 3.411828 |
| ENSMUSG00000063021 | *Hist1h2ak* | 2.45E-03 | 3.406041 |
| ENSMUSG00000095260 | *Gm25890* | 9.28E-10 | 3.34811 |
| ENSMUSG00000094812 | *Gm22614* | 9.28E-10 | 3.34811 |
| ENSMUSG00000064923 | *Gm22042* | 8.01E-10 | 3.347755 |
| ENSMUSG00000095580 | *Rnu1b1* | 2.34E-09 | 3.347511 |
| ENSMUSG00000093834 | *Rnu1b2* | 2.34E-09 | 3.347511 |
| ENSMUSG00000094306 | *Gm24924* | 7.72E-09 | 3.330976 |
| ENSMUSG00000095701 | *Gm24830* | 1.01E-09 | 3.296712 |
| ENSMUSG00000096838 | *Gm26232* | 1.01E-09 | 3.296712 |
| ENSMUSG00000065773 | *Rnu1b6* | 1.01E-09 | 3.296712 |
| ENSMUSG00000094826 | *Gm23804* | 3.33E-15 | 3.281135 |
| ENSMUSG00000097131 | *D230017M19Rik* | 7.32E-15 | 3.266318 |
| ENSMUSG00000069266 | *Hist1h4b* | 3.25E-06 | 3.259907 |
| ENSMUSG00000060981 | *Hist1h4h* | 4.37E-17 | 3.254606 |
| ENSMUSG00000101355 | *Hist1h3h* | 1.42E-03 | 3.179868 |
| ENSMUSG00000099517 | *Hist1h3g* | 1.06E-03 | 3.159837 |
| ENSMUSG00000010362 | *Rdm1* | 1.06E-11 | 3.14031 |
| ENSMUSG00000081684 | *Rps2-ps13* | 4.23E-24 | 3.13385 |
| ENSMUSG00000069305 | *Hist1h4n* | 4.43E-08 | 3.133044 |
| ENSMUSG00000065905 | *Gm26110* | 1.16E-04 | 3.130131 |
| ENSMUSG00000069306 | *Hist1h4m* | 5.27E-07 | 3.127821 |
| ENSMUSG00000095217 | *Hist1h2bn* | 2.95E-06 | 3.119423 |
| ENSMUSG00000064288 | *Hist1h4k* | 1.87E-08 | 3.117272 |
| ENSMUSG00000080465 | *Gm22486* | 2.90E-09 | 3.11249 |
| ENSMUSG00000047061 | *Gm9817* | 3.14E-02 | 3.110749 |
| ENSMUSG00000087963 | *Gm25394* | 2.00E-03 | 3.095042 |
| ENSMUSG00000115420 | *AL732506.1* | 2.92E-15 | 3.080505 |
| ENSMUSG00000088088 | *Rmrp* | 2.92E-15 | 3.080505 |
| ENSMUSG00000096205 | *Gm22068* | 3.77E-09 | 3.06075 |
| ENSMUSG00000094655 | *Gm25360* | 1.37E-14 | 3.058749 |
| ENSMUSG00000060678 | *Hist1h4c* | 2.01E-18 | 3.058247 |
| ENSMUSG00000065824 | *Gm26315* | 1.04E-11 | 3.053521 |
| ENSMUSG00000092837 | *Rpph1* | 4.58E-09 | 3.031887 |
| ENSMUSG00000088025 | *Rprl3* | 2.03E-07 | 3.027158 |
| ENSMUSG00000099583 | *Hist1h3d* | 3.33E-04 | 3.01984 |
| ENSMUSG00000071470 | *Ccnb1ip1* | 4.41E-09 | 3.011385 |
| ENSMUSG00000065911 | *Gm24447* | 2.34E-11 | 2.998729 |
| ENSMUSG00000064694 | *Gm24146* | 5.78E-07 | 2.99441 |
| ENSMUSG00000092805 | *Gm26461* | 4.04E-09 | 2.980919 |
| ENSMUSG00000069274 | *Hist1h4f* | 2.64E-07 | 2.963294 |
| ENSMUSG00000096214 | *Gm22634* | 3.07E-14 | 2.955618 |
| ENSMUSG00000096206 | *Gm22317* | 3.07E-14 | 2.955618 |
| ENSMUSG00000087775 | *Rprl2* | 5.59E-07 | 2.949971 |
| ENSMUSG00000061482 | *Hist1h4d* | 5.96E-25 | 2.947909 |

| | | | |
|---|---|---|---|
| ENSMUSG00000092819 | *Gm23639* | 1.31E-03 | 2.911866 |
| ENSMUSG00000023467 | *Tulp2* | 8.88E-03 | 2.887985 |
| ENSMUSG00000010342 | *Tex14* | 2.11E-12 | 2.887178 |
| ENSMUSG00000092702 | *Gm24514* | 3.30E-08 | 2.857933 |
| ENSMUSG00000105790 | *Gm24105* | 1.12E-10 | 2.857786 |
| ENSMUSG00000087943 | *Gm24245* | 4.70E-13 | 2.854799 |
| ENSMUSG00000100210 | *Hist1h3f* | 1.45E-07 | 2.846879 |
| ENSMUSG00000090625 | *Gm20721* | 5.27E-04 | 2.822061 |
| ENSMUSG00000092674 | *Gm24105* | 1.19E-10 | 2.820054 |
| ENSMUSG00000046242 | *Nme9* | 4.34E-05 | 2.796067 |
| ENSMUSG00000064604 | *Snora44* | 7.00E-06 | 2.781139 |
| ENSMUSG00000114279 | *Hist1h2bm* | 1.67E-03 | 2.777077 |
| ENSMUSG00000058385 | *Hist1h2bg* | 2.78E-09 | 2.77039 |
| ENSMUSG00000064901 | *Snora21* | 9.99E-04 | 2.75448 |
| ENSMUSG00000101972 | *Hist1h3i* | 1.08E-03 | 2.745319 |
| ENSMUSG00000070167 | *Snora57* | 2.32E-12 | 2.741946 |
| ENSMUSG00000069265 | *Hist1h3a* | 1.82E-03 | 2.720348 |
| ENSMUSG00000064451 | *Snora23* | 9.44E-04 | 2.713527 |
| ENSMUSG00000091383 | *Hist1h2al* | 5.55E-14 | 2.672474 |
| ENSMUSG00000114456 | *Hist1h2bh* | 2.26E-05 | 2.669249 |
| ENSMUSG00000068855 | *Hist2h2ac* | 1.06E-06 | 2.662418 |
| ENSMUSG00000088008 | *Gm25492* | 6.86E-05 | 2.660852 |
| ENSMUSG00000105145 | *Gm47299* | 1.14E-08 | 2.652374 |
| ENSMUSG00000105788 | *Gm47295* | 1.14E-08 | 2.652269 |
| ENSMUSG00000105399 | *Mir1843b* | 7.66E-03 | 2.649552 |
| ENSMUSG00000065686 | *Snora5c* | 9.51E-04 | 2.647534 |
| ENSMUSG00000062727 | *Hist1h2bk* | 1.21E-05 | 2.647047 |
| ENSMUSG00000090208 | *Gm15851* | 4.93E-02 | 2.646 |
| ENSMUSG00000096992 | *Gm26788* | 1.53E-02 | 2.640965 |
| ENSMUSG00000092386 | *Gm20536* | 1.21E-02 | 2.615154 |
| ENSMUSG00000057596 | *Trim30d* | 2.24E-12 | 2.613284 |
| ENSMUSG00000064999 | *Gm26035* | 7.18E-09 | 2.608373 |
| ENSMUSG00000069267 | *Hist1h3b* | 1.04E-03 | 2.564833 |
| ENSMUSG00000093355 | *Snora26* | 1.95E-02 | 2.562115 |
| ENSMUSG00000069273 | *Hist1h3e* | 5.25E-08 | 2.556766 |
| ENSMUSG00000064994 | *Gm22422* | 9.17E-05 | 2.54989 |
| ENSMUSG00000060988 | *Galnt13* | 1.31E-22 | 2.547254 |
| ENSMUSG00000081342 | *Gm11434* | 1.83E-02 | 2.528216 |
| ENSMUSG00000065649 | *Snora74a* | 1.80E-07 | 2.526072 |
| ENSMUSG00000064655 | *Gm25788* | 1.04E-03 | 2.525134 |
| ENSMUSG00000097799 | *Gm26899* | 2.35E-02 | 2.488561 |
| ENSMUSG00000065145 | *Vaultrc5* | 2.35E-06 | 2.476147 |
| ENSMUSG00000085465 | *Gm15347* | 1.42E-02 | 2.4729 |
| ENSMUSG00000075031 | *Hist1h2bb* | 2.05E-04 | 2.467707 |
| ENSMUSG00000087819 | *Gm25117* | 1.17E-09 | 2.467012 |
| ENSMUSG00000074403 | *Hist2h3b* | 1.52E-03 | 2.454729 |
| ENSMUSG00000077254 | *Gm26079* | 1.78E-05 | 2.438855 |
| ENSMUSG00000115783 | *Bc1* | 1.63E-04 | 2.432863 |
| ENSMUSG00000098178 | *Gm42418* | 3.40E-10 | 2.426788 |
| ENSMUSG00000065087 | *Snord22* | 4.68E-20 | 2.423107 |
| ENSMUSG00000094338 | *Hist1h2bl* | 5.96E-06 | 2.421661 |
| ENSMUSG00000096010 | *Hist4h4* | 1.32E-07 | 2.421203 |
| ENSMUSG00000076614 | *Ighg1* | 3.86E-02 | 2.417765 |

| | | | |
|---|---|---|---|
| ENSMUSG00000089536 | *Scarna3a* | 8.57E-03 | 2.411926 |
| ENSMUSG00000069268 | *Hist1h2bf* | 6.11E-03 | 2.41015 |
| ENSMUSG00000076258 | *Gm23935* | 6.42E-13 | 2.406898 |
| ENSMUSG00000097346 | *Gm26619* | 2.89E-02 | 2.388088 |
| ENSMUSG00000080365 | *Gm25776* | 3.68E-05 | 2.383555 |
| ENSMUSG00000076281 | *Gm24270* | 1.12E-10 | 2.373892 |
| ENSMUSG00000097815 | *Gm26809* | 2.33E-02 | 2.371395 |
| ENSMUSG00000114969 | *Hba-ps3* | 9.14E-04 | 2.369228 |
| ENSMUSG00000065778 | *Gm22154* | 1.68E-02 | 2.34964 |
| ENSMUSG00000105448 | *Gm25820* | 1.08E-02 | 2.348821 |
| ENSMUSG00000105288 | *Gm25820* | 1.08E-02 | 2.348821 |
| ENSMUSG00000064966 | *Snord15b* | 1.44E-06 | 2.333139 |
| ENSMUSG00000069308 | *Hist1h2bp* | 3.62E-03 | 2.322081 |
| ENSMUSG00000035299 | *Mid1* | 4.01E-07 | 2.316383 |
| ENSMUSG00000085790 | *Gm12729* | 2.53E-02 | 2.316346 |
| ENSMUSG00000088529 | *Gm26083* | 2.80E-05 | 2.314116 |
| ENSMUSG00000113512 | *Gm9063* | 3.83E-02 | 2.306271 |
| ENSMUSG00000069310 | *Hist1h3c* | 2.34E-03 | 2.302294 |
| ENSMUSG00000115432 | *D130009I18Rik* | 2.70E-03 | 2.300679 |
| ENSMUSG00000004814 | *Ccl24* | 9.80E-03 | 2.290894 |
| ENSMUSG00000091957 | *Rps2-ps10* | 1.37E-12 | 2.275718 |
| ENSMUSG00000077611 | *Gm23946* | 4.84E-02 | 2.264088 |
| ENSMUSG00000088609 | *Gm24187* | 3.72E-12 | 2.233571 |
| ENSMUSG00000099250 | *Rn7s2* | 5.82E-07 | 2.228017 |
| ENSMUSG00000099021 | *Rn7s1* | 5.82E-07 | 2.227961 |
| ENSMUSG00000095590 | *Gm24305* | 2.92E-02 | 2.221176 |
| ENSMUSG00000060639 | *Hist1h4i* | 5.98E-05 | 2.218385 |
| ENSMUSG00000082755 | *Gm8692* | 3.31E-04 | 2.214971 |
| ENSMUSG00000112246 | *Gm40761* | 4.75E-02 | 2.182331 |
| ENSMUSG00000105398 | *Gm47305* | 8.24E-07 | 2.175963 |
| ENSMUSG00000106279 | *Gm47287* | 8.24E-07 | 2.175954 |
| ENSMUSG00000063954 | *Hist2h2aa2* | 3.09E-07 | 2.164586 |
| ENSMUSG00000052305 | *Hbb-bs* | 1.32E-07 | 2.16149 |
| ENSMUSG00000104863 | *Gm49347* | 2.14E-04 | 2.145847 |
| ENSMUSG00000064380 | *Gm26448* | 4.40E-09 | 2.145839 |
| ENSMUSG00000088246 | *Gm25911* | 7.67E-12 | 2.145126 |
| ENSMUSG00000089255 | *Snora78* | 1.36E-02 | 2.137569 |
| ENSMUSG00000085297 | *Gm11651* | 5.25E-03 | 2.127159 |
| ENSMUSG00000058773 | *Hist1h1b* | 8.95E-05 | 2.120617 |
| ENSMUSG00000064220 | *Hist2h2aa1* | 3.10E-04 | 2.117751 |
| ENSMUSG00000104740 | *Mir6516* | 9.92E-04 | 2.113466 |
| ENSMUSG00000046229 | *Scand1* | 1.44E-02 | 2.111608 |
| ENSMUSG00000064387 | *Snora73a* | 7.47E-12 | 2.088608 |
| ENSMUSG00000085261 | *Gm13814* | 1.41E-02 | 2.077594 |
| ENSMUSG00000080542 | *Gm22710* | 2.43E-03 | 2.074884 |
| ENSMUSG00000091993 | *B930036N10Rik* | 2.07E-03 | 2.07144 |
| ENSMUSG00000112831 | *AC167229.1* | 2.82E-02 | 2.068995 |
| ENSMUSG00000091405 | *Hist2h4* | 9.28E-10 | 2.052495 |
| ENSMUSG00000046031 | *Calhm6* | 1.74E-06 | 2.048611 |
| ENSMUSG00000084708 | *Gm22988* | 5.06E-06 | 2.038794 |
| ENSMUSG00000031289 | *Il13ra2* | 4.46E-05 | 2.025167 |
| ENSMUSG00000088185 | *Scarna2* | 2.45E-08 | 2.024717 |
| ENSMUSG00000029635 | *Cdk8* | 3.25E-10 | 2.022129 |

| | | | |
|---|---|---|---|
| ENSMUSG00000088789 | *Scarna13* | 1.73E-06 | 2.017749 |
| ENSMUSG00000026573 | *Xcl1* | 3.45E-03 | 2.016054 |
| ENSMUSG00000100992 | *Gm23925* | 3.22E-02 | 2.01348 |
| ENSMUSG00000102189 | *Gm37194* | 5.60E-06 | 2.000652 |
| ENSMUSG00000077167 | *Gm24119* | 9.88E-03 | 1.997643 |
| ENSMUSG00000065637 | *Gm26397* | 4.01E-07 | 1.997637 |
| ENSMUSG00000084744 | *Gm25291* | 4.03E-03 | 1.99723 |
| ENSMUSG00000106649 | *Gm49349* | 1.43E-06 | 1.993359 |
| ENSMUSG00000064405 | *Gm23925* | 3.66E-02 | 1.98704 |
| ENSMUSG00000082516 | *Gm13342* | 1.02E-02 | 1.981125 |
| ENSMUSG00000027233 | *Patl2* | 3.08E-02 | 1.979645 |
| ENSMUSG00000092746 | *Rn7s6* | 1.82E-05 | 1.965498 |
| ENSMUSG00000073940 | *Hbb-bt* | 8.75E-06 | 1.965051 |
| ENSMUSG00000103336 | *Gm37993* | 4.98E-02 | 1.954663 |
| ENSMUSG00000100831 | *Gm17847* | 1.84E-02 | 1.954622 |
| ENSMUSG00000081058 | *Hist2h3c2* | 2.98E-03 | 1.954493 |
| ENSMUSG00000065118 | *Gm23297* | 1.62E-02 | 1.944291 |
| ENSMUSG00000030670 | *Cyp2r1* | 1.52E-02 | 1.939325 |
| ENSMUSG00000107722 | *2900060B14Rik* | 3.88E-03 | 1.935981 |
| ENSMUSG00000001983 | *Taco1* | 6.95E-08 | 1.933656 |
| ENSMUSG00000065226 | *Gm25791* | 3.01E-06 | 1.926078 |
| ENSMUSG00000062461 | *Gm5453* | 8.46E-10 | 1.924129 |
| ENSMUSG00000104690 | *Gm47304* | 3.05E-05 | 1.920177 |
| ENSMUSG00000065037 | *Rn7sk* | 2.04E-05 | 1.917124 |
| ENSMUSG00000069917 | *Hba-a2* | 3.52E-13 | 1.915594 |
| ENSMUSG00000055170 | *Ifng* | 5.97E-03 | 1.914606 |
| ENSMUSG00000064841 | *Gm26205* | 1.80E-02 | 1.910409 |
| ENSMUSG00000090101 | *Snhg9* | 2.49E-02 | 1.888943 |
| ENSMUSG00000065353 | *Snora73b* | 9.17E-05 | 1.888933 |
| ENSMUSG00000113136 | *Gm19951* | 1.01E-03 | 1.888218 |
| ENSMUSG00000113465 | *Gm40271* | 5.45E-03 | 1.872958 |
| ENSMUSG00000066629 | *Rpl36-ps3* | 5.45E-03 | 1.872958 |
| ENSMUSG00000024205 | *Rpl36-ps2* | 1.48E-07 | 1.856283 |
| ENSMUSG00000089617 | *Scarna10* | 3.70E-04 | 1.856044 |
| ENSMUSG00000099587 | *Gm28967* | 3.70E-04 | 1.856044 |
| ENSMUSG00000022658 | *Tagln3* | 3.59E-03 | 1.852149 |
| ENSMUSG00000117074 | *AC121299.1* | 4.22E-02 | 1.849145 |
| ENSMUSG00000087968 | *Gm25395* | 7.59E-06 | 1.841096 |
| ENSMUSG00000052565 | *Hist1h1d* | 8.37E-06 | 1.839687 |
| ENSMUSG00000091955 | *Gm9844* | 3.52E-18 | 1.837563 |
| ENSMUSG00000086196 | *Gm13571* | 3.25E-04 | 1.836858 |
| ENSMUSG00000064853 | *Gm23442* | 2.97E-04 | 1.832979 |
| ENSMUSG00000077714 | *Snord17* | 3.52E-04 | 1.822824 |
| ENSMUSG00000069919 | *Hba-a1* | 8.16E-13 | 1.822782 |
| ENSMUSG00000020641 | *Rsad2* | 3.37E-12 | 1.822486 |
| ENSMUSG00000049539 | *Hist1h1a* | 7.38E-03 | 1.821165 |
| ENSMUSG00000069307 | *Hist1h2bq* | 4.15E-05 | 1.818275 |
| ENSMUSG00000035528 | *Npffr2* | 7.77E-03 | 1.814241 |
| ENSMUSG00000027220 | *Syt13* | 1.11E-02 | 1.812055 |
| ENSMUSG00000067321 | *Gm7931* | 3.36E-02 | 1.81009 |
| ENSMUSG00000085348 | *Myhas* | 3.50E-03 | 1.803894 |
| ENSMUSG00000065822 | *Snord15a* | 1.55E-02 | 1.803286 |
| ENSMUSG00000063234 | *Gpr84* | 1.21E-08 | 1.800134 |

| | | | |
|---|---|---|---|
| ENSMUSG00000085241 | *Snhg3* | 6.58E-07 | 1.798898 |
| ENSMUSG00000085573 | *Gm15418* | 4.34E-02 | 1.79681 |
| ENSMUSG00000065304 | *Gm23245* | 2.53E-02 | 1.794865 |
| ENSMUSG00000003545 | *Fosb* | 2.44E-06 | 1.793447 |
| ENSMUSG00000089762 | *Ier5l* | 3.00E-05 | 1.79236 |
| ENSMUSG00000057465 | *Saa2* | 5.73E-09 | 1.789508 |
| ENSMUSG00000053553 | *3110082I17Rik* | 9.75E-15 | 1.769663 |
| ENSMUSG00000109875 | *Gm45456* | 4.92E-03 | 1.768914 |
| ENSMUSG00000069303 | *Hist1h2br* | 8.01E-05 | 1.766683 |
| ENSMUSG00000100969 | *1700030N03Rik* | 1.27E-04 | 1.764586 |
| ENSMUSG00000053626 | *Tll1* | 3.72E-29 | 1.754673 |
| ENSMUSG00000037190 | *Cyb561d2* | 9.84E-03 | 1.753648 |
| ENSMUSG00000046470 | *Sox18* | 1.23E-13 | 1.751676 |
| ENSMUSG00000044092 | *C130050O18Rik* | 1.22E-06 | 1.748316 |
| ENSMUSG00000062778 | *Chia1* | 4.57E-20 | 1.747009 |
| ENSMUSG00000035852 | *Misp* | 1.80E-10 | 1.74205 |
| ENSMUSG00000025946 | *Pth2r* | 3.71E-02 | 1.730767 |
| ENSMUSG00000113722 | *Snhg10* | 6.72E-05 | 1.728473 |
| ENSMUSG00000020638 | *Cmpk2* | 9.44E-27 | 1.721661 |
| ENSMUSG00000089281 | *Scarna6* | 1.62E-03 | 1.712381 |
| ENSMUSG00000077563 | *Snora68* | 2.00E-03 | 1.708197 |
| ENSMUSG00000114234 | *Gm3131* | 3.41E-02 | 1.706159 |
| ENSMUSG00000045932 | *Ifit2* | 2.48E-21 | 1.705861 |
| ENSMUSG00000028461 | *Ccdc107* | 1.38E-06 | 1.705554 |
| ENSMUSG00000102349 | *Gm37376* | 1.71E-02 | 1.701801 |
| ENSMUSG00000024810 | *Il33* | 4.65E-23 | 1.70107 |
| ENSMUSG00000070473 | *Cldn3* | 2.42E-10 | 1.696455 |
| ENSMUSG00000048709 | *Gm8666* | 3.20E-02 | 1.694683 |
| ENSMUSG00000028678 | *Kif2c* | 3.69E-02 | 1.693279 |
| ENSMUSG00000095098 | *Ccdc85b* | 4.36E-03 | 1.690404 |
| ENSMUSG00000074576 | *Mocs3* | 1.14E-02 | 1.689861 |
| ENSMUSG00000088948 | *Gm23262* | 2.79E-03 | 1.686819 |
| ENSMUSG00000075268 | *Gm10819* | 1.05E-04 | 1.686108 |
| ENSMUSG00000078141 | *Gm2399* | 4.75E-03 | 1.685731 |
| ENSMUSG00000022595 | *Lypd2* | 4.45E-03 | 1.685708 |
| ENSMUSG00000063171 | *Rps4l* | 2.87E-03 | 1.683014 |
| ENSMUSG00000110528 | *Gm1943* | 4.81E-02 | 1.681924 |
| ENSMUSG00000015880 | *Ncapg* | 1.05E-02 | 1.677303 |
| ENSMUSG00000056758 | *Hmga2* | 3.94E-02 | 1.676199 |
| ENSMUSG00000084783 | *Gm15419* | 2.22E-02 | 1.674643 |
| ENSMUSG00000116694 | *AC154667.2* | 1.48E-03 | 1.671371 |
| ENSMUSG00000083737 | *Prdx6-ps2* | 1.28E-02 | 1.666943 |
| ENSMUSG00000085431 | *4930440I19Rik* | 1.34E-03 | 1.66478 |
| ENSMUSG00000097971 | *Gm26917* | 1.21E-04 | 1.66206 |
| ENSMUSG00000094344 | *Gm11942* | 4.89E-10 | 1.661048 |
| ENSMUSG00000092267 | *Gm20417* | 6.05E-04 | 1.651296 |
| ENSMUSG00000042737 | *Dpm3* | 1.76E-02 | 1.647963 |
| ENSMUSG00000087247 | *Alkal1* | 1.04E-10 | 1.64795 |
| ENSMUSG00000064382 | *Gm26447* | 5.42E-04 | 1.647264 |
| ENSMUSG00000083337 | *Gm11539* | 4.30E-05 | 1.643919 |
| ENSMUSG00000025226 | *Fbxl15* | 3.44E-02 | 1.64349 |
| ENSMUSG00000082241 | *Gng5-ps* | 3.89E-02 | 1.641179 |
| ENSMUSG00000105827 | *Hist2h2bb* | 2.20E-03 | 1.637237 |

| | | | |
|---|---|---|---|
| ENSMUSG00000057160 | *Gm16372* | 4.55E-04 | 1.635812 |
| ENSMUSG00000106767 | *Gm42727* | 1.77E-02 | 1.635709 |
| ENSMUSG00000020493 | *Prr11* | 1.30E-02 | 1.633632 |
| ENSMUSG00000092116 | *Gm10320* | 3.42E-03 | 1.631539 |
| ENSMUSG00000117037 | *CT010433.2* | 2.53E-02 | 1.628923 |
| ENSMUSG00000024401 | *Tnf* | 3.17E-19 | 1.627752 |
| ENSMUSG00000057863 | *Rpl36* | 9.70E-06 | 1.626087 |
| ENSMUSG00000047370 | *Gm7367* | 9.82E-04 | 1.625595 |
| ENSMUSG00000020895 | *Tmem107* | 4.41E-02 | 1.624311 |
| ENSMUSG00000114304 | *Gm48099* | 1.30E-06 | 1.623936 |
| ENSMUSG00000028896 | *Rcc1* | 1.44E-11 | 1.622009 |
| ENSMUSG00000107349 | *Gm6655* | 8.45E-03 | 1.614486 |
| ENSMUSG00000080006 | *Rps19-ps7* | 3.14E-03 | 1.611362 |
| ENSMUSG00000060438 | *Rps10-ps1* | 9.88E-10 | 1.610232 |
| ENSMUSG00000080811 | *Gm14513* | 7.39E-05 | 1.609441 |
| ENSMUSG00000104862 | *Gm6520* | 2.83E-02 | 1.607007 |
| ENSMUSG00000028540 | *Dph2* | 7.43E-05 | 1.606169 |
| ENSMUSG00000022769 | *Sdf2l1* | 7.63E-04 | 1.604934 |
| ENSMUSG00000090942 | *F830016B08Rik* | 7.69E-06 | 1.603028 |
| ENSMUSG00000007892 | *Rplp1* | 2.35E-11 | 1.602848 |
| ENSMUSG00000022097 | *Sftpc* | 3.79E-08 | 1.601389 |
| ENSMUSG00000040936 | *Ulk4* | 1.79E-13 | 1.600562 |
| ENSMUSG00000097060 | *Gm26759* | 2.92E-04 | 1.600487 |
| ENSMUSG00000091845 | *Rpl36-ps12* | 6.25E-04 | 1.600167 |
| ENSMUSG00000030329 | *Pianp* | 8.93E-03 | 1.596449 |
| ENSMUSG00000035202 | *Lars2* | 9.76E-08 | 1.596336 |
| ENSMUSG00000104325 | *Gm37321* | 1.08E-02 | 1.595466 |
| ENSMUSG00000082662 | *Gm15516* | 4.17E-02 | 1.594564 |
| ENSMUSG00000063586 | *Gm5513* | 6.64E-03 | 1.592966 |
| ENSMUSG00000097679 | *Rps19-ps5* | 1.49E-02 | 1.590111 |
| ENSMUSG00000108414 | *Snhg1* | 2.93E-11 | 1.588096 |
| ENSMUSG00000085401 | *Slc39a1-ps* | 5.90E-10 | 1.587829 |
| ENSMUSG00000060183 | *Cxcl11* | 2.63E-04 | 1.587297 |
| ENSMUSG00000027447 | *Cst3* | 8.64E-19 | 1.585854 |
| ENSMUSG00000069515 | *Lyz1* | 3.15E-15 | 1.584793 |
| ENSMUSG00000071419 | *Rps15-ps2* | 2.45E-08 | 1.583911 |
| ENSMUSG00000102135 | *Gm37108* | 5.48E-03 | 1.582546 |
| ENSMUSG00000065126 | *Snord104* | 8.14E-04 | 1.581336 |
| ENSMUSG00000052837 | *Junb* | 4.96E-18 | 1.580061 |
| ENSMUSG00000117187 | *CT033749.1* | 1.02E-02 | 1.579981 |
| ENSMUSG00000000386 | *Mx1* | 2.32E-17 | 1.579725 |
| ENSMUSG00000036114 | *Rpp25l* | 1.62E-03 | 1.578918 |
| ENSMUSG00000079505 | *Gm11131* | 3.63E-03 | 1.576721 |
| ENSMUSG00000114321 | *Gm8971* | 3.66E-04 | 1.576165 |
| ENSMUSG00000046240 | *Hepacam* | 5.51E-04 | 1.575183 |
| ENSMUSG00000083669 | *Gm10169* | 5.08E-03 | 1.573737 |
| ENSMUSG00000094955 | *Gm3699* | 5.08E-03 | 1.573303 |
| ENSMUSG00000029410 | *Ppef2* | 5.25E-03 | 1.572807 |
| ENSMUSG00000068606 | *Gm4841* | 3.38E-02 | 1.572259 |
| ENSMUSG00000106352 | *5033403H07Rik* | 4.74E-06 | 1.571858 |
| ENSMUSG00000098534 | *Gm27167* | 3.43E-06 | 1.571239 |
| ENSMUSG00000111897 | *Gm19810* | 1.29E-04 | 1.568539 |
| ENSMUSG00000117269 | *AC151275.1* | 3.31E-02 | 1.567532 |

| | | | |
|---|---|---|---|
| ENSMUSG00000112290 | *Gm33782* | 4.90E-03 | 1.563678 |
| ENSMUSG00000015437 | *Gzmb* | 5.18E-03 | 1.561612 |
| ENSMUSG00000024165 | *Jpt2* | 9.07E-16 | 1.560746 |
| ENSMUSG00000047443 | *Erfe* | 1.12E-03 | 1.559959 |
| ENSMUSG00000089542 | *Gm25835* | 9.96E-04 | 1.558194 |
| ENSMUSG00000080747 | *Gm14016* | 4.05E-03 | 1.557656 |
| ENSMUSG00000064063 | *BC048507* | 1.80E-02 | 1.554995 |
| ENSMUSG00000117046 | *AC122746.1* | 2.71E-02 | 1.552568 |
| ENSMUSG00000021070 | *Bdkrb2* | 5.92E-05 | 1.552425 |
| ENSMUSG00000060419 | *Rps16-ps2* | 4.73E-10 | 1.551902 |
| ENSMUSG00000086324 | *Gm15564* | 3.09E-05 | 1.551082 |
| ENSMUSG00000086420 | *Gm8865* | 4.69E-03 | 1.548083 |
| ENSMUSG00000041378 | *Cldn5* | 1.39E-15 | 1.546994 |
| ENSMUSG00000027496 | *Aurka* | 4.17E-02 | 1.546663 |
| ENSMUSG00000095478 | *Gm9824* | 9.03E-03 | 1.546072 |
| ENSMUSG00000079330 | *Lemd1* | 1.45E-03 | 1.544978 |
| ENSMUSG00000076036 | *Gm22133* | 4.18E-13 | 1.544169 |
| ENSMUSG00000082062 | *Ftl2-ps* | 7.57E-11 | 1.542066 |
| ENSMUSG00000032425 | *Zfp949* | 7.47E-12 | 1.541584 |
| ENSMUSG00000070284 | *Gmppb* | 4.20E-05 | 1.539971 |
| ENSMUSG00000081382 | *Rpl18-ps1* | 1.43E-03 | 1.538547 |
| ENSMUSG00000099060 | *Mir7025* | 4.48E-02 | 1.53617 |
| ENSMUSG00000082609 | *Gm15464* | 3.50E-02 | 1.535836 |
| ENSMUSG00000030744 | *Rps3* | 6.04E-11 | 1.535727 |
| ENSMUSG00000037169 | *Mycn* | 3.45E-04 | 1.535438 |
| ENSMUSG00000094111 | *Nutf2-ps2* | 4.80E-02 | 1.534662 |
| ENSMUSG00000087006 | *Gm13889* | 3.61E-15 | 1.533387 |
| ENSMUSG00000113212 | *Gm47435* | 1.57E-02 | 1.532655 |
| ENSMUSG00000007872 | *Id3* | 6.85E-12 | 1.530174 |
| ENSMUSG00000015092 | *Edf1* | 3.80E-08 | 1.530159 |
| ENSMUSG00000072235 | *Tuba1a* | 2.93E-13 | 1.528993 |
| ENSMUSG00000073144 | *4930599N23Rik* | 1.03E-03 | 1.528627 |
| ENSMUSG00000087060 | *Eldr* | 9.87E-03 | 1.527571 |
| ENSMUSG00000028179 | *Cth* | 1.62E-04 | 1.52673 |
| ENSMUSG00000058900 | *Rsl1* | 9.35E-05 | 1.526622 |
| ENSMUSG00000053508 | *Gtsf2* | 3.10E-06 | 1.522863 |
| ENSMUSG00000050796 | *B3galt6* | 1.56E-02 | 1.522466 |
| ENSMUSG00000050370 | *Ch25h* | 2.02E-21 | 1.521802 |
| ENSMUSG00000059835 | *Rpl13-ps3* | 7.89E-04 | 1.521415 |
| ENSMUSG00000112003 | *Gm4864* | 3.60E-02 | 1.520838 |
| ENSMUSG00000043099 | *Hic1* | 1.04E-03 | 1.518516 |
| ENSMUSG00000033368 | *Trim69* | 4.03E-03 | 1.516764 |
| ENSMUSG00000046731 | *Kctd11* | 1.05E-14 | 1.516698 |
| ENSMUSG00000070392 | *Gm20634* | 2.49E-03 | 1.515844 |
| ENSMUSG00000093674 | *Rpl41* | 1.91E-12 | 1.515683 |
| ENSMUSG00000096942 | *Rps19-ps6* | 1.09E-06 | 1.515244 |
| ENSMUSG00000104560 | *Gm8115* | 7.31E-03 | 1.514801 |
| ENSMUSG00000070462 | *Tlnrd1* | 1.52E-09 | 1.513754 |
| ENSMUSG00000036992 | *Nxt1* | 1.64E-03 | 1.513731 |
| ENSMUSG00000061848 | *Gm5805* | 1.54E-10 | 1.512703 |
| ENSMUSG00000099544 | *Gm18699* | 2.13E-03 | 1.511865 |
| ENSMUSG00000059434 | *Gckr* | 1.64E-03 | 1.511175 |
| ENSMUSG00000042842 | *Serpinb6b* | 2.39E-16 | 1.510922 |

| | | | |
|---|---|---|---|
| ENSMUSG00000068240 | *Gm11808* | 9.62E-10 | 1.510582 |
| ENSMUSG00000109658 | *Gm45660* | 3.74E-05 | 1.510374 |
| ENSMUSG00000044197 | *Gpr146* | 8.83E-19 | 1.510371 |
| ENSMUSG00000105472 | *Gm43024* | 2.61E-02 | 1.509917 |
| ENSMUSG00000075020 | *Mir670hg* | 8.54E-03 | 1.509891 |
| ENSMUSG00000008140 | *Emc10* | 3.81E-09 | 1.509274 |
| ENSMUSG00000029371 | *Cxcl5* | 1.04E-11 | 1.507055 |
| ENSMUSG00000024925 | *Rnaseh2c* | 3.97E-02 | 1.506518 |
| ENSMUSG00000110275 | *Gm5905* | 2.92E-15 | 1.506379 |
| ENSMUSG00000115143 | *Fkbp1a-ps1* | 3.40E-04 | 1.504907 |
| ENSMUSG00000074115 | *Saa1* | 4.30E-08 | 1.504478 |
| ENSMUSG00000091639 | *Gm3756* | 5.43E-10 | 1.504454 |
| ENSMUSG00000101449 | *Gm17791* | 1.23E-02 | 1.504151 |
| ENSMUSG00000098181 | *Rps12-ps24* | 1.77E-02 | 1.503082 |
| ENSMUSG00000116358 | *Gm49450* | 2.31E-13 | 1.502451 |
| ENSMUSG00000059461 | *Gm7331* | 1.06E-06 | 1.502048 |
| ENSMUSG00000091269 | *Gm6682* | 9.38E-11 | 1.501853 |
| ENSMUSG00000022408 | Fam83f | 1.13E-02 | 1.500479 |
| ENSMUSG00000027843 | Ptpn22 | 4.55E-11 | 1.500715 |
| ENSMUSG00000026414 | Tnnt2 | 2.59E-14 | 1.500887 |
| ENSMUSG00000046961 | Gpr156 | 7.92E-04 | 1.503034 |
| ENSMUSG00000030468 | Siglecg | 1.69E-05 | 1.503144 |
| ENSMUSG00000059900 | Tmem40 | 3.18E-02 | 1.503359 |
| ENSMUSG00000015568 | Lpl | 1.57E-32 | 1.503627 |
| ENSMUSG00000048794 | Cfap100 | 1.97E-05 | 1.504035 |
| ENSMUSG00000043789 | Vwce | 1.10E-02 | 1.504138 |
| ENSMUSG00000021751 | Acox2 | 2.40E-04 | 1.504181 |
| ENSMUSG00000050600 | Zfp831 | 4.21E-09 | 1.504376 |
| ENSMUSG00000020892 | Aloxe3 | 1.83E-02 | 1.504792 |
| ENSMUSG00000078234 | Klhdc7a | 7.85E-04 | 1.505376 |
| ENSMUSG00000114937 | Gm30054 | 1.87E-05 | 1.505389 |
| ENSMUSG00000046186 | Cd109 | 2.33E-05 | 1.505728 |
| ENSMUSG00000038167 | Plekhg6 | 2.51E-03 | 1.506016 |
| ENSMUSG00000028017 | Egf | 4.01E-07 | 1.506454 |
| ENSMUSG00000019577 | Pdk4 | 8.43E-28 | 1.506585 |
| ENSMUSG00000025271 | Pfkfb1 | 7.66E-13 | 1.507316 |
| ENSMUSG00000029123 | Stk32b | 1.69E-02 | 1.507913 |
| ENSMUSG00000024049 | Myom1 | 1.17E-21 | 1.508449 |
| ENSMUSG00000007122 | Casq1 | 6.70E-07 | 1.509955 |
| ENSMUSG00000003476 | Crhr2 | 1.35E-04 | 1.511069 |
| ENSMUSG00000056366 | Fabp3-ps1 | 2.81E-06 | 1.512819 |
| ENSMUSG00000108955 | Gm44775 | 3.93E-02 | 1.513353 |
| ENSMUSG00000111907 | Gm48202 | 6.46E-03 | 1.514782 |
| ENSMUSG00000002007 | Srpk3 | 3.83E-03 | 1.514899 |
| ENSMUSG00000006345 | Ggt1 | 1.00E-04 | 1.514981 |
| ENSMUSG00000073968 | Trim68 | 1.66E-08 | 1.515157 |
| ENSMUSG00000112230 | Ifngas1 | 3.52E-02 | 1.515642 |
| ENSMUSG00000090124 | Ugt1a7c | 1.52E-11 | 1.516108 |
| ENSMUSG00000089943 | Ugt1a5 | 7.03E-06 | 1.516142 |
| ENSMUSG00000090165 | Ugt1a10 | 6.59E-12 | 1.516306 |
| ENSMUSG00000097579 | Gm26799 | 2.23E-03 | 1.517356 |
| ENSMUSG00000112023 | Lilr4b | 1.77E-04 | 1.517729 |
| ENSMUSG00000005951 | Shpk | 1.11E-04 | 1.51779 |

| | | | |
|---|---|---|---|
| ENSMUSG00000035296 | Sgcg | 1.16E-09 | 1.517805 |
| ENSMUSG00000032925 | Itgbl1 | 1.02E-17 | 1.518245 |
| ENSMUSG00000043873 | Chil5 | 7.73E-03 | 1.518859 |
| ENSMUSG00000109371 | Gm44626 | 2.32E-02 | 1.519236 |
| ENSMUSG00000090175 | Ugt1a9 | 7.00E-12 | 1.519384 |
| ENSMUSG00000060275 | Nrg2 | 6.87E-03 | 1.52007 |
| ENSMUSG00000089675 | Ugt1a8 | 6.80E-12 | 1.520697 |
| ENSMUSG00000041608 | Entpd3 | 1.12E-10 | 1.520924 |
| ENSMUSG00000069808 | Fam57a | 1.25E-06 | 1.521054 |
| ENSMUSG00000100510 | AV026068 | 5.30E-03 | 1.521178 |
| ENSMUSG00000086607 | 4930511M06Rik | 2.52E-02 | 1.521734 |
| ENSMUSG00000040247 | Tbc1d10c | 6.96E-07 | 1.522149 |
| ENSMUSG00000041642 | Kif21b | 3.24E-10 | 1.522398 |
| ENSMUSG00000090799 | Klhl33 | 8.11E-05 | 1.522621 |
| ENSMUSG00000070498 | Tmem132b | 3.43E-08 | 1.523157 |
| ENSMUSG00000018796 | Acsl1 | 8.24E-30 | 1.523655 |
| ENSMUSG00000078937 | Cpt1b | 5.04E-08 | 1.523826 |
| ENSMUSG00000034570 | Inpp5j | 8.55E-03 | 1.523892 |
| ENSMUSG00000044551 | 9930012K11Rik | 7.95E-04 | 1.524065 |
| ENSMUSG00000032053 | Pou2af1 | 6.57E-06 | 1.524753 |
| ENSMUSG00000089804 | Gm16136 | 3.66E-02 | 1.525379 |
| ENSMUSG00000031958 | Ldhd | 1.14E-03 | 1.526 |
| ENSMUSG00000073988 | Ttpa | 2.43E-06 | 1.52693 |
| ENSMUSG00000043639 | Rbm20 | 4.76E-16 | 1.527208 |
| ENSMUSG00000046971 | Pla2g4f | 1.67E-05 | 1.527252 |
| ENSMUSG00000058207 | Serpina3k | 5.08E-04 | 1.5276 |
| ENSMUSG00000062257 | Opcml | 5.34E-11 | 1.527897 |
| ENSMUSG00000066108 | Muc5b | 2.76E-21 | 1.528023 |
| ENSMUSG00000020256 | Aldh1l2 | 1.82E-04 | 1.530481 |
| ENSMUSG00000031303 | Map3k15 | 1.02E-08 | 1.532052 |
| ENSMUSG00000006014 | Prg4 | 3.62E-08 | 1.532224 |
| ENSMUSG00000007107 | Atp1a4 | 2.62E-06 | 1.53242 |
| ENSMUSG00000038777 | Sema6c | 8.09E-05 | 1.533598 |
| ENSMUSG00000030470 | Csrp3 | 6.81E-10 | 1.534077 |
| ENSMUSG00000024039 | Cbs | 3.49E-02 | 1.534538 |
| ENSMUSG00000000686 | Abhd15 | 8.63E-05 | 1.535291 |
| ENSMUSG00000051910 | Sox6 | 9.77E-26 | 1.536789 |
| ENSMUSG00000002997 | Prkar2b | 2.00E-17 | 1.537187 |
| ENSMUSG00000024803 | Ankrd1 | 1.13E-12 | 1.537284 |
| ENSMUSG00000110397 | Gm45540 | 4.65E-06 | 1.537736 |
| ENSMUSG00000029228 | Lnx1 | 1.73E-19 | 1.537867 |
| ENSMUSG00000031382 | Asb11 | 2.82E-04 | 1.538155 |
| ENSMUSG00000020303 | Stc2 | 3.41E-05 | 1.538946 |
| ENSMUSG00000026824 | Kcnj3 | 2.07E-10 | 1.539467 |
| ENSMUSG00000054545 | Ugt1a6a | 9.67E-10 | 1.539483 |
| ENSMUSG00000097193 | Gm26664 | 1.10E-02 | 1.539627 |
| ENSMUSG00000032066 | Bco2 | 3.66E-02 | 1.539965 |
| ENSMUSG00000090145 | Ugt1a6b | 6.45E-12 | 1.540465 |
| ENSMUSG00000030098 | Grip2 | 2.01E-05 | 1.541304 |
| ENSMUSG00000030650 | Tmc5 | 3.57E-04 | 1.543931 |
| ENSMUSG00000113165 | Gm47863 | 4.70E-02 | 1.547216 |
| ENSMUSG00000096981 | Gm16845 | 7.49E-05 | 1.547297 |
| ENSMUSG00000058498 | Rnf207 | 5.23E-03 | 1.547335 |

| | | | |
|---|---|---|---|
| ENSMUSG00000104094 | Gm37314 | 4.58E-03 | 1.547567 |
| ENSMUSG00000087172 | Gm12153 | 2.27E-09 | 1.547691 |
| ENSMUSG00000000197 | Nalcn | 4.06E-06 | 1.549339 |
| ENSMUSG00000097885 | 5031434O11Rik | 4.83E-03 | 1.549511 |
| ENSMUSG00000085779 | Atcayos | 8.31E-10 | 1.550724 |
| ENSMUSG00000038583 | Pln | 3.75E-10 | 1.550962 |
| ENSMUSG00000047940 | Stpg2 | 3.85E-03 | 1.55183 |
| ENSMUSG00000032495 | Lrrc2 | 1.65E-06 | 1.553009 |
| ENSMUSG00000046818 | Ddit4l | 1.98E-02 | 1.554266 |
| ENSMUSG00000031808 | Slc27a1 | 6.93E-12 | 1.555141 |
| ENSMUSG00000027796 | Smad9 | 1.44E-04 | 1.555153 |
| ENSMUSG00000064368 | mt-Nd6 | 4.83E-18 | 1.555459 |
| ENSMUSG00000074497 | A430078G23Rik | 5.03E-17 | 1.557086 |
| ENSMUSG00000087579 | Hectd2os | 3.22E-07 | 1.557447 |
| ENSMUSG00000097917 | Gm26839 | 3.13E-02 | 1.55764 |
| ENSMUSG00000113184 | AC125351.1 | 6.09E-05 | 1.558477 |
| ENSMUSG00000043531 | Sorcs1 | 2.00E-07 | 1.560471 |
| ENSMUSG00000112947 | Gm47493 | 2.90E-03 | 1.560632 |
| ENSMUSG00000042474 | Fcmr | 3.47E-04 | 1.561669 |
| ENSMUSG00000039521 | Foxp3 | 3.68E-03 | 1.562481 |
| ENSMUSG00000030219 | Erp27 | 2.46E-02 | 1.563639 |
| ENSMUSG00000029158 | Yipf7 | 1.40E-02 | 1.563821 |
| ENSMUSG00000079015 | Serpina1c | 2.41E-03 | 1.56468 |
| ENSMUSG00000038132 | Rbm24 | 4.06E-11 | 1.564762 |
| ENSMUSG00000028116 | Myoz2 | 2.18E-12 | 1.566472 |
| ENSMUSG00000114138 | Gm36423 | 1.97E-02 | 1.566955 |
| ENSMUSG00000044938 | Klhl31 | 1.10E-08 | 1.567813 |
| ENSMUSG00000012705 | Retn | 1.11E-02 | 1.567825 |
| ENSMUSG00000030278 | Cidec | 1.15E-06 | 1.56902 |
| ENSMUSG00000075307 | Klhl41 | 1.95E-09 | 1.569759 |
| ENSMUSG00000043165 | Lor | 7.00E-06 | 1.56992 |
| ENSMUSG00000060962 | Dmkn | 6.45E-08 | 1.570534 |
| ENSMUSG00000038242 | Aox4 | 5.04E-03 | 1.571204 |
| ENSMUSG00000091712 | Sec14l5 | 1.06E-03 | 1.572691 |
| ENSMUSG00000013523 | Bcas1 | 3.93E-02 | 1.572801 |
| ENSMUSG00000092201 | A530058N18Rik | 1.98E-06 | 1.572952 |
| ENSMUSG00000083120 | Gm15854 | 3.07E-03 | 1.573014 |
| ENSMUSG00000034009 | Rxfp1 | 2.53E-03 | 1.573695 |
| ENSMUSG00000029499 | Pxmp2 | 6.54E-04 | 1.574213 |
| ENSMUSG00000097062 | Gm17586 | 4.09E-02 | 1.574318 |
| ENSMUSG00000078486 | Perm1 | 7.00E-10 | 1.575601 |
| ENSMUSG00000053024 | Cntn2 | 3.82E-02 | 1.576448 |
| ENSMUSG00000044694 | 2010007H06Rik | 3.28E-04 | 1.576597 |
| ENSMUSG00000101640 | Gm28376 | 4.52E-02 | 1.576948 |
| ENSMUSG00000093938 | Evi2b | 2.47E-13 | 1.577235 |
| ENSMUSG00000029838 | Ptn | 3.62E-03 | 1.578484 |
| ENSMUSG00000074433 | Lce3e | 4.14E-02 | 1.579193 |
| ENSMUSG00000113178 | Mylf-ps | 5.90E-07 | 1.579344 |
| ENSMUSG00000043629 | 1700019D03Rik | 8.68E-03 | 1.58036 |
| ENSMUSG00000099966 | 2810402E24Rik | 1.76E-03 | 1.580758 |
| ENSMUSG00000026989 | Dapl1 | 4.53E-02 | 1.581376 |
| ENSMUSG00000030598 | Fbxo17 | 3.90E-05 | 1.582412 |
| ENSMUSG00000054598 | 9130230L23Rik | 5.99E-08 | 1.582807 |

| | | | |
|---|---|---|---|
| ENSMUSG00000033508 | Asprv1 | 6.88E-04 | 1.583878 |
| ENSMUSG00000038295 | Atg9b | 1.40E-02 | 1.58412 |
| ENSMUSG00000096351 | Samd11 | 9.99E-04 | 1.584324 |
| ENSMUSG00000039084 | Chad | 9.78E-06 | 1.586243 |
| ENSMUSG00000076617 | Ighm | 5.17E-12 | 1.586353 |
| ENSMUSG00000086968 | 4933431E20Rik | 3.14E-02 | 1.586354 |
| ENSMUSG00000040086 | Tnni3k | 4.57E-09 | 1.587296 |
| ENSMUSG00000035606 | Ky | 2.23E-04 | 1.588323 |
| ENSMUSG00000038170 | Pde4dip | 3.86E-39 | 1.588667 |
| ENSMUSG00000029055 | Plch2 | 6.80E-08 | 1.588932 |
| ENSMUSG00000061013 | Mkx | 2.08E-04 | 1.589186 |
| ENSMUSG00000041782 | Lad1 | 1.14E-06 | 1.589714 |
| ENSMUSG00000027456 | Sdcbp2 | 1.49E-07 | 1.5906 |
| ENSMUSG00000020836 | Coro6 | 8.35E-11 | 1.591197 |
| ENSMUSG00000060180 | Myh13 | 3.21E-04 | 1.591427 |
| ENSMUSG00000053093 | Myh7 | 2.88E-07 | 1.593422 |
| ENSMUSG00000109510 | Gm42417 | 7.99E-14 | 1.595001 |
| ENSMUSG00000057092 | Fxyd3 | 3.96E-03 | 1.596509 |
| ENSMUSG00000117297 | CT571247.2 | 9.90E-03 | 1.597064 |
| ENSMUSG00000090173 | Fbxw10 | 2.39E-04 | 1.598188 |
| ENSMUSG00000054889 | Dsp | 3.20E-46 | 1.598377 |
| ENSMUSG00000041616 | Nppa | 1.28E-03 | 1.59881 |
| ENSMUSG00000056271 | Lman1l | 4.16E-02 | 1.599281 |
| ENSMUSG00000057280 | Musk | 5.50E-08 | 1.600621 |
| ENSMUSG00000031377 | Bmx | 1.66E-10 | 1.601368 |
| ENSMUSG00000064340 | mt-Tl1 | 1.92E-09 | 1.601856 |
| ENSMUSG00000027559 | Car3 | 1.93E-38 | 1.60232 |
| ENSMUSG00000061462 | Obscn | 1.92E-25 | 1.602458 |
| ENSMUSG00000102305 | Gm38192 | 2.46E-02 | 1.603866 |
| ENSMUSG00000038173 | Enpp6 | 2.21E-02 | 1.604535 |
| ENSMUSG00000028150 | Rorc | 9.93E-07 | 1.604951 |
| ENSMUSG00000029167 | Ppargc1a | 2.14E-18 | 1.605113 |
| ENSMUSG00000092397 | C130080G10Rik | 7.30E-06 | 1.605123 |
| ENSMUSG00000103984 | Gm37447 | 3.69E-02 | 1.605238 |
| ENSMUSG00000040055 | Gjb6 | 2.24E-03 | 1.605389 |
| ENSMUSG00000024347 | Psd2 | 2.70E-02 | 1.606216 |
| ENSMUSG00000068885 | Lce3f | 8.58E-03 | 1.607541 |
| ENSMUSG00000037989 | Wnk2 | 1.81E-14 | 1.608343 |
| ENSMUSG00000085184 | 4933439K11Rik | 3.15E-03 | 1.608853 |
| ENSMUSG00000031097 | Tnni2 | 2.26E-08 | 1.6098 |
| ENSMUSG00000017400 | Stac2 | 4.29E-02 | 1.612166 |
| ENSMUSG00000024673 | Ms4a1 | 8.11E-06 | 1.612963 |
| ENSMUSG00000033544 | Angptl1 | 1.17E-02 | 1.613351 |
| ENSMUSG00000097652 | Mhrt | 2.82E-07 | 1.613894 |
| ENSMUSG00000085971 | Gm15411 | 3.02E-05 | 1.613896 |
| ENSMUSG00000075289 | Carns1 | 1.74E-06 | 1.615629 |
| ENSMUSG00000026163 | Sphkap | 7.15E-09 | 1.617694 |
| ENSMUSG00000040694 | Apobec2 | 6.16E-08 | 1.620042 |
| ENSMUSG00000043008 | Klhl6 | 3.96E-14 | 1.62231 |
| ENSMUSG00000018893 | Mb | 2.71E-08 | 1.623243 |
| ENSMUSG00000027861 | Casq2 | 1.05E-11 | 1.623312 |
| ENSMUSG00000020067 | Mypn | 1.05E-14 | 1.623658 |
| ENSMUSG00000028115 | Bnipl | 1.13E-02 | 1.623683 |

| | | | |
|---|---|---|---|
| ENSMUSG00000054659 | Pm20d2 | 1.90E-02 | 1.623767 |
| ENSMUSG00000055415 | Atp10b | 9.65E-03 | 1.624498 |
| ENSMUSG00000039891 | Txlnb | 6.59E-16 | 1.625928 |
| ENSMUSG00000020169 | Best3 | 3.97E-06 | 1.62632 |
| ENSMUSG00000085194 | Platr32 | 1.32E-02 | 1.628265 |
| ENSMUSG00000063297 | Luzp2 | 9.47E-06 | 1.63024 |
| ENSMUSG00000025815 | Dhtkd1 | 2.73E-06 | 1.630385 |
| ENSMUSG00000021123 | Rdh12 | 1.62E-12 | 1.631252 |
| ENSMUSG00000046056 | Sbsn | 8.20E-05 | 1.631281 |
| ENSMUSG00000076275 | Mir675 | 3.45E-04 | 1.631535 |
| ENSMUSG00000036560 | Lgi4 | 6.45E-08 | 1.632499 |
| ENSMUSG00000111818 | Gm17749 | 8.46E-06 | 1.632708 |
| ENSMUSG00000052496 | Pkdrej | 1.85E-02 | 1.635592 |
| ENSMUSG00000085417 | Gm13919 | 1.14E-02 | 1.637562 |
| ENSMUSG00000031700 | Gpt2 | 2.20E-10 | 1.638222 |
| ENSMUSG00000086564 | Cd101 | 2.04E-15 | 1.638923 |
| ENSMUSG00000067149 | Jchain | 6.64E-03 | 1.639225 |
| ENSMUSG00000049123 | Catsperg2 | 3.63E-02 | 1.639878 |
| ENSMUSG00000057719 | Sh3rf2 | 1.37E-13 | 1.640066 |
| ENSMUSG00000027887 | Sypl2 | 7.30E-06 | 1.641809 |
| ENSMUSG00000066510 | Ankdd1a | 8.15E-07 | 1.642186 |
| ENSMUSG00000102191 | Gm36569 | 1.33E-02 | 1.642411 |
| ENSMUSG00000022523 | Fgf12 | 9.16E-19 | 1.642983 |
| ENSMUSG00000058470 | Gm8369 | 1.21E-10 | 1.644013 |
| ENSMUSG00000096646 | CR974586.6 | 4.85E-02 | 1.647968 |
| ENSMUSG00000079853 | Klra1 | 4.74E-04 | 1.648672 |
| ENSMUSG00000029862 | Clcn1 | 3.16E-06 | 1.649099 |
| ENSMUSG00000007279 | Scube2 | 3.52E-19 | 1.651431 |
| ENSMUSG00000032564 | Cpne4 | 1.36E-05 | 1.651458 |
| ENSMUSG00000043487 | Acot6 | 1.05E-02 | 1.65192 |
| ENSMUSG00000014453 | Blk | 1.77E-09 | 1.652393 |
| ENSMUSG00000059857 | Ntng1 | 2.28E-05 | 1.653485 |
| ENSMUSG00000057246 | BC051142 | 2.48E-04 | 1.653565 |
| ENSMUSG00000033569 | Adgrb3 | 4.62E-07 | 1.653669 |
| ENSMUSG00000041538 | H2-Ob | 1.37E-05 | 1.653876 |
| ENSMUSG00000029361 | Nos1 | 1.56E-11 | 1.653885 |
| ENSMUSG00000106547 | B230303O12Rik | 1.21E-08 | 1.654582 |
| ENSMUSG00000072720 | Myo18b | 4.14E-27 | 1.655567 |
| ENSMUSG00000086962 | Gm12248 | 3.67E-04 | 1.656171 |
| ENSMUSG00000020374 | Rasgef1c | 1.48E-09 | 1.656757 |
| ENSMUSG00000033182 | Kbtbd12 | 1.02E-07 | 1.656789 |
| ENSMUSG00000105646 | Gm30211 | 1.89E-10 | 1.657828 |
| ENSMUSG00000054582 | Pabpc1l | 1.93E-07 | 1.660152 |
| ENSMUSG00000047228 | A2ml1 | 2.62E-03 | 1.660287 |
| ENSMUSG00000098381 | Mir7225 | 3.46E-03 | 1.660541 |
| ENSMUSG00000085532 | B430319H21Rik | 1.21E-03 | 1.661812 |
| ENSMUSG00000079764 | AC087559.1 | 4.29E-02 | 1.662417 |
| ENSMUSG00000108436 | Gm44851 | 1.77E-02 | 1.663035 |
| ENSMUSG00000078817 | Nlrp12 | 1.01E-15 | 1.663158 |
| ENSMUSG00000110195 | Pde2a | 7.09E-18 | 1.665411 |
| ENSMUSG00000075555 | Gm10863 | 1.92E-04 | 1.665852 |
| ENSMUSG00000037572 | Wdhd1 | 5.63E-09 | 1.666104 |
| ENSMUSG00000021506 | Pitx1 | 2.48E-02 | 1.668955 |

| | | | |
|---|---|---|---|
| ENSMUSG00000026564 | Dusp27 | 2.96E-09 | 1.67013 |
| ENSMUSG00000037188 | Grhl3 | 5.01E-04 | 1.670721 |
| ENSMUSG00000028834 | Trim63 | 5.36E-07 | 1.67122 |
| ENSMUSG00000038077 | Kcna6 | 3.83E-05 | 1.672132 |
| ENSMUSG00000063651 | Cnfn | 3.01E-05 | 1.673857 |
| ENSMUSG00000031245 | Hmgn5 | 1.75E-06 | 1.674099 |
| ENSMUSG00000028544 | Slc5a9 | 4.27E-03 | 1.67494 |
| ENSMUSG00000058966 | Fam57b | 2.45E-04 | 1.676352 |
| ENSMUSG00000022416 | Cacna1i | 7.73E-05 | 1.678167 |
| ENSMUSG00000036731 | Cysrt1 | 1.69E-02 | 1.678982 |
| ENSMUSG00000046352 | Gjb2 | 7.16E-07 | 1.679256 |
| ENSMUSG00000028459 | Cd72 | 3.05E-05 | 1.679816 |
| ENSMUSG00000045534 | Kcna5 | 2.06E-03 | 1.681084 |
| ENSMUSG00000000244 | Tspan32 | 1.65E-11 | 1.681161 |
| ENSMUSG00000108985 | A930030B08Rik | 2.28E-02 | 1.682419 |
| ENSMUSG00000032942 | Ucp3 | 6.10E-07 | 1.684317 |
| ENSMUSG00000087336 | Gm15860 | 2.35E-02 | 1.684872 |
| ENSMUSG00000002831 | Plin4 | 1.20E-31 | 1.686763 |
| ENSMUSG00000020848 | Doc2b | 1.14E-10 | 1.686804 |
| ENSMUSG00000052726 | Kcnt2 | 2.57E-21 | 1.687119 |
| ENSMUSG00000024105 | Themis3 | 1.57E-02 | 1.68772 |
| ENSMUSG00000112707 | D830005E20Rik | 1.41E-09 | 1.688501 |
| ENSMUSG00000030317 | Timp4 | 1.18E-08 | 1.688584 |
| ENSMUSG00000095438 | Mir133a-1hg | 2.49E-05 | 1.688705 |
| ENSMUSG00000034308 | Sdr42e1 | 1.55E-04 | 1.688804 |
| ENSMUSG00000070385 | Ampd1 | 9.23E-08 | 1.689044 |
| ENSMUSG00000086405 | 9330198N18Rik | 9.76E-07 | 1.689856 |
| ENSMUSG00000029660 | Tex26 | 5.91E-07 | 1.690097 |
| ENSMUSG00000032262 | Elovl4 | 6.32E-05 | 1.690396 |
| ENSMUSG00000031461 | Myom2 | 2.01E-27 | 1.69079 |
| ENSMUSG00000040740 | Slc25a34 | 2.14E-05 | 1.690811 |
| ENSMUSG00000000049 | Apoh | 2.58E-02 | 1.69197 |
| ENSMUSG00000023019 | Gpd1 | 5.00E-16 | 1.693018 |
| ENSMUSG00000030510 | Cers3 | 7.37E-07 | 1.693354 |
| ENSMUSG00000076937 | Iglc2 | 2.70E-03 | 1.693807 |
| ENSMUSG00000038763 | Alpk3 | 6.16E-26 | 1.693831 |
| ENSMUSG00000067889 | Sptbn2 | 2.00E-07 | 1.694539 |
| ENSMUSG00000106245 | Gm43824 | 1.15E-02 | 1.694809 |
| ENSMUSG00000042567 | Nek10 | 1.91E-12 | 1.695091 |
| ENSMUSG00000048764 | Tmprss11f | 8.49E-03 | 1.696489 |
| ENSMUSG00000022878 | Adipoq | 5.10E-07 | 1.69666 |
| ENSMUSG00000026527 | Rgs7 | 1.36E-33 | 1.696693 |
| ENSMUSG00000090206 | Tepp | 1.43E-03 | 1.696892 |
| ENSMUSG00000002500 | Rpl3l | 1.67E-04 | 1.698266 |
| ENSMUSG00000054057 | A930004D18Rik | 1.98E-02 | 1.699027 |
| ENSMUSG00000079644 | Gm1110 | 1.53E-02 | 1.699165 |
| ENSMUSG00000059201 | Lep | 5.30E-03 | 1.70052 |
| ENSMUSG00000061816 | Myl1 | 3.42E-12 | 1.701209 |
| ENSMUSG00000043366 | Olfr78 | 4.74E-03 | 1.701217 |
| ENSMUSG00000039092 | Sptlc3 | 1.10E-03 | 1.702114 |
| ENSMUSG00000053964 | Lgals4 | 6.56E-05 | 1.702508 |
| ENSMUSG00000025892 | Gria4 | 3.55E-03 | 1.703464 |
| ENSMUSG00000019906 | Lin7a | 1.80E-05 | 1.704483 |

| | | | |
|---|---|---|---|
| ENSMUSG00000041020 | Map7d2 | 2.72E-02 | 1.705429 |
| ENSMUSG00000100147 | 1700047M11Rik | 4.18E-08 | 1.706711 |
| ENSMUSG00000028773 | Fabp3 | 1.13E-08 | 1.706932 |
| ENSMUSG00000031385 | Plxnb3 | 2.72E-03 | 1.709339 |
| ENSMUSG00000110411 | Gm45457 | 9.03E-03 | 1.710479 |
| ENSMUSG00000034282 | Evpl | 1.07E-08 | 1.710621 |
| ENSMUSG00000087628 | Gm13028 | 1.40E-03 | 1.71063 |
| ENSMUSG00000036960 | Clca2 | 2.27E-02 | 1.712431 |
| ENSMUSG00000052013 | Btla | 1.86E-06 | 1.713815 |
| ENSMUSG00000025754 | Agbl1 | 1.29E-20 | 1.714057 |
| ENSMUSG00000034634 | Ly6d | 7.90E-04 | 1.715269 |
| ENSMUSG00000061126 | Cyp4f39 | 3.92E-03 | 1.718439 |
| ENSMUSG00000035580 | Kcnh8 | 9.67E-06 | 1.718443 |
| ENSMUSG00000021340 | Gpld1 | 5.88E-10 | 1.718732 |
| ENSMUSG00000040350 | Trim7 | 7.94E-08 | 1.718782 |
| ENSMUSG00000116987 | AC150035.3 | 1.76E-02 | 1.719502 |
| ENSMUSG00000038204 | Asb10 | 2.36E-02 | 1.719976 |
| ENSMUSG00000020334 | Slc22a4 | 1.93E-07 | 1.720261 |
| ENSMUSG00000064346 | mt-Tw | 2.44E-04 | 1.721955 |
| ENSMUSG00000007033 | Hspa1l | 2.73E-02 | 1.722982 |
| ENSMUSG00000038239 | Hrc | 6.28E-14 | 1.724801 |
| ENSMUSG00000073492 | Gm10521 | 2.63E-03 | 1.725675 |
| ENSMUSG00000020279 | Il9r | 3.97E-02 | 1.726651 |
| ENSMUSG00000040899 | Ccr6 | 4.51E-09 | 1.728272 |
| ENSMUSG00000073985 | Gm10602 | 5.77E-04 | 1.7291 |
| ENSMUSG00000069733 | Ube2u | 5.69E-03 | 1.72999 |
| ENSMUSG00000036123 | Slc9a3 | 8.61E-03 | 1.729994 |
| ENSMUSG00000019787 | Trdn | 2.22E-33 | 1.730141 |
| ENSMUSG00000099364 | 5730419F03Rik | 1.15E-02 | 1.732264 |
| ENSMUSG00000093577 | Gm20632 | 4.01E-02 | 1.733244 |
| ENSMUSG00000027077 | Smtnl1 | 4.61E-06 | 1.734518 |
| ENSMUSG00000057003 | Myh4 | 3.19E-14 | 1.734608 |
| ENSMUSG00000000031 | H19 | 5.10E-18 | 1.735173 |
| ENSMUSG00000003271 | Sult2b1 | 1.84E-03 | 1.735974 |
| ENSMUSG00000038354 | Ankrd35 | 3.79E-03 | 1.736278 |
| ENSMUSG00000015879 | Fam184b | 6.83E-03 | 1.737161 |
| ENSMUSG00000079243 | Xirp1 | 1.34E-16 | 1.737887 |
| ENSMUSG00000079588 | Tmem182 | 9.64E-13 | 1.738319 |
| ENSMUSG00000031376 | Atp2b3 | 9.23E-04 | 1.73926 |
| ENSMUSG00000042010 | Acacb | 9.27E-37 | 1.739339 |
| ENSMUSG00000022871 | Fetub | 5.57E-05 | 1.739701 |
| ENSMUSG00000010066 | Cacna2d2 | 4.85E-04 | 1.74003 |
| ENSMUSG00000027831 | Veph1 | 7.22E-11 | 1.743301 |
| ENSMUSG00000005268 | Prlr | 1.86E-05 | 1.743362 |
| ENSMUSG00000038665 | Dgki | 7.51E-11 | 1.743848 |
| ENSMUSG00000040536 | Necab1 | 6.32E-03 | 1.744057 |
| ENSMUSG00000062713 | Sim2 | 3.32E-09 | 1.745706 |
| ENSMUSG00000037139 | Myom3 | 3.04E-14 | 1.746519 |
| ENSMUSG00000014030 | Pax5 | 3.89E-07 | 1.746801 |
| ENSMUSG00000031626 | Sorbs2 | 8.87E-27 | 1.746865 |
| ENSMUSG00000048142 | Nat8l | 3.69E-04 | 1.747147 |
| ENSMUSG00000067341 | H2-Eb2 | 8.84E-03 | 1.748009 |
| ENSMUSG00000025172 | Ankrd2 | 1.45E-06 | 1.749393 |

| | | | |
|---|---|---|---|
| ENSMUSG00000096974 | Gm26881 | 1.04E-02 | 1.749441 |
| ENSMUSG00000020000 | Moxd1 | 4.65E-03 | 1.750501 |
| ENSMUSG00000070423 | Olfr558 | 4.01E-02 | 1.750691 |
| ENSMUSG00000099449 | Gm28401 | 4.06E-02 | 1.754089 |
| ENSMUSG00000067653 | Ankrd23 | 1.06E-15 | 1.754635 |
| ENSMUSG00000019102 | Aldh3a1 | 7.57E-08 | 1.754898 |
| ENSMUSG00000027868 | Tbx15 | 4.21E-14 | 1.755594 |
| ENSMUSG00000051747 | Ttn | 1.12E-46 | 1.757967 |
| ENSMUSG00000032033 | Barx2 | 1.39E-07 | 1.759194 |
| ENSMUSG00000022622 | Acr | 1.21E-08 | 1.760657 |
| ENSMUSG00000057054 | Inca1 | 1.12E-04 | 1.76172 |
| ENSMUSG00000115610 | Gm31251 | 3.90E-14 | 1.761914 |
| ENSMUSG00000061259 | Tmprss11d | 1.42E-02 | 1.761956 |
| ENSMUSG00000069830 | Nlrp1a | 1.11E-03 | 1.76325 |
| ENSMUSG00000110010 | Gm45629 | 1.53E-03 | 1.764151 |
| ENSMUSG00000038663 | Fsd2 | 1.08E-14 | 1.765708 |
| ENSMUSG00000100627 | A830008E24Rik | 1.45E-02 | 1.766465 |
| ENSMUSG00000001095 | Slc13a2 | 5.92E-04 | 1.767172 |
| ENSMUSG00000024827 | Gldc | 3.59E-02 | 1.767944 |
| ENSMUSG00000027913 | Crct1 | 4.51E-05 | 1.768545 |
| ENSMUSG00000013483 | Card14 | 4.06E-05 | 1.769133 |
| ENSMUSG00000030592 | Ryr1 | 3.35E-10 | 1.769178 |
| ENSMUSG00000102759 | Gm10463 | 4.04E-02 | 1.770039 |
| ENSMUSG00000074254 | Cyp2a4 | 3.39E-02 | 1.770248 |
| ENSMUSG00000058975 | Kcnc1 | 6.09E-05 | 1.770841 |
| ENSMUSG00000054325 | Lce3a | 5.92E-04 | 1.771249 |
| ENSMUSG00000022040 | Ephx2 | 1.34E-07 | 1.77423 |
| ENSMUSG00000069670 | Nkain2 | 1.14E-30 | 1.775621 |
| ENSMUSG00000099994 | 1700024B18Rik | 3.27E-02 | 1.775772 |
| ENSMUSG00000039873 | Neurl2 | 3.62E-03 | 1.776821 |
| ENSMUSG00000086765 | Gm11827 | 4.14E-05 | 1.779138 |
| ENSMUSG00000093765 | Gm20658 | 4.32E-02 | 1.779401 |
| ENSMUSG00000044708 | Kcnj10 | 2.20E-02 | 1.780045 |
| ENSMUSG00000010476 | Ebf3 | 2.43E-10 | 1.782939 |
| ENSMUSG00000043681 | Fam25c | 5.16E-03 | 1.783067 |
| ENSMUSG00000026220 | Slc16a14 | 1.02E-02 | 1.783575 |
| ENSMUSG00000048078 | Tenm4 | 8.65E-33 | 1.786182 |
| ENSMUSG00000074625 | Arhgap40 | 1.61E-05 | 1.786986 |
| ENSMUSG00000024334 | H2-Oa | 2.83E-02 | 1.78792 |
| ENSMUSG00000005373 | Mlxipl | 1.48E-14 | 1.792638 |
| ENSMUSG00000031757 | Mt4 | 1.02E-03 | 1.792943 |
| ENSMUSG00000104927 | Gm43388 | 8.61E-03 | 1.794412 |
| ENSMUSG00000031594 | Fgl1 | 3.86E-02 | 1.796159 |
| ENSMUSG00000017300 | Tnnc2 | 5.41E-14 | 1.797829 |
| ENSMUSG00000087307 | Gm12925 | 4.11E-02 | 1.79825 |
| ENSMUSG00000100199 | Gm20324 | 5.05E-04 | 1.800659 |
| ENSMUSG00000109754 | Gm39214 | 4.56E-02 | 1.800915 |
| ENSMUSG00000040490 | Lrfn2 | 2.65E-03 | 1.803413 |
| ENSMUSG00000110631 | Gm42047 | 2.58E-02 | 1.804324 |
| ENSMUSG00000111809 | Gm49334 | 5.99E-03 | 1.805073 |
| ENSMUSG00000048070 | Pirt | 4.90E-02 | 1.805324 |
| ENSMUSG00000111299 | 4930540M03Rik | 3.88E-02 | 1.805841 |
| ENSMUSG00000026489 | Coq8a | 5.21E-32 | 1.80787 |

| | | | |
|---|---|---|---|
| ENSMUSG00000085171 | D830026I12Rik | 1.96E-02 | 1.808535 |
| ENSMUSG00000000673 | Haao | 2.83E-04 | 1.808707 |
| ENSMUSG00000037661 | Gpr160 | 1.16E-06 | 1.8137 |
| ENSMUSG00000074215 | Gm10643 | 2.95E-02 | 1.814076 |
| ENSMUSG00000096449 | Gm4076 | 4.29E-02 | 1.815678 |
| ENSMUSG00000022383 | Ppara | 2.50E-24 | 1.817544 |
| ENSMUSG00000109157 | Gm44829 | 3.21E-05 | 1.818829 |
| ENSMUSG00000020698 | Cct6b | 6.42E-04 | 1.823548 |
| ENSMUSG00000049336 | Tenm2 | 1.31E-16 | 1.8238 |
| ENSMUSG00000074199 | Krtdap | 2.27E-03 | 1.824717 |
| ENSMUSG00000005547 | Cyp2a5 | 1.12E-09 | 1.826543 |
| ENSMUSG00000027261 | Hao1 | 1.24E-02 | 1.82689 |
| ENSMUSG00000037446 | Tulp1 | 5.73E-03 | 1.828912 |
| ENSMUSG00000086547 | Gm11755 | 1.77E-02 | 1.832213 |
| ENSMUSG00000087090 | Nctc1 | 7.99E-18 | 1.832231 |
| ENSMUSG00000047419 | Cmya5 | 3.32E-33 | 1.832861 |
| ENSMUSG00000031425 | Plp1 | 8.47E-11 | 1.833127 |
| ENSMUSG00000067616 | Klk11 | 1.67E-03 | 1.835648 |
| ENSMUSG00000079025 | Gsdmc | 4.96E-03 | 1.835798 |
| ENSMUSG00000085479 | 9430073C21Rik | 4.93E-02 | 1.836068 |
| ENSMUSG00000027674 | Pex5l | 3.20E-11 | 1.836523 |
| ENSMUSG00000001027 | Scn4a | 8.93E-13 | 1.838172 |
| ENSMUSG00000038020 | Rapgefl1 | 3.35E-07 | 1.842817 |
| ENSMUSG00000037509 | Arhgef4 | 1.01E-11 | 1.84526 |
| ENSMUSG00000056569 | Mpz | 2.39E-15 | 1.845464 |
| ENSMUSG00000018862 | Otop3 | 2.40E-03 | 1.847993 |
| ENSMUSG00000027022 | Xirp2 | 2.79E-35 | 1.849448 |
| ENSMUSG00000116508 | Gm49463 | 1.89E-02 | 1.850853 |
| ENSMUSG00000044788 | Fads6 | 2.44E-04 | 1.85333 |
| ENSMUSG00000031372 | Trex2 | 3.78E-02 | 1.854308 |
| ENSMUSG00000028573 | Fggy | 2.26E-05 | 1.854817 |
| ENSMUSG00000029683 | Lmod2 | 1.32E-16 | 1.857354 |
| ENSMUSG00000022491 | Glycam1 | 8.74E-05 | 1.858915 |
| ENSMUSG00000030724 | Cd19 | 1.17E-10 | 1.860844 |
| ENSMUSG00000087242 | C78197 | 4.25E-02 | 1.86137 |
| ENSMUSG00000040340 | Tex45 | 7.93E-03 | 1.861819 |
| ENSMUSG00000021613 | Hapln1 | 2.16E-04 | 1.864932 |
| ENSMUSG00000112895 | Gm47567 | 7.34E-05 | 1.86531 |
| ENSMUSG00000085743 | 8430419K02Rik | 6.18E-22 | 1.866081 |
| ENSMUSG00000101930 | Gm5441 | 1.11E-09 | 1.866612 |
| ENSMUSG00000042251 | Pm20d1 | 6.12E-05 | 1.867473 |
| ENSMUSG00000103965 | Gm30173 | 2.07E-04 | 1.869946 |
| ENSMUSG00000028996 | Rbp7 | 6.66E-03 | 1.871717 |
| ENSMUSG00000042031 | Lce3b | 1.73E-04 | 1.872678 |
| ENSMUSG00000042895 | Abra | 1.62E-09 | 1.874549 |
| ENSMUSG00000026697 | Myoc | 1.02E-02 | 1.875415 |
| ENSMUSG00000026100 | Mstn | 8.60E-03 | 1.877754 |
| ENSMUSG00000091119 | Ccdc152 | 2.42E-05 | 1.879304 |
| ENSMUSG00000062515 | Fabp4 | 8.53E-23 | 1.879539 |
| ENSMUSG00000062372 | Otof | 6.43E-03 | 1.881054 |
| ENSMUSG00000021622 | Ckmt2 | 1.81E-18 | 1.882596 |
| ENSMUSG00000096960 | A230028O05Rik | 2.40E-09 | 1.883268 |
| ENSMUSG00000104453 | Gm37829 | 6.61E-05 | 1.884749 |

| | | | |
|---|---|---|---|
| ENSMUSG00000039496 | Cdnf | 4.66E-03 | 1.887309 |
| ENSMUSG00000034645 | Zyg11a | 2.17E-02 | 1.88839 |
| ENSMUSG00000074862 | BC025920 | 2.90E-03 | 1.888707 |
| ENSMUSG00000030577 | Cd22 | 5.61E-10 | 1.888805 |
| ENSMUSG00000046480 | Scn4b | 2.33E-10 | 1.889073 |
| ENSMUSG00000061723 | Tnnt3 | 3.36E-12 | 1.895792 |
| ENSMUSG00000073602 | Serpinb3b | 8.46E-04 | 1.896022 |
| ENSMUSG00000042250 | Pglyrp4 | 1.68E-02 | 1.896106 |
| ENSMUSG00000037977 | 6430571L13Rik | 1.82E-05 | 1.897294 |
| ENSMUSG00000071858 | Gm94 | 1.11E-03 | 1.898142 |
| ENSMUSG00000043461 | Sptssb | 3.69E-04 | 1.8982 |
| ENSMUSG00000027470 | Mylk2 | 8.39E-14 | 1.898313 |
| ENSMUSG00000084929 | Foxo6os | 1.25E-02 | 1.898699 |
| ENSMUSG00000059832 | Kprp | 6.59E-08 | 1.901066 |
| ENSMUSG00000020061 | Mybpc1 | 4.60E-23 | 1.903022 |
| ENSMUSG00000000223 | Drp2 | 6.47E-08 | 1.90562 |
| ENSMUSG00000042514 | Klhl14 | 9.78E-05 | 1.906063 |
| ENSMUSG00000097286 | Gm26684 | 2.80E-02 | 1.914855 |
| ENSMUSG00000038602 | Slc35f1 | 6.46E-23 | 1.915687 |
| ENSMUSG00000029602 | Rasal1 | 7.61E-04 | 1.918457 |
| ENSMUSG00000025431 | Crisp1 | 6.65E-04 | 1.918683 |
| ENSMUSG00000047586 | Nccrp1 | 3.55E-02 | 1.920616 |
| ENSMUSG00000086980 | Gm13791 | 1.98E-02 | 1.922479 |
| ENSMUSG00000049134 | Nrap | 2.40E-35 | 1.926123 |
| ENSMUSG00000044737 | Klk14 | 1.25E-07 | 1.926874 |
| ENSMUSG00000071540 | 3425401B19Rik | 1.95E-20 | 1.927593 |
| ENSMUSG00000039763 | Dnajc28 | 1.34E-03 | 1.92807 |
| ENSMUSG00000041423 | Paqr6 | 9.00E-03 | 1.931307 |
| ENSMUSG00000045475 | Lce3c | 5.69E-05 | 1.931506 |
| ENSMUSG00000032311 | Nrg4 | 1.59E-17 | 1.932314 |
| ENSMUSG00000039269 | 2300002M23Rik | 5.11E-03 | 1.934349 |
| ENSMUSG00000050840 | Cdh20 | 3.44E-04 | 1.936624 |
| ENSMUSG00000032278 | Paqr5 | 8.26E-10 | 1.937115 |
| ENSMUSG00000039376 | Synpo2l | 3.96E-21 | 1.938741 |
| ENSMUSG00000007653 | Gabrb2 | 6.55E-03 | 1.938974 |
| ENSMUSG00000056328 | Myh1 | 2.30E-34 | 1.944869 |
| ENSMUSG00000032503 | Arpp21 | 1.12E-06 | 1.948018 |
| ENSMUSG00000090118 | Gm16163 | 7.49E-08 | 1.950295 |
| ENSMUSG00000005355 | Casp14 | 2.78E-02 | 1.950333 |
| ENSMUSG00000019851 | Perp | 2.82E-14 | 1.951709 |
| ENSMUSG00000026950 | Neb | 2.63E-19 | 1.958489 |
| ENSMUSG00000041476 | Smpx | 7.49E-08 | 1.958962 |
| ENSMUSG00000055333 | Fat2 | 8.85E-09 | 1.964371 |
| ENSMUSG00000032281 | Acsbg1 | 6.04E-11 | 1.96499 |
| ENSMUSG00000026450 | Chit1 | 3.30E-03 | 1.966851 |
| ENSMUSG00000110580 | D830024N08Rik | 3.00E-09 | 1.967915 |
| ENSMUSG00000039070 | Cpa4 | 2.42E-05 | 1.968184 |
| ENSMUSG00000021223 | Papln | 5.90E-05 | 1.970017 |
| ENSMUSG00000050808 | Muc15 | 3.15E-02 | 1.970166 |
| ENSMUSG00000096169 | Olfr1564 | 4.84E-02 | 1.970537 |
| ENSMUSG00000067242 | Lgi1 | 1.81E-04 | 1.972543 |
| ENSMUSG00000001497 | Pax9 | 7.05E-07 | 1.973159 |
| ENSMUSG00000041737 | Tmem45b | 1.68E-09 | 1.973324 |

| | | | |
|---|---|---|---|
| ENSMUSG00000055561 | Spink5 | 1.65E-11 | 1.975498 |
| ENSMUSG00000037129 | Tmprss13 | 2.20E-05 | 1.978708 |
| ENSMUSG00000112471 | Gm5779 | 6.88E-03 | 1.980462 |
| ENSMUSG00000027833 | Shox2 | 1.87E-02 | 1.981285 |
| ENSMUSG00000043286 | Pnpla1 | 1.95E-03 | 1.981508 |
| ENSMUSG00000105986 | Gm43065 | 5.52E-04 | 1.983525 |
| ENSMUSG00000030935 | Acsm3 | 9.82E-03 | 1.984411 |
| ENSMUSG00000059742 | Kcnh7 | 1.36E-03 | 1.986178 |
| ENSMUSG00000050650 | Mrgpra1 | 3.24E-02 | 1.986835 |
| ENSMUSG00000034730 | Adgrb1 | 1.54E-02 | 1.986876 |
| ENSMUSG00000050663 | Trhde | 2.20E-03 | 1.987786 |
| ENSMUSG00000034774 | Dsg1c | 2.60E-02 | 1.98819 |
| ENSMUSG00000030178 | Klra13-ps | 3.18E-03 | 1.989497 |
| ENSMUSG00000030730 | Atp2a1 | 2.28E-32 | 1.98969 |
| ENSMUSG00000027376 | Prom2 | 3.25E-06 | 1.991537 |
| ENSMUSG00000025105 | Bnc1 | 5.33E-04 | 1.99208 |
| ENSMUSG00000039956 | Mrap | 5.20E-09 | 1.993265 |
| ENSMUSG00000104213 | Ighd | 3.20E-10 | 1.994516 |
| ENSMUSG00000024331 | Dsc2 | 6.73E-10 | 1.997784 |
| ENSMUSG00000103243 | Lce1d | 3.79E-03 | 1.999738 |
| ENSMUSG00000050463 | Krt78 | 3.05E-12 | 2.000453 |
| ENSMUSG00000021359 | Tfap2a | 1.09E-03 | 2.000592 |
| ENSMUSG00000025329 | Padi1 | 1.62E-02 | 2.00394 |
| ENSMUSG00000031204 | Asb12 | 4.92E-02 | 2.004544 |
| ENSMUSG00000039518 | Cdsn | 8.05E-07 | 2.005848 |
| ENSMUSG00000031844 | Hsd17b2 | 1.89E-04 | 2.007414 |
| ENSMUSG00000099906 | Gm28653 | 1.27E-02 | 2.010149 |
| ENSMUSG00000075296 | Aldh3b2 | 1.77E-03 | 2.010847 |
| ENSMUSG00000031936 | Hephl1 | 1.35E-02 | 2.013642 |
| ENSMUSG00000102590 | Mannr | 1.14E-05 | 2.01519 |
| ENSMUSG00000026834 | Acvr1c | 4.87E-08 | 2.015446 |
| ENSMUSG00000030834 | Abcc6 | 6.84E-03 | 2.015784 |
| ENSMUSG00000025425 | St8sia5 | 2.17E-02 | 2.018317 |
| ENSMUSG00000036854 | Hspb6 | 1.10E-06 | 2.019132 |
| ENSMUSG00000019853 | Hebp2 | 4.55E-03 | 2.019308 |
| ENSMUSG00000050359 | Sprr1a | 4.50E-07 | 2.019357 |
| ENSMUSG00000110390 | Gm45869 | 4.89E-02 | 2.02195 |
| ENSMUSG00000090356 | Teddm3 | 1.30E-03 | 2.026459 |
| ENSMUSG00000050296 | Abca12 | 8.96E-08 | 2.026805 |
| ENSMUSG00000047746 | Fbxo40 | 8.92E-18 | 2.029312 |
| ENSMUSG00000032726 | Bmp8a | 5.87E-03 | 2.030065 |
| ENSMUSG00000029154 | Cwh43 | 1.88E-02 | 2.030933 |
| ENSMUSG00000055489 | Ano5 | 8.68E-17 | 2.033201 |
| ENSMUSG00000017204 | Gsdma | 3.20E-05 | 2.033369 |
| ENSMUSG00000044086 | Lmod3 | 3.08E-06 | 2.045815 |
| ENSMUSG00000110843 | Gm35552 | 4.00E-02 | 2.047718 |
| ENSMUSG00000033579 | Fa2h | 3.55E-03 | 2.048412 |
| ENSMUSG00000028989 | Angptl7 | 3.95E-11 | 2.049447 |
| ENSMUSG00000114493 | Gm47071 | 9.81E-10 | 2.052629 |
| ENSMUSG00000074489 | Bglap3 | 1.66E-03 | 2.058281 |
| ENSMUSG00000060621 | Nkpd1 | 1.03E-02 | 2.065488 |
| ENSMUSG00000062826 | Ces2f | 1.92E-08 | 2.067273 |
| ENSMUSG00000030546 | Plin1 | 8.52E-14 | 2.067798 |

| | | | |
|---|---|---|---|
| ENSMUSG00000109311 | AI314278 | 4.74E-04 | 2.069279 |
| ENSMUSG00000023473 | Celsr3 | 1.86E-12 | 2.073428 |
| ENSMUSG00000096001 | 2610528A11Rik | 3.36E-02 | 2.073506 |
| ENSMUSG00000089941 | Gm16168 | 2.39E-07 | 2.073697 |
| ENSMUSG00000074647 | Fam83c | 2.18E-02 | 2.076138 |
| ENSMUSG00000067081 | Asb18 | 2.76E-08 | 2.077407 |
| ENSMUSG00000044594 | Serpinb3a | 5.80E-06 | 2.079432 |
| ENSMUSG00000044499 | Hs3st5 | 3.23E-11 | 2.080027 |
| ENSMUSG00000030713 | Klk7 | 1.62E-02 | 2.081978 |
| ENSMUSG00000043472 | Lce3d | 2.62E-02 | 2.084031 |
| ENSMUSG00000045019 | Acer1 | 2.71E-02 | 2.084468 |
| ENSMUSG00000047976 | Kcna1 | 2.86E-12 | 2.088442 |
| ENSMUSG00000032013 | Trim29 | 1.72E-10 | 2.088868 |
| ENSMUSG00000026327 | Serpinb11 | 1.47E-04 | 2.090204 |
| ENSMUSG00000026592 | Tex35 | 4.71E-09 | 2.091662 |
| ENSMUSG00000047216 | Cdh19 | 9.22E-24 | 2.093022 |
| ENSMUSG00000050830 | Vwc2 | 6.27E-05 | 2.093227 |
| ENSMUSG00000078907 | Fam186b | 4.10E-02 | 2.094674 |
| ENSMUSG00000016179 | Camk1g | 3.72E-02 | 2.098511 |
| ENSMUSG00000026407 | Cacna1s | 4.35E-21 | 2.098587 |
| ENSMUSG00000059994 | Fcrl1 | 2.55E-10 | 2.10039 |
| ENSMUSG00000027961 | Lrrc39 | 1.44E-08 | 2.10099 |
| ENSMUSG00000010064 | Slc38a3 | 3.35E-10 | 2.106878 |
| ENSMUSG00000031448 | Adprhl1 | 1.43E-16 | 2.1087 |
| ENSMUSG00000045776 | Lrtm1 | 1.67E-11 | 2.109699 |
| ENSMUSG00000053522 | Lgals7 | 5.24E-05 | 2.111441 |
| ENSMUSG00000069873 | 4930438A08Rik | 3.96E-05 | 2.11536 |
| ENSMUSG00000022053 | Ebf2 | 2.57E-28 | 2.116937 |
| ENSMUSG00000049598 | Vsig8 | 5.27E-03 | 2.118056 |
| ENSMUSG00000056966 | Gjc3 | 1.05E-02 | 2.118632 |
| ENSMUSG00000094814 | Gm21973 | 5.63E-03 | 2.122571 |
| ENSMUSG00000026621 | 1-Mar | 2.60E-02 | 2.123251 |
| ENSMUSG00000021255 | Esrrb | 5.97E-07 | 2.124457 |
| ENSMUSG00000109293 | Dcst2 | 1.99E-02 | 2.125564 |
| ENSMUSG00000026147 | Col9a1 | 2.36E-02 | 2.125738 |
| ENSMUSG00000067001 | Serpinb7 | 1.24E-02 | 2.128035 |
| ENSMUSG00000024076 | Vit | 7.49E-08 | 2.128736 |
| ENSMUSG00000048399 | Tprg | 5.27E-05 | 2.129159 |
| ENSMUSG00000048721 | Fndc9 | 4.81E-02 | 2.129168 |
| ENSMUSG00000028865 | Cd164l2 | 6.80E-05 | 2.131367 |
| ENSMUSG00000029195 | Klb | 1.90E-02 | 2.135638 |
| ENSMUSG00000032401 | Lctl | 4.28E-02 | 2.141541 |
| ENSMUSG00000051965 | Nanos2 | 2.43E-02 | 2.142158 |
| ENSMUSG00000028736 | Pax7 | 1.19E-02 | 2.142682 |
| ENSMUSG00000044041 | Krt13 | 2.13E-07 | 2.145789 |
| ENSMUSG00000035686 | Thrsp | 1.00E-21 | 2.14587 |
| ENSMUSG00000059956 | Serpinb12 | 6.27E-06 | 2.148141 |
| ENSMUSG00000030484 | Lypd5 | 1.18E-07 | 2.149653 |
| ENSMUSG00000057454 | Lypd3 | 8.72E-05 | 2.150403 |
| ENSMUSG00000025479 | Cyp2e1 | 2.34E-37 | 2.150779 |
| ENSMUSG00000097768 | 2310043M15Rik | 2.17E-03 | 2.151001 |
| ENSMUSG00000020007 | Il20ra | 8.82E-04 | 2.163851 |
| ENSMUSG00000027359 | Slc27a2 | 3.11E-04 | 2.163887 |

| | | | |
|---|---|---|---|
| ENSMUSG00000045539 | Sprr3 | 1.12E-09 | 2.16411 |
| ENSMUSG00000032807 | Alox12b | 2.37E-06 | 2.164244 |
| ENSMUSG00000032454 | Rbp2 | 2.18E-02 | 2.166851 |
| ENSMUSG00000050108 | Bpifc | 6.18E-03 | 2.167485 |
| ENSMUSG00000043460 | Elfn2 | 2.40E-02 | 2.16877 |
| ENSMUSG00000041984 | Rptn | 1.76E-11 | 2.170781 |
| ENSMUSG00000043468 | Adam30 | 5.96E-03 | 2.173031 |
| ENSMUSG00000073601 | Serpinb3c | 2.50E-03 | 2.175747 |
| ENSMUSG00000024471 | Myot | 2.51E-26 | 2.176575 |
| ENSMUSG00000065445 | Mir143 | 2.74E-02 | 2.17762 |
| ENSMUSG00000005716 | Pvalb | 2.86E-05 | 2.18435 |
| ENSMUSG00000022510 | Trp63 | 1.40E-18 | 2.189235 |
| ENSMUSG00000026984 | Il1f6 | 3.02E-03 | 2.189606 |
| ENSMUSG00000062044 | Lmtk3 | 3.45E-02 | 2.190846 |
| ENSMUSG00000052581 | Lrrtm4 | 1.15E-11 | 2.192309 |
| ENSMUSG00000045569 | Mc2r | 3.78E-02 | 2.192408 |
| ENSMUSG00000017309 | Cd300lg | 6.73E-18 | 2.195374 |
| ENSMUSG00000024526 | Cidea | 2.31E-06 | 2.199377 |
| ENSMUSG00000081689 | Gm9166 | 4.77E-02 | 2.201972 |
| ENSMUSG00000102574 | Gm33206 | 1.64E-02 | 2.204692 |
| ENSMUSG00000025064 | Col17a1 | 1.23E-11 | 2.208855 |
| ENSMUSG00000074445 | Sprr2a3 | 3.39E-03 | 2.210458 |
| ENSMUSG00000116659 | AC158985.1 | 1.19E-02 | 2.215346 |
| ENSMUSG00000027401 | Tgm3 | 2.06E-08 | 2.229316 |
| ENSMUSG00000026985 | Il1f8 | 3.43E-03 | 2.233766 |
| ENSMUSG00000028186 | Uox | 1.47E-02 | 2.235586 |
| ENSMUSG00000041991 | Hrnr | 2.86E-07 | 2.235634 |
| ENSMUSG00000024972 | Lgals12 | 4.44E-07 | 2.236885 |
| ENSMUSG00000054146 | Krt15 | 4.97E-10 | 2.240443 |
| ENSMUSG00000021509 | Slc25a48 | 9.46E-04 | 2.243699 |
| ENSMUSG00000044322 | Dsc1 | 1.20E-04 | 2.247548 |
| ENSMUSG00000059668 | Krt4 | 3.30E-08 | 2.252859 |
| ENSMUSG00000102698 | Gm37777 | 4.88E-02 | 2.254948 |
| ENSMUSG00000031725 | Ces1f | 2.39E-03 | 2.264564 |
| ENSMUSG00000028396 | 2310002L09Rik | 6.45E-05 | 2.264786 |
| ENSMUSG00000105128 | Gm42870 | 1.67E-03 | 2.265372 |
| ENSMUSG00000117222 | AC060761.1 | 1.12E-09 | 2.268436 |
| ENSMUSG00000039238 | Zfp750 | 1.91E-05 | 2.270028 |
| ENSMUSG00000028148 | Them5 | 4.51E-09 | 2.271649 |
| ENSMUSG00000027966 | Col11a1 | 1.37E-12 | 2.271918 |
| ENSMUSG00000117029 | CT030155.1 | 3.50E-04 | 2.274179 |
| ENSMUSG00000085615 | A330035P11Rik | 3.82E-03 | 2.27617 |
| ENSMUSG00000097364 | Gm26719 | 1.09E-03 | 2.281662 |
| ENSMUSG00000046764 | A530053G22Rik | 3.40E-04 | 2.285877 |
| ENSMUSG00000069441 | Dsg1a | 1.19E-12 | 2.293298 |
| ENSMUSG00000106871 | Gm3289 | 3.38E-02 | 2.300456 |
| ENSMUSG00000026413 | Pkp1 | 1.59E-11 | 2.301027 |
| ENSMUSG00000043430 | Psapl1 | 6.34E-09 | 2.302115 |
| ENSMUSG00000026983 | Il1f5 | 7.17E-04 | 2.303818 |
| ENSMUSG00000044359 | P2ry4 | 3.22E-04 | 2.308913 |
| ENSMUSG00000023041 | Krt6b | 2.05E-02 | 2.310622 |
| ENSMUSG00000094335 | Igkv1-117 | 4.70E-02 | 2.317249 |
| ENSMUSG00000040154 | Wfdc5 | 1.56E-04 | 2.317891 |

| | | | |
|---|---|---|---|
| ENSMUSG00000044505 | Lingo4 | 4.11E-02 | 2.324002 |
| ENSMUSG00000086155 | 9430041J12Rik | 1.78E-02 | 2.328883 |
| ENSMUSG00000051596 | Otop1 | 5.62E-03 | 2.333939 |
| ENSMUSG00000040283 | Btnl9 | 4.98E-05 | 2.341489 |
| ENSMUSG00000071019 | Sdr16c6 | 6.69E-04 | 2.347186 |
| ENSMUSG00000061928 | Dsg1b | 2.79E-10 | 2.356478 |
| ENSMUSG00000044453 | Ffar1 | 6.83E-03 | 2.359339 |
| ENSMUSG00000074771 | Ankef1 | 3.32E-11 | 2.367063 |
| ENSMUSG00000102713 | Gm29994 | 4.96E-02 | 2.37116 |
| ENSMUSG00000033268 | Duox1 | 1.02E-07 | 2.373757 |
| ENSMUSG00000087382 | Ctcflos | 3.00E-06 | 2.374574 |
| ENSMUSG00000022860 | Chodl | 2.17E-03 | 2.383702 |
| ENSMUSG00000097335 | Gm26563 | 3.11E-04 | 2.388002 |
| ENSMUSG00000074113 | Gm10629 | 2.59E-02 | 2.396626 |
| ENSMUSG00000061527 | Krt5 | 1.53E-10 | 2.400635 |
| ENSMUSG00000102788 | Gm38255 | 2.20E-02 | 2.411338 |
| ENSMUSG00000063130 | Calml3 | 1.42E-10 | 2.41536 |
| ENSMUSG00000027761 | Aadac | 1.39E-02 | 2.416545 |
| ENSMUSG00000021565 | Slc6a19 | 1.11E-07 | 2.422981 |
| ENSMUSG00000017607 | Tns4 | 1.21E-08 | 2.42721 |
| ENSMUSG00000086584 | Gm12002 | 9.27E-09 | 2.431887 |
| ENSMUSG00000056078 | Lipm | 3.36E-06 | 2.434023 |
| ENSMUSG00000048455 | Sprr1b | 7.22E-04 | 2.435726 |
| ENSMUSG00000074623 | Gm826 | 2.01E-05 | 2.447222 |
| ENSMUSG00000117322 | AC154766.4 | 1.42E-03 | 2.450311 |
| ENSMUSG00000074652 | Myh7b | 3.23E-04 | 2.464603 |
| ENSMUSG00000097074 | 4833428L15Rik | 3.94E-03 | 2.46861 |
| ENSMUSG00000044748 | Defb1 | 6.87E-04 | 2.475512 |
| ENSMUSG00000109656 | Gm45548 | 1.18E-02 | 2.476315 |
| ENSMUSG00000028236 | Sdr16c5 | 2.53E-03 | 2.481761 |
| ENSMUSG00000030972 | Acsm5 | 4.43E-02 | 2.484046 |
| ENSMUSG00000082141 | Gm11212 | 1.17E-02 | 2.488085 |
| ENSMUSG00000114898 | Gm49390 | 2.70E-02 | 2.494279 |
| ENSMUSG00000043795 | Prr33 | 9.14E-05 | 2.508198 |
| ENSMUSG00000095079 | Igha | 3.83E-18 | 2.559163 |
| ENSMUSG00000070719 | Pla2g4d | 2.65E-03 | 2.562518 |
| ENSMUSG00000057913 | Gm10032 | 3.14E-02 | 2.585521 |
| ENSMUSG00000023176 | Cpn2 | 4.45E-03 | 2.587907 |
| ENSMUSG00000068893 | Sprr2a2 | 1.16E-09 | 2.589158 |
| ENSMUSG00000042662 | Dusp15 | 9.84E-03 | 2.602635 |
| ENSMUSG00000067006 | Serpinb5 | 9.95E-09 | 2.602844 |
| ENSMUSG00000086032 | Gm15929 | 3.41E-02 | 2.608879 |
| ENSMUSG00000092222 | Gm20506 | 6.46E-03 | 2.617156 |
| ENSMUSG00000055301 | Adh7 | 5.78E-18 | 2.63083 |
| ENSMUSG00000102762 | 4930403P22Rik | 1.44E-03 | 2.635338 |
| ENSMUSG00000028427 | Aqp7 | 2.36E-05 | 2.650459 |
| ENSMUSG00000078664 | Sprr2a1 | 9.93E-05 | 2.651024 |
| ENSMUSG00000019936 | Epyc | 2.77E-02 | 2.652037 |
| ENSMUSG00000046095 | Krt32 | 1.20E-03 | 2.667941 |
| ENSMUSG00000114091 | Gm40655 | 5.12E-07 | 2.678247 |
| ENSMUSG00000099798 | Gm29168 | 1.20E-02 | 2.686459 |
| ENSMUSG00000027048 | Abcb11 | 3.96E-02 | 2.686539 |
| ENSMUSG00000032373 | Car12 | 1.07E-30 | 2.712374 |

| | | | |
|---|---|---|---|
| ENSMUSG00000035861 | Tmprss11b | 5.06E-03 | 2.728767 |
| ENSMUSG00000097850 | 4631405K08Rik | 6.26E-04 | 2.731215 |
| ENSMUSG00000030046 | Bmp10 | 2.22E-02 | 2.748688 |
| ENSMUSG00000079451 | Tmprss11g | 2.47E-07 | 2.766776 |
| ENSMUSG00000033765 | Calm4 | 2.29E-07 | 2.770286 |
| ENSMUSG00000074156 | Ces1h | 1.20E-08 | 2.790568 |
| ENSMUSG00000078964 | Ces1b | 7.10E-05 | 2.800191 |
| ENSMUSG00000029019 | Nppb | 4.80E-12 | 2.809285 |
| ENSMUSG00000074183 | Gsta1 | 1.83E-03 | 2.825282 |
| ENSMUSG00000045545 | Krt14 | 8.07E-21 | 2.845235 |
| ENSMUSG00000033849 | B3galt2 | 1.28E-06 | 2.854439 |
| ENSMUSG00000037942 | Crp | 4.26E-04 | 2.872187 |
| ENSMUSG00000032852 | Rspo4 | 3.66E-02 | 2.874898 |
| ENSMUSG00000028011 | Tdo2 | 5.25E-03 | 2.883112 |
| ENSMUSG00000053675 | Tgm5 | 1.57E-02 | 2.88705 |
| ENSMUSG00000114922 | Ppifos | 6.49E-06 | 2.892979 |
| ENSMUSG00000092586 | Ly6g6c | 8.58E-10 | 2.894076 |
| ENSMUSG00000075217 | 4833423E24Rik | 5.12E-09 | 2.895953 |
| ENSMUSG00000016327 | Atp1b4 | 2.46E-06 | 2.972894 |
| ENSMUSG00000028356 | Ambp | 1.17E-04 | 3.000966 |
| ENSMUSG00000031710 | Ucp1 | 1.67E-14 | 3.002674 |
| ENSMUSG00000116482 | Gm34152 | 1.61E-02 | 3.020639 |
| ENSMUSG00000033196 | Myh2 | 1.40E-20 | 3.128999 |
| ENSMUSG00000013936 | Myl2 | 1.78E-08 | 3.151791 |
| ENSMUSG00000111709 | Gm3776 | 1.40E-03 | 3.198663 |
| ENSMUSG00000059898 | Dsc3 | 1.21E-15 | 3.210725 |
| ENSMUSG00000111556 | Gm19299 | 4.63E-06 | 3.219853 |
| ENSMUSG00000058354 | Krt6a | 5.91E-07 | 3.268707 |
| ENSMUSG00000004872 | Pax3 | 4.21E-04 | 3.300239 |
| ENSMUSG00000106056 | 4930500L23Rik | 1.18E-03 | 3.320195 |
| ENSMUSG00000115207 | Gm49132 | 2.01E-04 | 3.358344 |
| ENSMUSG00000028001 | Fga | 2.45E-06 | 3.395634 |
| ENSMUSG00000056632 | Dsg3 | 1.17E-26 | 3.443911 |
| ENSMUSG00000027513 | Pck1 | 2.05E-42 | 3.544075 |
| ENSMUSG00000017950 | Hnf4a | 3.71E-04 | 3.786185 |
| ENSMUSG00000025003 | Cyp2c39 | 1.57E-03 | 4.069422 |
| ENSMUSG00000102098 | 2310016D03Rik | 8.49E-06 | 4.610042 |
| ENSMUSG00000056656 | Apol8 | 1.42E-18 | 4.685594 |
| ENSMUSG00000033860 | Fgg | 2.32E-12 | 4.837005 |
| ENSMUSG00000022875 | Kng1 | 3.51E-06 | 4.98915 |
| ENSMUSG00000025194 | Abcc2 | 1.51E-05 | 5.208255 |
| ENSMUSG00000054422 | Fabp1 | 9.90E-07 | 5.384976 |
| ENSMUSG00000021364 | Elovl2 | 1.66E-05 | 5.766093 |
| ENSMUSG00000001670 | Tat | 8.99E-06 | 8.225412 |
| ENSMUSG00000030131 | Mug2 | 4.90E-08 | 8.940782 |
| ENSMUSG00000030359 | Pzp | 5.53E-19 | 10.7158 |
| ENSMUSG00000028307 | Aldob | 4.66E-11 | 13.80692 |
| ENSMUSG00000020609 | Apob | 3.96E-21 | 14.75844 |
| ENSMUSG00000085154 | C130046K22Rik | 5.61E-09 | 14.96049 |
| ENSMUSG00000022868 | Ahsg | 1.17E-21 | 15.70773 |
| ENSMUSG00000035540 | Gc | 1.39E-15 | 17.68534 |
| ENSMUSG00000019932 | Kera | 9.46E-13 | 17.79842 |
| ENSMUSG00000003053 | Cyp2c29 | 3.02E-11 | 21.15165 |

| | | | |
|---|---|---|---|
| ENSMUSG00000059481 | Plg | 1.20E-08 | 22.09324 |
| ENSMUSG00000056035 | Cyp3a11 | 1.28E-13 | 54.83161 |
| ENSMUSG00000032083 | Apoa1 | 5.63E-16 | 55.84116 |
| ENSMUSG00000061808 | Ttr | 6.66E-07 | 82.413 |
| ENSMUSG00000059908 | Mug1 | 1.60E-17 | 99.55708 |
| ENSMUSG00000029630 | Cyp3a25 | 1.25E-06 | 123.9654 |
| ENSMUSG00000029368 | Alb | 2.26E-43 | 153.3651 |
| ENSMUSG00000025991 | Cps1 | 2.28E-07 | 180.3232 |
| ENSMUSG00000033831 | Fgb | 4.13E-09 | 409.5904 |

**Supplementary Table 9.** Murine differential gene expression in the lungs 6 h post-infection, 9-47-Ear vs 4559M. Genes with fold change (FC) greater than 1.5 and $p < 0.05$ are shown. FC values highlighted in blue = upregulated in 9-47-Ear, while values highlighted in red = upregulated in 4559M.

| **Gene Stable ID** | **Gene Name** | **padj** | **FC** |
|---|---|---|---|
| ENSMUSG00000096349 | *Gm22513* | 4.55E-03 | 2.606374 |
| ENSMUSG00000045932 | *Ifit2* | 6.67E-41 | 2.120213 |
| ENSMUSG00000003545 | *Fosb* | 1.28E-08 | 2.101081 |
| ENSMUSG00000094392 | *Gm3788* | 2.89E-05 | 2.057367 |
| ENSMUSG00000057596 | *Trim30d* | 5.07E-06 | 1.986075 |
| ENSMUSG00000000386 | *Mx1* | 2.11E-34 | 1.933773 |
| ENSMUSG00000020641 | *Rsad2* | 1.35E-13 | 1.923719 |
| ENSMUSG00000059434 | *Gckr* | 5.94E-06 | 1.899378 |
| ENSMUSG00000097312 | *Gm26870* | 9.81E-06 | 1.880663 |
| ENSMUSG00000060183 | *Cxcl11* | 4.50E-06 | 1.864222 |
| ENSMUSG00000035852 | *Misp* | 1.64E-10 | 1.805939 |
| ENSMUSG00000015437 | *Gzmb* | 7.02E-04 | 1.803683 |
| ENSMUSG00000108289 | *Gm44174* | 1.88E-06 | 1.793558 |
| ENSMUSG00000048003 | *Catsper4* | 2.11E-03 | 1.765394 |
| ENSMUSG00000052353 | *Cemip* | 5.51E-17 | 1.762085 |
| ENSMUSG00000084823 | *Gm17216* | 1.45E-02 | 1.726759 |
| ENSMUSG00000020638 | *Cmpk2* | 4.84E-24 | 1.693086 |
| ENSMUSG00000103824 | *Gm38177* | 1.68E-02 | 1.688716 |
| ENSMUSG00000031757 | *Mt4* | 3.02E-02 | 1.675168 |
| ENSMUSG00000037962 | *Rflna* | 2.29E-02 | 1.671596 |
| ENSMUSG00000114553 | *Gm7644* | 1.01E-02 | 1.623832 |
| ENSMUSG00000026826 | *Nr4a2* | 1.18E-04 | 1.600445 |
| ENSMUSG00000059832 | *Kprp* | 2.90E-03 | 1.592978 |
| ENSMUSG00000014543 | *Klra17* | 2.61E-06 | 1.58641 |
| ENSMUSG00000063651 | *Cnfn* | 4.88E-03 | 1.573752 |
| ENSMUSG00000116660 | *CT030736.1* | 1.20E-05 | 1.572765 |
| ENSMUSG00000028031 | *Dkk2* | 3.03E-04 | 1.572515 |
| ENSMUSG00000059230 | *Defb4* | 2.16E-02 | 1.569063 |
| ENSMUSG00000086196 | *Gm13571* | 2.78E-02 | 1.568524 |
| ENSMUSG00000067767 | *Clec4b2* | 1.74E-02 | 1.568218 |
| ENSMUSG00000073902 | *Gm1966* | 8.80E-07 | 1.554048 |
| ENSMUSG00000111229 | *Gm39323* | 1.70E-02 | 1.52393 |

| | | | |
|---|---|---|---|
| ENSMUSG00000034855 | *Cxcl10* | 3.16E-04 | 1.523329 |
| ENSMUSG00000070866 | *Zfp804a* | 2.31E-02 | 1.521761 |
| ENSMUSG00000097134 | *1110002J07Rik* | 1.21E-02 | 1.519142 |
| ENSMUSG00000032502 | *Stac* | 4.38E-02 | 1.517901 |
| ENSMUSG00000043664 | *Tmem221* | 3.80E-02 | 1.516904 |
| ENSMUSG00000086213 | *A330040F15Rik* | 1.39E-04 | 1.503579 |
| ENSMUSG00000096349 | *Gm22513* | 4.55E-03 | 2.606374 |
| ENSMUSG00000045932 | *Ifit2* | 6.67E-41 | 2.120213 |
| ENSMUSG00000003545 | *Fosb* | 1.28E-08 | 2.101081 |
| ENSMUSG00000094392 | *Gm3788* | 2.89E-05 | 2.057367 |
| ENSMUSG00000057596 | *Trim30d* | 5.07E-06 | 1.986075 |
| ENSMUSG00000000386 | *Mx1* | 2.11E-34 | 1.933773 |
| ENSMUSG00000020641 | *Rsad2* | 1.35E-13 | 1.923719 |
| ENSMUSG00000059434 | *Gckr* | 5.94E-06 | 1.899378 |
| ENSMUSG00000097312 | *Gm26870* | 9.81E-06 | 1.880663 |
| ENSMUSG00000060183 | *Cxcl11* | 4.50E-06 | 1.864222 |
| ENSMUSG00000035852 | *Misp* | 1.64E-10 | 1.805939 |
| ENSMUSG00000015437 | *Gzmb* | 7.02E-04 | 1.803683 |
| ENSMUSG00000108289 | *Gm44174* | 1.88E-06 | 1.793558 |
| ENSMUSG00000048003 | *Catsper4* | 2.11E-03 | 1.765394 |
| ENSMUSG00000052353 | *Cemip* | 5.51E-17 | 1.762085 |
| ENSMUSG00000084823 | *Gm17216* | 1.45E-02 | 1.726759 |
| ENSMUSG00000020638 | *Cmpk2* | 4.84E-24 | 1.693086 |
| ENSMUSG00000103824 | *Gm38177* | 1.68E-02 | 1.688716 |
| ENSMUSG00000031757 | *Mt4* | 3.02E-02 | 1.675168 |
| ENSMUSG00000037962 | *Rflna* | 2.29E-02 | 1.671596 |
| ENSMUSG00000114553 | *Gm7644* | 1.01E-02 | 1.623832 |
| ENSMUSG00000026826 | *Nr4a2* | 1.18E-04 | 1.600445 |
| ENSMUSG00000059832 | *Kprp* | 2.90E-03 | 1.592978 |
| ENSMUSG00000014543 | *Klra17* | 2.61E-06 | 1.58641 |
| ENSMUSG00000063651 | *Cnfn* | 4.88E-03 | 1.573752 |
| ENSMUSG00000116660 | *CT030736.1* | 1.20E-05 | 1.572765 |
| ENSMUSG00000028031 | *Dkk2* | 3.03E-04 | 1.572515 |
| ENSMUSG00000059230 | *Defb4* | 2.16E-02 | 1.569063 |
| ENSMUSG00000086196 | *Gm13571* | 2.78E-02 | 1.568524 |
| ENSMUSG00000067767 | *Clec4b2* | 1.74E-02 | 1.568218 |
| ENSMUSG00000073902 | *Gm1966* | 8.80E-07 | 1.554048 |
| ENSMUSG00000111229 | *Gm39323* | 1.70E-02 | 1.52393 |
| ENSMUSG00000034855 | *Cxcl10* | 3.16E-04 | 1.523329 |
| ENSMUSG00000070866 | *Zfp804a* | 2.31E-02 | 1.521761 |
| ENSMUSG00000097134 | *1110002J07Rik* | 1.21E-02 | 1.519142 |
| ENSMUSG00000032502 | *Stac* | 4.38E-02 | 1.517901 |
| ENSMUSG00000043664 | *Tmem221* | 3.80E-02 | 1.516904 |
| ENSMUSG00000086213 | *A330040F15Rik* | 1.39E-04 | 1.503579 |
| ENSMUSG00000101930 | *Gm5441* | 8.04E-04 | 1.501768 |
| ENSMUSG00000116031 | *D030024E09Rik* | 1.14E-03 | 1.505026 |
| ENSMUSG00000109157 | *Gm44829* | 2.31E-02 | 1.513042 |
| ENSMUSG00000047976 | *Kcna1* | 1.32E-03 | 1.515831 |
| ENSMUSG00000098708 | *Gm27252* | 9.38E-03 | 1.516922 |
| ENSMUSG00000066510 | *Ankdd1a* | 2.65E-04 | 1.517129 |
| ENSMUSG00000096084 | *Gm21850* | 3.46E-03 | 1.51884 |
| ENSMUSG00000021259 | *Cyp46a1* | 4.91E-02 | 1.520911 |
| ENSMUSG00000056569 | *Mpz* | 1.47E-06 | 1.521436 |

| | | | |
|---|---|---|---|
| ENSMUSG00000092267 | *Gm20417* | 4.50E-03 | 1.523107 |
| ENSMUSG00000030350 | *Prmt8* | 2.14E-06 | 1.527589 |
| ENSMUSG00000108929 | *Cc2d2b* | 2.94E-02 | 1.536074 |
| ENSMUSG00000000197 | *Nalcn* | 3.23E-05 | 1.543563 |
| ENSMUSG00000089844 | *A530032D15Rik* | 2.07E-05 | 1.544398 |
| ENSMUSG00000054582 | *Pabpc1l* | 5.95E-05 | 1.549878 |
| ENSMUSG00000090877 | *Hspa1b* | 1.31E-04 | 1.554299 |
| ENSMUSG00000094125 | *Gm13698* | 1.01E-02 | 1.558384 |
| ENSMUSG00000094336 | *Gm13693* | 1.01E-02 | 1.558384 |
| ENSMUSG00000096484 | *Gm13696* | 1.01E-02 | 1.558384 |
| ENSMUSG00000096337 | *Gm13694* | 1.01E-02 | 1.558384 |
| ENSMUSG00000095824 | *Gm13697* | 1.01E-02 | 1.558384 |
| ENSMUSG00000079247 | *Gm13691* | 1.01E-02 | 1.558384 |
| ENSMUSG00000020698 | *Cct6b* | 4.95E-02 | 1.558508 |
| ENSMUSG00000097071 | *4930544I03Rik* | 2.71E-03 | 1.558902 |
| ENSMUSG00000111818 | *Gm17749* | 3.50E-04 | 1.559008 |
| ENSMUSG00000096729 | *Gm13695* | 9.65E-03 | 1.560095 |
| ENSMUSG00000000157 | *Itgb2l* | 1.35E-10 | 1.563713 |
| ENSMUSG00000102590 | *Mannr* | 3.05E-02 | 1.566127 |
| ENSMUSG00000100457 | *D830032E09Rik* | 4.66E-02 | 1.566802 |
| ENSMUSG00000069830 | *Nlrp1a* | 3.96E-02 | 1.567104 |
| ENSMUSG00000027513 | *Pck1* | 9.84E-05 | 1.579451 |
| ENSMUSG00000109002 | *Gm38405* | 3.97E-02 | 1.583287 |
| ENSMUSG00000087113 | *Gm11714* | 7.53E-03 | 1.586987 |
| ENSMUSG00000019577 | *Pdk4* | 7.99E-35 | 1.589981 |
| ENSMUSG00000109032 | *Gm7972* | 1.78E-03 | 1.592848 |
| ENSMUSG00000017309 | *Cd300lg* | 1.27E-05 | 1.593726 |
| ENSMUSG00000042895 | *Abra* | 1.07E-04 | 1.59373 |
| ENSMUSG00000093482 | *Gm20619* | 4.17E-04 | 1.595111 |
| ENSMUSG00000025038 | *Efhc2* | 1.41E-03 | 1.595115 |
| ENSMUSG00000028989 | *Angptl7* | 2.82E-04 | 1.597595 |
| ENSMUSG00000028573 | *Fggy* | 6.11E-03 | 1.598726 |
| ENSMUSG00000096351 | *Samd11* | 2.77E-03 | 1.599425 |
| ENSMUSG00000001482 | *Def8* | 1.63E-08 | 1.601844 |
| ENSMUSG00000103620 | *Gm37359* | 6.80E-03 | 1.602521 |
| ENSMUSG00000034755 | *Pcdh11x* | 2.76E-04 | 1.607959 |
| ENSMUSG00000112707 | *D830005E20Rik* | 3.55E-07 | 1.612262 |
| ENSMUSG00000025716 | *Myo3a* | 1.00E-02 | 1.616705 |
| ENSMUSG00000039376 | *Synpo2l* | 3.29E-10 | 1.617835 |
| ENSMUSG00000043487 | *Acot6* | 4.05E-02 | 1.619862 |
| ENSMUSG00000040740 | *Slc25a34* | 5.61E-04 | 1.620673 |
| ENSMUSG00000083898 | *Gm7340* | 2.52E-05 | 1.62131 |
| ENSMUSG00000047216 | *Cdh19* | 3.15E-09 | 1.621717 |
| ENSMUSG00000087436 | *Gm16156* | 3.54E-02 | 1.622122 |
| ENSMUSG00000046748 | *Tmem45a2* | 5.95E-04 | 1.63198 |
| ENSMUSG00000042451 | *Mybph* | 3.29E-11 | 1.633192 |
| ENSMUSG00000112895 | *Gm47567* | 9.03E-03 | 1.63877 |
| ENSMUSG00000031626 | *Sorbs2* | 2.96E-20 | 1.6396 |
| ENSMUSG00000030607 | *Acan* | 1.15E-02 | 1.640047 |
| ENSMUSG00000004668 | *Abca13* | 4.50E-22 | 1.64663 |
| ENSMUSG00000025892 | *Gria4* | 2.09E-02 | 1.648351 |
| ENSMUSG00000079243 | *Xirp1* | 1.13E-12 | 1.649967 |
| ENSMUSG00000025172 | *Ankrd2* | 1.14E-04 | 1.655491 |

| | | | |
|---|---|---|---|
| ENSMUSG00000032496 | *Ltf* | 1.64E-24 | 1.662871 |
| ENSMUSG00000087353 | *Gm13727* | 9.58E-05 | 1.669227 |
| ENSMUSG00000084783 | *Gm15419* | 2.41E-02 | 1.67799 |
| ENSMUSG00000043789 | *Vwce* | 2.45E-03 | 1.682827 |
| ENSMUSG00000092397 | *C130080G10Rik* | 3.31E-06 | 1.687017 |
| ENSMUSG00000052581 | *Lrrtm4* | 1.05E-04 | 1.698365 |
| ENSMUSG00000041476 | *Smpx* | 1.39E-04 | 1.709086 |
| ENSMUSG00000103288 | *Gm37273* | 2.47E-03 | 1.713415 |
| ENSMUSG00000091971 | *Hspa1a* | 2.89E-05 | 1.735291 |
| ENSMUSG00000027014 | *Cwc22* | 4.55E-06 | 1.737258 |
| ENSMUSG00000038357 | *Camp* | 1.72E-04 | 1.743837 |
| ENSMUSG00000086354 | *Gm13938* | 1.32E-12 | 1.744173 |
| ENSMUSG00000048388 | *Fam171b* | 3.75E-15 | 1.772987 |
| ENSMUSG00000091119 | *Ccdc152* | 7.27E-04 | 1.775671 |
| ENSMUSG00000092201 | *A530058N18Rik* | 3.96E-09 | 1.775799 |
| ENSMUSG00000027966 | *Col11a1* | 1.22E-05 | 1.791424 |
| ENSMUSG00000087362 | *Gm13710* | 7.49E-07 | 1.797954 |
| ENSMUSG00000028865 | *Cd164l2* | 7.33E-03 | 1.840898 |
| ENSMUSG00000026592 | *Tex35* | 4.20E-06 | 1.902222 |
| ENSMUSG00000050830 | *Vwc2* | 2.01E-03 | 1.925254 |
| ENSMUSG00000020848 | *Doc2b* | 7.33E-16 | 1.935334 |
| ENSMUSG00000021071 | *Trim9* | 7.02E-07 | 1.951751 |
| ENSMUSG00000031710 | *Ucp1* | 7.62E-05 | 1.957941 |
| ENSMUSG00000042567 | *Nek10* | 5.83E-20 | 1.985917 |
| ENSMUSG00000105361 | *AY036118* | 2.16E-04 | 2.072411 |
| ENSMUSG00000032484 | *Ngp* | 7.92E-53 | 2.07311 |
| ENSMUSG00000096960 | *A230028O05Rik* | 1.84E-11 | 2.079473 |
| ENSMUSG00000029019 | *Nppb* | 1.68E-06 | 2.216413 |
| ENSMUSG00000020609 | *Apob* | 5.06E-03 | 2.792234 |
| ENSMUSG00000056656 | *Apol8* | 3.96E-09 | 3.055558 |
| ENSMUSG00000013936 | *Myl2* | 4.89E-18 | 5.494857 |
| ENSMUSG00000029368 | *Alb* | 2.47E-14 | 19.45594 |

**Supplementary Table 10.** Murine differential gene expression in the lungs 6 h post-infection, 9-47M vs 4559. Genes with fold change (FC) greater than 1.5 and $p < 0.05$ are shown. FC values highlighted in blue = upregulated in 9-47M, while values highlighted in red = upregulated in 4559.

| Gene Stable ID | Gene Name | padj | FC |
|---|---|---|---|
| ENSMUSG00000096349 | *Gm22513* | 4.24E-09 | 5.203286 |
| ENSMUSG00000086468 | *Etaa1os* | 3.06E-03 | 4.383019 |
| ENSMUSG00000071679 | *Rtl4* | 2.82E-18 | 4.145063 |
| ENSMUSG00000097554 | *Gm26825* | 3.15E-02 | 3.419807 |
| ENSMUSG00000031551 | *Ido1* | 1.65E-05 | 3.222093 |
| ENSMUSG00000085465 | *Gm15347* | 7.57E-04 | 3.20346 |
| ENSMUSG00000089874 | *9230117E06Rik* | 4.64E-02 | 3.119926 |
| ENSMUSG00000112246 | *Gm40761* | 1.67E-03 | 3.067346 |
| ENSMUSG00000046242 | *Nme9* | 5.99E-06 | 3.059333 |
| ENSMUSG00000105361 | *AY036118* | 7.94E-06 | 3.053392 |
| ENSMUSG00000001494 | *Sost* | 8.97E-05 | 2.742959 |
| ENSMUSG00000030114 | *Klrg1* | 6.18E-03 | 2.73067 |

| | | | |
|---|---|---|---|
| ENSMUSG00000092591 | *Gm20429* | 2.74E-03 | 2.725606 |
| ENSMUSG00000102608 | *Gm37267* | 3.18E-02 | 2.662128 |
| ENSMUSG00000064941 | *Gm23238* | 4.50E-07 | 2.561069 |
| ENSMUSG00000091694 | *Apol11b* | 3.42E-02 | 2.557668 |
| ENSMUSG00000076612 | *Ighg2c* | 1.66E-02 | 2.526225 |
| ENSMUSG00000117074 | *AC121299.1* | 9.05E-04 | 2.484582 |
| ENSMUSG00000063594 | *Gng8* | 1.64E-02 | 2.483359 |
| ENSMUSG00000102795 | *Gm6236* | 3.35E-02 | 2.457189 |
| ENSMUSG00000095616 | *Gm26244* | 1.72E-04 | 2.370041 |
| ENSMUSG00000025578 | *Cbx8* | 3.26E-04 | 2.32664 |
| ENSMUSG00000086843 | *E030013I19Rik* | 1.04E-07 | 2.322747 |
| ENSMUSG00000094306 | *Gm24924* | 2.98E-04 | 2.262227 |
| ENSMUSG00000115432 | *D130009I18Rik* | 4.24E-03 | 2.229274 |
| ENSMUSG00000070530 | *Wfdc16* | 2.71E-02 | 2.177114 |
| ENSMUSG00000095969 | *Rnu1a1* | 3.65E-04 | 2.169236 |
| ENSMUSG00000101603 | *Gm28730* | 4.48E-02 | 2.160864 |
| ENSMUSG00000115307 | *Gm49313* | 2.36E-04 | 2.155958 |
| ENSMUSG00000096659 | *Gm25679* | 1.87E-06 | 2.155429 |
| ENSMUSG00000093815 | *Gm26444* | 1.87E-06 | 2.155429 |
| ENSMUSG00000090667 | *Gm765* | 4.68E-02 | 2.133759 |
| ENSMUSG00000082292 | *Gm12250* | 2.39E-09 | 2.123222 |
| ENSMUSG00000080797 | *Gm15760* | 1.81E-02 | 2.120402 |
| ENSMUSG00000108481 | *Gm33248* | 1.01E-19 | 2.098355 |
| ENSMUSG00000070720 | *Tmem200b* | 5.47E-05 | 2.088404 |
| ENSMUSG00000069300 | *Hist1h2bj* | 3.40E-02 | 2.08459 |
| ENSMUSG00000065778 | *Gm22154* | 4.98E-02 | 2.079887 |
| ENSMUSG00000094826 | *Gm23804* | 9.42E-06 | 2.058273 |
| ENSMUSG00000091956 | *C2cd4b* | 2.21E-02 | 2.046617 |
| ENSMUSG00000052026 | *Slc6a7* | 1.35E-06 | 2.03982 |
| ENSMUSG00000114969 | *Hba-ps3* | 8.89E-03 | 2.039639 |
| ENSMUSG00000109491 | *Gm15396* | 4.44E-03 | 2.0348 |
| ENSMUSG00000046470 | *Sox18* | 7.42E-22 | 2.034619 |
| ENSMUSG00000087943 | *Gm24245* | 4.06E-06 | 2.030591 |
| ENSMUSG00000112831 | *AC167229.1* | 4.48E-02 | 1.969926 |
| ENSMUSG00000044952 | *Kctd21* | 6.08E-12 | 1.954796 |
| ENSMUSG00000116809 | *AC122538.1* | 1.49E-02 | 1.952285 |
| ENSMUSG00000072599 | *Ear-ps2* | 4.25E-04 | 1.942615 |
| ENSMUSG00000095260 | *Gm25890* | 2.93E-03 | 1.936872 |
| ENSMUSG00000094812 | *Gm22614* | 2.93E-03 | 1.936872 |
| ENSMUSG00000048988 | *Elfn1* | 3.07E-16 | 1.934531 |
| ENSMUSG00000104965 | *Gm43437* | 2.42E-02 | 1.930614 |
| ENSMUSG00000071661 | *Zbtb3* | 1.99E-02 | 1.929765 |
| ENSMUSG00000035299 | *Mid1* | 1.10E-04 | 1.921743 |
| ENSMUSG00000079330 | *Lemd1* | 3.53E-07 | 1.913457 |
| ENSMUSG00000060981 | *Hist1h4h* | 2.00E-05 | 1.909585 |
| ENSMUSG00000068696 | *Gpr88* | 1.30E-02 | 1.908856 |
| ENSMUSG00000081684 | *Rps2-ps13* | 6.39E-08 | 1.908057 |
| ENSMUSG00000064063 | *BC048507* | 1.73E-04 | 1.907851 |
| ENSMUSG00000096205 | *Gm22068* | 2.30E-03 | 1.907755 |
| ENSMUSG00000082192 | *Gm14719* | 7.60E-03 | 1.90406 |
| ENSMUSG00000092118 | *Fancf* | 3.09E-02 | 1.903835 |
| ENSMUSG00000050578 | *Mmp13* | 1.52E-03 | 1.899788 |
| ENSMUSG00000064923 | *Gm22042* | 3.99E-03 | 1.899281 |

| | | | |
|---|---|---|---|
| ENSMUSG00000055945 | *Prr18* | 4.95E-05 | 1.895915 |
| ENSMUSG00000096214 | *Gm22634* | 3.91E-05 | 1.891642 |
| ENSMUSG00000096206 | *Gm22317* | 3.91E-05 | 1.891642 |
| ENSMUSG00000063681 | *Crb1* | 3.96E-02 | 1.887394 |
| ENSMUSG00000116097 | *Gm36738* | 2.84E-03 | 1.876667 |
| ENSMUSG00000095701 | *Gm24830* | 4.71E-03 | 1.8723 |
| ENSMUSG00000096838 | *Gm26232* | 4.71E-03 | 1.8723 |
| ENSMUSG00000065773 | *Rnu1b6* | 4.71E-03 | 1.8723 |
| ENSMUSG00000107794 | *Gm44095* | 1.32E-03 | 1.871248 |
| ENSMUSG00000083737 | *Prdx6-ps2* | 1.41E-03 | 1.870411 |
| ENSMUSG00000035435 | *Abca17* | 1.48E-13 | 1.866967 |
| ENSMUSG00000038233 | *Fam198a* | 3.68E-07 | 1.865708 |
| ENSMUSG00000069305 | *Hist1h4n* | 9.20E-03 | 1.855769 |
| ENSMUSG00000041479 | *Syt15* | 1.78E-07 | 1.853036 |
| ENSMUSG00000112969 | *Gm48496* | 9.76E-03 | 1.84879 |
| ENSMUSG00000032357 | *Tinag* | 3.44E-07 | 1.84595 |
| ENSMUSG00000085154 | *C130046K22Rik* | 1.34E-02 | 1.844771 |
| ENSMUSG00000114028 | *9630002D21Rik* | 4.31E-03 | 1.844134 |
| ENSMUSG00000036264 | *Fstl4* | 3.58E-03 | 1.839891 |
| ENSMUSG00000108580 | *Gm39094* | 5.20E-27 | 1.83939 |
| ENSMUSG00000095580 | *Rnu1b1* | 9.28E-03 | 1.834552 |
| ENSMUSG00000093834 | *Rnu1b2* | 9.28E-03 | 1.834552 |
| ENSMUSG00000060678 | *Hist1h4c* | 1.39E-05 | 1.826151 |
| ENSMUSG00000045410 | *Akr1e1* | 8.33E-04 | 1.825984 |
| ENSMUSG00000085297 | *Gm11651* | 3.62E-02 | 1.823187 |
| ENSMUSG00000110634 | *Gm45895* | 4.27E-02 | 1.812488 |
| ENSMUSG00000035528 | *Npffr2* | 8.19E-03 | 1.809944 |
| ENSMUSG00000088246 | *Gm25911* | 1.52E-07 | 1.809528 |
| ENSMUSG00000086368 | *Gm13830* | 4.51E-04 | 1.80628 |
| ENSMUSG00000042988 | *Notum* | 1.04E-02 | 1.800697 |
| ENSMUSG00000087247 | *Alkal1* | 1.02E-14 | 1.799935 |
| ENSMUSG00000037279 | *Ovol2* | 4.81E-03 | 1.799681 |
| ENSMUSG00000060093 | *Hist1h4a* | 1.67E-03 | 1.792799 |
| ENSMUSG00000088609 | *Gm24187* | 7.72E-07 | 1.790795 |
| ENSMUSG00000085982 | *9530051G07Rik* | 8.90E-03 | 1.790428 |
| ENSMUSG00000029193 | *Cckar* | 6.00E-06 | 1.790204 |
| ENSMUSG00000086822 | *5330413P13Rik* | 2.14E-02 | 1.781756 |
| ENSMUSG00000067455 | *Hist1h4j* | 4.84E-02 | 1.781086 |
| ENSMUSG00000009900 | *Wnt3a* | 7.74E-05 | 1.780263 |
| ENSMUSG00000092240 | *Gm4246* | 6.92E-04 | 1.769107 |
| ENSMUSG00000084998 | *Gm16279* | 3.51E-03 | 1.767919 |
| ENSMUSG00000053961 | *Ang5* | 3.61E-02 | 1.764068 |
| ENSMUSG00000027070 | *Lrp2* | 5.22E-34 | 1.760968 |
| ENSMUSG00000069306 | *Hist1h4m* | 3.39E-02 | 1.760173 |
| ENSMUSG00000085882 | *2610507I01Rik* | 1.69E-03 | 1.759883 |
| ENSMUSG00000092805 | *Gm26461* | 4.98E-03 | 1.756282 |
| ENSMUSG00000024925 | *Rnaseh2c* | 2.34E-03 | 1.756123 |
| ENSMUSG00000031831 | *Dnaaf1* | 3.63E-02 | 1.748301 |
| ENSMUSG00000052117 | *D630039A03Rik* | 1.92E-03 | 1.746534 |
| ENSMUSG00000031220 | *Awat2* | 4.93E-03 | 1.738885 |
| ENSMUSG00000101605 | *Ace3* | 9.36E-06 | 1.738835 |
| ENSMUSG00000106229 | *Gm19409* | 3.20E-02 | 1.738286 |
| ENSMUSG00000030043 | *Tacr1* | 5.14E-05 | 1.733899 |

| | | | |
|---|---|---|---|
| ENSMUSG00000101674 | *4930444A19Rik* | 4.33E-02 | 1.733428 |
| ENSMUSG00000103160 | *C130012C08Rik* | 4.54E-02 | 1.730458 |
| ENSMUSG00000047085 | *Lrrc4b* | 2.55E-04 | 1.729971 |
| ENSMUSG00000086596 | *Susd5* | 1.35E-05 | 1.729874 |
| ENSMUSG00000070167 | *Snora57* | 5.72E-04 | 1.729595 |
| ENSMUSG00000050188 | *Lsm10* | 1.05E-03 | 1.72668 |
| ENSMUSG00000037649 | *H2-DMa* | 1.45E-08 | 1.723012 |
| ENSMUSG00000086390 | *1810019D21Rik* | 4.92E-06 | 1.722576 |
| ENSMUSG00000020734 | *Grin2c* | 9.25E-03 | 1.722135 |
| ENSMUSG00000106373 | *Gm6522* | 1.37E-03 | 1.721283 |
| ENSMUSG00000100041 | *Gm5702* | 2.41E-02 | 1.71842 |
| ENSMUSG00000040936 | *Ulk4* | 1.20E-17 | 1.714171 |
| ENSMUSG00000046240 | *Hepacam* | 2.58E-05 | 1.711621 |
| ENSMUSG00000030041 | *M1ap* | 2.40E-08 | 1.711292 |
| ENSMUSG00000035504 | *Reep6* | 3.13E-04 | 1.70748 |
| ENSMUSG00000036295 | *Lrrn3* | 1.27E-03 | 1.70554 |
| ENSMUSG00000105790 | *Gm24105* | 3.99E-03 | 1.70285 |
| ENSMUSG00000053318 | *Slamf8* | 2.80E-02 | 1.700939 |
| ENSMUSG00000082571 | *Gm8737* | 7.77E-03 | 1.698276 |
| ENSMUSG00000034324 | *Tmem132c* | 9.33E-22 | 1.697915 |
| ENSMUSG00000064288 | *Hist1h4k* | 2.67E-02 | 1.697211 |
| ENSMUSG00000117037 | *CT010433.2* | 1.34E-02 | 1.69681 |
| ENSMUSG00000037628 | *Cdkn3* | 2.99E-02 | 1.696525 |
| ENSMUSG00000037962 | *Rflna* | 3.78E-03 | 1.692265 |
| ENSMUSG00000061482 | *Hist1h4d* | 3.50E-06 | 1.688716 |
| ENSMUSG00000050288 | *Fzd2* | 3.65E-04 | 1.688423 |
| ENSMUSG00000076258 | *Gm23935* | 4.28E-05 | 1.682922 |
| ENSMUSG00000092674 | *Gm24105* | 4.53E-03 | 1.682736 |
| ENSMUSG00000034532 | *Fbxo16* | 1.50E-02 | 1.679353 |
| ENSMUSG00000032502 | *Stac* | 5.86E-04 | 1.678126 |
| ENSMUSG00000072601 | *Ear1* | 1.06E-06 | 1.671809 |
| ENSMUSG00000105145 | *Gm47299* | 5.17E-03 | 1.670472 |
| ENSMUSG00000105788 | *Gm47295* | 5.18E-03 | 1.670415 |
| ENSMUSG00000035910 | *Dcdc2a* | 2.61E-02 | 1.666406 |
| ENSMUSG00000097399 | *Gm26555* | 2.67E-02 | 1.664266 |
| ENSMUSG00000060988 | *Galnt13* | 6.00E-07 | 1.662241 |
| ENSMUSG00000077323 | *Rnu11* | 3.05E-02 | 1.661727 |
| ENSMUSG00000039672 | *Kcne2* | 2.18E-02 | 1.658794 |
| ENSMUSG00000070873 | *Lilra5* | 5.27E-04 | 1.658722 |
| ENSMUSG00000030093 | *Wnt7a* | 1.03E-04 | 1.65745 |
| ENSMUSG00000022504 | *Ciita* | 2.11E-16 | 1.65614 |
| ENSMUSG00000042985 | *Upk3b* | 4.24E-09 | 1.655501 |
| ENSMUSG00000033053 | *1700028P14Rik* | 1.36E-02 | 1.654074 |
| ENSMUSG00000058057 | *Mettl7a3* | 3.23E-02 | 1.653564 |
| ENSMUSG00000049511 | *Htr1b* | 3.81E-04 | 1.649535 |
| ENSMUSG00000076281 | *Gm24270* | 3.95E-04 | 1.648636 |
| ENSMUSG00000079505 | *Gm11131* | 1.20E-03 | 1.646768 |
| ENSMUSG00000048696 | *Mex3d* | 4.84E-05 | 1.646018 |
| ENSMUSG00000027186 | *Elf5* | 3.61E-04 | 1.63763 |
| ENSMUSG00000106416 | *Gm5857* | 3.80E-02 | 1.636649 |
| ENSMUSG00000110380 | *Gm45332* | 1.47E-02 | 1.634963 |
| ENSMUSG00000097453 | *Gm26894* | 1.65E-02 | 1.633469 |
| ENSMUSG00000020493 | *Prr11* | 1.30E-02 | 1.633102 |

| | | | |
|---|---|---|---|
| ENSMUSG00000061171 | *Slc38a11* | 2.46E-02 | 1.631519 |
| ENSMUSG00000114566 | *Gm48500* | 2.22E-02 | 1.631147 |
| ENSMUSG00000080538 | *Gm25541* | 4.24E-02 | 1.631042 |
| ENSMUSG00000041431 | *Ccnb1* | 3.24E-02 | 1.628007 |
| ENSMUSG00000025789 | *St8sia2* | 7.94E-07 | 1.627729 |
| ENSMUSG00000090166 | *Ear10* | 5.45E-04 | 1.627571 |
| ENSMUSG00000063779 | *Chil4* | 4.43E-04 | 1.627486 |
| ENSMUSG00000062545 | *Tlr12* | 1.63E-02 | 1.627004 |
| ENSMUSG00000092702 | *Gm24514* | 2.88E-02 | 1.625445 |
| ENSMUSG00000030800 | *Prss8* | 1.50E-08 | 1.624743 |
| ENSMUSG00000060969 | *Irx1* | 1.52E-09 | 1.623286 |
| ENSMUSG00000027656 | *Wisp2* | 1.04E-26 | 1.618407 |
| ENSMUSG00000042210 | *Abhd14a* | 9.45E-03 | 1.615522 |
| ENSMUSG00000022097 | *Sftpc* | 1.49E-08 | 1.614898 |
| ENSMUSG00000037190 | *Cyb561d2* | 3.40E-02 | 1.614444 |
| ENSMUSG00000056665 | *Them6* | 2.04E-02 | 1.61301 |
| ENSMUSG00000085069 | *Gm13111* | 5.74E-03 | 1.611789 |
| ENSMUSG00000038094 | *Atp13a4* | 5.04E-06 | 1.611226 |
| ENSMUSG00000112468 | *Gm47720* | 4.04E-06 | 1.61105 |
| ENSMUSG00000106352 | *5033403H07Rik* | 1.21E-06 | 1.61053 |
| ENSMUSG00000023147 | *Wrb* | 8.63E-06 | 1.609411 |
| ENSMUSG00000043719 | *Col6a6* | 5.13E-12 | 1.607024 |
| ENSMUSG00000097805 | *Gm17473* | 1.31E-07 | 1.60527 |
| ENSMUSG00000099338 | *2810030D12Rik* | 6.11E-03 | 1.605182 |
| ENSMUSG00000098178 | *Gm42418* | 1.65E-03 | 1.604909 |
| ENSMUSG00000024810 | *Il33* | 1.71E-18 | 1.603739 |
| ENSMUSG00000026870 | *Cutal* | 1.49E-02 | 1.601874 |
| ENSMUSG00000036114 | *Rpp25l* | 1.15E-03 | 1.599792 |
| ENSMUSG00000074480 | *Mex3a* | 4.38E-02 | 1.598909 |
| ENSMUSG00000013089 | *Etv5* | 2.42E-22 | 1.598533 |
| ENSMUSG00000042766 | *Trim46* | 1.71E-03 | 1.598267 |
| ENSMUSG00000075020 | *Mir670hg* | 2.15E-03 | 1.596736 |
| ENSMUSG00000087424 | *5730405O15Rik* | 2.37E-02 | 1.595889 |
| ENSMUSG00000040841 | *Six5* | 3.01E-05 | 1.595572 |
| ENSMUSG00000027380 | *Acoxl* | 3.11E-06 | 1.593887 |
| ENSMUSG00000108218 | *Olfr1372-ps1* | 4.02E-05 | 1.592729 |
| ENSMUSG00000041380 | *Htr2c* | 1.13E-06 | 1.592272 |
| ENSMUSG00000019909 | *Fam162b* | 4.28E-02 | 1.591414 |
| ENSMUSG00000090704 | *Trp53-ps* | 2.03E-07 | 1.591127 |
| ENSMUSG00000022844 | *Pdia5* | 8.24E-13 | 1.590662 |
| ENSMUSG00000033595 | *Lgi3* | 2.28E-12 | 1.588201 |
| ENSMUSG00000065701 | *Rny1* | 4.19E-02 | 1.587797 |
| ENSMUSG00000037610 | *Kcnmb2* | 5.28E-08 | 1.587617 |
| ENSMUSG00000043895 | *S1pr2* | 1.57E-16 | 1.586409 |
| ENSMUSG00000102175 | *Gm6119* | 1.44E-02 | 1.583275 |
| ENSMUSG00000037846 | *Rtkn2* | 8.92E-07 | 1.58299 |
| ENSMUSG00000030329 | *Pianp* | 1.03E-02 | 1.58274 |
| ENSMUSG00000027932 | *Slc27a3* | 3.43E-03 | 1.582304 |
| ENSMUSG00000030732 | *Chrdl2* | 4.82E-02 | 1.581585 |
| ENSMUSG00000094786 | *Gm14403* | 1.30E-10 | 1.579975 |
| ENSMUSG00000107290 | *Gm43282* | 3.94E-02 | 1.578205 |
| ENSMUSG00000022615 | *Tymp* | 2.61E-03 | 1.57791 |
| ENSMUSG00000075389 | *2810410L24Rik* | 1.42E-04 | 1.575503 |

| | | | |
|---|---|---|---|
| ENSMUSG00000091780 | *Sco2* | 3.28E-03 | 1.574907 |
| ENSMUSG00000064945 | *Rny3* | 3.69E-02 | 1.573231 |
| ENSMUSG00000009114 | *2610028H24Rik* | 8.71E-05 | 1.569539 |
| ENSMUSG00000074283 | *Zfp109* | 6.83E-04 | 1.567793 |
| ENSMUSG00000019295 | *Tmem129* | 8.45E-07 | 1.564808 |
| ENSMUSG00000026941 | *Mamdc4* | 2.12E-05 | 1.563251 |
| ENSMUSG00000086549 | *Gm13648* | 7.89E-03 | 1.559935 |
| ENSMUSG00000055134 | *9130017K11Rik* | 1.15E-02 | 1.55658 |
| ENSMUSG00000033952 | *Aspm* | 2.58E-05 | 1.55636 |
| ENSMUSG00000003779 | *Kif20a* | 3.46E-02 | 1.556308 |
| ENSMUSG00000025035 | *Arl3* | 2.31E-07 | 1.556136 |
| ENSMUSG00000044030 | *Irf2bp1* | 2.69E-06 | 1.55373 |
| ENSMUSG00000093327 | *Mir5107* | 5.85E-09 | 1.552336 |
| ENSMUSG00000107349 | *Gm6655* | 1.81E-02 | 1.551154 |
| ENSMUSG00000106633 | *Gm43431* | 4.58E-02 | 1.551081 |
| ENSMUSG00000078794 | *Dact3* | 8.48E-05 | 1.549569 |
| ENSMUSG00000027954 | *Efna1* | 1.86E-10 | 1.54683 |
| ENSMUSG00000021700 | *Rab3c* | 3.56E-05 | 1.5459 |
| ENSMUSG00000025902 | *Sox17* | 9.62E-12 | 1.5454 |
| ENSMUSG00000078772 | *Gm12353* | 3.08E-08 | 1.545118 |
| ENSMUSG00000070473 | *Cldn3* | 4.38E-07 | 1.5441 |
| ENSMUSG00000069515 | *Lyz1* | 8.64E-14 | 1.543848 |
| ENSMUSG00000030796 | *Tead2* | 4.76E-10 | 1.543617 |
| ENSMUSG00000040809 | *Chil3* | 4.58E-05 | 1.543465 |
| ENSMUSG00000037101 | *Ttc29* | 7.26E-06 | 1.542686 |
| ENSMUSG00000040289 | *Hey1* | 6.55E-13 | 1.542366 |
| ENSMUSG00000022595 | *Lypd2* | 2.36E-02 | 1.542217 |
| ENSMUSG00000051817 | *Sox12* | 2.88E-02 | 1.54192 |
| ENSMUSG00000029053 | *Prkcz* | 4.18E-12 | 1.538465 |
| ENSMUSG00000113212 | *Gm47435* | 1.47E-02 | 1.538088 |
| ENSMUSG00000049625 | *Tifab* | 1.51E-07 | 1.537426 |
| ENSMUSG00000032224 | *Fam81a* | 6.07E-03 | 1.535556 |
| ENSMUSG00000112343 | *Sfta3-ps* | 1.96E-21 | 1.534461 |
| ENSMUSG00000027849 | *Syt6* | 2.09E-04 | 1.533147 |
| ENSMUSG00000024206 | *Rfx2* | 7.51E-12 | 1.530804 |
| ENSMUSG00000006567 | *Atp7b* | 5.51E-09 | 1.529838 |
| ENSMUSG00000073424 | *Cyp4f15* | 1.31E-02 | 1.529096 |
| ENSMUSG00000044081 | *Zfp85os* | 2.57E-02 | 1.528554 |
| ENSMUSG00000058396 | *Gpr182* | 1.82E-07 | 1.527896 |
| ENSMUSG00000086141 | *9030622O22Rik* | 1.39E-08 | 1.526052 |
| ENSMUSG00000029651 | *Mtus2* | 1.31E-14 | 1.524294 |
| ENSMUSG00000040009 | *Gnaz* | 1.48E-02 | 1.522347 |
| ENSMUSG00000089694 | *Nat8f7* | 3.13E-02 | 1.518666 |
| ENSMUSG00000034685 | *Fam171a2* | 4.78E-03 | 1.518575 |
| ENSMUSG00000041261 | *Car8* | 9.86E-22 | 1.517297 |
| ENSMUSG00000057123 | *Gja5* | 2.89E-12 | 1.517113 |
| ENSMUSG00000115420 | *AL732506.1* | 8.45E-03 | 1.516697 |
| ENSMUSG00000088088 | *Rmrp* | 8.45E-03 | 1.516697 |
| ENSMUSG00000081093 | *Gm12565* | 3.28E-02 | 1.515801 |
| ENSMUSG00000117046 | *AC122746.1* | 3.96E-02 | 1.515479 |
| ENSMUSG00000104114 | *Gm37297* | 5.73E-04 | 1.515439 |
| ENSMUSG00000021496 | *Pcbd2* | 8.78E-13 | 1.515151 |
| ENSMUSG00000028540 | *Dph2* | 7.17E-04 | 1.514882 |

| | | | |
|---|---|---|---|
| ENSMUSG00000043668 | *Tox3* | 3.57E-13 | 1.513681 |
| ENSMUSG00000111409 | *Gm49380* | 3.92E-07 | 1.511921 |
| ENSMUSG00000027424 | *Mgme1* | 1.40E-02 | 1.511053 |
| ENSMUSG00000103738 | *Gm37652* | 2.01E-02 | 1.50916 |
| ENSMUSG00000044217 | *Aqp5* | 2.95E-15 | 1.509144 |
| ENSMUSG00000027490 | *E2f1* | 4.24E-02 | 1.508243 |
| ENSMUSG00000001444 | *Tbx21* | 2.37E-04 | 1.504887 |
| ENSMUSG00000066319 | *Rtp3* | 3.35E-03 | 1.504256 |
| ENSMUSG00000025512 | *Chid1* | 8.49E-12 | 1.504081 |
| ENSMUSG00000009394 | *Syn2* | 4.75E-08 | 1.503605 |
| ENSMUSG00000115143 | *Fkbp1a-ps1* | 3.78E-04 | 1.502351 |
| ENSMUSG00000033282 | *Rpgrip1l* | 6.03E-09 | 1.502144 |
| ENSMUSG00000036192 | *Rorb* | 1.76E-04 | 1.50102 |
| ENSMUSG00000099553 | *Gm29538* | 1.36E-02 | 1.500991 |
| ENSMUSG00000072596 | *Ear2* | 2.39E-06 | 1.500831 |
| ENSMUSG00000090737 | *Gm6788* | 2.53E-03 | 1.500806 |
| ENSMUSG00000085043 | *Gm13184* | 3.64E-02 | 1.500172 |
| ENSMUSG00000030428 | *Ttyh1* | 1.19E-02 | 1.502203 |
| ENSMUSG00000050357 | *Carmil2* | 4.60E-05 | 1.502416 |
| ENSMUSG00000032089 | *Il10ra* | 4.81E-21 | 1.502903 |
| ENSMUSG00000030107 | *Usp18* | 1.51E-14 | 1.502976 |
| ENSMUSG00000001014 | *Icam4* | 1.79E-02 | 1.502992 |
| ENSMUSG00000030785 | *Cox6a2* | 2.61E-07 | 1.503108 |
| ENSMUSG00000079434 | *Neu2* | 3.87E-02 | 1.503483 |
| ENSMUSG00000105677 | *Gm43328* | 4.53E-05 | 1.503833 |
| ENSMUSG00000101751 | *Gm2427* | 5.37E-05 | 1.503916 |
| ENSMUSG00000113668 | *Gm47451* | 1.42E-03 | 1.50528 |
| ENSMUSG00000052374 | *Actn2* | 3.36E-22 | 1.50534 |
| ENSMUSG00000057897 | *Camk2b* | 6.39E-04 | 1.505487 |
| ENSMUSG00000105337 | *1700094M23Rik* | 2.95E-06 | 1.505662 |
| ENSMUSG00000042678 | *Myo15* | 2.07E-03 | 1.506355 |
| ENSMUSG00000033107 | *Rnf125* | 1.82E-24 | 1.50685 |
| ENSMUSG00000036390 | *Gadd45a* | 1.65E-09 | 1.507699 |
| ENSMUSG00000058470 | *Gm8369* | 1.45E-07 | 1.507965 |
| ENSMUSG00000032369 | *Plscr1* | 5.18E-17 | 1.508682 |
| ENSMUSG00000105355 | *Gm40117* | 4.51E-02 | 1.508939 |
| ENSMUSG00000040899 | *Ccr6* | 1.50E-05 | 1.509044 |
| ENSMUSG00000011305 | *Plin5* | 1.11E-03 | 1.509246 |
| ENSMUSG00000102343 | *Gm37381* | 4.52E-02 | 1.509423 |
| ENSMUSG00000078937 | *Cpt1b* | 1.25E-07 | 1.509518 |
| ENSMUSG00000021250 | *Fos* | 1.72E-12 | 1.509638 |
| ENSMUSG00000006763 | *Saal1* | 3.57E-13 | 1.509651 |
| ENSMUSG00000074452 | *Pate2* | 1.01E-06 | 1.509912 |
| ENSMUSG00000109700 | *Gm21123* | 5.59E-03 | 1.510051 |
| ENSMUSG00000030433 | *Sbk2* | 7.43E-06 | 1.510461 |
| ENSMUSG00000053040 | *Aph1c* | 2.31E-08 | 1.510621 |
| ENSMUSG00000101344 | *Gm29183* | 7.91E-11 | 1.510713 |
| ENSMUSG00000035208 | *Slfn8* | 1.32E-11 | 1.510878 |
| ENSMUSG00000024778 | *Fas* | 1.01E-25 | 1.5111 |
| ENSMUSG00000028862 | *Map3k6* | 1.35E-20 | 1.511475 |
| ENSMUSG00000034833 | *Tespa1* | 1.65E-09 | 1.512576 |
| ENSMUSG00000115344 | *Gm49364* | 3.50E-09 | 1.512709 |
| ENSMUSG00000057092 | *Fxyd3* | 1.24E-02 | 1.513268 |

| | | | |
|---|---|---|---|
| ENSMUSG00000034959 | *Rubcnl* | 4.29E-14 | 1.514234 |
| ENSMUSG00000034842 | *Art3* | 1.48E-18 | 1.514796 |
| ENSMUSG00000003134 | *Tbc1d8* | 2.32E-15 | 1.51505 |
| ENSMUSG00000048004 | *Tmem196* | 1.46E-02 | 1.516581 |
| ENSMUSG00000012123 | *Crybg2* | 6.60E-05 | 1.516896 |
| ENSMUSG00000112808 | *Gm4739* | 1.85E-05 | 1.516929 |
| ENSMUSG00000025271 | *Pfkfb1* | 3.17E-13 | 1.51768 |
| ENSMUSG00000091412 | *Gm2895* | 1.63E-06 | 1.518936 |
| ENSMUSG00000026535 | *Ifi202b* | 5.15E-11 | 1.518943 |
| ENSMUSG00000044244 | *Il20rb* | 2.54E-19 | 1.519567 |
| ENSMUSG00000056755 | *Grm7* | 9.05E-05 | 1.519614 |
| ENSMUSG00000085908 | *Gm14209* | 7.71E-04 | 1.519669 |
| ENSMUSG00000029377 | *Ereg* | 1.71E-06 | 1.520092 |
| ENSMUSG00000024590 | *Lmnb1* | 2.04E-16 | 1.521965 |
| ENSMUSG00000071342 | *Lsmem1* | 5.96E-06 | 1.522759 |
| ENSMUSG00000069808 | *Fam57a* | 1.08E-06 | 1.522864 |
| ENSMUSG00000109371 | *Gm44626* | 2.28E-02 | 1.523263 |
| ENSMUSG00000051650 | *B3gnt2* | 2.04E-28 | 1.524523 |
| ENSMUSG00000033065 | *Pfkm* | 5.41E-13 | 1.52463 |
| ENSMUSG00000087589 | *D430040D24Rik* | 2.69E-06 | 1.524663 |
| ENSMUSG00000092201 | *A530058N18Rik* | 1.19E-05 | 1.524752 |
| ENSMUSG00000030077 | *Chl1* | 1.64E-18 | 1.526046 |
| ENSMUSG00000038665 | *Dgki* | 1.02E-06 | 1.526881 |
| ENSMUSG00000032661 | *Oas3* | 2.11E-10 | 1.526922 |
| ENSMUSG00000104010 | *Gm37366* | 8.49E-03 | 1.527991 |
| ENSMUSG00000032338 | *Hcn4* | 1.08E-07 | 1.528197 |
| ENSMUSG00000021125 | *Arg2* | 6.25E-15 | 1.528694 |
| ENSMUSG00000109279 | *Gm45220* | 7.04E-07 | 1.52872 |
| ENSMUSG00000025804 | *Ccr1* | 1.42E-12 | 1.528934 |
| ENSMUSG00000058498 | *Rnf207* | 6.72E-03 | 1.530495 |
| ENSMUSG00000074625 | *Arhgap40* | 2.06E-03 | 1.530634 |
| ENSMUSG00000108929 | *Cc2d2b* | 1.62E-02 | 1.532023 |
| ENSMUSG00000014198 | *Zfp385c* | 4.22E-02 | 1.532252 |
| ENSMUSG00000035818 | *Plekhs1* | 3.36E-03 | 1.532308 |
| ENSMUSG00000035580 | *Kcnh8* | 6.34E-04 | 1.532597 |
| ENSMUSG00000051748 | *Wfdc21* | 7.29E-08 | 1.53281 |
| ENSMUSG00000038663 | *Fsd2* | 8.48E-09 | 1.533131 |
| ENSMUSG00000022523 | *Fgf12* | 2.76E-14 | 1.533583 |
| ENSMUSG00000037904 | *Ankrd9* | 6.45E-03 | 1.535904 |
| ENSMUSG00000104406 | *Gm38014* | 4.26E-03 | 1.536295 |
| ENSMUSG00000109764 | *Klkb1* | 2.91E-02 | 1.53632 |
| ENSMUSG00000026628 | *Atf3* | 2.70E-20 | 1.537981 |
| ENSMUSG00000029915 | *Clec5a* | 1.81E-14 | 1.538051 |
| ENSMUSG00000042254 | *Cilp* | 4.11E-06 | 1.538166 |
| ENSMUSG00000078963 | *Hsbp1l1* | 4.27E-02 | 1.538173 |
| ENSMUSG00000004360 | *9330159F19Rik* | 2.37E-04 | 1.53843 |
| ENSMUSG00000073489 | *Ifi204* | 9.86E-22 | 1.540317 |
| ENSMUSG00000024481 | *Lvrn* | 2.04E-11 | 1.541111 |
| ENSMUSG00000057246 | *BC051142* | 1.99E-03 | 1.541414 |
| ENSMUSG00000086564 | *Cd101* | 4.69E-12 | 1.541652 |
| ENSMUSG00000064340 | *mt-Tl1* | 4.10E-08 | 1.54264 |
| ENSMUSG00000032125 | *Robo4* | 1.62E-20 | 1.542785 |
| ENSMUSG00000044405 | *Adig* | 4.27E-02 | 1.543087 |

| | | | |
|---|---|---|---|
| ENSMUSG00000052212 | *Cd177* | 4.55E-16 | 1.543341 |
| ENSMUSG00000032355 | *Mlip* | 2.15E-13 | 1.543416 |
| ENSMUSG00000042828 | *Trim72* | 5.22E-08 | 1.543717 |
| ENSMUSG00000056300 | *Zfp981* | 1.45E-05 | 1.543885 |
| ENSMUSG00000004864 | *Mapk13* | 6.25E-10 | 1.546096 |
| ENSMUSG00000063129 | *Aldoart2* | 1.66E-05 | 1.54623 |
| ENSMUSG00000042010 | *Acacb* | 3.59E-23 | 1.54649 |
| ENSMUSG00000070354 | *Evi2* | 8.63E-14 | 1.546718 |
| ENSMUSG00000034990 | *Otoa* | 4.05E-03 | 1.547561 |
| ENSMUSG00000049093 | *Il23r* | 5.93E-07 | 1.54799 |
| ENSMUSG00000071691 | *Gm960* | 7.83E-05 | 1.548087 |
| ENSMUSG00000022026 | *Olfm4* | 7.52E-16 | 1.548559 |
| ENSMUSG00000026452 | *Syt2* | 1.23E-10 | 1.548817 |
| ENSMUSG00000061578 | *Ksr2* | 1.01E-03 | 1.54969 |
| ENSMUSG00000032816 | *Igdcc4* | 5.80E-05 | 1.549734 |
| ENSMUSG00000104728 | *Gm42462* | 1.51E-06 | 1.55063 |
| ENSMUSG00000098702 | *1500015A07Rik* | 1.10E-03 | 1.552174 |
| ENSMUSG00000064147 | *Rab44* | 7.19E-18 | 1.55308 |
| ENSMUSG00000092397 | *C130080G10Rik* | 3.48E-05 | 1.554863 |
| ENSMUSG00000003849 | *Nqo1* | 1.35E-03 | 1.555535 |
| ENSMUSG00000057054 | *Inca1* | 3.20E-03 | 1.557038 |
| ENSMUSG00000009292 | *Trpm2* | 8.56E-13 | 1.558139 |
| ENSMUSG00000001943 | *Vsig2* | 1.50E-12 | 1.560479 |
| ENSMUSG00000025701 | *Alox5* | 2.14E-20 | 1.561191 |
| ENSMUSG00000105053 | *Gm43064* | 2.35E-10 | 1.564013 |
| ENSMUSG00000068794 | *Col28a1* | 4.53E-05 | 1.564183 |
| ENSMUSG00000021950 | *Anxa8* | 2.41E-06 | 1.565581 |
| ENSMUSG00000113918 | *Gm6566* | 7.52E-03 | 1.565835 |
| ENSMUSG00000055027 | *Smyd1* | 9.41E-12 | 1.566011 |
| ENSMUSG00000047746 | *Fbxo40* | 6.40E-08 | 1.566184 |
| ENSMUSG00000016349 | *Eef1a2* | 2.02E-12 | 1.566216 |
| ENSMUSG00000034308 | *Sdr42e1* | 1.45E-03 | 1.566555 |
| ENSMUSG00000029204 | *Rhoh* | 1.15E-21 | 1.56695 |
| ENSMUSG00000021061 | *Sptb* | 7.50E-20 | 1.567256 |
| ENSMUSG00000020698 | *Cct6b* | 1.31E-02 | 1.567908 |
| ENSMUSG00000045102 | *Poln* | 7.37E-20 | 1.567926 |
| ENSMUSG00000020077 | *Srgn* | 3.93E-19 | 1.568811 |
| ENSMUSG00000090582 | *Gm17024* | 1.32E-06 | 1.570766 |
| ENSMUSG00000017830 | *Dhx58* | 5.45E-13 | 1.571673 |
| ENSMUSG00000044716 | *Dok7* | 1.96E-05 | 1.57206 |
| ENSMUSG00000070604 | *Vsig10l* | 5.39E-07 | 1.572936 |
| ENSMUSG00000053338 | *Tarm1* | 1.01E-19 | 1.573966 |
| ENSMUSG00000005800 | *Mmp8* | 2.88E-22 | 1.574036 |
| ENSMUSG00000113778 | *Gm40847* | 2.80E-02 | 1.574565 |
| ENSMUSG00000037572 | *Wdhd1* | 2.94E-07 | 1.574592 |
| ENSMUSG00000032561 | *Acpp* | 2.04E-16 | 1.575351 |
| ENSMUSG00000053914 | *Kdm4d* | 6.99E-07 | 1.57607 |
| ENSMUSG00000002769 | *Gnmt* | 4.11E-02 | 1.576947 |
| ENSMUSG00000021579 | *Lrrc14b* | 1.53E-02 | 1.577637 |
| ENSMUSG00000031452 | *1700029H14Rik* | 4.92E-11 | 1.578495 |
| ENSMUSG00000112707 | *D830005E20Rik* | 1.67E-07 | 1.578937 |
| ENSMUSG00000024517 | *Grp* | 1.13E-02 | 1.579229 |
| ENSMUSG00000030577 | *Cd22* | 1.23E-05 | 1.580173 |

| | | | |
|---|---|---|---|
| ENSMUSG00000051351 | *Zfp46* | 5.74E-27 | 1.580489 |
| ENSMUSG00000000204 | *Slfn4* | 1.72E-07 | 1.580569 |
| ENSMUSG00000062007 | *Hsh2d* | 6.70E-05 | 1.580819 |
| ENSMUSG00000070000 | *Fcho1* | 1.23E-04 | 1.580984 |
| ENSMUSG00000105572 | *Gm43300* | 8.23E-05 | 1.581967 |
| ENSMUSG00000027360 | *Hdc* | 5.77E-25 | 1.582111 |
| ENSMUSG00000024803 | *Ankrd1* | 1.68E-14 | 1.582117 |
| ENSMUSG00000031872 | *Bean1* | 2.56E-09 | 1.582908 |
| ENSMUSG00000108313 | *Gm45052* | 3.46E-02 | 1.583486 |
| ENSMUSG00000024210 | *Ip6k3* | 2.10E-09 | 1.583691 |
| ENSMUSG00000037849 | *Ifi206* | 8.40E-04 | 1.584723 |
| ENSMUSG00000050663 | *Trhde* | 4.90E-02 | 1.585677 |
| ENSMUSG00000039891 | *Txlnb* | 2.06E-14 | 1.58583 |
| ENSMUSG00000024593 | *Megf10* | 6.07E-05 | 1.585923 |
| ENSMUSG00000059901 | *Adamts14* | 1.44E-19 | 1.586361 |
| ENSMUSG00000053024 | *Cntn2* | 3.57E-02 | 1.587941 |
| ENSMUSG00000071561 | *BC100530* | 2.30E-07 | 1.588282 |
| ENSMUSG00000114138 | *Gm36423* | 1.66E-02 | 1.589727 |
| ENSMUSG00000043789 | *Vwce* | 3.78E-03 | 1.5899 |
| ENSMUSG00000042851 | *Zc3h6* | 1.02E-14 | 1.590111 |
| ENSMUSG00000116604 | *AC087898.2* | 3.16E-02 | 1.590457 |
| ENSMUSG00000010476 | *Ebf3* | 5.07E-07 | 1.590556 |
| ENSMUSG00000004609 | *Cd33* | 3.48E-28 | 1.592001 |
| ENSMUSG00000098875 | *Gm27211* | 1.13E-06 | 1.592609 |
| ENSMUSG00000029205 | *Chrna9* | 2.74E-02 | 1.592947 |
| ENSMUSG00000050840 | *Cdh20* | 1.45E-02 | 1.593388 |
| ENSMUSG00000006457 | *Actn3* | 7.83E-07 | 1.59497 |
| ENSMUSG00000104348 | *Gm37691* | 1.06E-04 | 1.595263 |
| ENSMUSG00000024076 | *Vit* | 1.11E-03 | 1.595268 |
| ENSMUSG00000024972 | *Lgals12* | 4.52E-03 | 1.595515 |
| ENSMUSG00000040055 | *Gjb6* | 2.71E-03 | 1.595537 |
| ENSMUSG00000049336 | *Tenm2* | 1.62E-10 | 1.596005 |
| ENSMUSG00000104213 | *Ighd* | 3.47E-05 | 1.596239 |
| ENSMUSG00000022383 | *Ppara* | 2.03E-15 | 1.597777 |
| ENSMUSG00000054169 | *Ceacam10* | 2.78E-06 | 1.598518 |
| ENSMUSG00000109232 | *Gm44577* | 1.35E-02 | 1.598735 |
| ENSMUSG00000035692 | *Isg15* | 6.47E-15 | 1.601526 |
| ENSMUSG00000032523 | *Hhatl* | 3.08E-03 | 1.601819 |
| ENSMUSG00000040747 | *Cd53* | 9.31E-18 | 1.602191 |
| ENSMUSG00000053469 | *Tg* | 4.33E-17 | 1.602498 |
| ENSMUSG00000059974 | *Ntm* | 1.81E-26 | 1.602733 |
| ENSMUSG00000018425 | *Dhx40* | 3.14E-21 | 1.604037 |
| ENSMUSG00000075122 | *Cd80* | 1.39E-21 | 1.605419 |
| ENSMUSG00000002831 | *Plin4* | 3.11E-26 | 1.606197 |
| ENSMUSG00000095079 | *Igha* | 1.40E-05 | 1.607185 |
| ENSMUSG00000078902 | *Gm14443* | 1.56E-02 | 1.607245 |
| ENSMUSG00000115855 | *Gm34643* | 1.33E-02 | 1.60811 |
| ENSMUSG00000073492 | *Gm10521* | 1.01E-02 | 1.608342 |
| ENSMUSG00000058818 | *Pirb* | 1.25E-16 | 1.608386 |
| ENSMUSG00000071562 | *Stfa1* | 1.20E-02 | 1.608568 |
| ENSMUSG00000058755 | *Osm* | 9.87E-06 | 1.608663 |
| ENSMUSG00000025479 | *Cyp2e1* | 1.75E-15 | 1.608811 |
| ENSMUSG00000116975 | *AC097366.1* | 4.70E-02 | 1.609642 |

| | | | |
|---|---|---|---|
| ENSMUSG00000061126 | *Cyp4f39* | 1.24E-02 | 1.611491 |
| ENSMUSG00000086756 | *Gm14051* | 4.26E-03 | 1.612421 |
| ENSMUSG00000054383 | *Pnma1* | 1.58E-02 | 1.612439 |
| ENSMUSG00000084788 | *Gm11342* | 1.75E-08 | 1.6151 |
| ENSMUSG00000102776 | *Gm38162* | 1.33E-04 | 1.615539 |
| ENSMUSG00000104453 | *Gm37829* | 3.14E-03 | 1.616164 |
| ENSMUSG00000049709 | *Nlrp10* | 1.69E-03 | 1.616218 |
| ENSMUSG00000047798 | *Cd300lf* | 2.12E-26 | 1.616545 |
| ENSMUSG00000093938 | *Evi2b* | 9.87E-15 | 1.61906 |
| ENSMUSG00000031303 | *Map3k15* | 8.44E-11 | 1.619429 |
| ENSMUSG00000110195 | *Pde2a* | 3.98E-16 | 1.622023 |
| ENSMUSG00000109819 | *B930018H19Rik* | 2.50E-04 | 1.622025 |
| ENSMUSG00000028860 | *Sytl1* | 2.85E-04 | 1.623336 |
| ENSMUSG00000034570 | *Inpp5j* | 2.35E-03 | 1.623616 |
| ENSMUSG00000026220 | *Slc16a14* | 3.53E-02 | 1.625516 |
| ENSMUSG00000116290 | *Gm49518* | 1.89E-02 | 1.626577 |
| ENSMUSG00000096957 | *E230013L22Rik* | 7.49E-09 | 1.626815 |
| ENSMUSG00000003153 | *Slc2a3* | 1.50E-16 | 1.627617 |
| ENSMUSG00000074419 | *Gm15448* | 4.65E-11 | 1.62794 |
| ENSMUSG00000067577 | *A430093F15Rik* | 6.34E-09 | 1.628234 |
| ENSMUSG00000079597 | *Gm5483* | 5.40E-13 | 1.628886 |
| ENSMUSG00000018507 | *Trpv2* | 5.02E-19 | 1.629285 |
| ENSMUSG00000032023 | *Jhy* | 7.50E-07 | 1.630081 |
| ENSMUSG00000042567 | *Nek10* | 7.69E-11 | 1.630143 |
| ENSMUSG00000022876 | *Samsn1* | 1.31E-21 | 1.6304 |
| ENSMUSG00000113998 | *Gm40909* | 5.76E-07 | 1.630854 |
| ENSMUSG00000022454 | *Nell2* | 1.66E-02 | 1.631056 |
| ENSMUSG00000107480 | *Gm44165* | 2.95E-03 | 1.631572 |
| ENSMUSG00000115355 | *4930445E18Rik* | 5.26E-04 | 1.631708 |
| ENSMUSG00000040287 | *Stac3* | 6.79E-06 | 1.632241 |
| ENSMUSG00000116747 | *AC144408.2* | 2.16E-02 | 1.633409 |
| ENSMUSG00000056071 | *S100a9* | 5.19E-11 | 1.634188 |
| ENSMUSG00000066800 | *Rnasel* | 7.15E-19 | 1.63539 |
| ENSMUSG00000000386 | *Mx1* | 3.52E-19 | 1.638433 |
| ENSMUSG00000030669 | *Calca* | 6.86E-06 | 1.640644 |
| ENSMUSG00000099364 | *5730419F03Rik* | 2.51E-02 | 1.642235 |
| ENSMUSG00000041754 | *Trem3* | 2.37E-12 | 1.64293 |
| ENSMUSG00000048416 | *Mlf1* | 3.35E-12 | 1.64339 |
| ENSMUSG00000056054 | *S100a8* | 5.09E-08 | 1.644216 |
| ENSMUSG00000011148 | *Adssl1* | 5.36E-13 | 1.645005 |
| ENSMUSG00000052726 | *Kcnt2* | 2.51E-19 | 1.645058 |
| ENSMUSG00000029228 | *Lnx1* | 2.29E-25 | 1.645255 |
| ENSMUSG00000116031 | *D030024E09Rik* | 2.72E-05 | 1.645676 |
| ENSMUSG00000099092 | *Mir7062* | 3.90E-02 | 1.646838 |
| ENSMUSG00000109096 | *Gm44888* | 1.16E-02 | 1.647995 |
| ENSMUSG00000038132 | *Rbm24* | 2.05E-13 | 1.648088 |
| ENSMUSG00000000673 | *Haao* | 2.61E-03 | 1.648454 |
| ENSMUSG00000026073 | *Il1r2* | 8.70E-09 | 1.64854 |
| ENSMUSG00000037731 | *Themis2* | 6.62E-24 | 1.649889 |
| ENSMUSG00000091438 | *Gm17088* | 1.47E-02 | 1.650306 |
| ENSMUSG00000064346 | *mt-Tw* | 8.31E-04 | 1.65172 |
| ENSMUSG00000034855 | *Cxcl10* | 1.23E-06 | 1.652425 |
| ENSMUSG00000031312 | *Itgb1bp2* | 3.28E-08 | 1.652551 |

| | | | |
|---|---|---|---|
| ENSMUSG00000059657 | *Stfa2l1* | 1.02E-14 | 1.652763 |
| ENSMUSG00000024379 | *Tslp* | 1.96E-07 | 1.652917 |
| ENSMUSG00000032648 | *Pygm* | 1.27E-25 | 1.653929 |
| ENSMUSG00000064360 | *mt-Nd3* | 6.22E-34 | 1.653955 |
| ENSMUSG00000005686 | *Ampd3* | 5.07E-44 | 1.654326 |
| ENSMUSG00000036067 | *Slc2a6* | 2.22E-21 | 1.655183 |
| ENSMUSG00000110125 | *Gm33148* | 4.23E-02 | 1.655315 |
| ENSMUSG00000021071 | *Trim9* | 4.06E-04 | 1.655679 |
| ENSMUSG00000012819 | *Cdh23* | 5.33E-08 | 1.655851 |
| ENSMUSG00000029602 | *Rasal1* | 1.09E-02 | 1.6569 |
| ENSMUSG00000038204 | *Asb10* | 3.79E-02 | 1.657364 |
| ENSMUSG00000034755 | *Pcdh11x* | 2.75E-05 | 1.65877 |
| ENSMUSG00000038067 | *Csf3* | 2.30E-17 | 1.659219 |
| ENSMUSG00000029059 | *Fam213b* | 2.03E-02 | 1.659422 |
| ENSMUSG00000032033 | *Barx2* | 2.86E-06 | 1.659698 |
| ENSMUSG00000034686 | *Prr7* | 4.58E-13 | 1.660366 |
| ENSMUSG00000101939 | *Gm28438* | 1.42E-34 | 1.661494 |
| ENSMUSG00000100510 | *AV026068* | 6.71E-04 | 1.66215 |
| ENSMUSG00000046610 | *Oacyl* | 7.18E-05 | 1.662776 |
| ENSMUSG00000028834 | *Trim63* | 7.79E-07 | 1.663212 |
| ENSMUSG00000003863 | *Ppfia3* | 3.09E-02 | 1.664438 |
| ENSMUSG00000031626 | *Sorbs2* | 1.22E-22 | 1.665918 |
| ENSMUSG00000047501 | *Cldn4* | 2.88E-03 | 1.66654 |
| ENSMUSG00000087579 | *Hectd2os* | 4.33E-09 | 1.666956 |
| ENSMUSG00000022821 | *Hgd* | 9.49E-04 | 1.667405 |
| ENSMUSG00000020067 | *Mypn* | 2.91E-16 | 1.668111 |
| ENSMUSG00000074071 | *Fam169b* | 1.03E-40 | 1.669004 |
| ENSMUSG00000093806 | *Asmt* | 3.89E-02 | 1.669364 |
| ENSMUSG00000031258 | *Xkrx* | 2.29E-02 | 1.669615 |
| ENSMUSG00000040435 | *Ppp1r15a* | 1.10E-25 | 1.669806 |
| ENSMUSG00000097643 | *A130051J06Rik* | 3.09E-02 | 1.669883 |
| ENSMUSG00000079644 | *Gm1110* | 2.01E-02 | 1.67036 |
| ENSMUSG00000042717 | *Ppp1r3a* | 8.51E-16 | 1.670789 |
| ENSMUSG00000075042 | *4930431P03Rik* | 6.43E-03 | 1.670913 |
| ENSMUSG00000054717 | *Hmgb2* | 8.73E-07 | 1.670984 |
| ENSMUSG00000082179 | *Gm11407* | 1.60E-02 | 1.671338 |
| ENSMUSG00000056133 | *Unc93a2* | 1.34E-02 | 1.672597 |
| ENSMUSG00000101640 | *Gm28376* | 2.21E-02 | 1.673873 |
| ENSMUSG00000030156 | *Cd69* | 2.53E-09 | 1.674983 |
| ENSMUSG00000113185 | *Gm47127* | 3.62E-10 | 1.675362 |
| ENSMUSG00000038239 | *Hrc* | 1.25E-12 | 1.67644 |
| ENSMUSG00000078486 | *Perm1* | 3.53E-12 | 1.677159 |
| ENSMUSG00000059741 | *Myl3* | 2.68E-08 | 1.677439 |
| ENSMUSG00000027843 | *Ptpn22* | 1.42E-17 | 1.677529 |
| ENSMUSG00000098670 | *Gm27818* | 6.52E-03 | 1.678782 |
| ENSMUSG00000003476 | *Crhr2* | 1.34E-06 | 1.679035 |
| ENSMUSG00000091712 | *Sec14l5* | 1.80E-04 | 1.679152 |
| ENSMUSG00000103308 | *Gm37800* | 3.21E-05 | 1.680186 |
| ENSMUSG00000026358 | *Rgs1* | 3.15E-07 | 1.680205 |
| ENSMUSG00000109864 | *Eid3* | 9.84E-08 | 1.682789 |
| ENSMUSG00000073771 | *Btbd19* | 4.74E-06 | 1.683039 |
| ENSMUSG00000090439 | *Gm17455* | 2.54E-02 | 1.683429 |
| ENSMUSG00000035653 | *Lrfn5* | 2.04E-02 | 1.684683 |

| | | | |
|---|---|---|---|
| ENSMUSG00000086335 | *Gm12107* | 4.07E-02 | 1.684829 |
| ENSMUSG00000111809 | *Gm49334* | 1.68E-02 | 1.687123 |
| ENSMUSG00000006724 | *Cyp27b1* | 2.49E-02 | 1.688563 |
| ENSMUSG00000097028 | *Ptgs2os* | 4.00E-03 | 1.688812 |
| ENSMUSG00000027966 | *Col11a1* | 6.48E-06 | 1.689108 |
| ENSMUSG00000042251 | *Pm20d1* | 9.34E-04 | 1.689267 |
| ENSMUSG00000104554 | *Gm4610* | 2.85E-03 | 1.689388 |
| ENSMUSG00000028544 | *Slc5a9* | 3.63E-03 | 1.690419 |
| ENSMUSG00000027861 | *Casq2* | 1.46E-13 | 1.690798 |
| ENSMUSG00000087684 | *1200007C13Rik* | 9.26E-19 | 1.692709 |
| ENSMUSG00000046223 | *Plaur* | 3.12E-25 | 1.69343 |
| ENSMUSG00000086727 | *4931428L18Rik* | 4.68E-02 | 1.694028 |
| ENSMUSG00000018470 | *Kcnab3* | 4.44E-02 | 1.695106 |
| ENSMUSG00000020722 | *Cacng1* | 3.42E-03 | 1.695444 |
| ENSMUSG00000050966 | *Lin28a* | 3.70E-02 | 1.696226 |
| ENSMUSG00000031245 | *Hmgn5* | 1.10E-06 | 1.696266 |
| ENSMUSG00000084760 | *Gm15472* | 4.94E-02 | 1.699259 |
| ENSMUSG00000029379 | *Cxcl3* | 3.14E-19 | 1.699755 |
| ENSMUSG00000106741 | *4930513D17Rik* | 2.78E-04 | 1.700156 |
| ENSMUSG00000033676 | *Gabrb3* | 9.53E-03 | 1.700516 |
| ENSMUSG00000111827 | *Gm8131* | 2.03E-02 | 1.701357 |
| ENSMUSG00000026989 | *Dapl1* | 1.87E-02 | 1.702816 |
| ENSMUSG00000021298 | *Gpr132* | 1.17E-13 | 1.705667 |
| ENSMUSG00000005373 | *Mlxipl* | 2.58E-12 | 1.705754 |
| ENSMUSG00000022651 | *Retnlg* | 1.15E-21 | 1.705788 |
| ENSMUSG00000036557 | *Stpg4* | 7.22E-03 | 1.706336 |
| ENSMUSG00000060600 | *Eno3* | 1.40E-17 | 1.70718 |
| ENSMUSG00000092134 | *Gm17089* | 2.08E-03 | 1.708295 |
| ENSMUSG00000115610 | *Gm31251* | 8.64E-13 | 1.708416 |
| ENSMUSG00000027824 | *Vmn2r1* | 3.39E-02 | 1.71058 |
| ENSMUSG00000057580 | *Gm10012* | 5.95E-03 | 1.711245 |
| ENSMUSG00000090785 | *Gm17116* | 3.94E-02 | 1.712965 |
| ENSMUSG00000109005 | *Gm45221* | 9.18E-13 | 1.713519 |
| ENSMUSG00000097652 | *Mhrt* | 5.45E-09 | 1.713729 |
| ENSMUSG00000039376 | *Synpo2l* | 2.08E-14 | 1.714714 |
| ENSMUSG00000030935 | *Acsm3* | 4.85E-02 | 1.714851 |
| ENSMUSG00000085792 | *Gm15414* | 4.12E-02 | 1.715017 |
| ENSMUSG00000032845 | *Alpk2* | 8.22E-25 | 1.71506 |
| ENSMUSG00000024215 | *Spdef* | 3.57E-05 | 1.715667 |
| ENSMUSG00000040490 | *Lrfn2* | 6.64E-03 | 1.717782 |
| ENSMUSG00000067049 | *Unc93a* | 3.95E-04 | 1.719453 |
| ENSMUSG00000020836 | *Coro6* | 4.06E-14 | 1.719576 |
| ENSMUSG00000025236 | *Adpgk* | 9.28E-08 | 1.720655 |
| ENSMUSG00000069670 | *Nkain2* | 2.61E-27 | 1.720935 |
| ENSMUSG00000054204 | *Alkal2* | 9.31E-03 | 1.721026 |
| ENSMUSG00000050439 | *Enthd1* | 1.61E-03 | 1.722314 |
| ENSMUSG00000058966 | *Fam57b* | 1.21E-04 | 1.722847 |
| ENSMUSG00000039521 | *Foxp3* | 3.73E-04 | 1.722974 |
| ENSMUSG00000042451 | *Mybph* | 4.32E-14 | 1.725736 |
| ENSMUSG00000025746 | *Il6* | 5.63E-08 | 1.726006 |
| ENSMUSG00000087382 | *Ctcflos* | 3.89E-03 | 1.729287 |
| ENSMUSG00000024521 | *Pmaip1* | 5.17E-21 | 1.731247 |
| ENSMUSG00000113321 | *Gm8075* | 1.52E-03 | 1.732541 |

| | | | |
|---|---|---|---|
| ENSMUSG00000028125 | *Abca4* | 1.40E-15 | 1.733301 |
| ENSMUSG00000109510 | *Gm42417* | 2.03E-18 | 1.733587 |
| ENSMUSG00000056643 | *Chst13* | 3.89E-02 | 1.737568 |
| ENSMUSG00000109002 | *Gm38405* | 3.68E-03 | 1.738016 |
| ENSMUSG00000066366 | *Serpina1a* | 4.81E-04 | 1.739086 |
| ENSMUSG00000030144 | *Clec4d* | 1.24E-21 | 1.739304 |
| ENSMUSG00000074899 | *Sptbn5* | 8.93E-15 | 1.741636 |
| ENSMUSG00000019933 | *Mrln* | 6.47E-04 | 1.74233 |
| ENSMUSG00000117297 | *CT571247.2* | 2.12E-03 | 1.742747 |
| ENSMUSG00000029838 | *Ptn* | 3.58E-04 | 1.745315 |
| ENSMUSG00000110626 | *Gm45805* | 3.93E-03 | 1.746875 |
| ENSMUSG00000114638 | *Gm31834* | 1.32E-02 | 1.748994 |
| ENSMUSG00000004902 | *Slc25a18* | 9.40E-11 | 1.748994 |
| ENSMUSG00000021886 | *Gpr65* | 1.38E-16 | 1.751875 |
| ENSMUSG00000111334 | *AC162938.1* | 2.19E-03 | 1.751922 |
| ENSMUSG00000090141 | *Gm614* | 1.79E-07 | 1.752594 |
| ENSMUSG00000028116 | *Myoz2* | 1.41E-18 | 1.753998 |
| ENSMUSG00000024912 | *Fosl1* | 4.81E-11 | 1.757523 |
| ENSMUSG00000047281 | *Sfn* | 1.96E-12 | 1.758427 |
| ENSMUSG00000048489 | *Depp1* | 5.70E-10 | 1.75859 |
| ENSMUSG00000052769 | *Gm9889* | 3.49E-03 | 1.758749 |
| ENSMUSG00000038242 | *Aox4* | 3.63E-04 | 1.759686 |
| ENSMUSG00000047940 | *Stpg2* | 1.76E-04 | 1.761496 |
| ENSMUSG00000067242 | *Lgi1* | 2.15E-03 | 1.76182 |
| ENSMUSG00000079015 | *Serpina1c* | 9.81E-05 | 1.762405 |
| ENSMUSG00000028573 | *Fggy* | 9.93E-05 | 1.763622 |
| ENSMUSG00000106547 | *B230303O12Rik* | 1.91E-10 | 1.763742 |
| ENSMUSG00000110010 | *Gm45629* | 1.72E-03 | 1.764186 |
| ENSMUSG00000106478 | *Gm36551* | 2.69E-03 | 1.766268 |
| ENSMUSG00000034919 | *Ttc22* | 1.21E-03 | 1.767889 |
| ENSMUSG00000028773 | *Fabp3* | 1.25E-09 | 1.769041 |
| ENSMUSG00000026564 | *Dusp27* | 4.71E-11 | 1.770954 |
| ENSMUSG00000079429 | *Mroh2a* | 4.95E-02 | 1.771987 |
| ENSMUSG00000030510 | *Cers3* | 8.43E-08 | 1.773692 |
| ENSMUSG00000094334 | *Fabp5l2* | 1.94E-02 | 1.773759 |
| ENSMUSG00000030235 | *Slco1c1* | 6.91E-16 | 1.775853 |
| ENSMUSG00000037989 | *Wnk2* | 2.70E-20 | 1.777419 |
| ENSMUSG00000027674 | *Pex5l* | 4.64E-10 | 1.777644 |
| ENSMUSG00000004668 | *Abca13* | 5.69E-29 | 1.778516 |
| ENSMUSG00000062713 | *Sim2* | 1.19E-09 | 1.779309 |
| ENSMUSG00000116961 | *AC117662.4* | 3.90E-05 | 1.7808 |
| ENSMUSG00000026669 | *Mcm10* | 2.70E-08 | 1.780809 |
| ENSMUSG00000049598 | *Vsig8* | 3.66E-02 | 1.781692 |
| ENSMUSG00000051747 | *Ttn* | 1.20E-50 | 1.787185 |
| ENSMUSG00000085417 | *Gm13919* | 2.58E-03 | 1.789173 |
| ENSMUSG00000033544 | *Angptl1* | 2.01E-03 | 1.789715 |
| ENSMUSG00000020000 | *Moxd1* | 3.38E-03 | 1.789741 |
| ENSMUSG00000001249 | *Hpn* | 8.32E-15 | 1.79272 |
| ENSMUSG00000034648 | *Lrrn1* | 4.21E-04 | 1.792909 |
| ENSMUSG00000090173 | *Fbxw10* | 4.07E-06 | 1.793362 |
| ENSMUSG00000110580 | *D830024N08Rik* | 3.86E-07 | 1.79383 |
| ENSMUSG00000027559 | *Car3* | 1.14E-58 | 1.794165 |
| ENSMUSG00000049173 | *Myoz3* | 3.68E-04 | 1.794511 |

| | | | |
|---|---|---|---|
| ENSMUSG00000097579 | *Gm26799* | 1.22E-05 | 1.798591 |
| ENSMUSG00000108322 | *5430431A17Rik* | 9.26E-04 | 1.79877 |
| ENSMUSG00000041954 | *Tnfrsf18* | 1.91E-04 | 1.798838 |
| ENSMUSG00000086236 | *5830418P13Rik* | 3.97E-02 | 1.799096 |
| ENSMUSG00000107667 | *C530044C16Rik* | 2.55E-03 | 1.799886 |
| ENSMUSG00000032484 | *Ngp* | 7.22E-33 | 1.802514 |
| ENSMUSG00000097543 | *Gm805* | 3.14E-03 | 1.804272 |
| ENSMUSG00000042045 | *Sln* | 1.59E-14 | 1.805054 |
| ENSMUSG00000031382 | *Asb11* | 5.07E-07 | 1.807798 |
| ENSMUSG00000039653 | *Baat* | 1.72E-02 | 1.807968 |
| ENSMUSG00000109921 | *Gm33326* | 4.27E-02 | 1.808594 |
| ENSMUSG00000046818 | *Ddit4l* | 1.37E-03 | 1.809116 |
| ENSMUSG00000044083 | *Efcab8* | 1.54E-22 | 1.80916 |
| ENSMUSG00000101639 | *Gm8597* | 2.68E-02 | 1.810383 |
| ENSMUSG00000000223 | *Drp2* | 8.23E-07 | 1.810905 |
| ENSMUSG00000085584 | *Rtl9* | 4.99E-02 | 1.814506 |
| ENSMUSG00000006345 | *Ggt1* | 2.21E-08 | 1.817124 |
| ENSMUSG00000086448 | *9330162012Rik* | 3.73E-02 | 1.817998 |
| ENSMUSG00000067889 | *Sptbn2* | 4.24E-09 | 1.818248 |
| ENSMUSG00000038763 | *Alpk3* | 1.96E-32 | 1.81872 |
| ENSMUSG00000085194 | *Platr32* | 2.24E-03 | 1.819714 |
| ENSMUSG00000030470 | *Csrp3* | 2.08E-18 | 1.819847 |
| ENSMUSG00000043008 | *Klhl6* | 2.15E-21 | 1.820648 |
| ENSMUSG00000019987 | *Arg1* | 2.20E-03 | 1.822152 |
| ENSMUSG00000039264 | *Gimap3* | 3.51E-14 | 1.824217 |
| ENSMUSG00000060470 | *Adgrg3* | 1.14E-17 | 1.824816 |
| ENSMUSG00000025488 | *Cox8b* | 1.08E-07 | 1.825697 |
| ENSMUSG00000097071 | *4930544I03Rik* | 5.59E-06 | 1.830008 |
| ENSMUSG00000028680 | *Plk3* | 8.61E-23 | 1.830772 |
| ENSMUSG00000020007 | *Il20ra* | 1.06E-02 | 1.831433 |
| ENSMUSG00000074340 | *Ovgp1* | 2.09E-12 | 1.832669 |
| ENSMUSG00000112148 | *Lilrb4a* | 1.06E-12 | 1.832695 |
| ENSMUSG00000034145 | *Tmem63c* | 7.44E-03 | 1.833488 |
| ENSMUSG00000031837 | *Necab2* | 4.40E-02 | 1.833535 |
| ENSMUSG00000040694 | *Apobec2* | 1.11E-11 | 1.834504 |
| ENSMUSG00000021255 | *Esrrb* | 6.95E-05 | 1.835067 |
| ENSMUSG00000092008 | *Cyp2c69* | 1.42E-02 | 1.83538 |
| ENSMUSG00000103485 | *Gm38001* | 2.57E-02 | 1.836426 |
| ENSMUSG00000029660 | *Tex26* | 9.36E-09 | 1.837627 |
| ENSMUSG00000086405 | *9330198N18Rik* | 1.69E-08 | 1.837669 |
| ENSMUSG00000037509 | *Arhgef4* | 1.46E-11 | 1.84023 |
| ENSMUSG00000041737 | *Tmem45b* | 8.44E-08 | 1.840717 |
| ENSMUSG00000027359 | *Slc27a2* | 5.04E-03 | 1.841241 |
| ENSMUSG00000047419 | *Cmya5* | 5.22E-34 | 1.843129 |
| ENSMUSG00000033182 | *Kbtbd12* | 1.30E-10 | 1.843376 |
| ENSMUSG00000044938 | *Klhl31* | 4.21E-15 | 1.843504 |
| ENSMUSG00000109675 | *Nxpe1-ps* | 2.49E-02 | 1.843549 |
| ENSMUSG00000000049 | *Apoh* | 9.36E-03 | 1.843946 |
| ENSMUSG00000105107 | *Gm43412* | 3.93E-05 | 1.845797 |
| ENSMUSG00000023341 | *Mx2* | 2.68E-43 | 1.846495 |
| ENSMUSG00000053641 | *Dennd4a* | 7.82E-42 | 1.846884 |
| ENSMUSG00000029158 | *Yipf7* | 5.09E-04 | 1.849072 |
| ENSMUSG00000032278 | *Paqr5* | 1.41E-08 | 1.850269 |

| | | | |
|---|---|---|---|
| ENSMUSG00000031098 | *Syt8* | 9.62E-03 | 1.851676 |
| ENSMUSG00000090118 | *Gm16163* | 8.91E-07 | 1.853032 |
| ENSMUSG00000024526 | *Cidea* | 2.58E-04 | 1.853747 |
| ENSMUSG00000115919 | *Gm31583* | 2.47E-02 | 1.853826 |
| ENSMUSG00000114372 | *2310067P03Rik* | 2.35E-03 | 1.854639 |
| ENSMUSG00000081700 | *Atp5k-ps2* | 8.27E-05 | 1.855005 |
| ENSMUSG00000066491 | *Cox6c2* | 5.55E-11 | 1.856124 |
| ENSMUSG00000031461 | *Myom2* | 1.24E-37 | 1.857642 |
| ENSMUSG00000104956 | *4930429D17Rik* | 1.29E-02 | 1.859359 |
| ENSMUSG00000041616 | *Nppa* | 8.40E-06 | 1.859817 |
| ENSMUSG00000030834 | *Abcc6* | 1.85E-02 | 1.86125 |
| ENSMUSG00000044367 | *Slc16a13* | 1.02E-23 | 1.863682 |
| ENSMUSG00000026100 | *Mstn* | 1.03E-02 | 1.863901 |
| ENSMUSG00000058975 | *Kcnc1* | 1.48E-05 | 1.864086 |
| ENSMUSG00000044086 | *Lmod3* | 5.83E-05 | 1.864089 |
| ENSMUSG00000032262 | *Elovl4* | 1.84E-06 | 1.865299 |
| ENSMUSG00000022824 | *Muc13* | 2.46E-06 | 1.871007 |
| ENSMUSG00000105986 | *Gm43065* | 1.82E-03 | 1.871109 |
| ENSMUSG00000091119 | *Ccdc152* | 3.30E-05 | 1.87189 |
| ENSMUSG00000109198 | *D7Bwg0826e* | 6.88E-03 | 1.872806 |
| ENSMUSG00000087187 | *Gm13431* | 2.85E-04 | 1.873198 |
| ENSMUSG00000106245 | *Gm43824* | 2.54E-03 | 1.874635 |
| ENSMUSG00000110411 | *Gm45457* | 2.24E-03 | 1.87632 |
| ENSMUSG00000025069 | *Gsto2* | 7.14E-04 | 1.878743 |
| ENSMUSG00000090066 | *1110002E22Rik* | 4.80E-18 | 1.881563 |
| ENSMUSG00000110498 | *A630001O12Rik* | 2.05E-05 | 1.881948 |
| ENSMUSG00000032174 | *Icam5* | 1.60E-03 | 1.884311 |
| ENSMUSG00000040350 | *Trim7* | 3.16E-10 | 1.886702 |
| ENSMUSG00000063001 | *Gm9701* | 2.00E-02 | 1.886992 |
| ENSMUSG00000028017 | *Egf* | 2.98E-15 | 1.887283 |
| ENSMUSG00000070498 | *Tmem132b* | 7.38E-17 | 1.887425 |
| ENSMUSG00000028427 | *Aqp7* | 6.69E-03 | 1.888129 |
| ENSMUSG00000038508 | *Gdf15* | 6.53E-03 | 1.888853 |
| ENSMUSG00000027513 | *Pck1* | 9.43E-13 | 1.890906 |
| ENSMUSG00000059900 | *Tmem40* | 4.20E-04 | 1.893524 |
| ENSMUSG00000115518 | *Gm10791* | 1.65E-05 | 1.895648 |
| ENSMUSG00000090293 | *Gm17034* | 2.96E-02 | 1.897849 |
| ENSMUSG00000025004 | *Cyp2c40* | 1.94E-02 | 1.898265 |
| ENSMUSG00000112895 | *Gm47567* | 5.03E-05 | 1.899642 |
| ENSMUSG00000031958 | *Ldhd* | 4.49E-07 | 1.899952 |
| ENSMUSG00000022582 | *Ly6g* | 4.01E-07 | 1.902278 |
| ENSMUSG00000062077 | *Trim54* | 1.10E-15 | 1.902342 |
| ENSMUSG00000024617 | *Camk2a* | 4.89E-09 | 1.903979 |
| ENSMUSG00000007122 | *Casq1* | 2.89E-15 | 1.904008 |
| ENSMUSG00000069733 | *Ube2u* | 1.19E-03 | 1.904037 |
| ENSMUSG00000024430 | *Cabyr* | 1.01E-03 | 1.904429 |
| ENSMUSG00000056569 | *Mpz* | 1.14E-16 | 1.905577 |
| ENSMUSG00000032942 | *Ucp3* | 9.12E-10 | 1.909391 |
| ENSMUSG00000044165 | *Bcl2l15* | 4.24E-09 | 1.911503 |
| ENSMUSG00000079588 | *Tmem182* | 5.78E-17 | 1.91235 |
| ENSMUSG00000007877 | *Tcap* | 1.05E-20 | 1.912751 |
| ENSMUSG00000025473 | *Adam8* | 9.12E-15 | 1.913184 |
| ENSMUSG00000031376 | *Atp2b3* | 1.10E-04 | 1.915267 |

| | | | |
|---|---|---|---|
| ENSMUSG00000099760 | *Gm28800* | 1.04E-02 | 1.918089 |
| ENSMUSG00000085249 | *Gm12847* | 6.44E-04 | 1.921004 |
| ENSMUSG00000105826 | *Gm42141* | 5.75E-03 | 1.921732 |
| ENSMUSG00000112082 | *Gm35035* | 1.26E-03 | 1.922335 |
| ENSMUSG00000033213 | *AA467197* | 2.17E-05 | 1.926523 |
| ENSMUSG00000046764 | *A530053G22Rik* | 5.14E-03 | 1.927384 |
| ENSMUSG00000074623 | *Gm826* | 1.94E-03 | 1.931273 |
| ENSMUSG00000030399 | *Ckm* | 2.68E-12 | 1.931828 |
| ENSMUSG00000101930 | *Gm5441* | 1.33E-10 | 1.932682 |
| ENSMUSG00000027456 | *Sdcbp2* | 5.13E-14 | 1.933121 |
| ENSMUSG00000086765 | *Gm11827* | 2.95E-06 | 1.933308 |
| ENSMUSG00000027505 | *Fam209* | 8.82E-04 | 1.935233 |
| ENSMUSG00000054905 | *Stfa3* | 8.40E-04 | 1.935879 |
| ENSMUSG00000062515 | *Fabp4* | 8.09E-25 | 1.936282 |
| ENSMUSG00000024223 | *Armc12* | 3.96E-02 | 1.94095 |
| ENSMUSG00000026068 | *Il18rap* | 6.22E-34 | 1.941935 |
| ENSMUSG00000026489 | *Coq8a* | 2.76E-39 | 1.94642 |
| ENSMUSG00000105542 | *Gm34248* | 2.05E-02 | 1.947483 |
| ENSMUSG00000022468 | *Endou* | 3.09E-13 | 1.95126 |
| ENSMUSG00000062372 | *Otof* | 4.14E-03 | 1.953074 |
| ENSMUSG00000060962 | *Dmkn* | 4.46E-16 | 1.954338 |
| ENSMUSG00000075307 | *Klhl41* | 3.80E-19 | 1.954528 |
| ENSMUSG00000013936 | *Myl2* | 8.34E-04 | 1.957148 |
| ENSMUSG00000074218 | *Cox7a1* | 4.66E-09 | 1.957494 |
| ENSMUSG00000103965 | *Gm30173* | 8.23E-05 | 1.957865 |
| ENSMUSG00000015879 | *Fam184b* | 9.58E-04 | 1.958074 |
| ENSMUSG00000030236 | *Slco1b2* | 6.11E-03 | 1.958235 |
| ENSMUSG00000116641 | *AC165271.1* | 3.93E-02 | 1.960352 |
| ENSMUSG00000029683 | *Lmod2* | 2.29E-19 | 1.962142 |
| ENSMUSG00000058427 | *Cxcl2* | 7.52E-16 | 1.965001 |
| ENSMUSG00000097468 | *Mdrl* | 1.18E-02 | 1.9652 |
| ENSMUSG00000085739 | *Gm15862* | 4.50E-02 | 1.966473 |
| ENSMUSG00000099906 | *Gm28653* | 1.68E-02 | 1.966573 |
| ENSMUSG00000097622 | *A330033J07Rik* | 9.21E-03 | 1.967677 |
| ENSMUSG00000025929 | *Il17a* | 6.98E-03 | 1.968258 |
| ENSMUSG00000101969 | *Gm20125* | 7.68E-08 | 1.970172 |
| ENSMUSG00000109881 | *Gm45507* | 3.61E-02 | 1.972472 |
| ENSMUSG00000025892 | *Gria4* | 1.74E-04 | 1.973108 |
| ENSMUSG00000028786 | *Tmem54* | 2.69E-02 | 1.973491 |
| ENSMUSG00000037188 | *Grhl3* | 3.13E-06 | 1.976128 |
| ENSMUSG00000028736 | *Pax7* | 2.68E-02 | 1.978015 |
| ENSMUSG00000117029 | *CT030155.1* | 3.35E-03 | 1.980083 |
| ENSMUSG00000072720 | *Myo18b* | 5.67E-49 | 1.980402 |
| ENSMUSG00000001027 | *Scn4a* | 1.63E-15 | 1.982106 |
| ENSMUSG00000020848 | *Doc2b* | 7.17E-17 | 1.983107 |
| ENSMUSG00000071540 | *3425401B19Rik* | 1.11E-21 | 1.983567 |
| ENSMUSG00000059108 | *Ifitm6* | 1.39E-27 | 1.986147 |
| ENSMUSG00000026166 | *Ccl20* | 1.87E-24 | 1.988087 |
| ENSMUSG00000032021 | *Crtam* | 1.62E-20 | 1.988346 |
| ENSMUSG00000048939 | *Atp13a5* | 8.98E-03 | 1.988773 |
| ENSMUSG00000027961 | *Lrrc39* | 1.97E-07 | 1.989628 |
| ENSMUSG00000050211 | *Pla2g4e* | 2.57E-15 | 1.990622 |
| ENSMUSG00000050612 | *Txndc2* | 9.64E-03 | 1.991405 |

| | | | |
|---|---|---|---|
| ENSMUSG00000097476 | *Gm26583* | 2.67E-02 | 1.992121 |
| ENSMUSG00000018893 | *Mb* | 2.96E-16 | 1.995476 |
| ENSMUSG00000001497 | *Pax9* | 5.45E-07 | 1.99584 |
| ENSMUSG00000022871 | *Fetub* | 5.14E-07 | 1.996529 |
| ENSMUSG00000040536 | *Necab1* | 6.95E-04 | 1.9986 |
| ENSMUSG00000109831 | *Gm36431* | 7.50E-03 | 1.99936 |
| ENSMUSG00000102762 | *4930403P22Rik* | 2.46E-02 | 2.001551 |
| ENSMUSG00000041782 | *Lad1* | 3.08E-13 | 2.002946 |
| ENSMUSG00000043873 | *Chil5* | 7.40E-06 | 2.00396 |
| ENSMUSG00000085806 | *Gm12023* | 2.03E-05 | 2.005786 |
| ENSMUSG00000035686 | *Thrsp* | 3.08E-18 | 2.008526 |
| ENSMUSG00000067653 | *Ankrd23* | 7.18E-23 | 2.012683 |
| ENSMUSG00000021622 | *Ckmt2* | 2.18E-22 | 2.016924 |
| ENSMUSG00000046480 | *Scn4b* | 3.61E-12 | 2.018601 |
| ENSMUSG00000114656 | *2810403G07Rik* | 2.60E-02 | 2.018932 |
| ENSMUSG00000022525 | *Hrasls* | 1.48E-02 | 2.019224 |
| ENSMUSG00000070644 | *Etnk2* | 8.41E-03 | 2.020491 |
| ENSMUSG00000042306 | *S100a14* | 4.36E-12 | 2.021351 |
| ENSMUSG00000098132 | *Rassf10* | 1.04E-02 | 2.021448 |
| ENSMUSG00000062593 | *Gm49339* | 1.64E-14 | 2.024042 |
| ENSMUSG00000111299 | *4930540M03Rik* | 1.34E-02 | 2.024137 |
| ENSMUSG00000021340 | *Gpld1* | 6.91E-16 | 2.028941 |
| ENSMUSG00000046352 | *Gjb2* | 1.21E-11 | 2.030651 |
| ENSMUSG00000103620 | *Gm37359* | 4.58E-06 | 2.031625 |
| ENSMUSG00000085743 | *8430419K02Rik* | 1.91E-27 | 2.032356 |
| ENSMUSG00000089854 | *Gm16133* | 2.61E-06 | 2.035203 |
| ENSMUSG00000006204 | *5430419D17Rik* | 3.20E-02 | 2.035247 |
| ENSMUSG00000021509 | *Slc25a48* | 4.04E-03 | 2.035611 |
| ENSMUSG00000061462 | *Obscn* | 5.68E-58 | 2.038968 |
| ENSMUSG00000059994 | *Fcrl1* | 1.30E-09 | 2.039878 |
| ENSMUSG00000076275 | *Mir675* | 1.77E-07 | 2.041447 |
| ENSMUSG00000069372 | *Ctxn3* | 3.18E-08 | 2.043908 |
| ENSMUSG00000037129 | *Tmprss13* | 8.77E-06 | 2.050404 |
| ENSMUSG00000021223 | *Papln* | 2.43E-05 | 2.052027 |
| ENSMUSG00000085779 | *Atcayos* | 9.38E-24 | 2.05238 |
| ENSMUSG00000031231 | *Cox7b* | 6.58E-27 | 2.052756 |
| ENSMUSG00000049100 | *Pcdh10* | 8.51E-03 | 2.052949 |
| ENSMUSG00000085996 | *A830012C17Rik* | 1.30E-03 | 2.057668 |
| ENSMUSG00000034282 | *Evpl* | 8.05E-15 | 2.062518 |
| ENSMUSG00000027887 | *Sypl2* | 3.80E-11 | 2.066455 |
| ENSMUSG00000032503 | *Arpp21* | 1.46E-07 | 2.066522 |
| ENSMUSG00000047976 | *Kcna1* | 7.38E-12 | 2.070587 |
| ENSMUSG00000032066 | *Bco2* | 2.78E-04 | 2.073257 |
| ENSMUSG00000044359 | *P2ry4* | 1.91E-03 | 2.076374 |
| ENSMUSG00000093577 | *Gm20632* | 6.14E-03 | 2.077013 |
| ENSMUSG00000013483 | *Card14* | 1.31E-07 | 2.078412 |
| ENSMUSG00000000157 | *Itgb2l* | 9.04E-27 | 2.079323 |
| ENSMUSG00000043029 | *Trpv3* | 1.44E-02 | 2.081377 |
| ENSMUSG00000039238 | *Zfp750* | 1.47E-04 | 2.082281 |
| ENSMUSG00000110752 | *D730003K21Rik* | 4.64E-02 | 2.082774 |
| ENSMUSG00000028778 | *Hcrtr1* | 3.59E-02 | 2.085093 |
| ENSMUSG00000058207 | *Serpina3k* | 1.26E-09 | 2.085453 |
| ENSMUSG00000067081 | *Asb18* | 2.65E-08 | 2.090381 |

| | | | |
|---|---|---|---|
| ENSMUSG00000086628 | *Gm16157* | 1.73E-02 | 2.090745 |
| ENSMUSG00000024105 | *Themis3* | 5.35E-04 | 2.093314 |
| ENSMUSG00000061259 | *Tmprss11d* | 1.09E-03 | 2.098869 |
| ENSMUSG00000090698 | *Apold1* | 9.53E-29 | 2.09911 |
| ENSMUSG00000019102 | *Aldh3a1* | 7.05E-13 | 2.102843 |
| ENSMUSG00000083832 | *Gm13771* | 1.13E-07 | 2.104884 |
| ENSMUSG00000102630 | *Gm37289* | 1.55E-03 | 2.105392 |
| ENSMUSG00000110278 | *Gm5608* | 5.85E-19 | 2.108069 |
| ENSMUSG00000034768 | *Asb16* | 1.07E-02 | 2.111324 |
| ENSMUSG00000039956 | *Mrap* | 4.45E-10 | 2.112034 |
| ENSMUSG00000103923 | *Gm37896* | 8.60E-03 | 2.11783 |
| ENSMUSG00000097335 | *Gm26563* | 2.10E-03 | 2.118682 |
| ENSMUSG00000078907 | *Fam186b* | 3.97E-02 | 2.120724 |
| ENSMUSG00000059742 | *Kcnh7* | 5.20E-04 | 2.121118 |
| ENSMUSG00000069581 | *Tspear* | 4.61E-03 | 2.12407 |
| ENSMUSG00000111582 | *Gm47347* | 3.71E-02 | 2.125209 |
| ENSMUSG00000110101 | *Gm6249* | 2.05E-02 | 2.12537 |
| ENSMUSG00000114838 | *Gm48703* | 2.07E-02 | 2.125697 |
| ENSMUSG00000110622 | *Iqcn* | 5.56E-03 | 2.126382 |
| ENSMUSG00000032013 | *Trim29* | 7.42E-11 | 2.126553 |
| ENSMUSG00000019787 | *Trdn* | 6.78E-61 | 2.127457 |
| ENSMUSG00000103502 | *9330121J05Rik* | 4.80E-02 | 2.127771 |
| ENSMUSG00000059201 | *Lep* | 5.78E-05 | 2.128026 |
| ENSMUSG00000072573 | *Gm10369* | 3.96E-02 | 2.129896 |
| ENSMUSG00000035923 | *Myf6* | 2.65E-02 | 2.130758 |
| ENSMUSG00000028989 | *Angptl7* | 5.47E-12 | 2.131128 |
| ENSMUSG00000055775 | *Myh8* | 4.26E-21 | 2.138598 |
| ENSMUSG00000108436 | *Gm44851* | 3.51E-04 | 2.138805 |
| ENSMUSG00000100551 | *Gm19503* | 1.04E-09 | 2.138828 |
| ENSMUSG00000038670 | *Mybpc2* | 7.68E-24 | 2.148925 |
| ENSMUSG00000079243 | *Xirp1* | 5.72E-30 | 2.150145 |
| ENSMUSG00000104340 | *Gm10522* | 2.43E-02 | 2.151948 |
| ENSMUSG00000031448 | *Adprhl1* | 2.72E-17 | 2.153714 |
| ENSMUSG00000021359 | *Tfap2a* | 3.33E-04 | 2.154076 |
| ENSMUSG00000021702 | *Thbs4* | 2.51E-18 | 2.160411 |
| ENSMUSG00000082884 | *Gm13339* | 7.91E-11 | 2.160938 |
| ENSMUSG00000072845 | *Tmprss11a* | 1.18E-02 | 2.165947 |
| ENSMUSG00000025370 | *Cdh9* | 1.48E-02 | 2.168399 |
| ENSMUSG00000028356 | *Ambp* | 6.80E-03 | 2.16969 |
| ENSMUSG00000019851 | *Perp* | 7.61E-19 | 2.171313 |
| ENSMUSG00000108985 | *A930030B08Rik* | 5.86E-04 | 2.171742 |
| ENSMUSG00000089948 | *Far2os1* | 3.02E-02 | 2.174011 |
| ENSMUSG00000024973 | *Hrasls5* | 6.29E-03 | 2.174329 |
| ENSMUSG00000038751 | *Ptk6* | 2.90E-04 | 2.175162 |
| ENSMUSG00000055489 | *Ano5* | 2.65E-19 | 2.176035 |
| ENSMUSG00000036960 | *Clca2* | 8.46E-04 | 2.179341 |
| ENSMUSG00000064371 | *mt-Tt* | 3.00E-02 | 2.180972 |
| ENSMUSG00000111514 | *E230014E18Rik* | 6.75E-12 | 2.182891 |
| ENSMUSG00000086777 | *Far2os2* | 7.84E-04 | 2.185863 |
| ENSMUSG00000020892 | *Aloxe3* | 1.75E-06 | 2.187315 |
| ENSMUSG00000032726 | *Bmp8a* | 2.54E-03 | 2.1889 |
| ENSMUSG00000117092 | *AC166110.1* | 2.83E-03 | 2.190222 |
| ENSMUSG00000023176 | *Cpn2* | 2.07E-02 | 2.192113 |

| | | | |
|---|---|---|---|
| ENSMUSG00000027196 | *Alkbh3os1* | 3.46E-02 | 2.193943 |
| ENSMUSG00000032292 | *Nr2e3* | 4.12E-03 | 2.196921 |
| ENSMUSG00000042918 | *Mamstr* | 1.20E-37 | 2.197597 |
| ENSMUSG00000046056 | *Sbsn* | 2.83E-10 | 2.198103 |
| ENSMUSG00000027902 | *Chil6* | 1.73E-02 | 2.200008 |
| ENSMUSG00000091238 | *Gm17103* | 7.14E-04 | 2.200115 |
| ENSMUSG00000096351 | *Samd11* | 9.26E-09 | 2.20062 |
| ENSMUSG00000109157 | *Gm44829* | 6.38E-08 | 2.201996 |
| ENSMUSG00000042250 | *Pglyrp4* | 3.18E-03 | 2.203176 |
| ENSMUSG00000102759 | *Gm10463* | 4.03E-03 | 2.205769 |
| ENSMUSG00000007653 | *Gabrb2* | 1.16E-03 | 2.208491 |
| ENSMUSG00000047216 | *Cdh19* | 1.60E-26 | 2.20978 |
| ENSMUSG00000096887 | *Gm20594* | 7.85E-12 | 2.211799 |
| ENSMUSG00000045027 | *Prss22* | 4.56E-13 | 2.221258 |
| ENSMUSG00000041872 | *Il17f* | 3.11E-11 | 2.222913 |
| ENSMUSG00000104314 | *Gm37907* | 4.45E-02 | 2.226011 |
| ENSMUSG00000025754 | *Agbl1* | 1.47E-44 | 2.226428 |
| ENSMUSG00000019853 | *Hebp2* | 1.26E-03 | 2.230586 |
| ENSMUSG00000111818 | *Gm17749* | 3.92E-13 | 2.231279 |
| ENSMUSG00000047686 | *Rtl3* | 3.27E-02 | 2.237085 |
| ENSMUSG00000017607 | *Tns4* | 2.61E-07 | 2.237262 |
| ENSMUSG00000028435 | *Aqp3* | 1.87E-10 | 2.240723 |
| ENSMUSG00000034683 | *Ppp1r1c* | 4.89E-04 | 2.242066 |
| ENSMUSG00000031936 | *Hephl1* | 4.37E-03 | 2.253413 |
| ENSMUSG00000087477 | *Gm13822* | 2.34E-04 | 2.260881 |
| ENSMUSG00000046961 | *Gpr156* | 1.14E-11 | 2.261469 |
| ENSMUSG00000068890 | *Lce1a2* | 5.61E-03 | 2.26277 |
| ENSMUSG00000112023 | *Lilr4b* | 2.80E-15 | 2.264239 |
| ENSMUSG00000038020 | *Rapgefl1* | 7.31E-12 | 2.268906 |
| ENSMUSG00000042662 | *Dusp15* | 2.80E-02 | 2.282184 |
| ENSMUSG00000036123 | *Slc9a3* | 7.66E-05 | 2.282388 |
| ENSMUSG00000070385 | *Ampd1* | 2.67E-16 | 2.283597 |
| ENSMUSG00000111765 | *Gm10635* | 3.06E-03 | 2.287427 |
| ENSMUSG00000042895 | *Abra* | 4.53E-15 | 2.29493 |
| ENSMUSG00000040340 | *Tex45* | 2.95E-04 | 2.297836 |
| ENSMUSG00000027022 | *Xirp2* | 1.00E-64 | 2.303434 |
| ENSMUSG00000032281 | *Acsbg1* | 1.63E-15 | 2.307473 |
| ENSMUSG00000106746 | *2900064F13Rik* | 3.81E-02 | 2.307491 |
| ENSMUSG00000085564 | *Gm12198* | 3.19E-02 | 2.309238 |
| ENSMUSG00000000031 | *H19* | 9.37E-41 | 2.316057 |
| ENSMUSG00000017697 | *Ada* | 2.19E-32 | 2.316335 |
| ENSMUSG00000108964 | *Gm18600* | 2.02E-02 | 2.316741 |
| ENSMUSG00000040258 | *Nxph4* | 1.61E-02 | 2.322015 |
| ENSMUSG00000034634 | *Ly6d* | 7.55E-08 | 2.32215 |
| ENSMUSG00000085434 | *Gm11725* | 1.18E-05 | 2.32389 |
| ENSMUSG00000041827 | *Oasl1* | 1.27E-48 | 2.333376 |
| ENSMUSG00000046049 | *Rp1l1* | 8.88E-06 | 2.335651 |
| ENSMUSG00000029361 | *Nos1* | 2.34E-29 | 2.336301 |
| ENSMUSG00000022860 | *Chodl* | 3.02E-03 | 2.338522 |
| ENSMUSG00000025329 | *Padi1* | 3.46E-03 | 2.339451 |
| ENSMUSG00000004885 | *Crabp2* | 2.14E-05 | 2.342472 |
| ENSMUSG00000097076 | *Platr7* | 7.34E-03 | 2.342649 |
| ENSMUSG00000073602 | *Serpinb3b* | 7.91E-06 | 2.343508 |

| | | | |
|---|---|---|---|
| ENSMUSG00000049134 | *Nrap* | 7.61E-59 | 2.345431 |
| ENSMUSG00000048764 | *Tmprss11f* | 1.67E-05 | 2.348341 |
| ENSMUSG00000056966 | *Gjc3* | 3.67E-03 | 2.35148 |
| ENSMUSG00000068697 | *Myoz1* | 3.99E-14 | 2.351495 |
| ENSMUSG00000009070 | *Rsph14* | 5.67E-15 | 2.352471 |
| ENSMUSG00000022622 | *Acr* | 2.57E-18 | 2.353769 |
| ENSMUSG00000114457 | *Gm36346* | 1.55E-02 | 2.357304 |
| ENSMUSG00000037469 | *Acp7* | 6.67E-04 | 2.357911 |
| ENSMUSG00000096084 | *Gm21850* | 1.15E-11 | 2.358208 |
| ENSMUSG00000031972 | *Acta1* | 3.59E-28 | 2.359297 |
| ENSMUSG00000026983 | *Il1f5* | 5.52E-04 | 2.359573 |
| ENSMUSG00000059832 | *Kprp* | 5.63E-13 | 2.360512 |
| ENSMUSG00000043286 | *Pnpla1* | 1.03E-04 | 2.366505 |
| ENSMUSG00000033508 | *Asprv1* | 8.12E-12 | 2.36952 |
| ENSMUSG00000085511 | *Gm11738* | 1.72E-03 | 2.370938 |
| ENSMUSG00000044737 | *Klk14* | 4.19E-12 | 2.372258 |
| ENSMUSG00000111556 | *Gm19299* | 6.49E-04 | 2.375603 |
| ENSMUSG00000024771 | *Lipk* | 3.83E-03 | 2.37619 |
| ENSMUSG00000027376 | *Prom2* | 7.96E-09 | 2.376333 |
| ENSMUSG00000021123 | *Rdh12* | 6.55E-37 | 2.378531 |
| ENSMUSG00000043165 | *Lor* | 6.90E-19 | 2.380809 |
| ENSMUSG00000097069 | *Gm16998* | 3.57E-02 | 2.384026 |
| ENSMUSG00000060180 | *Myh13* | 2.82E-11 | 2.384598 |
| ENSMUSG00000026621 | *1-Mar* | 1.07E-02 | 2.385001 |
| ENSMUSG00000024331 | *Dsc2* | 7.84E-15 | 2.385649 |
| ENSMUSG00000095438 | *Mir133a-1hg* | 5.55E-12 | 2.386779 |
| ENSMUSG00000069873 | *4930438A08Rik* | 2.56E-06 | 2.387168 |
| ENSMUSG00000041476 | *Smpx* | 1.20E-12 | 2.388318 |
| ENSMUSG00000097281 | *Gm26685* | 4.14E-02 | 2.39049 |
| ENSMUSG00000022510 | *Trp63* | 1.37E-22 | 2.394718 |
| ENSMUSG00000034730 | *Adgrb1* | 2.15E-03 | 2.399098 |
| ENSMUSG00000108448 | *Gm21269* | 1.77E-02 | 2.402324 |
| ENSMUSG00000081619 | *Gm13351* | 4.66E-02 | 2.402361 |
| ENSMUSG00000097850 | *4631405K08Rik* | 3.02E-03 | 2.406628 |
| ENSMUSG00000109372 | *Gm19410* | 2.96E-02 | 2.407345 |
| ENSMUSG00000092222 | *Gm20506* | 1.38E-02 | 2.407749 |
| ENSMUSG00000064179 | *Tnnt1* | 6.04E-08 | 2.41389 |
| ENSMUSG00000050808 | *Muc15* | 4.42E-03 | 2.423088 |
| ENSMUSG00000030672 | *Mylpf* | 4.00E-11 | 2.425658 |
| ENSMUSG00000037942 | *Crp* | 3.22E-03 | 2.427037 |
| ENSMUSG00000075296 | *Aldh3b2* | 7.70E-05 | 2.42977 |
| ENSMUSG00000044499 | *Hs3st5* | 1.87E-15 | 2.429999 |
| ENSMUSG00000038357 | *Camp* | 4.41E-10 | 2.437799 |
| ENSMUSG00000081163 | *Defa-ps12* | 4.11E-03 | 2.439291 |
| ENSMUSG00000097413 | *A830052D11Rik* | 6.08E-05 | 2.440113 |
| ENSMUSG00000040283 | *Btnl9* | 2.24E-05 | 2.447538 |
| ENSMUSG00000074695 | *Il22* | 2.53E-04 | 2.451827 |
| ENSMUSG00000026592 | *Tex35* | 3.15E-12 | 2.456187 |
| ENSMUSG00000109311 | *AI314278* | 2.12E-05 | 2.458668 |
| ENSMUSG00000042092 | *Lce1c* | 1.91E-03 | 2.46598 |
| ENSMUSG00000061816 | *Myl1* | 1.65E-32 | 2.468997 |
| ENSMUSG00000027868 | *Tbx15* | 3.60E-32 | 2.473286 |
| ENSMUSG00000067001 | *Serpinb7* | 2.67E-03 | 2.476115 |

| | | | |
|---|---|---|---|
| ENSMUSG00000043461 | *Sptssb* | 4.59E-07 | 2.477048 |
| ENSMUSG00000067616 | *Klk11* | 2.13E-06 | 2.484335 |
| ENSMUSG00000032807 | *Alox12b* | 2.99E-08 | 2.487194 |
| ENSMUSG00000050830 | *Vwc2* | 9.16E-07 | 2.489155 |
| ENSMUSG00000057280 | *Musk* | 7.35E-26 | 2.491914 |
| ENSMUSG00000052581 | *Lrrtm4* | 1.35E-14 | 2.496839 |
| ENSMUSG00000026984 | *Il1f6* | 6.08E-04 | 2.49812 |
| ENSMUSG00000075394 | *Hoxc4* | 3.49E-02 | 2.499153 |
| ENSMUSG00000042761 | *Mrap2* | 3.01E-04 | 2.502625 |
| ENSMUSG00000027077 | *Smtnl1* | 6.53E-14 | 2.503067 |
| ENSMUSG00000023153 | *Tmem52* | 1.52E-03 | 2.503879 |
| ENSMUSG00000028865 | *Cd164l2* | 1.88E-06 | 2.505749 |
| ENSMUSG00000100147 | *1700047M11Rik* | 1.89E-21 | 2.506465 |
| ENSMUSG00000045019 | *Acer1* | 5.56E-03 | 2.51065 |
| ENSMUSG00000050359 | *Sprr1a* | 6.00E-11 | 2.511127 |
| ENSMUSG00000113178 | *Mylf-ps* | 7.37E-25 | 2.513042 |
| ENSMUSG00000028186 | *Uox* | 5.38E-03 | 2.521096 |
| ENSMUSG00000021506 | *Pitx1* | 4.13E-05 | 2.524928 |
| ENSMUSG00000100410 | *2310020H05Rik* | 5.52E-04 | 2.525092 |
| ENSMUSG00000099798 | *Gm29168* | 1.95E-02 | 2.530757 |
| ENSMUSG00000023473 | *Celsr3* | 5.94E-19 | 2.534816 |
| ENSMUSG00000042124 | *Lce1f* | 2.44E-03 | 2.541445 |
| ENSMUSG00000027470 | *Mylk2* | 2.72E-26 | 2.549178 |
| ENSMUSG00000044322 | *Dsc1* | 5.79E-06 | 2.554197 |
| ENSMUSG00000037139 | *Myom3* | 1.14E-36 | 2.554969 |
| ENSMUSG00000026327 | *Serpinb11* | 1.65E-06 | 2.557138 |
| ENSMUSG00000096960 | *A230028O05Rik* | 1.73E-18 | 2.567768 |
| ENSMUSG00000057609 | *Lce1a1* | 1.62E-04 | 2.572862 |
| ENSMUSG00000025064 | *Col17a1* | 5.34E-16 | 2.577045 |
| ENSMUSG00000024365 | *Cyp21a1* | 1.60E-02 | 2.581649 |
| ENSMUSG00000025317 | *Car5a* | 3.01E-02 | 2.589695 |
| ENSMUSG00000026418 | *Tnni1* | 3.68E-06 | 2.58989 |
| ENSMUSG00000061928 | *Dsg1b* | 2.34E-12 | 2.593438 |
| ENSMUSG00000114493 | *Gm47071* | 2.06E-15 | 2.5951 |
| ENSMUSG00000081689 | *Gm9166* | 1.74E-02 | 2.598563 |
| ENSMUSG00000028148 | *Them5* | 1.44E-11 | 2.59946 |
| ENSMUSG00000048455 | *Sprr1b* | 3.10E-04 | 2.603137 |
| ENSMUSG00000054582 | *Pabpc1l* | 3.75E-22 | 2.605057 |
| ENSMUSG00000047586 | *Nccrp1* | 1.96E-03 | 2.608167 |
| ENSMUSG00000103382 | *Gm37755* | 1.50E-03 | 2.610909 |
| ENSMUSG00000021768 | *Dusp13* | 4.51E-06 | 2.617201 |
| ENSMUSG00000021565 | *Slc6a19* | 1.45E-08 | 2.620258 |
| ENSMUSG00000069441 | *Dsg1a* | 1.77E-16 | 2.627138 |
| ENSMUSG00000029862 | *Clcn1* | 1.98E-18 | 2.631589 |
| ENSMUSG00000066897 | *Olfr872* | 7.98E-03 | 2.634158 |
| ENSMUSG00000038354 | *Ankrd35* | 3.43E-07 | 2.634991 |
| ENSMUSG00000016942 | *Tmprss6* | 2.91E-02 | 2.635952 |
| ENSMUSG00000074199 | *Krtdap* | 1.84E-07 | 2.63948 |
| ENSMUSG00000026450 | *Chit1* | 2.83E-05 | 2.645391 |
| ENSMUSG00000034774 | *Dsg1c* | 1.69E-03 | 2.648044 |
| ENSMUSG00000055561 | *Spink5* | 2.81E-21 | 2.649507 |
| ENSMUSG00000049593 | *Lce1h* | 1.12E-02 | 2.651785 |
| ENSMUSG00000087410 | *2310065F04Rik* | 1.84E-02 | 2.652839 |

| | | | |
|---|---|---|---|
| ENSMUSG00000031710 | *Ucp1* | 8.66E-12 | 2.653905 |
| ENSMUSG00000074001 | *Klhl40* | 2.22E-08 | 2.659211 |
| ENSMUSG00000031343 | *Gabra3* | 8.60E-12 | 2.660099 |
| ENSMUSG00000040154 | *Wfdc5* | 1.24E-05 | 2.663952 |
| ENSMUSG00000112041 | *9530020I12Rik* | 3.40E-05 | 2.669075 |
| ENSMUSG00000116482 | *Gm34152* | 3.36E-02 | 2.674711 |
| ENSMUSG00000059956 | *Serpinb12* | 2.21E-09 | 2.67661 |
| ENSMUSG00000060621 | *Nkpd1* | 5.88E-04 | 2.680095 |
| ENSMUSG00000063651 | *Cnfn* | 1.68E-15 | 2.693832 |
| ENSMUSG00000086980 | *Gm13791* | 4.89E-04 | 2.702456 |
| ENSMUSG00000040127 | *Sdr9c7* | 7.30E-03 | 2.70979 |
| ENSMUSG00000059230 | *Defb4* | 4.63E-11 | 2.713218 |
| ENSMUSG00000053675 | *Tgm5* | 2.41E-02 | 2.71792 |
| ENSMUSG00000112471 | *Gm5779* | 8.23E-05 | 2.719729 |
| ENSMUSG00000057003 | *Myh4* | 8.95E-42 | 2.71992 |
| ENSMUSG00000097768 | *2310043M15Rik* | 6.60E-05 | 2.725258 |
| ENSMUSG00000100781 | *Gm29481* | 3.64E-04 | 2.726074 |
| ENSMUSG00000026950 | *Neb* | 4.90E-43 | 2.726723 |
| ENSMUSG00000030592 | *Ryr1* | 3.70E-30 | 2.727653 |
| ENSMUSG00000034362 | *Csta1* | 4.08E-04 | 2.728502 |
| ENSMUSG00000111821 | *Gm48545* | 1.08E-07 | 2.729446 |
| ENSMUSG00000050296 | *Abca12* | 2.65E-14 | 2.739696 |
| ENSMUSG00000002500 | *Rpl3l* | 1.16E-12 | 2.745902 |
| ENSMUSG00000084228 | *Gm8080* | 1.10E-02 | 2.75149 |
| ENSMUSG00000041991 | *Hrnr* | 4.02E-11 | 2.752579 |
| ENSMUSG00000091376 | *Aadacl2* | 1.73E-03 | 2.759068 |
| ENSMUSG00000028011 | *Tdo2* | 8.06E-03 | 2.760958 |
| ENSMUSG00000084929 | *Foxo6os* | 9.14E-05 | 2.765202 |
| ENSMUSG00000090356 | *Teddm3* | 5.80E-06 | 2.767835 |
| ENSMUSG00000074489 | *Bglap3* | 9.79E-06 | 2.770307 |
| ENSMUSG00000061762 | *Tac1* | 9.92E-03 | 2.773699 |
| ENSMUSG00000031844 | *Hsd17b2* | 7.25E-08 | 2.776301 |
| ENSMUSG00000027401 | *Tgm3* | 2.87E-13 | 2.777067 |
| ENSMUSG00000041984 | *Rptn* | 1.46E-19 | 2.778259 |
| ENSMUSG00000048806 | *Ifnb1* | 9.77E-04 | 2.781116 |
| ENSMUSG00000058260 | *Serpina9* | 4.93E-03 | 2.790483 |
| ENSMUSG00000045539 | *Sprr3* | 1.26E-16 | 2.802851 |
| ENSMUSG00000030484 | *Lypd5* | 2.30E-12 | 2.803817 |
| ENSMUSG00000028396 | *2310002L09Rik* | 8.35E-07 | 2.810987 |
| ENSMUSG00000061723 | *Tnnt3* | 7.78E-31 | 2.811141 |
| ENSMUSG00000108371 | *Gm38832* | 4.21E-03 | 2.817575 |
| ENSMUSG00000074113 | *Gm10629* | 9.39E-03 | 2.826311 |
| ENSMUSG00000066510 | *Ankdd1a* | 1.97E-24 | 2.828707 |
| ENSMUSG00000031097 | *Tnni2* | 5.06E-35 | 2.82918 |
| ENSMUSG00000039518 | *Cdsn* | 2.66E-13 | 2.831018 |
| ENSMUSG00000043468 | *Adam30* | 2.93E-04 | 2.837999 |
| ENSMUSG00000106056 | *4930500L23Rik* | 4.77E-03 | 2.840989 |
| ENSMUSG00000045776 | *Lrtm1* | 1.50E-19 | 2.850709 |
| ENSMUSG00000043460 | *Elfn2* | 2.64E-03 | 2.850978 |
| ENSMUSG00000057454 | *Lypd3* | 1.13E-07 | 2.855151 |
| ENSMUSG00000018862 | *Otop3* | 4.00E-07 | 2.861507 |
| ENSMUSG00000068885 | *Lce3f* | 3.57E-09 | 2.863271 |
| ENSMUSG00000032401 | *Lctl* | 5.17E-03 | 2.865397 |

| | | | |
|---|---|---|---|
| ENSMUSG00000032373 | *Car12* | 2.44E-33 | 2.866191 |
| ENSMUSG00000046748 | *Tmem45a2* | 3.29E-15 | 2.875771 |
| ENSMUSG00000116659 | *AC158985.1* | 1.03E-03 | 2.879651 |
| ENSMUSG00000111555 | *Gm47326* | 4.82E-02 | 2.882505 |
| ENSMUSG00000079025 | *Gsdmc* | 9.10E-07 | 2.885077 |
| ENSMUSG00000056328 | *Myh1* | 2.08E-87 | 2.886996 |
| ENSMUSG00000027913 | *Crct1* | 2.20E-14 | 2.894169 |
| ENSMUSG00000020216 | *Jsrp1* | 1.18E-04 | 2.896039 |
| ENSMUSG00000071019 | *Sdr16c6* | 3.06E-05 | 2.897926 |
| ENSMUSG00000025172 | *Ankrd2* | 3.28E-18 | 2.90344 |
| ENSMUSG00000074433 | *Lce3e* | 9.33E-07 | 2.913285 |
| ENSMUSG00000036731 | *Cysrt1* | 8.18E-07 | 2.915053 |
| ENSMUSG00000010064 | *Slc38a3* | 5.21E-18 | 2.918093 |
| ENSMUSG00000030730 | *Atp2a1* | 5.75E-78 | 2.932671 |
| ENSMUSG00000085479 | *9430073C21Rik* | 5.71E-04 | 2.934549 |
| ENSMUSG00000059668 | *Krt4* | 4.18E-14 | 2.937284 |
| ENSMUSG00000114163 | *Gm32296* | 5.99E-04 | 2.941019 |
| ENSMUSG00000038295 | *Atg9b* | 6.02E-09 | 2.942834 |
| ENSMUSG00000099449 | *Gm28401* | 1.09E-04 | 2.950106 |
| ENSMUSG00000026413 | *Pkp1* | 5.96E-19 | 2.956086 |
| ENSMUSG00000028386 | *Slc46a2* | 8.03E-03 | 2.960449 |
| ENSMUSG00000039070 | *Cpa4* | 3.59E-11 | 2.967827 |
| ENSMUSG00000037977 | *6430571L13Rik* | 1.15E-12 | 2.986157 |
| ENSMUSG00000110426 | *Gm45757* | 2.83E-02 | 2.990865 |
| ENSMUSG00000032454 | *Rbp2* | 9.49E-04 | 2.991761 |
| ENSMUSG00000073601 | *Serpinb3c* | 2.89E-05 | 2.992627 |
| ENSMUSG00000029154 | *Cwh43* | 3.21E-04 | 2.995754 |
| ENSMUSG00000054325 | *Lce3a* | 3.84E-11 | 2.996653 |
| ENSMUSG00000105160 | *A530030E21Rik* | 7.94E-03 | 3.00639 |
| ENSMUSG00000074183 | *Gsta1* | 1.18E-03 | 3.011134 |
| ENSMUSG00000068889 | *Lce1e* | 3.36E-03 | 3.016024 |
| ENSMUSG00000020061 | *Mybpc1* | 4.58E-66 | 3.020577 |
| ENSMUSG00000027048 | *Abcb11* | 2.16E-02 | 3.040284 |
| ENSMUSG00000039269 | *2300002M23Rik* | 1.76E-06 | 3.045743 |
| ENSMUSG00000117222 | *AC060761.1* | 8.51E-16 | 3.048258 |
| ENSMUSG00000025129 | *Ppp1r27* | 6.54E-04 | 3.050154 |
| ENSMUSG00000020908 | *Myh3* | 2.90E-04 | 3.053928 |
| ENSMUSG00000097364 | *Gm26719* | 2.09E-05 | 3.072513 |
| ENSMUSG00000054422 | *Fabp1* | 5.44E-04 | 3.078014 |
| ENSMUSG00000044041 | *Krt13* | 2.04E-15 | 3.079995 |
| ENSMUSG00000063821 | *Dupd1* | 1.16E-02 | 3.080693 |
| ENSMUSG00000043430 | *Psapl1* | 2.65E-14 | 3.082218 |
| ENSMUSG00000044594 | *Serpinb3a* | 5.68E-13 | 3.087602 |
| ENSMUSG00000085532 | *B430319H21Rik* | 1.32E-14 | 3.09774 |
| ENSMUSG00000042031 | *Lce3b* | 1.13E-11 | 3.112196 |
| ENSMUSG00000005355 | *Casp14* | 2.13E-04 | 3.113601 |
| ENSMUSG00000087038 | *2900079G21Rik* | 4.50E-03 | 3.1195 |
| ENSMUSG00000114922 | *Ppifos* | 2.43E-06 | 3.120941 |
| ENSMUSG00000031372 | *Trex2* | 1.04E-04 | 3.124283 |
| ENSMUSG00000086584 | *Gm12002* | 6.34E-13 | 3.127654 |
| ENSMUSG00000071858 | *Gm94* | 2.60E-09 | 3.127964 |
| ENSMUSG00000092586 | *Ly6g6c* | 8.86E-11 | 3.133452 |
| ENSMUSG00000101133 | *Gm29050* | 2.40E-04 | 3.136949 |

| | | | |
|---|---|---|---|
| ENSMUSG00000048399 | *Tprg* | 1.51E-09 | 3.169198 |
| ENSMUSG00000045475 | *Lce3c* | 3.14E-12 | 3.171884 |
| ENSMUSG00000110735 | *Gm47173* | 8.51E-03 | 3.184631 |
| ENSMUSG00000054146 | *Krt15* | 8.74E-19 | 3.18484 |
| ENSMUSG00000103243 | *Lce1d* | 2.93E-06 | 3.187473 |
| ENSMUSG00000026407 | *Cacna1s* | 1.08E-45 | 3.18982 |
| ENSMUSG00000024471 | *Myot* | 5.42E-54 | 3.196349 |
| ENSMUSG00000030713 | *Klk7* | 1.82E-04 | 3.197459 |
| ENSMUSG00000097214 | *Gm19791* | 2.77E-02 | 3.223603 |
| ENSMUSG00000068893 | *Sprr2a2* | 2.58E-13 | 3.229494 |
| ENSMUSG00000050108 | *Bpifc* | 5.73E-05 | 3.240719 |
| ENSMUSG00000028236 | *Sdr16c5* | 1.03E-04 | 3.243863 |
| ENSMUSG00000078664 | *Sprr2a1* | 4.79E-06 | 3.253984 |
| ENSMUSG00000033268 | *Duox1* | 2.09E-12 | 3.257037 |
| ENSMUSG00000063130 | *Calml3* | 1.60E-17 | 3.258145 |
| ENSMUSG00000050463 | *Krt78* | 7.67E-32 | 3.271363 |
| ENSMUSG00000096269 | *Rpl31-ps21* | 2.93E-03 | 3.274694 |
| ENSMUSG00000074652 | *Myh7b* | 5.39E-06 | 3.28996 |
| ENSMUSG00000061527 | *Krt5* | 7.51E-19 | 3.292118 |
| ENSMUSG00000030155 | *Clec2e* | 8.29E-03 | 3.314901 |
| ENSMUSG00000047884 | *Klk9* | 1.29E-04 | 3.323011 |
| ENSMUSG00000114091 | *Gm40655* | 1.54E-09 | 3.328511 |
| ENSMUSG00000053522 | *Lgals7* | 1.35E-10 | 3.329974 |
| ENSMUSG00000017204 | *Gsdma* | 4.28E-12 | 3.335837 |
| ENSMUSG00000087090 | *Nctc1* | 2.38E-59 | 3.337836 |
| ENSMUSG00000096001 | *2610528A11Rik* | 4.77E-04 | 3.356603 |
| ENSMUSG00000062826 | *Ces2f* | 1.27E-20 | 3.364975 |
| ENSMUSG00000031344 | *Gabrq* | 9.36E-03 | 3.374101 |
| ENSMUSG00000085257 | *Gm13264* | 2.59E-02 | 3.380709 |
| ENSMUSG00000082141 | *Gm11212* | 1.11E-03 | 3.415103 |
| ENSMUSG00000067006 | *Serpinb5* | 3.69E-13 | 3.437795 |
| ENSMUSG00000074156 | *Ces1h* | 2.13E-11 | 3.444764 |
| ENSMUSG00000046676 | *Lce1l* | 1.08E-02 | 3.449777 |
| ENSMUSG00000021364 | *Elovl2* | 1.25E-03 | 3.464127 |
| ENSMUSG00000074445 | *Sprr2a3* | 8.74E-06 | 3.468969 |
| ENSMUSG00000078964 | *Ces1b* | 2.78E-06 | 3.472679 |
| ENSMUSG00000045545 | *Krt14* | 4.41E-27 | 3.472881 |
| ENSMUSG00000059898 | *Dsc3* | 2.86E-17 | 3.473919 |
| ENSMUSG00000033765 | *Calm4* | 5.17E-10 | 3.476147 |
| ENSMUSG00000105961 | *Gm40123* | 8.89E-03 | 3.493003 |
| ENSMUSG00000048721 | *Fndc9* | 1.16E-03 | 3.493393 |
| ENSMUSG00000017300 | *Tnnc2* | 2.61E-56 | 3.493517 |
| ENSMUSG00000101840 | *Gm28294* | 1.22E-02 | 3.494083 |
| ENSMUSG00000046095 | *Krt32* | 6.38E-05 | 3.511431 |
| ENSMUSG00000043795 | *Prr33* | 3.14E-07 | 3.513074 |
| ENSMUSG00000074647 | *Fam83c* | 1.30E-04 | 3.523902 |
| ENSMUSG00000091172 | *Gm17120* | 7.31E-03 | 3.529014 |
| ENSMUSG00000110207 | *Gm33586* | 2.10E-02 | 3.530274 |
| ENSMUSG00000051965 | *Nanos2* | 3.22E-04 | 3.536107 |
| ENSMUSG00000051431 | *Gpr87* | 7.37E-05 | 3.549606 |
| ENSMUSG00000025229 | *Pitx3* | 6.86E-03 | 3.553927 |
| ENSMUSG00000005716 | *Pvalb* | 2.81E-11 | 3.631134 |
| ENSMUSG00000031757 | *Mt4* | 1.87E-13 | 3.699945 |

| | | | |
|---|---|---|---|
| ENSMUSG00000058354 | *Krt6a* | 8.73E-08 | 3.701633 |
| ENSMUSG00000026985 | *Il1f8* | 4.85E-06 | 3.747629 |
| ENSMUSG00000055301 | *Adh7* | 9.72E-31 | 3.762133 |
| ENSMUSG00000097770 | *Gm26776* | 2.69E-05 | 3.767514 |
| ENSMUSG00000115270 | *5430430K15Rik* | 1.03E-02 | 3.813184 |
| ENSMUSG00000075217 | *4833423E24Rik* | 7.79E-13 | 3.828324 |
| ENSMUSG00000043472 | *Lce3d* | 9.57E-05 | 3.852596 |
| ENSMUSG00000030244 | *Gys2* | 3.13E-03 | 3.866888 |
| ENSMUSG00000056078 | *Lipm* | 5.55E-12 | 3.878651 |
| ENSMUSG00000047428 | *Dlk2* | 6.06E-04 | 3.942878 |
| ENSMUSG00000013766 | *Ly6g6e* | 6.75E-03 | 3.95354 |
| ENSMUSG00000017950 | *Hnf4a* | 2.61E-04 | 4.02923 |
| ENSMUSG00000025194 | *Abcc2* | 1.91E-04 | 4.03213 |
| ENSMUSG00000030046 | *Bmp10* | 5.70E-04 | 4.173749 |
| ENSMUSG00000056632 | *Dsg3* | 2.00E-33 | 4.176681 |
| ENSMUSG00000030237 | *Slco1a4* | 4.96E-03 | 4.192899 |
| ENSMUSG00000028001 | *Fga* | 1.73E-08 | 4.260873 |
| ENSMUSG00000109685 | *Gm45912* | 2.62E-03 | 4.273195 |
| ENSMUSG00000111709 | *Gm3776* | 1.46E-04 | 4.284462 |
| ENSMUSG00000043681 | *Fam25c* | 7.65E-12 | 4.287186 |
| ENSMUSG00000079451 | *Tmprss11g* | 6.62E-12 | 4.355436 |
| ENSMUSG00000097189 | *Gm26594* | 1.21E-02 | 4.459896 |
| ENSMUSG00000025431 | *Crisp1* | 1.13E-13 | 4.600625 |
| ENSMUSG00000023041 | *Krt6b* | 7.88E-05 | 4.619968 |
| ENSMUSG00000011632 | *Pinlyp* | 1.13E-03 | 4.658043 |
| ENSMUSG00000022875 | *Kng1* | 9.90E-06 | 4.667574 |
| ENSMUSG00000035031 | *C8a* | 2.65E-03 | 4.905581 |
| ENSMUSG00000044748 | *Defb1* | 2.48E-08 | 5.134939 |
| ENSMUSG00000102098 | *2310016D03Rik* | 3.16E-06 | 5.282878 |
| ENSMUSG00000035861 | *Tmprss11b* | 1.93E-05 | 5.434747 |
| ENSMUSG00000027761 | *Aadac* | 2.12E-05 | 5.483728 |
| ENSMUSG00000029019 | *Nppb* | 2.05E-26 | 5.500836 |
| ENSMUSG00000033196 | *Myh2* | 1.31E-45 | 5.509976 |
| ENSMUSG00000027261 | *Hao1* | 4.23E-11 | 5.543845 |
| ENSMUSG00000070719 | *Pla2g4d* | 3.51E-07 | 5.701431 |
| ENSMUSG00000025003 | *Cyp2c39* | 1.58E-04 | 6.085577 |
| ENSMUSG00000004872 | *Pax3* | 1.87E-06 | 6.162959 |
| ENSMUSG00000016327 | *Atp1b4* | 4.64E-12 | 6.441707 |
| ENSMUSG00000027249 | *F2* | 5.25E-04 | 6.44469 |
| ENSMUSG00000033860 | *Fgg* | 1.30E-15 | 6.79552 |
| ENSMUSG00000056656 | *Apol8* | 2.35E-27 | 6.857002 |
| ENSMUSG00000073375 | *Lrrc30* | 2.39E-04 | 7.131156 |
| ENSMUSG00000024512 | *Dynap* | 2.79E-05 | 7.653523 |
| ENSMUSG00000001670 | *Tat* | 1.65E-05 | 8.104061 |
| ENSMUSG00000029671 | *Wnt16* | 5.12E-05 | 8.122169 |
| ENSMUSG00000019932 | *Kera* | 9.32E-11 | 9.197208 |
| ENSMUSG00000115207 | *Gm49132* | 6.77E-08 | 9.321712 |
| ENSMUSG00000030131 | *Mug2* | 3.68E-08 | 10.01166 |
| ENSMUSG00000020609 | *Apob* | 1.70E-16 | 10.07445 |
| ENSMUSG00000003053 | *Cyp2c29* | 1.58E-09 | 11.55749 |
| ENSMUSG00000022868 | *Ahsg* | 1.79E-19 | 12.68437 |
| ENSMUSG00000035540 | *Gc* | 2.58E-14 | 14.12206 |
| ENSMUSG00000030359 | *Pzp* | 3.60E-20 | 14.37129 |

| | | | |
|---|---|---|---|
| ENSMUSG00000059481 | *Plg* | 3.52E-08 | 19.5621 |
| ENSMUSG00000057400 | *Ces1c* | 8.80E-07 | 19.90682 |
| ENSMUSG00000028307 | *Aldob* | 1.46E-10 | 20.84021 |
| ENSMUSG00000005681 | *Apoa2* | 3.99E-05 | 38.11525 |
| ENSMUSG00000059908 | *Mug1* | 1.99E-19 | 49.75901 |
| ENSMUSG00000029630 | *Cyp3a25* | 3.05E-06 | 60.85914 |
| ENSMUSG00000056035 | *Cyp3a11* | 2.89E-12 | 61.11608 |
| ENSMUSG00000025991 | *Cps1* | 1.52E-07 | 68.002 |
| ENSMUSG00000061808 | *Ttr* | 9.12E-07 | 72.79417 |
| ENSMUSG00000033831 | *Fgb* | 2.27E-10 | 80.34186 |
| ENSMUSG00000032083 | *Apoa1* | 1.86E-13 | 85.92226 |
| ENSMUSG00000029368 | *Alb* | 8.71E-39 | 89.99856 |

**Supplementary Table 11.** Murine differential gene expression in the lungs 6 h post-infection, 9-47M vs 4559M. Genes with fold change (FC) greater than 1.5 and $p < 0.05$ are shown. FC values highlighted in blue = upregulated in 9-47M, while values highlighted in red = upregulated in 4559M.

| **Gene Stable ID** | **Gene Name** | **padj** | **FC** |
|---|---|---|---|
| ENSMUSG00000085261 | *Gm13814* | 1.01E-05 | 5.162232 |
| ENSMUSG00000031551 | *Ido1* | 1.63E-07 | 4.116956 |
| ENSMUSG00000071679 | *Rtl4* | 1.67E-17 | 4.081086 |
| ENSMUSG00000063594 | *Gng8* | 7.78E-04 | 3.837051 |
| ENSMUSG00000009900 | *Wnt3a* | 3.97E-16 | 3.498319 |
| ENSMUSG00000092591 | *Gm20429* | 6.56E-04 | 3.202419 |
| ENSMUSG00000115574 | *Gm49245* | 6.79E-04 | 3.163799 |
| ENSMUSG00000115307 | *Gm49313* | 7.11E-07 | 2.940982 |
| ENSMUSG00000032502 | *Stac* | 7.16E-13 | 2.939761 |
| ENSMUSG00000085982 | *9530051G07Rik* | 4.37E-06 | 2.908872 |
| ENSMUSG00000085235 | *Gm12576* | 1.23E-05 | 2.830949 |
| ENSMUSG00000030114 | *Klrg1* | 5.25E-03 | 2.807412 |
| ENSMUSG00000068428 | *Gmnc* | 8.48E-03 | 2.716893 |
| ENSMUSG00000001494 | *Sost* | 1.53E-04 | 2.681953 |
| ENSMUSG00000020081 | *Tacr2* | 7.50E-03 | 2.609488 |
| ENSMUSG00000105402 | *Gm3716* | 8.05E-03 | 2.563662 |
| ENSMUSG00000108617 | *Gm31749* | 3.03E-03 | 2.550466 |
| ENSMUSG00000060275 | *Nrg2* | 1.89E-11 | 2.543613 |
| ENSMUSG00000114028 | *9630002D21Rik* | 2.18E-05 | 2.514966 |
| ENSMUSG00000048485 | *Zbtb8b* | 1.10E-02 | 2.476257 |
| ENSMUSG00000076613 | *Ighg2b* | 8.81E-04 | 2.473764 |
| ENSMUSG00000053961 | *Ang5* | 8.04E-04 | 2.455475 |
| ENSMUSG00000092389 | *Gm20483* | 1.60E-02 | 2.454367 |
| ENSMUSG00000090166 | *Ear10* | 5.22E-10 | 2.447762 |
| ENSMUSG00000114566 | *Gm48500* | 3.34E-05 | 2.443691 |
| ENSMUSG00000087500 | *Gm12426* | 1.87E-03 | 2.430623 |
| ENSMUSG00000086843 | *E030013I19Rik* | 9.29E-08 | 2.368294 |
| ENSMUSG00000061171 | *Slc38a11* | 6.50E-05 | 2.368149 |
| ENSMUSG00000087355 | *Gm13187* | 2.64E-02 | 2.35886 |
| ENSMUSG00000076612 | *Ighg2c* | 2.96E-02 | 2.329747 |

| | | | |
|---|---|---|---|
| ENSMUSG00000079436 | *Kcnj13* | 3.92E-03 | 2.327287 |
| ENSMUSG00000052026 | *Slc6a7* | 2.97E-08 | 2.315711 |
| ENSMUSG00000070530 | *Wfdc16* | 1.76E-02 | 2.314729 |
| ENSMUSG00000109056 | *A630009H07Rik* | 2.27E-02 | 2.292291 |
| ENSMUSG00000056824 | *Zfp663* | 6.59E-04 | 2.279782 |
| ENSMUSG00000072599 | *Ear-ps2* | 2.14E-05 | 2.246685 |
| ENSMUSG00000009394 | *Syn2* | 3.23E-27 | 2.246363 |
| ENSMUSG00000051969 | *Tlr11* | 1.09E-02 | 2.239618 |
| ENSMUSG00000086313 | *Gm15940* | 1.71E-03 | 2.238922 |
| ENSMUSG00000036295 | *Lrrn3* | 1.33E-06 | 2.227998 |
| ENSMUSG00000085882 | *2610507I01Rik* | 1.42E-05 | 2.211385 |
| ENSMUSG00000116097 | *Gm36738* | 2.22E-04 | 2.2066 |
| ENSMUSG00000030041 | *M1ap* | 6.18E-16 | 2.204481 |
| ENSMUSG00000011263 | *Exoc3l2* | 4.87E-04 | 2.194716 |
| ENSMUSG00000086822 | *5330413P13Rik* | 1.81E-03 | 2.193201 |
| ENSMUSG00000102082 | *Gm28675* | 9.97E-04 | 2.179205 |
| ENSMUSG00000099338 | *2810030D12Rik* | 7.38E-06 | 2.170948 |
| ENSMUSG00000030495 | *Slc7a10* | 8.33E-20 | 2.14737 |
| ENSMUSG00000087042 | *Gm11611* | 2.21E-02 | 2.145158 |
| ENSMUSG00000099553 | *Gm29538* | 2.37E-06 | 2.13579 |
| ENSMUSG00000066319 | *Rtp3* | 5.27E-08 | 2.127259 |
| ENSMUSG00000041479 | *Syt15* | 4.13E-10 | 2.125265 |
| ENSMUSG00000110427 | *4933406B17Rik* | 1.98E-02 | 2.10939 |
| ENSMUSG00000025789 | *St8sia2* | 8.02E-14 | 2.104873 |
| ENSMUSG00000028755 | *Cda* | 1.82E-03 | 2.099281 |
| ENSMUSG00000026304 | *Rab17* | 1.54E-02 | 2.091565 |
| ENSMUSG00000029193 | *Cckar* | 1.80E-08 | 2.087954 |
| ENSMUSG00000019909 | *Fam162b* | 9.91E-04 | 2.085813 |
| ENSMUSG00000039179 | *Tekt5* | 3.66E-03 | 2.083175 |
| ENSMUSG00000044633 | *B530045E10Rik* | 2.46E-24 | 2.082099 |
| ENSMUSG00000108084 | *Gm7932* | 3.36E-02 | 2.079352 |
| ENSMUSG00000060615 | *Ang4* | 3.19E-02 | 2.073621 |
| ENSMUSG00000000901 | *Mmp11* | 1.21E-09 | 2.071062 |
| ENSMUSG00000051920 | *Rspo2* | 3.77E-02 | 2.060566 |
| ENSMUSG00000024968 | *Rcor2* | 7.64E-03 | 2.054541 |
| ENSMUSG00000048988 | *Elfn1* | 2.26E-18 | 2.049999 |
| ENSMUSG00000028919 | *Arhgef19* | 1.84E-03 | 2.045006 |
| ENSMUSG00000085282 | *Gm15663* | 1.26E-03 | 2.042585 |
| ENSMUSG00000101605 | *Ace3* | 1.64E-08 | 2.033861 |
| ENSMUSG00000063730 | *Hsd3b2* | 4.33E-03 | 2.031267 |
| ENSMUSG00000037962 | *Rflna* | 9.51E-05 | 2.029917 |
| ENSMUSG00000048455 | *Sprr1b* | 4.70E-02 | 2.029009 |
| ENSMUSG00000054622 | *D730045B01Rik* | 3.09E-03 | 2.018059 |
| ENSMUSG00000092353 | *Gm20539* | 1.07E-02 | 2.004409 |
| ENSMUSG00000029601 | *Iqcd* | 9.80E-03 | 2.001456 |
| ENSMUSG00000026109 | *Tmeff2* | 3.23E-11 | 1.998603 |
| ENSMUSG00000009114 | *2610028H24Rik* | 2.03E-09 | 1.997699 |
| ENSMUSG00000029477 | *Morn3* | 2.44E-03 | 1.99663 |
| ENSMUSG00000074505 | *Fat3* | 3.95E-40 | 1.986612 |
| ENSMUSG00000108481 | *Gm33248* | 9.66E-17 | 1.983101 |
| ENSMUSG00000026494 | *Kif26b* | 6.07E-29 | 1.976902 |
| ENSMUSG00000103857 | *Gm37249* | 2.44E-02 | 1.97662 |
| ENSMUSG00000055945 | *Prr18* | 2.04E-05 | 1.976009 |

| | | | |
|---|---|---|---|
| ENSMUSG00000087142 | *Gm12454* | 1.40E-03 | 1.974069 |
| ENSMUSG00000022491 | *Glycam1* | 9.58E-05 | 1.972925 |
| ENSMUSG00000070720 | *Tmem200b* | 2.56E-04 | 1.963692 |
| ENSMUSG00000116660 | *CT030736.1* | 1.35E-13 | 1.960853 |
| ENSMUSG00000020218 | *Wif1* | 3.37E-14 | 1.957565 |
| ENSMUSG00000047085 | *Lrrc4b* | 1.22E-05 | 1.939777 |
| ENSMUSG00000027656 | *Wisp2* | 6.19E-47 | 1.938229 |
| ENSMUSG00000027380 | *Acoxl* | 1.13E-11 | 1.935006 |
| ENSMUSG00000017007 | *Rbpjl* | 2.17E-14 | 1.933213 |
| ENSMUSG00000033533 | *Acsm1* | 1.08E-07 | 1.933132 |
| ENSMUSG00000079039 | *Gm11037* | 3.17E-02 | 1.927416 |
| ENSMUSG00000092004 | *Gm17482* | 3.73E-05 | 1.925619 |
| ENSMUSG00000052353 | *Cemip* | 1.08E-23 | 1.921299 |
| ENSMUSG00000067714 | *Lpar5* | 4.75E-02 | 1.916894 |
| ENSMUSG00000086368 | *Gm13830* | 1.34E-04 | 1.915922 |
| ENSMUSG00000107230 | *Gm19265* | 2.82E-03 | 1.914934 |
| ENSMUSG00000022504 | *Ciita* | 3.98E-26 | 1.914004 |
| ENSMUSG00000050288 | *Fzd2* | 1.07E-05 | 1.912167 |
| ENSMUSG00000086585 | *Gm16126* | 3.81E-02 | 1.91092 |
| ENSMUSG00000040841 | *Six5* | 6.58E-09 | 1.910496 |
| ENSMUSG00000084998 | *Gm16279* | 9.76E-04 | 1.908554 |
| ENSMUSG00000072596 | *Ear2* | 7.41E-14 | 1.907457 |
| ENSMUSG00000063681 | *Crb1* | 3.66E-02 | 1.90717 |
| ENSMUSG00000026870 | *Cutal* | 7.86E-04 | 1.907001 |
| ENSMUSG00000105547 | *Iglc3* | 3.49E-02 | 1.904731 |
| ENSMUSG00000103824 | *Gm38177* | 4.20E-04 | 1.893794 |
| ENSMUSG00000034324 | *Tmem132c* | 6.54E-30 | 1.893682 |
| ENSMUSG00000108390 | *Gm39038* | 1.42E-02 | 1.893336 |
| ENSMUSG00000097591 | *A330032B11Rik* | 2.92E-03 | 1.889618 |
| ENSMUSG00000108195 | *Gm44415* | 1.81E-02 | 1.887889 |
| ENSMUSG00000038233 | *Fam198a* | 3.94E-07 | 1.883042 |
| ENSMUSG00000103839 | *Gm37607* | 3.89E-02 | 1.879446 |
| ENSMUSG00000030043 | *Tacr1* | 4.32E-06 | 1.878906 |
| ENSMUSG00000074480 | *Mex3a* | 5.57E-03 | 1.877201 |
| ENSMUSG00000063260 | *Syt10* | 2.50E-02 | 1.87586 |
| ENSMUSG00000072601 | *Ear1* | 3.55E-09 | 1.87466 |
| ENSMUSG00000025577 | *Cbx2* | 1.79E-04 | 1.874541 |
| ENSMUSG00000043110 | *Lrrn4* | 8.40E-06 | 1.874015 |
| ENSMUSG00000045087 | *S1pr5* | 2.29E-03 | 1.872791 |
| ENSMUSG00000085069 | *Gm13111* | 2.88E-04 | 1.871815 |
| ENSMUSG00000014543 | *Klra17* | 1.24E-12 | 1.871249 |
| ENSMUSG00000092515 | *C87198* | 2.73E-02 | 1.870875 |
| ENSMUSG00000114161 | *Gm48662* | 1.37E-02 | 1.868726 |
| ENSMUSG00000108289 | *Gm44174* | 3.34E-08 | 1.868318 |
| ENSMUSG00000034467 | *Dynlrb2* | 1.07E-04 | 1.864633 |
| ENSMUSG00000030125 | *Lrrc23* | 3.21E-07 | 1.862472 |
| ENSMUSG00000037492 | *Zmat4* | 1.44E-11 | 1.858985 |
| ENSMUSG00000089713 | *Gm16564* | 3.70E-02 | 1.858329 |
| ENSMUSG00000110380 | *Gm45332* | 2.12E-03 | 1.856669 |
| ENSMUSG00000026616 | *Cr2* | 1.36E-07 | 1.856455 |
| ENSMUSG00000027831 | *Veph1* | 6.53E-14 | 1.85639 |
| ENSMUSG00000108580 | *Gm39094* | 9.91E-27 | 1.847514 |
| ENSMUSG00000085419 | *Gm11734* | 9.18E-03 | 1.8441 |

| | | | |
|---|---|---|---|
| ENSMUSG00000024440 | *Pcdh12* | 1.29E-20 | 1.843863 |
| ENSMUSG00000044952 | *Kctd21* | 5.91E-10 | 1.842085 |
| ENSMUSG00000116609 | *AC122141.1* | 1.30E-02 | 1.840951 |
| ENSMUSG00000086154 | *Gm16196* | 1.33E-02 | 1.840383 |
| ENSMUSG00000046242 | *Nme9* | 1.23E-02 | 1.838804 |
| ENSMUSG00000063320 | *1190007I07Rik* | 1.48E-02 | 1.836405 |
| ENSMUSG00000086390 | *1810019D21Rik* | 4.43E-07 | 1.836126 |
| ENSMUSG00000087518 | *Gm13561* | 5.48E-03 | 1.835776 |
| ENSMUSG00000100041 | *Gm5702* | 1.11E-02 | 1.835727 |
| ENSMUSG00000102175 | *Gm6119* | 1.10E-03 | 1.833252 |
| ENSMUSG00000087514 | *Gm16076* | 7.80E-03 | 1.831068 |
| ENSMUSG00000079355 | *Ackr4* | 1.42E-05 | 1.830347 |
| ENSMUSG00000021234 | *Fam161b* | 1.93E-04 | 1.828797 |
| ENSMUSG00000097339 | *Gm26671* | 6.23E-05 | 1.828632 |
| ENSMUSG00000046182 | *Gsg1l* | 2.43E-05 | 1.826043 |
| ENSMUSG00000047787 | *Flrt1* | 4.73E-02 | 1.825861 |
| ENSMUSG00000021708 | *Rasgrf2* | 7.04E-24 | 1.822346 |
| ENSMUSG00000029651 | *Mtus2* | 3.04E-27 | 1.820688 |
| ENSMUSG00000079173 | *Zan* | 1.05E-03 | 1.82063 |
| ENSMUSG00000042256 | *Ptchd4* | 8.22E-05 | 1.819936 |
| ENSMUSG00000063430 | *Wscd2* | 1.07E-09 | 1.81492 |
| ENSMUSG00000104168 | *Gm38250* | 8.48E-04 | 1.810521 |
| ENSMUSG00000103738 | *Gm37652* | 6.73E-04 | 1.809033 |
| ENSMUSG00000056155 | *Nanos3* | 8.78E-04 | 1.806709 |
| ENSMUSG00000044338 | *Aplnr* | 1.37E-34 | 1.806317 |
| ENSMUSG00000038402 | *Foxf2* | 1.43E-09 | 1.806283 |
| ENSMUSG00000110660 | *Gm5362* | 1.46E-02 | 1.804937 |
| ENSMUSG00000020642 | *Rnf144a* | 4.67E-29 | 1.804772 |
| ENSMUSG00000086283 | *2810433D01Rik* | 4.15E-02 | 1.804678 |
| ENSMUSG00000033249 | *Hsf4* | 2.62E-04 | 1.798369 |
| ENSMUSG00000105376 | *Gm36535* | 8.45E-03 | 1.797718 |
| ENSMUSG00000030098 | *Grip2* | 2.83E-09 | 1.795977 |
| ENSMUSG00000027796 | *Smad9* | 7.70E-08 | 1.788007 |
| ENSMUSG00000052563 | *D930048N14Rik* | 1.75E-06 | 1.786953 |
| ENSMUSG00000045555 | *Mettl24* | 4.87E-17 | 1.786089 |
| ENSMUSG00000039037 | *St6galnac5* | 9.72E-33 | 1.782983 |
| ENSMUSG00000084866 | *A930006K02Rik* | 7.02E-03 | 1.779173 |
| ENSMUSG00000050022 | *Amz1* | 1.30E-06 | 1.779075 |
| ENSMUSG00000115333 | *Gm34590* | 6.00E-03 | 1.77317 |
| ENSMUSG00000091415 | *Ak9* | 7.15E-10 | 1.77086 |
| ENSMUSG00000036923 | *Stox1* | 1.49E-08 | 1.769288 |
| ENSMUSG00000025902 | *Sox17* | 1.61E-18 | 1.76508 |
| ENSMUSG00000026301 | *Iqca* | 1.77E-05 | 1.764688 |
| ENSMUSG00000079330 | *Lemd1* | 1.10E-05 | 1.764372 |
| ENSMUSG00000057123 | *Gja5* | 5.68E-21 | 1.76359 |
| ENSMUSG00000112468 | *Gm47720* | 6.31E-08 | 1.762303 |
| ENSMUSG00000027932 | *Slc27a3* | 3.01E-04 | 1.757901 |
| ENSMUSG00000027318 | *Adam33* | 5.00E-08 | 1.756879 |
| ENSMUSG00000034584 | *Exph5* | 9.73E-20 | 1.754362 |
| ENSMUSG00000042359 | *Osbpl6* | 1.37E-20 | 1.752867 |
| ENSMUSG00000086141 | *9030622O22Rik* | 6.75E-14 | 1.748517 |
| ENSMUSG00000087413 | *Gm11266* | 1.49E-10 | 1.7464 |
| ENSMUSG00000043088 | *Il17re* | 8.79E-06 | 1.745302 |

| | | | |
|---|---|---|---|
| ENSMUSG00000073877 | *Gm13306* | 1.50E-02 | 1.741028 |
| ENSMUSG00000058420 | *Syt17* | 5.21E-07 | 1.739965 |
| ENSMUSG00000054966 | *Lmntd1* | 3.10E-21 | 1.739876 |
| ENSMUSG00000044156 | *Hepacam2* | 8.78E-03 | 1.7394 |
| ENSMUSG00000112343 | *Sfta3-ps* | 6.02E-35 | 1.73883 |
| ENSMUSG00000025578 | *Cbx8* | 2.05E-02 | 1.738267 |
| ENSMUSG00000049420 | *Tmem200a* | 4.62E-16 | 1.737708 |
| ENSMUSG00000094786 | *Gm14403* | 2.31E-14 | 1.73598 |
| ENSMUSG00000052516 | *Robo2* | 3.01E-29 | 1.732225 |
| ENSMUSG00000052117 | *D630039A03Rik* | 2.49E-03 | 1.730316 |
| ENSMUSG00000095247 | *Ccl27a* | 1.67E-02 | 1.729601 |
| ENSMUSG00000035829 | *Ppp1r26* | 4.19E-05 | 1.727533 |
| ENSMUSG00000031489 | *Adrb3* | 1.72E-07 | 1.727349 |
| ENSMUSG00000022844 | *Pdia5* | 7.32E-17 | 1.727186 |
| ENSMUSG00000030732 | *Chrdl2* | 1.63E-02 | 1.727151 |
| ENSMUSG00000034685 | *Fam171a2* | 2.13E-04 | 1.726734 |
| ENSMUSG00000063535 | *Zfp773* | 5.88E-03 | 1.725168 |
| ENSMUSG00000030302 | *Atp2b2* | 1.69E-08 | 1.723946 |
| ENSMUSG00000024063 | *Lbh* | 4.98E-36 | 1.722771 |
| ENSMUSG00000068246 | *Apol9b* | 3.02E-02 | 1.721537 |
| ENSMUSG00000027849 | *Syt6* | 2.77E-06 | 1.721018 |
| ENSMUSG00000023011 | *Faim2* | 1.27E-05 | 1.718468 |
| ENSMUSG00000037709 | *Fam13a* | 1.35E-08 | 1.718152 |
| ENSMUSG00000037973 | *Ccdc129* | 1.14E-09 | 1.717358 |
| ENSMUSG00000030093 | *Wnt7a* | 3.68E-05 | 1.717134 |
| ENSMUSG00000054459 | *Vsnl1* | 3.49E-14 | 1.71657 |
| ENSMUSG00000106749 | *Gm43281* | 1.17E-06 | 1.71649 |
| ENSMUSG00000038453 | *Srcin1* | 3.73E-09 | 1.71294 |
| ENSMUSG00000020169 | *Best3* | 4.21E-07 | 1.7127 |
| ENSMUSG00000070802 | *Pnmal2* | 2.73E-03 | 1.712669 |
| ENSMUSG00000108767 | *Gm29763* | 4.23E-02 | 1.712211 |
| ENSMUSG00000106617 | *Gm36266* | 4.87E-02 | 1.708878 |
| ENSMUSG00000037846 | *Rtkn2* | 5.59E-09 | 1.707006 |
| ENSMUSG00000046916 | *Myct1* | 1.96E-11 | 1.7066 |
| ENSMUSG00000020734 | *Grin2c* | 1.11E-02 | 1.70189 |
| ENSMUSG00000020135 | *Apc2* | 5.57E-05 | 1.700572 |
| ENSMUSG00000029510 | *Gpc2* | 3.36E-02 | 1.699228 |
| ENSMUSG00000011658 | *Fuz* | 2.30E-07 | 1.696259 |
| ENSMUSG00000028871 | *Rspo1* | 5.07E-03 | 1.694801 |
| ENSMUSG00000059323 | *Tonsl* | 8.20E-04 | 1.693202 |
| ENSMUSG00000051727 | *Kctd14* | 3.92E-02 | 1.692769 |
| ENSMUSG00000021496 | *Pcbd2* | 5.81E-19 | 1.692202 |
| ENSMUSG00000024206 | *Rfx2* | 4.60E-17 | 1.691263 |
| ENSMUSG00000034771 | *Tle2* | 8.58E-09 | 1.690916 |
| ENSMUSG00000105637 | *Gm42480* | 4.36E-02 | 1.690621 |
| ENSMUSG00000110206 | *Flt3l* | 3.41E-08 | 1.689626 |
| ENSMUSG00000020566 | *Atp6v1c2* | 6.88E-19 | 1.689203 |
| ENSMUSG00000026347 | *Tmem163* | 2.61E-23 | 1.688979 |
| ENSMUSG00000038422 | *Hdhd3* | 1.93E-02 | 1.688234 |
| ENSMUSG00000110893 | *Gm47392* | 3.19E-02 | 1.684205 |
| ENSMUSG00000020546 | *Stxbp4* | 8.52E-19 | 1.682897 |
| ENSMUSG00000031147 | *Magix* | 1.99E-02 | 1.682316 |
| ENSMUSG00000097399 | *Gm26555* | 2.40E-02 | 1.681825 |

| | | | |
|---|---|---|---|
| ENSMUSG00000057719 | *Sh3rf2* | 5.96E-15 | 1.681035 |
| ENSMUSG00000078794 | *Dact3* | 3.49E-06 | 1.680713 |
| ENSMUSG00000026676 | *Ccdc3* | 9.32E-13 | 1.678664 |
| ENSMUSG00000113585 | *Gm19605* | 3.31E-04 | 1.676504 |
| ENSMUSG00000112714 | *Gm4798* | 2.16E-06 | 1.675832 |
| ENSMUSG00000082192 | *Gm14719* | 3.57E-02 | 1.675599 |
| ENSMUSG00000104244 | *2900018N21Rik* | 2.55E-02 | 1.67496 |
| ENSMUSG00000098022 | *Zfp82* | 2.20E-02 | 1.672994 |
| ENSMUSG00000040543 | *Pitpnm3* | 2.42E-15 | 1.670307 |
| ENSMUSG00000040009 | *Gnaz* | 2.72E-03 | 1.670102 |
| ENSMUSG00000074282 | *Zfp94* | 1.02E-04 | 1.669767 |
| ENSMUSG00000041431 | *Ccnb1* | 2.46E-02 | 1.668727 |
| ENSMUSG00000037101 | *Ttc29* | 1.48E-07 | 1.668254 |
| ENSMUSG00000044694 | *2010007H06Rik* | 4.57E-05 | 1.664324 |
| ENSMUSG00000074283 | *Zfp109* | 1.27E-04 | 1.664311 |
| ENSMUSG00000042750 | *Bex2* | 2.12E-02 | 1.660556 |
| ENSMUSG00000037610 | *Kcnmb2* | 3.48E-09 | 1.660213 |
| ENSMUSG00000029206 | *Nsun7* | 8.95E-03 | 1.65966 |
| ENSMUSG00000048096 | *Lmod1* | 4.51E-06 | 1.658969 |
| ENSMUSG00000087424 | *5730405O15Rik* | 1.41E-02 | 1.658718 |
| ENSMUSG00000043664 | *Tmem221* | 1.85E-03 | 1.657821 |
| ENSMUSG00000036962 | *Cfap221* | 1.50E-04 | 1.657207 |
| ENSMUSG00000104658 | *Gm30275* | 9.01E-04 | 1.656842 |
| ENSMUSG00000059588 | *Calcrl* | 5.46E-25 | 1.656551 |
| ENSMUSG00000032606 | *Nicn1* | 4.61E-05 | 1.655171 |
| ENSMUSG00000042988 | *Notum* | 3.14E-02 | 1.651852 |
| ENSMUSG00000090135 | *Gm15809* | 7.06E-05 | 1.651829 |
| ENSMUSG00000026785 | *Pkn3* | 3.37E-07 | 1.651277 |
| ENSMUSG00000036864 | *Proser3* | 1.85E-04 | 1.650066 |
| ENSMUSG00000021573 | *Tppp* | 1.72E-15 | 1.64999 |
| ENSMUSG00000047146 | *Tet1* | 9.98E-15 | 1.649532 |
| ENSMUSG00000022763 | *Aifm3* | 4.57E-02 | 1.648956 |
| ENSMUSG00000029074 | *Ttll10* | 4.54E-08 | 1.648169 |
| ENSMUSG00000106951 | *5930430L01Rik* | 2.79E-03 | 1.648167 |
| ENSMUSG00000066129 | *Kndc1* | 1.48E-05 | 1.648069 |
| ENSMUSG00000036218 | *Pdzrn4* | 1.20E-13 | 1.64797 |
| ENSMUSG00000049349 | *Gm5105* | 4.33E-02 | 1.647966 |
| ENSMUSG00000034522 | *Zfp395* | 2.88E-12 | 1.647198 |
| ENSMUSG00000004996 | *Mri1* | 6.19E-05 | 1.646009 |
| ENSMUSG00000030209 | *Grin2b* | 1.06E-03 | 1.644894 |
| ENSMUSG00000083640 | *Gm8876* | 2.25E-02 | 1.644745 |
| ENSMUSG00000022199 | *Slc22a17* | 2.43E-06 | 1.644577 |
| ENSMUSG00000097115 | *1810019N24Rik* | 4.68E-02 | 1.644237 |
| ENSMUSG00000053730 | *Tmem39b* | 5.53E-05 | 1.642946 |
| ENSMUSG00000022615 | *Tymp* | 9.68E-04 | 1.641539 |
| ENSMUSG00000038605 | *Samd10* | 3.89E-05 | 1.640188 |
| ENSMUSG00000028801 | *Stpg1* | 5.81E-04 | 1.639582 |
| ENSMUSG00000024579 | *Pcyox1l* | 2.73E-06 | 1.639339 |
| ENSMUSG00000109394 | *A230057D06Rik* | 1.24E-15 | 1.639166 |
| ENSMUSG00000038576 | *Susd4* | 5.59E-13 | 1.639016 |
| ENSMUSG00000097453 | *Gm26894* | 1.64E-02 | 1.637894 |
| ENSMUSG00000027186 | *Elf5* | 3.99E-04 | 1.637037 |
| ENSMUSG00000067629 | *Syngap1* | 1.04E-10 | 1.636623 |

| | | | |
|---|---|---|---|
| ENSMUSG00000042474 | *Fcmr* | 1.58E-04 | 1.636419 |
| ENSMUSG00000049625 | *Tifab* | 3.19E-09 | 1.636014 |
| ENSMUSG00000102437 | *Gm38048* | 9.87E-06 | 1.635702 |
| ENSMUSG00000032357 | *Tinag* | 5.76E-05 | 1.635216 |
| ENSMUSG00000033952 | *Aspm* | 3.64E-06 | 1.635025 |
| ENSMUSG00000106993 | *Gm43417* | 1.87E-02 | 1.634425 |
| ENSMUSG00000085682 | *Gm14267* | 1.56E-03 | 1.633309 |
| ENSMUSG00000050705 | *2310061I04Rik* | 1.46E-05 | 1.632802 |
| ENSMUSG00000035504 | *Reep6* | 1.13E-03 | 1.631873 |
| ENSMUSG00000029521 | *Chek2* | 1.57E-03 | 1.62588 |
| ENSMUSG00000085237 | *Gm15406* | 4.19E-08 | 1.6257 |
| ENSMUSG00000031220 | *Awat2* | 1.51E-02 | 1.624394 |
| ENSMUSG00000054555 | *Adam12* | 2.00E-23 | 1.623831 |
| ENSMUSG00000112694 | *Gm33037* | 1.78E-02 | 1.620917 |
| ENSMUSG00000042826 | *Fgf11* | 1.79E-03 | 1.618808 |
| ENSMUSG00000040282 | *BC052040* | 2.59E-16 | 1.618536 |
| ENSMUSG00000032064 | *Dixdc1* | 5.04E-09 | 1.617751 |
| ENSMUSG00000111229 | *Gm39323* | 1.15E-03 | 1.617613 |
| ENSMUSG00000052921 | *Arhgef15* | 1.73E-18 | 1.61749 |
| ENSMUSG00000031870 | *Pgr* | 3.58E-10 | 1.617384 |
| ENSMUSG00000051811 | *Cox6b2* | 1.16E-02 | 1.617084 |
| ENSMUSG00000035435 | *Abca17* | 2.04E-08 | 1.615687 |
| ENSMUSG00000022179 | *4931414P19Rik* | 3.70E-05 | 1.615439 |
| ENSMUSG00000028031 | *Dkk2* | 2.04E-05 | 1.613944 |
| ENSMUSG00000049811 | *Fam161a* | 3.97E-06 | 1.611627 |
| ENSMUSG00000097805 | *Gm17473* | 1.63E-07 | 1.610565 |
| ENSMUSG00000027560 | *Dok5* | 1.58E-03 | 1.609425 |
| ENSMUSG00000045875 | *Adra1a* | 3.66E-05 | 1.608373 |
| ENSMUSG00000074793 | *Hspa12b* | 3.94E-19 | 1.607954 |
| ENSMUSG00000068551 | *Zfp467* | 5.27E-10 | 1.605012 |
| ENSMUSG00000022442 | *Ttll1* | 1.44E-05 | 1.604733 |
| ENSMUSG00000058396 | *Gpr182* | 6.53E-09 | 1.603606 |
| ENSMUSG00000001496 | *Nkx2-1* | 1.73E-11 | 1.599734 |
| ENSMUSG00000116116 | *Gm671* | 1.39E-07 | 1.599663 |
| ENSMUSG00000030796 | *Tead2* | 3.21E-11 | 1.599608 |
| ENSMUSG00000057060 | *Slc35f3* | 1.38E-03 | 1.599499 |
| ENSMUSG00000046806 | *Cyren* | 2.91E-05 | 1.5994 |
| ENSMUSG00000003779 | *Kif20a* | 2.39E-02 | 1.599212 |
| ENSMUSG00000066235 | *Pomgnt2* | 2.51E-04 | 1.597016 |
| ENSMUSG00000055333 | *Fat2* | 2.60E-04 | 1.596568 |
| ENSMUSG00000030873 | *Scnn1b* | 2.04E-20 | 1.594799 |
| ENSMUSG00000097915 | *A330009N23Rik* | 3.03E-03 | 1.594615 |
| ENSMUSG00000020614 | *Fam20a* | 1.16E-14 | 1.594492 |
| ENSMUSG00000062542 | *Syt9* | 2.52E-02 | 1.59351 |
| ENSMUSG00000068263 | *Efcc1* | 1.34E-09 | 1.593299 |
| ENSMUSG00000097124 | *A530020G20Rik* | 5.09E-04 | 1.593002 |
| ENSMUSG00000078607 | *1810010H24Rik* | 7.09E-07 | 1.592582 |
| ENSMUSG00000043668 | *Tox3* | 5.03E-16 | 1.591389 |
| ENSMUSG00000022676 | *Snai2* | 1.80E-03 | 1.591354 |
| ENSMUSG00000042210 | *Abhd14a* | 1.25E-02 | 1.591316 |
| ENSMUSG00000024403 | *Atp6v1g2* | 5.74E-03 | 1.591076 |
| ENSMUSG00000026860 | *Sh3glb2* | 5.11E-12 | 1.590949 |
| ENSMUSG00000027217 | *Tspan18* | 4.38E-21 | 1.590887 |

| | | | |
|---|---|---|---|
| ENSMUSG00000082292 | *Gm12250* | 3.96E-04 | 1.59078 |
| ENSMUSG00000105135 | *Gm43667* | 1.64E-02 | 1.589917 |
| ENSMUSG00000031665 | *Sall1* | 9.47E-03 | 1.589292 |
| ENSMUSG00000108207 | *1810059H22Rik* | 2.37E-02 | 1.589229 |
| ENSMUSG00000087013 | *2610027K06Rik* | 5.70E-14 | 1.589185 |
| ENSMUSG00000044548 | *Dact1* | 2.47E-07 | 1.587675 |
| ENSMUSG00000110279 | *Gm45552* | 1.29E-02 | 1.585905 |
| ENSMUSG00000115301 | *Gm6330* | 2.91E-02 | 1.585784 |
| ENSMUSG00000055150 | *Zfp78* | 2.68E-03 | 1.584768 |
| ENSMUSG00000057751 | *Megf6* | 1.53E-14 | 1.584688 |
| ENSMUSG00000054517 | *Trim65* | 3.34E-06 | 1.584269 |
| ENSMUSG00000002771 | *Grin2d* | 3.01E-05 | 1.583762 |
| ENSMUSG00000107736 | *Gm44148* | 4.72E-09 | 1.583468 |
| ENSMUSG00000030680 | *Pagr1a* | 7.90E-04 | 1.5832 |
| ENSMUSG00000037025 | *Foxa2* | 1.28E-03 | 1.582839 |
| ENSMUSG00000103766 | *Gm38392* | 2.13E-04 | 1.580705 |
| ENSMUSG00000033233 | *Trim45* | 1.95E-02 | 1.578751 |
| ENSMUSG00000046240 | *Hepacam* | 4.39E-04 | 1.578062 |
| ENSMUSG00000098465 | *Mir1668* | 3.82E-05 | 1.577691 |
| ENSMUSG00000089769 | *Gm16574* | 3.22E-02 | 1.577572 |
| ENSMUSG00000029790 | *Cep41* | 1.21E-04 | 1.576274 |
| ENSMUSG00000046470 | *Sox18* | 1.24E-09 | 1.57479 |
| ENSMUSG00000036598 | *Ccdc113* | 2.12E-02 | 1.574563 |
| ENSMUSG00000084939 | *Gm830* | 2.06E-02 | 1.573865 |
| ENSMUSG00000069378 | *Prdm6* | 2.91E-16 | 1.572953 |
| ENSMUSG00000042766 | *Trim46* | 2.68E-03 | 1.572774 |
| ENSMUSG00000034848 | *Ttc21b* | 1.51E-11 | 1.572038 |
| ENSMUSG00000064284 | *Cdpf1* | 2.94E-03 | 1.571986 |
| ENSMUSG00000115095 | *Gm49311* | 8.92E-03 | 1.571257 |
| ENSMUSG00000020182 | *Ddc* | 2.97E-02 | 1.571232 |
| ENSMUSG00000032492 | *Pth1r* | 1.36E-04 | 1.570378 |
| ENSMUSG00000027574 | *Nkain4* | 3.42E-06 | 1.570249 |
| ENSMUSG00000021684 | *Pde8b* | 6.66E-17 | 1.570207 |
| ENSMUSG00000039552 | *Rsph4a* | 1.53E-02 | 1.570094 |
| ENSMUSG00000102305 | *Gm38192* | 4.40E-02 | 1.56988 |
| ENSMUSG00000041117 | *Ccdc8* | 7.59E-06 | 1.569825 |
| ENSMUSG00000033590 | *Myo5c* | 7.37E-13 | 1.569346 |
| ENSMUSG00000043932 | *Klri2* | 2.44E-03 | 1.5688 |
| ENSMUSG00000108218 | *Olfr1372-ps1* | 8.01E-05 | 1.567865 |
| ENSMUSG00000055134 | *9130017K11Rik* | 1.04E-02 | 1.567643 |
| ENSMUSG00000038725 | *Pkhd1l1* | 2.18E-08 | 1.56719 |
| ENSMUSG00000037279 | *Ovol2* | 3.58E-02 | 1.566143 |
| ENSMUSG00000050947 | *Amigo1* | 1.16E-07 | 1.566084 |
| ENSMUSG00000022416 | *Cacna1i* | 1.45E-03 | 1.565741 |
| ENSMUSG00000038379 | *Ttk* | 3.20E-02 | 1.564059 |
| ENSMUSG00000018395 | *Kif3a* | 5.66E-10 | 1.563864 |
| ENSMUSG00000078772 | *Gm12353* | 1.41E-08 | 1.563774 |
| ENSMUSG00000115157 | *Gm49397* | 8.53E-23 | 1.563261 |
| ENSMUSG00000079164 | *Tlr5* | 3.12E-06 | 1.563178 |
| ENSMUSG00000068323 | *Slc4a5* | 5.95E-07 | 1.56313 |
| ENSMUSG00000054967 | *Zfp647* | 3.16E-03 | 1.563113 |
| ENSMUSG00000086533 | *Mypopos* | 8.00E-09 | 1.560072 |
| ENSMUSG00000064294 | *Aox3* | 2.44E-09 | 1.560014 |

| | | | |
|---|---|---|---|
| ENSMUSG00000025933 | *Tmem14a* | 2.01E-02 | 1.559242 |
| ENSMUSG00000061397 | *Krt79* | 1.26E-02 | 1.559113 |
| ENSMUSG00000062861 | *Zfp28* | 1.43E-05 | 1.558974 |
| ENSMUSG00000114553 | *Gm7644* | 7.76E-03 | 1.558629 |
| ENSMUSG00000039577 | *Nphp4* | 5.79E-05 | 1.558615 |
| ENSMUSG00000085936 | *2610307P16Rik* | 1.35E-26 | 1.558192 |
| ENSMUSG00000041429 | *Nthl1* | 1.26E-02 | 1.55773 |
| ENSMUSG00000030351 | *Tspan11* | 1.91E-24 | 1.555549 |
| ENSMUSG00000017167 | *Cntnap1* | 5.58E-04 | 1.555233 |
| ENSMUSG00000030800 | *Prss8* | 4.00E-07 | 1.554977 |
| ENSMUSG00000026824 | *Kcnj3* | 1.18E-10 | 1.55393 |
| ENSMUSG00000022759 | *Lrrc74b* | 1.13E-02 | 1.553927 |
| ENSMUSG00000102691 | *Gm37780* | 9.59E-03 | 1.553868 |
| ENSMUSG00000027424 | *Mgme1* | 8.77E-03 | 1.553739 |
| ENSMUSG00000038797 | *Zscan2* | 3.13E-04 | 1.553465 |
| ENSMUSG00000112121 | *C230072F16Rik* | 1.48E-06 | 1.553102 |
| ENSMUSG00000039304 | *Tnfsf10* | 1.86E-18 | 1.552207 |
| ENSMUSG00000054074 | *Skida1* | 4.97E-02 | 1.551638 |
| ENSMUSG00000074646 | *6430550D23Rik* | 3.75E-02 | 1.550595 |
| ENSMUSG00000051550 | *Zfp579* | 4.82E-05 | 1.549596 |
| ENSMUSG00000029053 | *Prkcz* | 2.72E-12 | 1.549462 |
| ENSMUSG00000045410 | *Akr1e1* | 1.81E-02 | 1.547976 |
| ENSMUSG00000108365 | *Gm44951* | 3.56E-03 | 1.547705 |
| ENSMUSG00000083817 | *Gm14400* | 4.50E-02 | 1.547364 |
| ENSMUSG00000024299 | *Adamts10* | 6.54E-19 | 1.546832 |
| ENSMUSG00000029603 | *Dtx1* | 4.99E-03 | 1.546727 |
| ENSMUSG00000075592 | *Nynrin* | 8.46E-14 | 1.546558 |
| ENSMUSG00000031517 | *Gpm6a* | 1.10E-11 | 1.546528 |
| ENSMUSG00000099564 | *Gm28729* | 2.89E-02 | 1.546472 |
| ENSMUSG00000022701 | *Ccdc191* | 1.34E-10 | 1.546372 |
| ENSMUSG00000039058 | *Ak5* | 3.48E-05 | 1.546267 |
| ENSMUSG00000048498 | *Cd300e* | 4.94E-02 | 1.545969 |
| ENSMUSG00000106633 | *Gm43431* | 4.81E-02 | 1.544944 |
| ENSMUSG00000066438 | *Plekhd1* | 9.20E-06 | 1.544901 |
| ENSMUSG00000060548 | *Tnfrsf19* | 5.99E-12 | 1.544576 |
| ENSMUSG00000020682 | *Mmp28* | 3.42E-09 | 1.544554 |
| ENSMUSG00000042073 | *Abhd14b* | 8.46E-08 | 1.543843 |
| ENSMUSG00000074384 | *AI429214* | 3.33E-02 | 1.543505 |
| ENSMUSG00000082264 | *Gm12799* | 1.72E-04 | 1.543452 |
| ENSMUSG00000048897 | *Zfp710* | 6.90E-19 | 1.542688 |
| ENSMUSG00000109165 | *Gm45148* | 3.20E-02 | 1.542583 |
| ENSMUSG00000022543 | *4930451G09Rik* | 2.13E-02 | 1.542361 |
| ENSMUSG00000032609 | *Klhdc8b* | 1.78E-05 | 1.541974 |
| ENSMUSG00000062785 | *Kcnc3* | 7.41E-09 | 1.541955 |
| ENSMUSG00000034532 | *Fbxo16* | 4.66E-02 | 1.5419 |
| ENSMUSG00000020513 | *Tubd1* | 2.22E-06 | 1.541575 |
| ENSMUSG00000035547 | *Capn5* | 1.06E-11 | 1.541062 |
| ENSMUSG00000020268 | *Lyrm7* | 1.22E-03 | 1.541004 |
| ENSMUSG00000106758 | *Gm42731* | 3.30E-02 | 1.540776 |
| ENSMUSG00000107436 | *Gm44416* | 2.31E-02 | 1.540763 |
| ENSMUSG00000041460 | *Cacna2d4* | 2.80E-02 | 1.540666 |
| ENSMUSG00000037813 | *D630003M21Rik* | 1.59E-03 | 1.540376 |
| ENSMUSG00000037892 | *Pcdh18* | 2.52E-14 | 1.540206 |

| | | | |
|---|---|---|---|
| ENSMUSG00000078552 | *Dcdc2b* | 3.72E-02 | 1.539768 |
| ENSMUSG00000113543 | *Gm36264* | 2.91E-02 | 1.539705 |
| ENSMUSG00000029335 | *Bmp3* | 1.77E-12 | 1.539659 |
| ENSMUSG00000107068 | *Gm42742* | 4.89E-06 | 1.538758 |
| ENSMUSG00000095987 | *Gm13104* | 1.34E-02 | 1.538362 |
| ENSMUSG00000046456 | *Tmem150b* | 2.91E-03 | 1.538263 |
| ENSMUSG00000106247 | *Gm43720* | 4.73E-02 | 1.538062 |
| ENSMUSG00000070873 | *Lilra5* | 3.76E-03 | 1.53778 |
| ENSMUSG00000022818 | *Cyp2ab1* | 2.94E-02 | 1.537532 |
| ENSMUSG00000058656 | *Samd12* | 1.10E-19 | 1.536867 |
| ENSMUSG00000059851 | *Kmt5c* | 1.85E-08 | 1.536592 |
| ENSMUSG00000104114 | *Gm37297* | 4.16E-04 | 1.536347 |
| ENSMUSG00000018740 | *Slc25a35* | 2.79E-05 | 1.536163 |
| ENSMUSG00000103845 | *Gm19026* | 3.48E-03 | 1.536156 |
| ENSMUSG00000112627 | *4933412E12Rik* | 2.50E-08 | 1.535554 |
| ENSMUSG00000035863 | *Palm* | 2.73E-10 | 1.535105 |
| ENSMUSG00000033763 | *Mtss1l* | 3.13E-14 | 1.535088 |
| ENSMUSG00000024787 | *Snx15* | 1.16E-06 | 1.534706 |
| ENSMUSG00000028088 | *Fmo5* | 1.52E-06 | 1.534518 |
| ENSMUSG00000106892 | *Gm42791* | 4.56E-09 | 1.534363 |
| ENSMUSG00000051359 | *Ncald* | 1.38E-26 | 1.534218 |
| ENSMUSG00000049532 | *Sall2* | 6.86E-05 | 1.534187 |
| ENSMUSG00000112164 | *Gm19990* | 4.48E-02 | 1.533334 |
| ENSMUSG00000026888 | *Grb14* | 1.31E-24 | 1.5329 |
| ENSMUSG00000026586 | *Prrx1* | 1.15E-18 | 1.532838 |
| ENSMUSG00000023147 | *Wrb* | 8.70E-05 | 1.531827 |
| ENSMUSG00000022941 | *Ripply3* | 1.87E-10 | 1.531256 |
| ENSMUSG00000021055 | *Esr2* | 9.86E-08 | 1.531097 |
| ENSMUSG00000050069 | *Grem2* | 2.17E-14 | 1.53098 |
| ENSMUSG00000042784 | *Muc1* | 1.74E-15 | 1.530839 |
| ENSMUSG00000030361 | *Klrb1a* | 1.98E-02 | 1.530589 |
| ENSMUSG00000036278 | *Macrod1* | 1.31E-10 | 1.530463 |
| ENSMUSG00000027500 | *Stmn2* | 5.55E-12 | 1.529312 |
| ENSMUSG00000079671 | *2610203C22Rik* | 1.17E-16 | 1.529211 |
| ENSMUSG00000070737 | *Tmem35b* | 1.01E-03 | 1.528272 |
| ENSMUSG00000026443 | *Lrrn2* | 3.23E-03 | 1.52766 |
| ENSMUSG00000036882 | *Arhgap33* | 2.72E-04 | 1.527118 |
| ENSMUSG00000076431 | *Sox4* | 1.22E-10 | 1.526384 |
| ENSMUSG00000049107 | *Ntf3* | 3.97E-10 | 1.526123 |
| ENSMUSG00000097354 | *2310001H17Rik* | 1.98E-27 | 1.525662 |
| ENSMUSG00000102326 | *Gm37788* | 2.52E-02 | 1.525393 |
| ENSMUSG00000115970 | *8430426J06Rik* | 1.78E-06 | 1.52498 |
| ENSMUSG00000000627 | *Sema4f* | 1.29E-05 | 1.524743 |
| ENSMUSG00000051855 | *Mest* | 2.70E-08 | 1.524224 |
| ENSMUSG00000019312 | *Grb7* | 4.43E-03 | 1.524165 |
| ENSMUSG00000041361 | *Myzap* | 3.12E-18 | 1.523261 |
| ENSMUSG00000036983 | *Tfb1m* | 1.77E-06 | 1.521731 |
| ENSMUSG00000024673 | *Ms4a1* | 3.20E-04 | 1.521639 |
| ENSMUSG00000019945 | *Cabcoco1* | 2.38E-04 | 1.52083 |
| ENSMUSG00000026748 | *Plxdc2* | 3.63E-19 | 1.520549 |
| ENSMUSG00000107499 | *Ccdc142* | 4.86E-02 | 1.520041 |
| ENSMUSG00000029456 | *Acad10* | 4.39E-07 | 1.520036 |
| ENSMUSG00000023949 | *Tcte1* | 3.76E-02 | 1.5199 |

| | | | |
|---|---|---|---|
| ENSMUSG00000036555 | *Iqce* | 2.95E-11 | 1.518721 |
| ENSMUSG00000060862 | *Zbtb40* | 1.39E-13 | 1.517839 |
| ENSMUSG00000027490 | *E2f1* | 3.92E-02 | 1.517055 |
| ENSMUSG00000032680 | *6820408C15Rik* | 1.89E-02 | 1.516845 |
| ENSMUSG00000022103 | *Gfra2* | 1.63E-10 | 1.516841 |
| ENSMUSG00000000402 | *Egfl6* | 8.17E-08 | 1.516514 |
| ENSMUSG00000026390 | *Marco* | 1.23E-08 | 1.516135 |
| ENSMUSG00000107794 | *Gm44095* | 3.87E-02 | 1.515819 |
| ENSMUSG00000111761 | *9230112J17Rik* | 1.69E-02 | 1.51564 |
| ENSMUSG00000056228 | *Cars2* | 3.79E-12 | 1.514966 |
| ENSMUSG00000079003 | *Samd1* | 1.24E-04 | 1.514814 |
| ENSMUSG00000038302 | *Afg1l* | 7.32E-25 | 1.514245 |
| ENSMUSG00000022309 | *Angpt1* | 1.03E-07 | 1.514084 |
| ENSMUSG00000045114 | *Prrt2* | 9.94E-04 | 1.51398 |
| ENSMUSG00000037922 | *Bank1* | 1.10E-03 | 1.512972 |
| ENSMUSG00000034800 | *Zfp661* | 9.55E-03 | 1.512619 |
| ENSMUSG00000036411 | *9530077C05Rik* | 8.03E-05 | 1.512568 |
| ENSMUSG00000035459 | *Stab2* | 1.02E-04 | 1.512457 |
| ENSMUSG00000062960 | *Kdr* | 3.63E-21 | 1.511554 |
| ENSMUSG00000050936 | *Gm42743* | 1.63E-07 | 1.511039 |
| ENSMUSG00000021719 | *Rgs7bp* | 4.17E-07 | 1.510943 |
| ENSMUSG00000031802 | *Phxr4* | 1.18E-04 | 1.510821 |
| ENSMUSG00000085207 | *Gm11767* | 2.22E-03 | 1.50959 |
| ENSMUSG00000057396 | *Zfp759* | 1.29E-03 | 1.509269 |
| ENSMUSG00000031736 | *Crnde* | 1.62E-06 | 1.509225 |
| ENSMUSG00000027977 | *Ndst3* | 7.76E-05 | 1.508506 |
| ENSMUSG00000025395 | *Prim1* | 4.62E-03 | 1.507443 |
| ENSMUSG00000030551 | *Nr2f2* | 3.20E-10 | 1.507264 |
| ENSMUSG00000028003 | *Lrat* | 1.08E-05 | 1.507135 |
| ENSMUSG00000100707 | *Gm28523* | 2.77E-03 | 1.50701 |
| ENSMUSG00000050222 | *Il17d* | 1.67E-02 | 1.506503 |
| ENSMUSG00000056185 | *Snx32* | 3.36E-07 | 1.506257 |
| ENSMUSG00000098318 | *Lockd* | 1.28E-03 | 1.50611 |
| ENSMUSG00000031775 | *Pllp* | 1.05E-06 | 1.506048 |
| ENSMUSG00000002346 | *Slc25a42* | 8.80E-09 | 1.506014 |
| ENSMUSG00000007207 | *Stx1a* | 7.57E-06 | 1.5059 |
| ENSMUSG00000030084 | *Plxna1* | 1.91E-16 | 1.505539 |
| ENSMUSG00000059146 | *Ntrk3* | 4.22E-15 | 1.504398 |
| ENSMUSG00000073424 | *Cyp4f15* | 1.80E-02 | 1.503825 |
| ENSMUSG00000039062 | *Anpep* | 4.19E-13 | 1.503782 |
| ENSMUSG00000001288 | *Rarg* | 2.33E-13 | 1.503681 |
| ENSMUSG00000051817 | *Sox12* | 4.13E-02 | 1.502765 |
| ENSMUSG00000076609 | *Igkc* | 4.41E-05 | 1.502726 |
| ENSMUSG00000013033 | *Adgrl1* | 1.28E-14 | 1.502317 |
| ENSMUSG00000033948 | *Zswim5* | 6.73E-05 | 1.501944 |
| ENSMUSG00000079620 | *Muc4* | 1.41E-14 | 1.501612 |
| ENSMUSG00000024018 | *Ccdc167* | 3.57E-09 | 1.501095 |
| ENSMUSG00000045441 | *Gprin3* | 3.83E-11 | 1.500851 |
| ENSMUSG00000036959 | *Bcorl1* | 3.46E-14 | 1.500655 |
| ENSMUSG00000044701 | *Il27* | 1.05E-02 | 1.500073 |
| ENSMUSG00000028773 | *Fabp3* | 3.89E-05 | 1.501682 |
| ENSMUSG00000070031 | *Sp140* | 3.17E-21 | 1.501904 |
| ENSMUSG00000105053 | *Gm43064* | 1.55E-08 | 1.501933 |

| | | | |
|---|---|---|---|
| ENSMUSG00000029552 | *Tes* | 1.46E-22 | 1.502499 |
| ENSMUSG00000047976 | *Kcna1* | 4.87E-04 | 1.502872 |
| ENSMUSG00000045349 | *Sh2d5* | 1.76E-02 | 1.502987 |
| ENSMUSG00000058755 | *Osm* | 1.79E-04 | 1.503113 |
| ENSMUSG00000104348 | *Gm37691* | 1.01E-03 | 1.503512 |
| ENSMUSG00000113041 | *Gm48302* | 4.83E-03 | 1.503595 |
| ENSMUSG00000100969 | *1700030N03Rik* | 9.26E-03 | 1.503827 |
| ENSMUSG00000030595 | *Nfkbib* | 5.68E-12 | 1.503921 |
| ENSMUSG00000090141 | *Gm614* | 3.44E-04 | 1.504654 |
| ENSMUSG00000035208 | *Slfn8* | 3.12E-11 | 1.504893 |
| ENSMUSG00000004951 | *Hspb1* | 3.20E-15 | 1.505858 |
| ENSMUSG00000059498 | *Fcgr3* | 5.02E-25 | 1.506788 |
| ENSMUSG00000043953 | *Ccrl2* | 2.05E-15 | 1.507145 |
| ENSMUSG00000041754 | *Trem3* | 1.95E-08 | 1.507303 |
| ENSMUSG00000112707 | *D830005E20Rik* | 3.83E-06 | 1.507644 |
| ENSMUSG00000084208 | *Gm12792* | 3.21E-02 | 1.508358 |
| ENSMUSG00000053560 | *Ier2* | 6.51E-12 | 1.508916 |
| ENSMUSG00000021886 | *Gpr65* | 5.04E-09 | 1.509174 |
| ENSMUSG00000105337 | *1700094M23Rik* | 2.68E-06 | 1.510638 |
| ENSMUSG00000075042 | *4930431P03Rik* | 3.69E-02 | 1.511896 |
| ENSMUSG00000101969 | *Gm20125* | 2.94E-03 | 1.51201 |
| ENSMUSG00000039005 | *Tlr4* | 9.61E-33 | 1.512543 |
| ENSMUSG00000102476 | *Gm37245* | 3.69E-08 | 1.512732 |
| ENSMUSG00000036557 | *Stpg4* | 4.97E-02 | 1.513017 |
| ENSMUSG00000108368 | *Gm45053* | 8.49E-05 | 1.513592 |
| ENSMUSG00000034640 | *Tiparp* | 3.82E-23 | 1.514228 |
| ENSMUSG00000004609 | *Cd33* | 1.53E-22 | 1.514366 |
| ENSMUSG00000097558 | *Gm26902* | 2.40E-02 | 1.514485 |
| ENSMUSG00000030069 | *Prok2* | 3.12E-09 | 1.51466 |
| ENSMUSG00000079012 | *Serpina3m* | 9.97E-15 | 1.515028 |
| ENSMUSG00000037833 | *Sh2d4b* | 1.28E-10 | 1.515452 |
| ENSMUSG00000036181 | *Hist1h1c* | 8.16E-07 | 1.515838 |
| ENSMUSG00000032942 | *Ucp3* | 2.72E-04 | 1.515872 |
| ENSMUSG00000078915 | *Hsp25-ps1* | 1.89E-14 | 1.515918 |
| ENSMUSG00000045763 | *Basp1* | 8.22E-23 | 1.516202 |
| ENSMUSG00000023964 | *Calcr* | 2.13E-09 | 1.51776 |
| ENSMUSG00000074449 | *Gm15319* | 1.18E-02 | 1.518175 |
| ENSMUSG00000027748 | *Trpc4* | 4.48E-02 | 1.518209 |
| ENSMUSG00000084093 | *Gm16418* | 4.51E-02 | 1.518297 |
| ENSMUSG00000025716 | *Myo3a* | 1.15E-02 | 1.518517 |
| ENSMUSG00000019787 | *Trdn* | 5.00E-18 | 1.51858 |
| ENSMUSG00000089281 | *Scarna6* | 1.66E-02 | 1.519358 |
| ENSMUSG00000090248 | *Gm14027* | 1.67E-02 | 1.519727 |
| ENSMUSG00000086320 | *Gm12840* | 2.45E-10 | 1.519868 |
| ENSMUSG00000028573 | *Fggy* | 5.40E-03 | 1.520122 |
| ENSMUSG00000033730 | *Egr3* | 6.03E-09 | 1.520144 |
| ENSMUSG00000089716 | *Gm6264* | 6.83E-06 | 1.520708 |
| ENSMUSG00000051650 | *B3gnt2* | 7.54E-28 | 1.520857 |
| ENSMUSG00000104108 | *Gm37876* | 2.97E-03 | 1.520932 |
| ENSMUSG00000051439 | *Cd14* | 2.28E-24 | 1.521574 |
| ENSMUSG00000020732 | *Rab37* | 4.19E-04 | 1.521864 |
| ENSMUSG00000037904 | *Ankrd9* | 7.83E-03 | 1.522534 |
| ENSMUSG00000105195 | *Gm43584* | 5.16E-04 | 1.522603 |

| | | | | |
|---|---|---|---|---|
| ENSMUSG00000027360 | *Hdc* | 4.67E-21 | | 1.522604 |
| ENSMUSG00000001014 | *Icam4* | 1.36E-02 | | 1.522651 |
| ENSMUSG00000022126 | *Acod1* | 4.82E-25 | | 1.523696 |
| ENSMUSG00000031448 | *Adprhl1* | 1.56E-05 | | 1.524028 |
| ENSMUSG00000056328 | *Myh1* | 4.72E-14 | | 1.524211 |
| ENSMUSG00000087593 | *Gm16174* | 6.08E-03 | | 1.524313 |
| ENSMUSG00000020275 | *Rel* | 1.32E-22 | | 1.524463 |
| ENSMUSG00000016559 | *H3f3b* | 1.21E-21 | | 1.524786 |
| ENSMUSG00000116614 | *AC151299.1* | 5.83E-04 | | 1.526535 |
| ENSMUSG00000099250 | *Rn7s2* | 1.32E-02 | | 1.526701 |
| ENSMUSG00000099021 | *Rn7s1* | 1.32E-02 | | 1.526715 |
| ENSMUSG00000023067 | *Cdkn1a* | 1.39E-25 | | 1.527143 |
| ENSMUSG00000095294 | *Gm21119* | 1.32E-02 | | 1.527698 |
| ENSMUSG00000026073 | *Il1r2* | 1.36E-06 | | 1.527927 |
| ENSMUSG00000031231 | *Cox7b* | 1.64E-09 | | 1.528049 |
| ENSMUSG00000025161 | *Slc16a3* | 3.55E-18 | | 1.528109 |
| ENSMUSG00000097543 | *Gm805* | 4.90E-02 | | 1.528998 |
| ENSMUSG00000059974 | *Ntm* | 4.19E-21 | | 1.529013 |
| ENSMUSG00000103288 | *Gm37273* | 8.69E-03 | | 1.529304 |
| ENSMUSG00000052769 | *Gm9889* | 3.93E-02 | | 1.529381 |
| ENSMUSG00000024590 | *Lmnb1* | 8.45E-17 | | 1.529562 |
| ENSMUSG00000114277 | *Gm48583* | 4.03E-03 | | 1.529671 |
| ENSMUSG00000106497 | *Gm43195* | 3.51E-02 | | 1.529948 |
| ENSMUSG00000085147 | *Gm12609* | 2.37E-02 | | 1.529985 |
| ENSMUSG00000068697 | *Myoz1* | 6.25E-04 | | 1.530719 |
| ENSMUSG00000029660 | *Tex26* | 1.65E-04 | | 1.530841 |
| ENSMUSG00000046480 | *Scn4b* | 9.28E-05 | | 1.530902 |
| ENSMUSG00000115946 | *Mirt2* | 7.87E-13 | | 1.531478 |
| ENSMUSG00000021622 | *Ckmt2* | 1.46E-08 | | 1.531721 |
| ENSMUSG00000039753 | *Fbxl5* | 8.84E-17 | | 1.532351 |
| ENSMUSG00000108486 | *Gm44836* | 8.76E-04 | | 1.533293 |
| ENSMUSG00000082057 | *Gm15789* | 2.89E-02 | | 1.533491 |
| ENSMUSG00000028862 | *Map3k6* | 6.29E-22 | | 1.534354 |
| ENSMUSG00000061462 | *Obscn* | 6.82E-21 | | 1.535088 |
| ENSMUSG00000029204 | *Rhoh* | 1.11E-19 | | 1.535236 |
| ENSMUSG00000072620 | *Slfn2* | 1.20E-17 | | 1.535662 |
| ENSMUSG00000098670 | *Gm27818* | 3.10E-02 | | 1.536024 |
| ENSMUSG00000107029 | *Gm43123* | 2.37E-04 | | 1.536945 |
| ENSMUSG00000087066 | *Gm15518* | 2.95E-07 | | 1.537162 |
| ENSMUSG00000031682 | *1700011L22Rik* | 6.64E-04 | | 1.537235 |
| ENSMUSG00000078763 | *Slfn1* | 2.40E-16 | | 1.537518 |
| ENSMUSG00000074149 | *Gm10634* | 4.40E-06 | | 1.540107 |
| ENSMUSG00000030669 | *Calca* | 1.35E-04 | | 1.540221 |
| ENSMUSG00000030088 | *Aldh1l1* | 8.07E-17 | | 1.541343 |
| ENSMUSG00000030107 | *Usp18* | 3.01E-16 | | 1.541655 |
| ENSMUSG00000071342 | *Lsmem1* | 3.17E-06 | | 1.541988 |
| ENSMUSG00000027861 | *Casq2* | 2.44E-09 | | 1.542362 |
| ENSMUSG00000074419 | *Gm15448* | 1.08E-08 | | 1.542588 |
| ENSMUSG00000062077 | *Trim54* | 3.72E-07 | | 1.543231 |
| ENSMUSG00000031289 | *Il13ra2* | 2.50E-02 | | 1.544205 |
| ENSMUSG00000094344 | *Gm11942* | 5.03E-07 | | 1.545029 |
| ENSMUSG00000055602 | *Tcp10b* | 1.12E-04 | | 1.545251 |
| ENSMUSG00000022902 | *Stfa2* | 2.23E-06 | | 1.546076 |

| | | | |
|---|---|---|---|
| ENSMUSG00000097579 | *Gm26799* | 2.14E-03 | 1.546157 |
| ENSMUSG00000028116 | *Myoz2* | 3.48E-11 | 1.546592 |
| ENSMUSG00000029915 | *Clec5a* | 1.27E-14 | 1.546688 |
| ENSMUSG00000087135 | *Gm16096* | 3.13E-02 | 1.547248 |
| ENSMUSG00000026669 | *Mcm10* | 6.49E-05 | 1.547768 |
| ENSMUSG00000026358 | *Rgs1* | 2.94E-05 | 1.548109 |
| ENSMUSG00000114790 | *4921509O07Rik* | 1.41E-03 | 1.54851 |
| ENSMUSG00000115855 | *Gm34643* | 2.49E-02 | 1.548911 |
| ENSMUSG00000097062 | *Gm17586* | 4.91E-02 | 1.549771 |
| ENSMUSG00000037139 | *Myom3* | 5.04E-08 | 1.550368 |
| ENSMUSG00000028012 | *Rrh* | 5.89E-04 | 1.551343 |
| ENSMUSG00000052270 | *Fpr2* | 8.89E-17 | 1.551885 |
| ENSMUSG00000025017 | *Pik3ap1* | 2.80E-32 | 1.552126 |
| ENSMUSG00000042680 | *Garem1* | 2.43E-28 | 1.553264 |
| ENSMUSG00000083500 | *Gm15470* | 2.33E-02 | 1.553699 |
| ENSMUSG00000089617 | *Scarna10* | 1.33E-02 | 1.554248 |
| ENSMUSG00000099587 | *Gm28967* | 1.33E-02 | 1.554248 |
| ENSMUSG00000061723 | *Tnnt3* | 3.59E-06 | 1.554618 |
| ENSMUSG00000101930 | *Gm5441* | 4.71E-05 | 1.554924 |
| ENSMUSG00000047281 | *Sfn* | 1.18E-07 | 1.554958 |
| ENSMUSG00000003134 | *Tbc1d8* | 2.31E-17 | 1.555831 |
| ENSMUSG00000095098 | *Ccdc85b* | 1.46E-02 | 1.556789 |
| ENSMUSG00000105572 | *Gm43300* | 1.75E-04 | 1.557092 |
| ENSMUSG00000007877 | *Tcap* | 8.03E-10 | 1.557367 |
| ENSMUSG00000028896 | *Rcc1* | 6.05E-10 | 1.558192 |
| ENSMUSG00000104093 | *A330015K06Rik* | 2.94E-04 | 1.558524 |
| ENSMUSG00000049515 | *Espnl* | 1.85E-06 | 1.558599 |
| ENSMUSG00000105145 | *Gm47299* | 1.61E-02 | 1.559865 |
| ENSMUSG00000105788 | *Gm47295* | 1.61E-02 | 1.560047 |
| ENSMUSG00000030470 | *Csrp3* | 2.57E-10 | 1.560127 |
| ENSMUSG00000086350 | *B230369F24Rik* | 3.44E-02 | 1.561156 |
| ENSMUSG00000092134 | *Gm17089* | 1.34E-02 | 1.562976 |
| ENSMUSG00000087684 | *1200007C13Rik* | 1.65E-13 | 1.563081 |
| ENSMUSG00000031626 | *Sorbs2* | 1.55E-17 | 1.563623 |
| ENSMUSG00000109279 | *Gm45220* | 1.65E-07 | 1.564122 |
| ENSMUSG00000031780 | *Ccl17* | 2.29E-05 | 1.565128 |
| ENSMUSG00000110251 | *Gm2996* | 4.56E-09 | 1.565506 |
| ENSMUSG00000021591 | *Glrx* | 1.01E-19 | 1.565524 |
| ENSMUSG00000079597 | *Gm5483* | 5.00E-11 | 1.565997 |
| ENSMUSG00000097282 | *5031415H12Rik* | 2.32E-03 | 1.566965 |
| ENSMUSG00000086554 | *9530034E10Rik* | 1.85E-02 | 1.567387 |
| ENSMUSG00000109005 | *Gm45221* | 4.72E-09 | 1.567699 |
| ENSMUSG00000050439 | *Enthd1* | 1.23E-02 | 1.568498 |
| ENSMUSG00000020644 | *Id2* | 5.12E-18 | 1.570106 |
| ENSMUSG00000086584 | *Gm12002* | 1.57E-02 | 1.570339 |
| ENSMUSG00000079190 | *AC133103.1* | 6.72E-04 | 1.570523 |
| ENSMUSG00000107352 | *Gm43660* | 2.34E-04 | 1.570886 |
| ENSMUSG00000056569 | *Mpz* | 2.72E-08 | 1.570995 |
| ENSMUSG00000069873 | *4930438A08Rik* | 3.36E-02 | 1.571682 |
| ENSMUSG00000085249 | *Gm12847* | 2.96E-02 | 1.571826 |
| ENSMUSG00000052212 | *Cd177* | 3.33E-17 | 1.571901 |
| ENSMUSG00000031382 | *Asb11* | 2.67E-04 | 1.571915 |
| ENSMUSG00000087597 | *Gm15345* | 5.12E-03 | 1.573455 |

| | | | |
|---|---|---|---|
| ENSMUSG00000101751 | *Gm2427* | 5.74E-06 | 1.573571 |
| ENSMUSG00000024774 | *Ankrd22* | 7.98E-10 | 1.573578 |
| ENSMUSG00000078122 | *F630028O10Rik* | 7.10E-07 | 1.57378 |
| ENSMUSG00000104801 | *Gm43834* | 1.91E-08 | 1.574775 |
| ENSMUSG00000095260 | *Gm25890* | 2.05E-02 | 1.575215 |
| ENSMUSG00000094812 | *Gm22614* | 2.05E-02 | 1.575215 |
| ENSMUSG00000044997 | *E130304I02Rik* | 4.00E-02 | 1.575721 |
| ENSMUSG00000098702 | *1500015A07Rik* | 6.89E-04 | 1.576298 |
| ENSMUSG00000019933 | *Mrln* | 7.38E-03 | 1.577435 |
| ENSMUSG00000055972 | *2810407A14Rik* | 3.65E-09 | 1.577756 |
| ENSMUSG00000104406 | *Gm38014* | 2.17E-03 | 1.5782 |
| ENSMUSG00000046223 | *Plaur* | 3.97E-19 | 1.578531 |
| ENSMUSG00000031452 | *1700029H14Rik* | 7.05E-11 | 1.578888 |
| ENSMUSG00000044639 | *Gm11507* | 3.21E-02 | 1.580159 |
| ENSMUSG00000047604 | *Frat2* | 6.81E-03 | 1.580191 |
| ENSMUSG00000086765 | *Gm11827* | 2.45E-03 | 1.580289 |
| ENSMUSG00000090582 | *Gm17024* | 1.06E-06 | 1.58119 |
| ENSMUSG00000029377 | *Ereg* | 1.30E-07 | 1.581718 |
| ENSMUSG00000005686 | *Ampd3* | 3.84E-36 | 1.582624 |
| ENSMUSG00000055639 | *Dach1* | 2.45E-49 | 1.582772 |
| ENSMUSG00000048572 | *Tmem252* | 2.15E-30 | 1.583456 |
| ENSMUSG00000106741 | *4930513D17Rik* | 2.31E-03 | 1.583518 |
| ENSMUSG00000104449 | *Gm37255* | 6.66E-04 | 1.583657 |
| ENSMUSG00000034226 | *Rhov* | 1.51E-03 | 1.583825 |
| ENSMUSG00000034990 | *Otoa* | 2.27E-03 | 1.584135 |
| ENSMUSG00000038059 | *Smim3* | 2.50E-27 | 1.584279 |
| ENSMUSG00000064288 | *Hist1h4k* | 2.55E-02 | 1.584323 |
| ENSMUSG00000067049 | *Unc93a* | 3.55E-03 | 1.584342 |
| ENSMUSG00000114662 | *Gm31683* | 3.36E-03 | 1.584857 |
| ENSMUSG00000071561 | *BC100530* | 2.75E-07 | 1.585823 |
| ENSMUSG00000056025 | *Clca3a1* | 1.39E-09 | 1.586042 |
| ENSMUSG00000064923 | *Gm22042* | 1.82E-02 | 1.586766 |
| ENSMUSG00000087362 | *Gm13710* | 3.87E-05 | 1.588172 |
| ENSMUSG00000115764 | *Gm18811* | 4.77E-02 | 1.588766 |
| ENSMUSG00000102135 | *Gm37108* | 6.83E-03 | 1.589787 |
| ENSMUSG00000018930 | *Ccl4* | 3.25E-14 | 1.592018 |
| ENSMUSG00000092920 | *Mirt2* | 2.33E-03 | 1.593218 |
| ENSMUSG00000003484 | *Cyp4f18* | 1.28E-11 | 1.594494 |
| ENSMUSG00000068855 | *Hist2h2ac* | 2.26E-02 | 1.595197 |
| ENSMUSG00000077192 | *Snora17* | 1.22E-02 | 1.595366 |
| ENSMUSG00000112148 | *Lilrb4a* | 6.29E-08 | 1.595991 |
| ENSMUSG00000053914 | *Kdm4d* | 3.47E-07 | 1.598458 |
| ENSMUSG00000078905 | *Gm14393* | 2.09E-02 | 1.598898 |
| ENSMUSG00000055301 | *Adh7* | 3.36E-04 | 1.599546 |
| ENSMUSG00000025804 | *Ccr1* | 2.69E-15 | 1.600075 |
| ENSMUSG00000025236 | *Adpgk* | 4.54E-06 | 1.600129 |
| ENSMUSG00000028747 | *Htr6* | 3.61E-02 | 1.600441 |
| ENSMUSG00000026452 | *Syt2* | 4.75E-12 | 1.600466 |
| ENSMUSG00000095438 | *Mir133a-1hg* | 8.40E-04 | 1.60088 |
| ENSMUSG00000107092 | *Gm7993* | 3.05E-09 | 1.601292 |
| ENSMUSG00000099061 | *Gm28050* | 3.42E-06 | 1.601401 |
| ENSMUSG00000100182 | *1810006J02Rik* | 2.34E-03 | 1.601528 |
| ENSMUSG00000080845 | *Gm9115* | 9.28E-06 | 1.602056 |

| | | | |
|---|---|---|---|
| ENSMUSG00000021702 | *Thbs4* | 5.37E-07 | 1.602758 |
| ENSMUSG00000094733 | *Gm5416* | 2.35E-02 | 1.603974 |
| ENSMUSG00000044719 | *E230025N22Rik* | 7.14E-03 | 1.604274 |
| ENSMUSG00000031538 | *Plat* | 1.12E-27 | 1.6045 |
| ENSMUSG00000071562 | *Stfa1* | 1.25E-02 | 1.604593 |
| ENSMUSG00000116760 | *AC164314.1* | 1.55E-18 | 1.605019 |
| ENSMUSG00000062028 | *Irgc1* | 1.27E-03 | 1.605428 |
| ENSMUSG00000020108 | *Ddit4* | 3.87E-18 | 1.605556 |
| ENSMUSG00000097168 | *C230088H06Rik* | 1.53E-07 | 1.607501 |
| ENSMUSG00000059657 | *Stfa2l1* | 4.70E-13 | 1.608177 |
| ENSMUSG00000054383 | *Pnma1* | 1.59E-02 | 1.609741 |
| ENSMUSG00000030235 | *Slco1c1* | 8.01E-11 | 1.610186 |
| ENSMUSG00000079800 | *AC125149.3* | 5.67E-04 | 1.610687 |
| ENSMUSG00000047330 | *Kcne4* | 1.72E-12 | 1.611245 |
| ENSMUSG00000011179 | *Odc1* | 6.77E-19 | 1.611702 |
| ENSMUSG00000088185 | *Scarna2* | 2.29E-04 | 1.611916 |
| ENSMUSG00000029635 | *Cdk8* | 2.85E-05 | 1.613325 |
| ENSMUSG00000040747 | *Cd53* | 2.71E-18 | 1.613354 |
| ENSMUSG00000104740 | *Mir6516* | 3.89E-02 | 1.613694 |
| ENSMUSG00000086275 | *1700121C08Rik* | 1.98E-03 | 1.614578 |
| ENSMUSG00000046178 | *Nxph1* | 3.18E-02 | 1.614789 |
| ENSMUSG00000046101 | *Mcmdc2* | 2.24E-02 | 1.615786 |
| ENSMUSG00000114980 | *4933432I03Rik* | 3.86E-13 | 1.617126 |
| ENSMUSG00000112105 | *Rps7-ps2* | 2.09E-03 | 1.617464 |
| ENSMUSG00000024215 | *Spdef* | 3.27E-04 | 1.617571 |
| ENSMUSG00000087436 | *Gm16156* | 1.33E-02 | 1.618843 |
| ENSMUSG00000095701 | *Gm24830* | 1.23E-02 | 1.619865 |
| ENSMUSG00000096838 | *Gm26232* | 1.23E-02 | 1.619865 |
| ENSMUSG00000065773 | *Rnu1b6* | 1.23E-02 | 1.619865 |
| ENSMUSG00000029378 | *Areg* | 8.57E-05 | 1.620274 |
| ENSMUSG00000030144 | *Clec4d* | 1.13E-16 | 1.621129 |
| ENSMUSG00000089999 | *Gm6485* | 1.16E-11 | 1.621898 |
| ENSMUSG00000097060 | *Gm26759* | 2.64E-04 | 1.622397 |
| ENSMUSG00000043613 | *Mmp3* | 2.40E-04 | 1.624738 |
| ENSMUSG00000106607 | *Gm43178* | 2.36E-02 | 1.625189 |
| ENSMUSG00000064941 | *Gm23238* | 2.37E-03 | 1.625244 |
| ENSMUSG00000045102 | *Poln* | 6.51E-23 | 1.627181 |
| ENSMUSG00000074146 | *4930579C12Rik* | 9.25E-09 | 1.627321 |
| ENSMUSG00000061878 | *Sphk1* | 2.13E-23 | 1.628004 |
| ENSMUSG00000067555 | *Gm10212* | 4.20E-03 | 1.628146 |
| ENSMUSG00000086993 | *Rsf1os2* | 1.93E-02 | 1.629504 |
| ENSMUSG00000116747 | *AC144408.2* | 2.19E-02 | 1.630024 |
| ENSMUSG00000069273 | *Hist1h3e* | 8.64E-03 | 1.630123 |
| ENSMUSG00000085743 | *8430419K02Rik* | 5.99E-13 | 1.63082 |
| ENSMUSG00000053338 | *Tarm1* | 1.27E-22 | 1.63113 |
| ENSMUSG00000024471 | *Myot* | 2.74E-09 | 1.631316 |
| ENSMUSG00000046610 | *Oacyl* | 1.57E-04 | 1.631988 |
| ENSMUSG00000114125 | *1700037F03Rik* | 1.56E-02 | 1.632073 |
| ENSMUSG00000027832 | *Ptx3* | 2.36E-34 | 1.633264 |
| ENSMUSG00000092397 | *C130080G10Rik* | 3.11E-06 | 1.634192 |
| ENSMUSG00000099803 | *Gm28863* | 4.44E-02 | 1.634596 |
| ENSMUSG00000100779 | *Gm29292* | 5.73E-14 | 1.635041 |
| ENSMUSG00000032523 | *Hhatl* | 1.82E-03 | 1.63623 |

| | | | |
|---|---|---|---|
| ENSMUSG00000028125 | *Abca4* | 1.97E-12 | 1.637562 |
| ENSMUSG00000038037 | *Socs1* | 4.73E-08 | 1.638886 |
| ENSMUSG00000031872 | *Bean1* | 1.38E-10 | 1.639807 |
| ENSMUSG00000049734 | *Trex1* | 2.49E-11 | 1.640014 |
| ENSMUSG00000095580 | *Rnu1b1* | 1.33E-02 | 1.641977 |
| ENSMUSG00000093834 | *Rnu1b2* | 1.33E-02 | 1.641977 |
| ENSMUSG00000009633 | *G0s2* | 2.23E-07 | 1.642205 |
| ENSMUSG00000002847 | *Pla1a* | 3.33E-22 | 1.642753 |
| ENSMUSG00000043008 | *Klhl6* | 7.14E-15 | 1.642823 |
| ENSMUSG00000062461 | *Gm5453* | 8.90E-06 | 1.643537 |
| ENSMUSG00000045455 | *Gm9797* | 4.46E-02 | 1.644519 |
| ENSMUSG00000082884 | *Gm13339* | 5.60E-05 | 1.644853 |
| ENSMUSG00000052477 | *C130026I21Rik* | 6.00E-27 | 1.645118 |
| ENSMUSG00000040490 | *Lrfn2* | 1.41E-02 | 1.645149 |
| ENSMUSG00000113701 | *B230303A05Rik* | 4.29E-04 | 1.645667 |
| ENSMUSG00000104560 | *Gm8115* | 6.75E-04 | 1.645915 |
| ENSMUSG00000028396 | *2310002L09Rik* | 4.23E-02 | 1.649697 |
| ENSMUSG00000024521 | *Pmaip1* | 2.02E-17 | 1.652637 |
| ENSMUSG00000066361 | *Serpina3c* | 4.62E-21 | 1.652992 |
| ENSMUSG00000091971 | *Hspa1a* | 3.82E-05 | 1.65332 |
| ENSMUSG00000027014 | *Cwc22* | 6.70E-06 | 1.653834 |
| ENSMUSG00000085779 | *Atcayos* | 1.32E-11 | 1.655658 |
| ENSMUSG00000079462 | *Gm15737* | 4.27E-02 | 1.655772 |
| ENSMUSG00000069305 | *Hist1h4n* | 1.34E-02 | 1.65667 |
| ENSMUSG00000108414 | *Snhg1* | 6.02E-13 | 1.656675 |
| ENSMUSG00000023341 | *Mx2* | 6.24E-29 | 1.656679 |
| ENSMUSG00000096957 | *E230013L22Rik* | 2.31E-09 | 1.657441 |
| ENSMUSG00000082838 | *Gm14519* | 1.01E-05 | 1.657928 |
| ENSMUSG00000095500 | *AC132444.5* | 1.55E-02 | 1.658094 |
| ENSMUSG00000035692 | *Isg15* | 5.98E-17 | 1.659027 |
| ENSMUSG00000031972 | *Acta1* | 2.51E-10 | 1.660693 |
| ENSMUSG00000115518 | *Gm10791* | 1.14E-03 | 1.661218 |
| ENSMUSG00000028989 | *Angptl7* | 1.47E-05 | 1.661267 |
| ENSMUSG00000030592 | *Ryr1* | 3.10E-08 | 1.662936 |
| ENSMUSG00000087968 | *Gm25395* | 3.29E-04 | 1.663115 |
| ENSMUSG00000074218 | *Cox7a1* | 2.46E-05 | 1.664493 |
| ENSMUSG00000097927 | *Gm6999* | 1.40E-06 | 1.664539 |
| ENSMUSG00000024481 | *Lvrn* | 1.55E-15 | 1.664843 |
| ENSMUSG00000051748 | *Wfdc21* | 6.59E-11 | 1.665772 |
| ENSMUSG00000056133 | *Unc93a2* | 1.42E-02 | 1.666035 |
| ENSMUSG00000022821 | *Hgd* | 9.91E-04 | 1.666819 |
| ENSMUSG00000077391 | *Gm24336* | 9.21E-03 | 1.667557 |
| ENSMUSG00000112895 | *Gm47567* | 1.94E-03 | 1.668933 |
| ENSMUSG00000097414 | *B130046B21Rik* | 5.30E-06 | 1.670513 |
| ENSMUSG00000057003 | *Myh4* | 1.07E-10 | 1.67121 |
| ENSMUSG00000096010 | *Hist4h4* | 2.64E-03 | 1.673569 |
| ENSMUSG00000113960 | *4933412O06Rik* | 5.41E-03 | 1.673653 |
| ENSMUSG00000041616 | *Nppa* | 2.60E-04 | 1.674788 |
| ENSMUSG00000108436 | *Gm44851* | 2.65E-02 | 1.675059 |
| ENSMUSG00000052496 | *Pkdrej* | 8.35E-03 | 1.675102 |
| ENSMUSG00000032845 | *Alpk2* | 2.46E-22 | 1.67545 |
| ENSMUSG00000006345 | *Ggt1* | 2.80E-06 | 1.675688 |
| ENSMUSG00000030672 | *Mylpf* | 2.51E-04 | 1.677392 |

| | | | |
|---|---|---|---|
| ENSMUSG00000075122 | *Cd80* | 1.39E-25 | 1.678083 |
| ENSMUSG00000100301 | *6030407O03Rik* | 1.46E-17 | 1.678651 |
| ENSMUSG00000091952 | *Gm17709* | 3.50E-11 | 1.679019 |
| ENSMUSG00000022876 | *Samsn1* | 2.79E-24 | 1.680546 |
| ENSMUSG00000113178 | *Mylf-ps* | 3.93E-08 | 1.680637 |
| ENSMUSG00000095620 | *2010005H15Rik* | 3.96E-03 | 1.680691 |
| ENSMUSG00000054258 | *Gm5082* | 3.59E-02 | 1.681204 |
| ENSMUSG00000070498 | *Tmem132b* | 3.01E-11 | 1.68225 |
| ENSMUSG00000056856 | *Jakmip3* | 3.89E-03 | 1.683214 |
| ENSMUSG00000111324 | *Gm31410* | 6.04E-03 | 1.68436 |
| ENSMUSG00000054204 | *Alkal2* | 1.32E-02 | 1.6846 |
| ENSMUSG00000078902 | *Gm14443* | 6.61E-03 | 1.684691 |
| ENSMUSG00000076135 | *Gm24276* | 3.81E-02 | 1.685494 |
| ENSMUSG00000000204 | *Slfn4* | 1.36E-09 | 1.686 |
| ENSMUSG00000005800 | *Mmp8* | 3.48E-29 | 1.686464 |
| ENSMUSG00000001249 | *Hpn* | 7.62E-12 | 1.687075 |
| ENSMUSG00000041827 | *Oasl1* | 9.65E-18 | 1.687086 |
| ENSMUSG00000074452 | *Pate2* | 1.83E-10 | 1.688618 |
| ENSMUSG00000045551 | *Fpr1* | 1.35E-39 | 1.689129 |
| ENSMUSG00000053641 | *Dennd4a* | 2.32E-30 | 1.689402 |
| ENSMUSG00000104818 | *Gm43661* | 1.29E-03 | 1.689966 |
| ENSMUSG00000109232 | *Gm44577* | 4.63E-03 | 1.690024 |
| ENSMUSG00000048489 | *Depp1* | 1.22E-08 | 1.690058 |
| ENSMUSG00000093378 | *Gm20663* | 1.10E-02 | 1.691727 |
| ENSMUSG00000049134 | *Nrap* | 6.44E-22 | 1.692374 |
| ENSMUSG00000051832 | *E230016K23Rik* | 8.52E-07 | 1.694895 |
| ENSMUSG00000052837 | *Junb* | 1.10E-23 | 1.695927 |
| ENSMUSG00000069274 | *Hist1h4f* | 1.44E-02 | 1.701594 |
| ENSMUSG00000086836 | *Gm13748* | 1.50E-03 | 1.701696 |
| ENSMUSG00000028680 | *Plk3* | 1.48E-17 | 1.701976 |
| ENSMUSG00000032174 | *Icam5* | 1.01E-02 | 1.705483 |
| ENSMUSG00000087113 | *Gm11714* | 3.79E-04 | 1.706526 |
| ENSMUSG00000111514 | *E230014E18Rik* | 7.88E-06 | 1.708494 |
| ENSMUSG00000064382 | *Gm26447* | 1.31E-04 | 1.709249 |
| ENSMUSG00000107667 | *C530044C16Rik* | 6.95E-03 | 1.709767 |
| ENSMUSG00000057580 | *Gm10012* | 5.96E-03 | 1.711919 |
| ENSMUSG00000062593 | *Gm49339* | 8.13E-09 | 1.712107 |
| ENSMUSG00000047216 | *Cdh19* | 5.39E-12 | 1.712183 |
| ENSMUSG00000092626 | *9130230N09Rik* | 3.71E-02 | 1.712613 |
| ENSMUSG00000109032 | *Gm7972* | 5.34E-05 | 1.712639 |
| ENSMUSG00000087159 | *Gm15246* | 5.79E-06 | 1.712843 |
| ENSMUSG00000071637 | *Cebpd* | 2.78E-33 | 1.713579 |
| ENSMUSG00000027737 | *Slc7a11* | 7.04E-24 | 1.713985 |
| ENSMUSG00000109729 | *Gm45418* | 1.79E-06 | 1.714339 |
| ENSMUSG00000026068 | *Il18rap* | 1.63E-22 | 1.714997 |
| ENSMUSG00000097263 | *Gm26804* | 1.62E-02 | 1.715908 |
| ENSMUSG00000109052 | *Gm45012* | 1.67E-02 | 1.716023 |
| ENSMUSG00000029862 | *Clcn1* | 8.39E-06 | 1.716184 |
| ENSMUSG00000022582 | *Ly6g* | 3.58E-05 | 1.718298 |
| ENSMUSG00000042918 | *Mamstr* | 7.60E-18 | 1.719004 |
| ENSMUSG00000027505 | *Fam209* | 8.93E-03 | 1.720522 |
| ENSMUSG00000092201 | *A530058N18Rik* | 6.08E-09 | 1.721383 |
| ENSMUSG00000102349 | *Gm37376* | 7.33E-03 | 1.722299 |

| | | | |
|---|---|---|---|
| ENSMUSG00000102051 | *I830127L07Rik* | 2.66E-06 | 1.722921 |
| ENSMUSG00000094306 | *Gm24924* | 3.54E-03 | 1.723532 |
| ENSMUSG00000027470 | *Mylk2* | 8.88E-09 | 1.72462 |
| ENSMUSG00000088008 | *Gm25492* | 2.74E-02 | 1.727829 |
| ENSMUSG00000029384 | *2010109A12Rik* | 9.32E-06 | 1.729412 |
| ENSMUSG00000104010 | *Gm37366* | 3.43E-04 | 1.729563 |
| ENSMUSG00000031710 | *Ucp1* | 4.71E-04 | 1.730521 |
| ENSMUSG00000064841 | *Gm26205* | 3.76E-02 | 1.732023 |
| ENSMUSG00000087059 | *Gm12339* | 3.57E-10 | 1.73504 |
| ENSMUSG00000059741 | *Myl3* | 2.57E-09 | 1.735257 |
| ENSMUSG00000030399 | *Ckm* | 6.58E-09 | 1.737508 |
| ENSMUSG00000072720 | *Myo18b* | 2.53E-31 | 1.738448 |
| ENSMUSG00000100733 | *4932411K12Rik* | 8.48E-06 | 1.74245 |
| ENSMUSG00000013483 | *Card14* | 1.29E-04 | 1.743775 |
| ENSMUSG00000026407 | *Cacna1s* | 4.58E-10 | 1.743939 |
| ENSMUSG00000085241 | *Snhg3* | 2.11E-06 | 1.74396 |
| ENSMUSG00000097122 | *Gm26624* | 2.89E-10 | 1.744094 |
| ENSMUSG00000032661 | *Oas3* | 1.63E-17 | 1.74446 |
| ENSMUSG00000060639 | *Hist1h4i* | 6.34E-03 | 1.746403 |
| ENSMUSG00000092539 | *Gm20468* | 3.84E-02 | 1.747708 |
| ENSMUSG00000100147 | *1700047M11Rik* | 3.91E-08 | 1.748016 |
| ENSMUSG00000070524 | *Fcrlb* | 1.24E-02 | 1.748075 |
| ENSMUSG00000095616 | *Gm26244* | 3.02E-03 | 1.749841 |
| ENSMUSG00000069306 | *Hist1h4m* | 1.13E-02 | 1.750899 |
| ENSMUSG00000101191 | *Gm28809* | 3.20E-02 | 1.7511 |
| ENSMUSG00000097358 | *Gm26773* | 2.59E-03 | 1.753117 |
| ENSMUSG00000061816 | *Myl1* | 1.60E-12 | 1.754682 |
| ENSMUSG00000054169 | *Ceacam10* | 9.98E-09 | 1.754693 |
| ENSMUSG00000105826 | *Gm42141* | 2.16E-02 | 1.754897 |
| ENSMUSG00000105790 | *Gm24105* | 3.07E-04 | 1.75526 |
| ENSMUSG00000092674 | *Gm24105* | 2.58E-04 | 1.75676 |
| ENSMUSG00000056071 | *S100a9* | 2.01E-14 | 1.760172 |
| ENSMUSG00000069792 | *Wfdc17* | 2.66E-11 | 1.760299 |
| ENSMUSG00000022651 | *Retnlg* | 1.87E-24 | 1.766124 |
| ENSMUSG00000023473 | *Celsr3* | 3.06E-07 | 1.767531 |
| ENSMUSG00000101344 | *Gm29183* | 4.83E-20 | 1.767788 |
| ENSMUSG00000050931 | *Sgms2* | 5.11E-55 | 1.768559 |
| ENSMUSG00000091119 | *Ccdc152* | 2.18E-04 | 1.768666 |
| ENSMUSG00000097413 | *A830052D11Rik* | 2.01E-02 | 1.771034 |
| ENSMUSG00000091383 | *Hist1h2al* | 9.26E-06 | 1.771647 |
| ENSMUSG00000064853 | *Gm23442* | 3.06E-04 | 1.773453 |
| ENSMUSG00000099092 | *Mir7062* | 1.37E-02 | 1.773986 |
| ENSMUSG00000052767 | *Gm12703* | 5.22E-04 | 1.775187 |
| ENSMUSG00000111752 | *Gm38575* | 1.61E-15 | 1.776614 |
| ENSMUSG00000043789 | *Vwce* | 1.79E-04 | 1.778777 |
| ENSMUSG00000109831 | *Gm36431* | 3.33E-02 | 1.779631 |
| ENSMUSG00000039653 | *Baat* | 2.08E-02 | 1.779818 |
| ENSMUSG00000050553 | *Gk2* | 1.02E-02 | 1.780074 |
| ENSMUSG00000022824 | *Muc13* | 2.08E-05 | 1.781872 |
| ENSMUSG00000027855 | *Sycp1* | 2.52E-02 | 1.785086 |
| ENSMUSG00000102343 | *Gm37381* | 2.22E-03 | 1.785986 |
| ENSMUSG00000058427 | *Cxcl2* | 6.05E-12 | 1.786152 |
| ENSMUSG00000031097 | *Tnni2* | 6.59E-11 | 1.786684 |

| | | | |
|---|---|---|---|
| ENSMUSG00000072875 | *Gpr27* | 2.99E-05 | 1.787666 |
| ENSMUSG00000026100 | *Mstn* | 1.84E-02 | 1.787836 |
| ENSMUSG00000090698 | *Apold1* | 1.30E-17 | 1.788991 |
| ENSMUSG00000027824 | *Vmn2r1* | 1.85E-02 | 1.78902 |
| ENSMUSG00000116508 | *Gm49463* | 2.15E-02 | 1.789422 |
| ENSMUSG00000088529 | *Gm26083* | 3.34E-03 | 1.790452 |
| ENSMUSG00000081857 | *Gm13624* | 1.87E-03 | 1.79269 |
| ENSMUSG00000077323 | *Rnu11* | 2.17E-03 | 1.793382 |
| ENSMUSG00000022602 | *Arc* | 6.32E-03 | 1.79405 |
| ENSMUSG00000044367 | *Slc16a13* | 8.06E-21 | 1.795745 |
| ENSMUSG00000109096 | *Gm44888* | 2.17E-03 | 1.796222 |
| ENSMUSG00000085792 | *Gm15414* | 2.28E-02 | 1.796476 |
| ENSMUSG00000115432 | *D130009I18Rik* | 1.25E-02 | 1.7966 |
| ENSMUSG00000074001 | *Klhl40* | 2.39E-03 | 1.804207 |
| ENSMUSG00000024827 | *Gldc* | 2.59E-02 | 1.808096 |
| ENSMUSG00000110626 | *Gm45805* | 1.93E-03 | 1.809283 |
| ENSMUSG00000102759 | *Gm10463* | 4.37E-02 | 1.811725 |
| ENSMUSG00000108929 | *Cc2d2b* | 3.12E-04 | 1.814541 |
| ENSMUSG00000048388 | *Fam171b* | 3.36E-17 | 1.816202 |
| ENSMUSG00000089844 | *A530032D15Rik* | 3.03E-10 | 1.816284 |
| ENSMUSG00000043795 | *Prr33* | 4.01E-02 | 1.817973 |
| ENSMUSG00000027456 | *Sdcbp2* | 1.73E-11 | 1.818137 |
| ENSMUSG00000095969 | *Rnu1a1* | 7.72E-04 | 1.823663 |
| ENSMUSG00000041872 | *Il17f* | 1.92E-06 | 1.828096 |
| ENSMUSG00000001482 | *Def8* | 2.17E-14 | 1.828763 |
| ENSMUSG00000006724 | *Cyp27b1* | 7.32E-03 | 1.82998 |
| ENSMUSG00000101279 | *Gm18342* | 1.51E-05 | 1.831412 |
| ENSMUSG00000109157 | *Gm44829* | 8.56E-05 | 1.83179 |
| ENSMUSG00000056300 | *Zfp981* | 3.27E-10 | 1.83268 |
| ENSMUSG00000077714 | *Snord17* | 2.21E-04 | 1.835137 |
| ENSMUSG00000034683 | *Ppp1r1c* | 1.37E-02 | 1.836685 |
| ENSMUSG00000087700 | *Gm15283* | 3.63E-04 | 1.838693 |
| ENSMUSG00000096214 | *Gm22634* | 4.41E-06 | 1.840548 |
| ENSMUSG00000096206 | *Gm22317* | 4.41E-06 | 1.840548 |
| ENSMUSG00000110411 | *Gm45457* | 3.21E-03 | 1.842368 |
| ENSMUSG00000115420 | *AL732506.1* | 3.41E-05 | 1.84363 |
| ENSMUSG00000088088 | *Rmrp* | 3.41E-05 | 1.84363 |
| ENSMUSG00000070167 | *Snora57* | 6.49E-06 | 1.843722 |
| ENSMUSG00000044083 | *Efcab8* | 8.89E-24 | 1.847127 |
| ENSMUSG00000096205 | *Gm22068* | 4.44E-04 | 1.848083 |
| ENSMUSG00000034755 | *Pcdh11x* | 1.69E-07 | 1.849981 |
| ENSMUSG00000064655 | *Gm25788* | 3.42E-02 | 1.850018 |
| ENSMUSG00000112082 | *Gm35035* | 2.70E-03 | 1.850562 |
| ENSMUSG00000096659 | *Gm25679* | 6.20E-06 | 1.851418 |
| ENSMUSG00000093815 | *Gm26444* | 6.20E-06 | 1.851418 |
| ENSMUSG00000044092 | *C130050O18Rik* | 4.16E-07 | 1.85322 |
| ENSMUSG00000060093 | *Hist1h4a* | 6.83E-05 | 1.854758 |
| ENSMUSG00000112023 | *Lilr4b* | 3.87E-09 | 1.856512 |
| ENSMUSG00000030730 | *Atp2a1* | 2.57E-25 | 1.856765 |
| ENSMUSG00000113185 | *Gm47127* | 2.52E-14 | 1.857191 |
| ENSMUSG00000087203 | *Gm13986* | 9.23E-38 | 1.857802 |
| ENSMUSG00000017300 | *Tnnc2* | 2.32E-13 | 1.860495 |
| ENSMUSG00000005716 | *Pvalb* | 5.17E-03 | 1.862103 |

| | | | |
|---|---|---|---|
| ENSMUSG00000064179 | *Tnnt1* | 3.30E-04 | 1.865691 |
| ENSMUSG00000103907 | *Gm37498* | 2.41E-19 | 1.868144 |
| ENSMUSG00000094826 | *Gm23804* | 6.28E-06 | 1.870232 |
| ENSMUSG00000107480 | *Gm44165* | 6.63E-05 | 1.871722 |
| ENSMUSG00000050304 | *Slc25a2* | 1.14E-02 | 1.87573 |
| ENSMUSG00000065862 | *Gm24029* | 6.61E-04 | 1.875777 |
| ENSMUSG00000064453 | *Snord21* | 3.67E-02 | 1.876243 |
| ENSMUSG00000096972 | *Gm26883* | 3.02E-04 | 1.878023 |
| ENSMUSG00000002500 | *Rpl3l* | 4.07E-05 | 1.878812 |
| ENSMUSG00000065145 | *Vaultrc5* | 9.73E-04 | 1.881338 |
| ENSMUSG00000097052 | *Snora43* | 4.79E-02 | 1.882312 |
| ENSMUSG00000025473 | *Adam8* | 3.65E-14 | 1.883209 |
| ENSMUSG00000025929 | *Il17a* | 1.27E-02 | 1.884795 |
| ENSMUSG00000069733 | *Ube2u* | 1.49E-03 | 1.885786 |
| ENSMUSG00000060988 | *Galnt13* | 6.96E-12 | 1.886176 |
| ENSMUSG00000064387 | *Snora73a* | 4.95E-09 | 1.88777 |
| ENSMUSG00000110125 | *Gm33148* | 6.45E-03 | 1.888934 |
| ENSMUSG00000082069 | *Gm16242* | 1.52E-03 | 1.890196 |
| ENSMUSG00000042451 | *Mybph* | 7.08E-19 | 1.89042 |
| ENSMUSG00000074071 | *Fam169b* | 1.22E-62 | 1.892733 |
| ENSMUSG00000062393 | *Dgkk* | 3.95E-02 | 1.894875 |
| ENSMUSG00000104554 | *Gm4610* | 1.51E-04 | 1.897808 |
| ENSMUSG00000032021 | *Crtam* | 1.06E-17 | 1.89973 |
| ENSMUSG00000092805 | *Gm26461* | 7.94E-04 | 1.903632 |
| ENSMUSG00000060470 | *Adgrg3* | 5.50E-20 | 1.903686 |
| ENSMUSG00000097081 | *Gm10425* | 2.22E-02 | 1.906599 |
| ENSMUSG00000032066 | *Bco2* | 1.74E-03 | 1.906705 |
| ENSMUSG00000092267 | *Gm20417* | 5.68E-07 | 1.908725 |
| ENSMUSG00000025892 | *Gria4* | 4.06E-04 | 1.909271 |
| ENSMUSG00000042567 | *Nek10* | 1.49E-18 | 1.909826 |
| ENSMUSG00000115426 | *Gm19510* | 3.61E-02 | 1.911947 |
| ENSMUSG00000116641 | *AC165271.1* | 4.77E-02 | 1.912683 |
| ENSMUSG00000108332 | *D530033B14Rik* | 3.38E-17 | 1.912716 |
| ENSMUSG00000038067 | *Csf3* | 5.24E-28 | 1.912958 |
| ENSMUSG00000043681 | *Fam25c* | 9.06E-03 | 1.913562 |
| ENSMUSG00000064999 | *Gm26035* | 1.27E-04 | 1.915529 |
| ENSMUSG00000084929 | *Foxo6os* | 2.44E-02 | 1.916095 |
| ENSMUSG00000113321 | *Gm8075* | 1.08E-04 | 1.9166 |
| ENSMUSG00000080542 | *Gm22710* | 9.10E-03 | 1.916605 |
| ENSMUSG00000097622 | *A330033J07Rik* | 1.30E-02 | 1.916889 |
| ENSMUSG00000111971 | *Gm48678* | 1.01E-02 | 1.918064 |
| ENSMUSG00000065353 | *Snora73b* | 4.31E-05 | 1.92082 |
| ENSMUSG00000079451 | *Tmprss11g* | 1.05E-02 | 1.922677 |
| ENSMUSG00000057280 | *Musk* | 4.21E-13 | 1.924024 |
| ENSMUSG00000026166 | *Ccl20* | 6.82E-22 | 1.924662 |
| ENSMUSG00000043873 | *Chil5* | 3.16E-05 | 1.926813 |
| ENSMUSG00000099470 | *Gm29340* | 6.43E-03 | 1.928433 |
| ENSMUSG00000038508 | *Gdf15* | 4.48E-03 | 1.930522 |
| ENSMUSG00000085434 | *Gm11725* | 1.19E-03 | 1.931465 |
| ENSMUSG00000049173 | *Myoz3* | 4.24E-05 | 1.93273 |
| ENSMUSG00000097417 | *Gm26669* | 3.24E-02 | 1.932952 |
| ENSMUSG00000052581 | *Lrrtm4* | 1.64E-07 | 1.934282 |
| ENSMUSG00000116975 | *AC097366.1* | 2.80E-03 | 1.934645 |

| | | | |
|---|---|---|---|
| ENSMUSG00000107722 | *2900060B14Rik* | 4.30E-03 | 1.93608 |
| ENSMUSG00000044165 | *Bcl2l15* | 2.49E-09 | 1.936183 |
| ENSMUSG00000104728 | *Gm42462* | 3.00E-14 | 1.936263 |
| ENSMUSG00000022622 | *Acr* | 4.06E-11 | 1.941514 |
| ENSMUSG00000065226 | *Gm25791* | 1.83E-06 | 1.942482 |
| ENSMUSG00000109002 | *Gm38405* | 2.93E-04 | 1.943877 |
| ENSMUSG00000055523 | *Gucy2g* | 2.09E-02 | 1.944978 |
| ENSMUSG00000024912 | *Fosl1* | 4.08E-15 | 1.946069 |
| ENSMUSG00000049100 | *Pcdh10* | 1.63E-02 | 1.947579 |
| ENSMUSG00000100334 | *C230024C17Rik* | 1.73E-02 | 1.947699 |
| ENSMUSG00000102416 | *4933424G06Rik* | 2.21E-02 | 1.951006 |
| ENSMUSG00000042895 | *Abra* | 1.14E-09 | 1.951135 |
| ENSMUSG00000102630 | *Gm37289* | 5.47E-03 | 1.953604 |
| ENSMUSG00000037705 | *Tecta* | 1.30E-02 | 1.953664 |
| ENSMUSG00000085750 | *Gm14009* | 2.26E-02 | 1.959056 |
| ENSMUSG00000106392 | *Gm5870* | 2.74E-02 | 1.959897 |
| ENSMUSG00000116961 | *AC117662.4* | 7.69E-07 | 1.970064 |
| ENSMUSG00000029379 | *Cxcl3* | 5.85E-31 | 1.970535 |
| ENSMUSG00000056054 | *S100a8* | 2.08E-14 | 1.971878 |
| ENSMUSG00000009070 | *Rsph14* | 1.08E-09 | 1.971919 |
| ENSMUSG00000106649 | *Gm49349* | 1.69E-06 | 1.97517 |
| ENSMUSG00000115355 | *4930445E18Rik* | 3.15E-07 | 1.98206 |
| ENSMUSG00000084783 | *Gm15419* | 3.73E-04 | 1.984549 |
| ENSMUSG00000048806 | *Ifnb1* | 4.47E-02 | 1.985866 |
| ENSMUSG00000112683 | *Gm18904* | 5.10E-03 | 1.988548 |
| ENSMUSG00000103626 | *Gm37358* | 4.92E-04 | 1.992354 |
| ENSMUSG00000087090 | *Nctc1* | 3.78E-18 | 1.993635 |
| ENSMUSG00000110498 | *A630001O12Rik* | 2.63E-06 | 1.993703 |
| ENSMUSG00000097643 | *A130051J06Rik* | 1.72E-03 | 1.997776 |
| ENSMUSG00000024430 | *Cabyr* | 3.39E-04 | 1.999224 |
| ENSMUSG00000065701 | *Rny1* | 7.04E-04 | 2.000426 |
| ENSMUSG00000104956 | *4930429D17Rik* | 4.22E-03 | 2.000856 |
| ENSMUSG00000046049 | *Rp1l1* | 5.00E-04 | 2.002651 |
| ENSMUSG00000089647 | *Gm2245* | 4.25E-03 | 2.00392 |
| ENSMUSG00000034145 | *Tmem63c* | 1.54E-03 | 2.004762 |
| ENSMUSG00000097300 | *Gm26835* | 2.99E-02 | 2.005741 |
| ENSMUSG00000058740 | *Kcnt1* | 1.78E-11 | 2.006095 |
| ENSMUSG00000032496 | *Ltf* | 1.13E-45 | 2.006145 |
| ENSMUSG00000065126 | *Snord104* | 6.05E-07 | 2.006554 |
| ENSMUSG00000097071 | *4930544I03Rik* | 1.01E-07 | 2.006732 |
| ENSMUSG00000077167 | *Gm24119* | 1.08E-02 | 2.009462 |
| ENSMUSG00000110622 | *Iqcn* | 1.16E-02 | 2.011367 |
| ENSMUSG00000071470 | *Ccnb1ip1* | 2.89E-04 | 2.016759 |
| ENSMUSG00000037977 | *6430571L13Rik* | 2.38E-05 | 2.018356 |
| ENSMUSG00000053469 | *Tg* | 9.23E-38 | 2.022512 |
| ENSMUSG00000080950 | *Gm7278* | 9.59E-05 | 2.024914 |
| ENSMUSG00000092837 | *Rpph1* | 2.95E-04 | 2.025336 |
| ENSMUSG00000040340 | *Tex45* | 2.80E-03 | 2.026678 |
| ENSMUSG00000086754 | *Gm16098* | 3.57E-02 | 2.033401 |
| ENSMUSG00000088025 | *Rprl3* | 1.21E-03 | 2.034205 |
| ENSMUSG00000116031 | *D030024E09Rik* | 3.10E-10 | 2.034724 |
| ENSMUSG00000079243 | *Xirp1* | 1.63E-25 | 2.041369 |
| ENSMUSG00000054905 | *Stfa3* | 2.46E-04 | 2.044116 |

| | | | |
|---|---|---|---|
| ENSMUSG00000032292 | *Nr2e3* | 1.06E-02 | 2.046336 |
| ENSMUSG00000089854 | *Gm16133* | 2.44E-06 | 2.047282 |
| ENSMUSG00000116606 | *AC163633.2* | 3.18E-02 | 2.047702 |
| ENSMUSG00000103382 | *Gm37755* | 2.66E-02 | 2.049867 |
| ENSMUSG00000077563 | *Snora68* | 1.19E-05 | 2.052854 |
| ENSMUSG00000106224 | *Gm43823* | 1.06E-02 | 2.054673 |
| ENSMUSG00000103308 | *Gm37800* | 1.61E-09 | 2.055329 |
| ENSMUSG00000115151 | *Gm19276* | 5.37E-03 | 2.055894 |
| ENSMUSG00000086649 | *Gm15286* | 3.26E-02 | 2.064189 |
| ENSMUSG00000020216 | *Jsrp1* | 1.65E-02 | 2.064977 |
| ENSMUSG00000097036 | *1110036E04Rik* | 2.16E-02 | 2.065813 |
| ENSMUSG00000089536 | *Scarna3a* | 2.33E-02 | 2.0728 |
| ENSMUSG00000113447 | *Gm48538* | 3.67E-05 | 2.078097 |
| ENSMUSG00000087477 | *Gm13822* | 1.26E-03 | 2.082278 |
| ENSMUSG00000041476 | *Smpx* | 3.41E-09 | 2.083676 |
| ENSMUSG00000065824 | *Gm26315* | 1.20E-05 | 2.084426 |
| ENSMUSG00000103485 | *Gm38001* | 4.48E-03 | 2.088313 |
| ENSMUSG00000109311 | *AI314278* | 8.28E-04 | 2.092515 |
| ENSMUSG00000065911 | *Gm24447* | 1.06E-05 | 2.092602 |
| ENSMUSG00000108985 | *A930030B08Rik* | 1.21E-03 | 2.092679 |
| ENSMUSG00000094088 | *Ighv1-64* | 4.84E-02 | 2.093497 |
| ENSMUSG00000093577 | *Gm20632* | 5.25E-03 | 2.099577 |
| ENSMUSG00000097468 | *Mdrl* | 4.60E-03 | 2.100337 |
| ENSMUSG00000103923 | *Gm37896* | 9.28E-03 | 2.102126 |
| ENSMUSG00000112471 | *Gm5779* | 5.98E-03 | 2.104841 |
| ENSMUSG00000093908 | *Gm5784* | 1.59E-06 | 2.105369 |
| ENSMUSG00000047501 | *Cldn4* | 3.24E-06 | 2.10583 |
| ENSMUSG00000108677 | *Gm44759* | 1.85E-02 | 2.11069 |
| ENSMUSG00000087775 | *Rprl2* | 7.23E-04 | 2.112469 |
| ENSMUSG00000033213 | *AA467197* | 7.56E-07 | 2.116442 |
| ENSMUSG00000117092 | *AC166110.1* | 4.57E-03 | 2.122036 |
| ENSMUSG00000110278 | *Gm5608* | 4.80E-19 | 2.124723 |
| ENSMUSG00000033196 | *Myh2* | 2.12E-09 | 2.129399 |
| ENSMUSG00000021123 | *Rdh12* | 1.23E-27 | 2.13007 |
| ENSMUSG00000111818 | *Gm17749* | 1.56E-11 | 2.130559 |
| ENSMUSG00000022026 | *Olfm4* | 1.74E-47 | 2.133503 |
| ENSMUSG00000097131 | *D230017M19Rik* | 8.10E-07 | 2.14415 |
| ENSMUSG00000113998 | *Gm40909* | 1.98E-16 | 2.150507 |
| ENSMUSG00000090101 | *Snhg9* | 6.89E-03 | 2.150509 |
| ENSMUSG00000074398 | *Gm15441* | 1.46E-03 | 2.151127 |
| ENSMUSG00000035653 | *Lrfn5* | 2.00E-04 | 2.153773 |
| ENSMUSG00000086220 | *Gm16599* | 1.86E-03 | 2.158383 |
| ENSMUSG00000116237 | *Gm30339* | 3.53E-03 | 2.162938 |
| ENSMUSG00000028865 | *Cd164l2* | 1.12E-04 | 2.164257 |
| ENSMUSG00000114457 | *Gm36346* | 3.31E-02 | 2.164377 |
| ENSMUSG00000064604 | *Snora44* | 5.26E-04 | 2.16931 |
| ENSMUSG00000084708 | *Gm22988* | 4.87E-07 | 2.171425 |
| ENSMUSG00000089542 | *Gm25835* | 1.60E-09 | 2.171658 |
| ENSMUSG00000024907 | *Gal* | 4.18E-02 | 2.171949 |
| ENSMUSG00000103620 | *Gm37359* | 3.97E-07 | 2.173343 |
| ENSMUSG00000093355 | *Snora26* | 4.17E-02 | 2.191494 |
| ENSMUSG00000043468 | *Adam30* | 1.06E-02 | 2.195772 |
| ENSMUSG00000101845 | *Gm28198* | 4.01E-02 | 2.207421 |

| | | | |
|---|---|---|---|
| ENSMUSG00000064380 | *Gm26448* | 9.55E-10 | 2.208699 |
| ENSMUSG00000025431 | *Crisp1* | 6.59E-04 | 2.208969 |
| ENSMUSG00000111867 | *Gm31013* | 2.68E-02 | 2.214693 |
| ENSMUSG00000087819 | *Gm25117* | 1.67E-07 | 2.214707 |
| ENSMUSG00000091366 | *Gm17040* | 4.44E-02 | 2.216824 |
| ENSMUSG00000064694 | *Gm24146* | 4.82E-04 | 2.217621 |
| ENSMUSG00000009551 | *6330409D20Rik* | 4.87E-02 | 2.221007 |
| ENSMUSG00000096351 | *Samd11* | 6.92E-09 | 2.221597 |
| ENSMUSG00000080365 | *Gm25776* | 1.17E-04 | 2.222028 |
| ENSMUSG00000065259 | *Snora30* | 5.73E-03 | 2.223572 |
| ENSMUSG00000107841 | *Gm44185* | 1.38E-02 | 2.224563 |
| ENSMUSG00000046961 | *Gpr156* | 4.17E-11 | 2.227765 |
| ENSMUSG00000026592 | *Tex35* | 1.15E-09 | 2.233732 |
| ENSMUSG00000103502 | *9330121J05Rik* | 3.03E-02 | 2.239685 |
| ENSMUSG00000086354 | *Gm13938* | 8.15E-26 | 2.24405 |
| ENSMUSG00000114697 | *Gm30806* | 2.67E-02 | 2.246899 |
| ENSMUSG00000004668 | *Abca13* | 2.61E-57 | 2.24728 |
| ENSMUSG00000085573 | *Gm15418* | 3.19E-04 | 2.250028 |
| ENSMUSG00000092819 | *Gm23639* | 1.25E-02 | 2.258412 |
| ENSMUSG00000104863 | *Gm49347* | 6.55E-05 | 2.266581 |
| ENSMUSG00000096887 | *Gm20594* | 2.13E-12 | 2.268158 |
| ENSMUSG00000113868 | *Gm34719* | 1.46E-03 | 2.270756 |
| ENSMUSG00000096243 | *Gm24265* | 3.92E-05 | 2.270994 |
| ENSMUSG00000085599 | *Gm13449* | 3.60E-04 | 2.273353 |
| ENSMUSG00000020848 | *Doc2b* | 4.53E-24 | 2.275294 |
| ENSMUSG00000088948 | *Gm23262* | 4.47E-07 | 2.276029 |
| ENSMUSG00000092702 | *Gm24514* | 2.89E-06 | 2.276899 |
| ENSMUSG00000109921 | *Gm33326* | 2.14E-03 | 2.281901 |
| ENSMUSG00000111765 | *Gm10635* | 3.03E-03 | 2.28596 |
| ENSMUSG00000053615 | *Gm9913* | 3.32E-02 | 2.288715 |
| ENSMUSG00000050830 | *Vwc2* | 1.21E-05 | 2.28941 |
| ENSMUSG00000045027 | *Prss22* | 4.65E-14 | 2.295806 |
| ENSMUSG00000059108 | *Ifitm6* | 3.62E-40 | 2.296863 |
| ENSMUSG00000111639 | *1700019L13Rik* | 1.07E-02 | 2.297301 |
| ENSMUSG00000006764 | *Tph2* | 1.53E-02 | 2.300399 |
| ENSMUSG00000099760 | *Gm28800* | 5.16E-04 | 2.300754 |
| ENSMUSG00000108171 | *Gm43915* | 1.82E-02 | 2.304507 |
| ENSMUSG00000051965 | *Nanos2* | 3.15E-02 | 2.309161 |
| ENSMUSG00000065649 | *Snora74a* | 1.83E-06 | 2.319528 |
| ENSMUSG00000111821 | *Gm48545* | 1.45E-05 | 2.341304 |
| ENSMUSG00000074652 | *Myh7b* | 2.41E-03 | 2.354961 |
| ENSMUSG00000023153 | *Tmem52* | 3.20E-03 | 2.37334 |
| ENSMUSG00000113778 | *Gm40847* | 2.82E-06 | 2.385725 |
| ENSMUSG00000102189 | *Gm37194* | 2.59E-09 | 2.389093 |
| ENSMUSG00000091238 | *Gm17103* | 1.31E-04 | 2.395524 |
| ENSMUSG00000110237 | *Gm18258* | 2.84E-02 | 2.417341 |
| ENSMUSG00000088252 | *Snord13* | 5.28E-05 | 2.418714 |
| ENSMUSG00000098975 | *Gm27177* | 1.24E-02 | 2.419022 |
| ENSMUSG00000097346 | *Gm26619* | 8.29E-03 | 2.422694 |
| ENSMUSG00000054582 | *Pabpc1l* | 8.18E-19 | 2.43202 |
| ENSMUSG00000094377 | *Gm24407* | 5.41E-07 | 2.44395 |
| ENSMUSG00000090625 | *Gm20721* | 5.29E-04 | 2.452874 |
| ENSMUSG00000010342 | *Tex14* | 2.41E-09 | 2.454352 |

| | | | |
|---|---|---|---|
| ENSMUSG00000033860 | *Fgg* | 1.06E-03 | 2.454493 |
| ENSMUSG00000043460 | *Elfn2* | 1.29E-02 | 2.455388 |
| ENSMUSG00000110114 | *Gm45471* | 1.99E-02 | 2.456678 |
| ENSMUSG00000064945 | *Rny3* | 2.32E-06 | 2.465394 |
| ENSMUSG00000085297 | *Gm11651* | 3.64E-05 | 2.473195 |
| ENSMUSG00000089655 | *Gm16004* | 2.40E-03 | 2.475649 |
| ENSMUSG00000108720 | *Gm44672* | 7.98E-03 | 2.479163 |
| ENSMUSG00000000157 | *Itgb2l* | 5.28E-41 | 2.481365 |
| ENSMUSG00000086980 | *Gm13791* | 1.78E-03 | 2.484391 |
| ENSMUSG00000024729 | *1700017D01Rik* | 1.57E-02 | 2.495512 |
| ENSMUSG00000069310 | *Hist1h3c* | 5.35E-04 | 2.501758 |
| ENSMUSG00000065087 | *Snord22* | 1.02E-20 | 2.504791 |
| ENSMUSG00000077254 | *Gm26079* | 5.09E-05 | 2.508432 |
| ENSMUSG00000023467 | *Tulp2* | 3.92E-03 | 2.545698 |
| ENSMUSG00000084744 | *Gm25291* | 1.27E-04 | 2.547348 |
| ENSMUSG00000080538 | *Gm25541* | 1.65E-07 | 2.547886 |
| ENSMUSG00000086628 | *Gm16157* | 1.31E-03 | 2.549909 |
| ENSMUSG00000116657 | *AC127341.3* | 6.32E-03 | 2.552284 |
| ENSMUSG00000085532 | *B430319H21Rik* | 2.25E-10 | 2.571664 |
| ENSMUSG00000095590 | *Gm24305* | 5.01E-03 | 2.571674 |
| ENSMUSG00000040258 | *Nxph4* | 5.24E-03 | 2.575445 |
| ENSMUSG00000097069 | *Gm16998* | 1.82E-02 | 2.578575 |
| ENSMUSG00000010362 | *Rdm1* | 1.63E-08 | 2.583826 |
| ENSMUSG00000107179 | *Gm43069* | 7.95E-03 | 2.585741 |
| ENSMUSG00000064966 | *Snord15b* | 2.43E-08 | 2.605397 |
| ENSMUSG00000066510 | *Ankdd1a* | 1.64E-20 | 2.613295 |
| ENSMUSG00000031343 | *Gabra3* | 2.60E-11 | 2.613416 |
| ENSMUSG00000096084 | *Gm21850* | 1.82E-14 | 2.614388 |
| ENSMUSG00000090208 | *Gm15851* | 1.89E-02 | 2.614614 |
| ENSMUSG00000074113 | *Gm10629* | 1.77E-02 | 2.621575 |
| ENSMUSG00000064901 | *Snora21* | 1.20E-03 | 2.626929 |
| ENSMUSG00000090293 | *Gm17034* | 1.90E-04 | 2.681395 |
| ENSMUSG00000037942 | *Crp* | 7.71E-04 | 2.683734 |
| ENSMUSG00000111582 | *Gm47347* | 3.20E-03 | 2.685612 |
| ENSMUSG00000085511 | *Gm11738* | 1.89E-04 | 2.70767 |
| ENSMUSG00000064371 | *mt-Tt* | 2.94E-03 | 2.714465 |
| ENSMUSG00000114649 | *3110006O06Rik* | 8.08E-03 | 2.716208 |
| ENSMUSG00000065822 | *Snord15a* | 1.62E-05 | 2.722509 |
| ENSMUSG00000085806 | *Gm12023* | 8.06E-11 | 2.725972 |
| ENSMUSG00000021071 | *Trim9* | 6.06E-15 | 2.732965 |
| ENSMUSG00000025172 | *Ankrd2* | 4.58E-16 | 2.747592 |
| ENSMUSG00000064899 | *Snord118* | 3.85E-04 | 2.765016 |
| ENSMUSG00000110790 | *Gm47079* | 8.78E-03 | 2.791718 |
| ENSMUSG00000016327 | *Atp1b4* | 7.76E-04 | 2.794331 |
| ENSMUSG00000096960 | *A230028O05Rik* | 2.22E-22 | 2.835285 |
| ENSMUSG00000089255 | *Snora78* | 7.85E-04 | 2.851741 |
| ENSMUSG00000032484 | *Ngp* | 3.65E-107 | 2.860267 |
| ENSMUSG00000064943 | *Gm23240* | 8.10E-03 | 2.878259 |
| ENSMUSG00000114922 | *Ppifos* | 1.68E-05 | 2.879757 |
| ENSMUSG00000022249 | *Ttc23l* | 3.38E-03 | 2.910968 |
| ENSMUSG00000030359 | *Pzp* | 1.62E-03 | 2.922478 |
| ENSMUSG00000065118 | *Gm23297* | 4.76E-05 | 2.968259 |

| | | | |
|---|---|---|---|
| ENSMUSG00000111734 | *Gm29825* | 3.88E-03 | 2.997991 |
| ENSMUSG00000115532 | *Gm41206* | 2.62E-03 | 3.003723 |
| ENSMUSG00000106056 | *4930500L23Rik* | 2.55E-03 | 3.004773 |
| ENSMUSG00000093843 | *Gm25939* | 6.49E-07 | 3.015148 |
| ENSMUSG00000065176 | *Rnu12* | 9.49E-18 | 3.015627 |
| ENSMUSG00000094050 | *Gm23472* | 2.00E-08 | 3.0215 |
| ENSMUSG00000064856 | *Gm23444* | 2.61E-08 | 3.025159 |
| ENSMUSG00000093956 | *Gm24497* | 3.05E-08 | 3.03474 |
| ENSMUSG00000059908 | *Mug1* | 4.53E-02 | 3.062787 |
| ENSMUSG00000064994 | *Gm22422* | 7.84E-06 | 3.063734 |
| ENSMUSG00000109464 | *Gm34350* | 1.03E-02 | 3.073399 |
| ENSMUSG00000042761 | *Mrap2* | 4.40E-06 | 3.081864 |
| ENSMUSG00000089948 | *Far2os1* | 4.72E-04 | 3.104791 |
| ENSMUSG00000065944 | *Rnu2-10* | 7.80E-09 | 3.107609 |
| ENSMUSG00000064702 | *Gm24950* | 9.28E-09 | 3.114592 |
| ENSMUSG00000065251 | *Gm23971* | 9.28E-09 | 3.114592 |
| ENSMUSG00000065767 | *Gm23849* | 7.91E-09 | 3.116902 |
| ENSMUSG00000087038 | *2900079G21Rik* | 4.45E-03 | 3.117026 |
| ENSMUSG00000091993 | *B930036N10Rik* | 9.73E-07 | 3.120615 |
| ENSMUSG00000022868 | *Ahsg* | 3.12E-04 | 3.161068 |
| ENSMUSG00000095892 | *Rnu5g* | 3.38E-09 | 3.162782 |
| ENSMUSG00000065820 | *Gm26316* | 3.08E-09 | 3.189586 |
| ENSMUSG00000046748 | *Tmem45a2* | 1.25E-18 | 3.232248 |
| ENSMUSG00000097554 | *Gm26825* | 1.50E-03 | 3.233552 |
| ENSMUSG00000095738 | *Gm25313* | 4.08E-09 | 3.26894 |
| ENSMUSG00000110735 | *Gm47173* | 6.45E-03 | 3.273965 |
| ENSMUSG00000099291 | *Rnu3b3* | 2.67E-15 | 3.306299 |
| ENSMUSG00000104856 | *Rnu3b3* | 2.67E-15 | 3.306299 |
| ENSMUSG00000105115 | *Rnu3b2* | 5.42E-16 | 3.31235 |
| ENSMUSG00000098925 | *Rnu3b2* | 5.42E-16 | 3.31235 |
| ENSMUSG00000104896 | *Rnu3b4* | 7.98E-16 | 3.316405 |
| ENSMUSG00000098641 | *Rnu3b4* | 8.89E-16 | 3.316683 |
| ENSMUSG00000027233 | *Patl2* | 9.28E-06 | 3.317931 |
| ENSMUSG00000098943 | *Rnu3b1* | 9.63E-16 | 3.32264 |
| ENSMUSG00000105025 | *Rnu3b1* | 9.63E-16 | 3.32264 |
| ENSMUSG00000085348 | *Myhas* | 2.21E-09 | 3.342376 |
| ENSMUSG00000065254 | *Gm23973* | 3.63E-04 | 3.375179 |
| ENSMUSG00000088675 | *Rprl1* | 9.25E-05 | 3.390244 |
| ENSMUSG00000106222 | *Rprl1* | 9.25E-05 | 3.390244 |
| ENSMUSG00000086777 | *Far2os2* | 8.53E-09 | 3.410703 |
| ENSMUSG00000013936 | *Myl2* | 1.38E-11 | 3.412107 |
| ENSMUSG00000065870 | *Rnu3a* | 8.20E-11 | 3.438606 |
| ENSMUSG00000106147 | *Rnu3a* | 8.20E-11 | 3.438606 |
| ENSMUSG00000094655 | *Gm25360* | 9.25E-18 | 3.476566 |
| ENSMUSG00000099449 | *Gm28401* | 3.66E-06 | 3.512495 |
| ENSMUSG00000080465 | *Gm22486* | 3.79E-11 | 3.525781 |
| ENSMUSG00000097770 | *Gm26776* | 7.15E-05 | 3.556066 |
| ENSMUSG00000095676 | *Gm25099* | 1.48E-18 | 3.569557 |
| ENSMUSG00000038357 | *Camp* | 2.10E-20 | 3.574791 |
| ENSMUSG00000110588 | *Gm45774* | 2.37E-05 | 3.651499 |
| ENSMUSG00000105361 | *AY036118* | 4.75E-13 | 3.706614 |
| ENSMUSG00000114163 | *Gm32296* | 1.06E-05 | 3.753819 |
| ENSMUSG00000097815 | *Gm26809* | 4.14E-06 | 3.770845 |

| | | | |
|---|---|---|---|
| ENSMUSG00000101133 | *Gm29050* | 4.66E-06 | 3.935451 |
| ENSMUSG00000094668 | *Gm24871* | 3.50E-05 | 3.990369 |
| ENSMUSG00000065232 | *Gm22973* | 1.47E-13 | 4.080121 |
| ENSMUSG00000096269 | *Rpl31-ps21* | 1.07E-04 | 4.321736 |
| ENSMUSG00000029019 | *Nppb* | 3.13E-19 | 4.33994 |
| ENSMUSG00000097364 | *Gm26719* | 2.54E-09 | 4.440093 |
| ENSMUSG00000056656 | *Apol8* | 1.21E-16 | 4.471575 |
| ENSMUSG00000064682 | *Gm25813* | 4.91E-11 | 4.908819 |
| ENSMUSG00000115207 | *Gm49132* | 2.67E-04 | 5.036502 |
| ENSMUSG00000028307 | *Aldob* | 2.14E-03 | 5.215422 |
| ENSMUSG00000027261 | *Hao1* | 2.14E-10 | 5.270442 |
| ENSMUSG00000056035 | *Cyp3a11* | 1.02E-03 | 8.674899 |
| ENSMUSG00000025991 | *Cps1* | 9.04E-03 | 9.941545 |
| ENSMUSG00000032083 | *Apoa1* | 5.02E-04 | 10.21736 |
| ENSMUSG00000029368 | *Alb* | 1.23E-11 | 11.41724 |
| ENSMUSG00000033831 | *Fgb* | 1.09E-04 | 16.81357 |

**Supplementary Table 12.** Murine differential gene expression in the lungs 6 h post-infection, 4559-Blood vs 4559M. Genes with fold change (FC) greater than 1.5 and $p < 0.05$ are shown. FC values highlighted in blue = upregulated in 4559-Blood, while values highlighted in red = upregulated in 4559M.

| **Gene Stable ID** | **Gene Name** | **padj** | **FC** |
|---|---|---|---|
| ENSMUSG00000029630 | *Cyp3a25* | 1.75E-05 | 49.7977 |
| ENSMUSG00000059908 | *Mug1* | 3.28E-16 | 16.2463 |
| ENSMUSG00000061808 | *Ttr* | 2.84E-06 | 14.48 |
| ENSMUSG00000032083 | *Apoa1* | 1.40E-09 | 8.40944 |
| ENSMUSG00000029368 | *Alb* | 7.42E-10 | 7.88269 |
| ENSMUSG00000056035 | *Cyp3a11* | 2.46E-09 | 7.04516 |
| ENSMUSG00000003053 | *Cyp2c29* | 1.29E-06 | 6.94961 |
| ENSMUSG00000025991 | *Cps1* | 1.02E-02 | 6.84018 |
| ENSMUSG00000059481 | *Plg* | 5.17E-05 | 6.64669 |
| ENSMUSG00000035540 | *Gc* | 4.99E-09 | 6.39712 |
| ENSMUSG00000019932 | *Kera* | 1.93E-07 | 5.71283 |
| ENSMUSG00000058354 | *Krt6a* | 2.39E-09 | 5.47428 |
| ENSMUSG00000020609 | *Apob* | 7.58E-09 | 5.28553 |
| ENSMUSG00000048455 | *Sprr1b* | 8.94E-08 | 5.28178 |
| ENSMUSG00000030131 | *Mug2* | 7.90E-05 | 4.9728 |
| ENSMUSG00000030359 | *Pzp* | 5.04E-10 | 4.9175 |
| ENSMUSG00000074647 | *Fam83c* | 3.54E-05 | 4.88673 |
| ENSMUSG00000033831 | *Fgb* | 8.00E-03 | 4.77839 |
| ENSMUSG00000023041 | *Krt6b* | 6.28E-04 | 4.23661 |
| ENSMUSG00000046095 | *Krt32* | 6.56E-05 | 4.06888 |
| ENSMUSG00000022875 | *Kng1* | 1.83E-04 | 4.04028 |
| ENSMUSG00000022868 | *Ahsg* | 6.92E-08 | 4.01269 |
| ENSMUSG00000028307 | *Aldob* | 4.31E-05 | 3.99588 |
| ENSMUSG00000068888 | *Lce1i* | 1.87E-03 | 3.93124 |
| ENSMUSG00000035861 | *Tmprss11b* | 9.18E-04 | 3.92883 |

| | | | |
|---|---|---|---|
| ENSMUSG00000025194 | *Abcc2* | 8.61E-04 | 3.90941 |
| ENSMUSG00000027761 | *Aadac* | 1.07E-03 | 3.88939 |
| ENSMUSG00000092586 | *Ly6g6c* | 2.26E-12 | 3.86378 |
| ENSMUSG00000074445 | *Sprr2a3* | 1.99E-05 | 3.71115 |
| ENSMUSG00000059898 | *Dsc3* | 2.90E-17 | 3.70907 |
| ENSMUSG00000054422 | *Fabp1* | 3.27E-04 | 3.66856 |
| ENSMUSG00000053675 | *Tgm5* | 1.06E-02 | 3.59152 |
| ENSMUSG00000032373 | *Car12* | 2.24E-43 | 3.56925 |
| ENSMUSG00000021364 | *Elovl2* | 3.03E-03 | 3.52501 |
| ENSMUSG00000026450 | *Chit1* | 1.24E-06 | 3.51671 |
| ENSMUSG00000033268 | *Duox1* | 7.07E-12 | 3.43887 |
| ENSMUSG00000056632 | *Dsg3* | 8.74E-24 | 3.40573 |
| ENSMUSG00000067006 | *Serpinb5* | 8.21E-12 | 3.40296 |
| ENSMUSG00000017950 | *Hnf4a* | 3.10E-03 | 3.39187 |
| ENSMUSG00000040127 | *Sdr9c7* | 3.52E-03 | 3.33689 |
| ENSMUSG00000045545 | *Krt14* | 1.09E-22 | 3.26379 |
| ENSMUSG00000061527 | *Krt5* | 1.31E-17 | 3.25295 |
| ENSMUSG00000097850 | *4631405K08Rik* | 3.80E-04 | 3.17772 |
| ENSMUSG00000026985 | *Il1f8* | 1.88E-04 | 3.17313 |
| ENSMUSG00000078664 | *Sprr2a1* | 4.06E-05 | 3.14672 |
| ENSMUSG00000054146 | *Krt15* | 8.77E-17 | 3.1094 |
| ENSMUSG00000028236 | *Sdr16c5* | 5.99E-04 | 3.09328 |
| ENSMUSG00000026413 | *Pkp1* | 2.42E-19 | 3.08859 |
| ENSMUSG00000102098 | *2310016D03Rik* | 2.45E-03 | 3.03012 |
| ENSMUSG00000059832 | *Kprp* | 2.49E-18 | 3.02836 |
| ENSMUSG00000068893 | *Sprr2a2* | 5.63E-11 | 3.01925 |
| ENSMUSG00000031757 | *Mt4* | 2.82E-09 | 3.00348 |
| ENSMUSG00000038295 | *Atg9b* | 3.71E-08 | 3.00088 |
| ENSMUSG00000097768 | *2310043M15Rik* | 6.14E-05 | 2.99467 |
| ENSMUSG00000048399 | *Tprg* | 8.10E-08 | 2.97043 |
| ENSMUSG00000071858 | *Gm94* | 8.27E-08 | 2.95766 |
| ENSMUSG00000042031 | *Lce3b* | 8.70E-10 | 2.94911 |
| ENSMUSG00000085261 | *Gm13814* | 1.65E-02 | 2.93578 |
| ENSMUSG00000075217 | *4833423E24Rik* | 3.24E-08 | 2.93359 |
| ENSMUSG00000041984 | *Rptn* | 2.63E-20 | 2.90552 |
| ENSMUSG00000033765 | *Calm4* | 4.66E-07 | 2.90141 |
| ENSMUSG00000005355 | *Casp14* | 1.53E-03 | 2.89567 |
| ENSMUSG00000026327 | *Serpinb11* | 7.21E-07 | 2.86034 |
| ENSMUSG00000044748 | *Defb1* | 4.03E-04 | 2.84373 |
| ENSMUSG00000079025 | *Gsdmc* | 6.93E-06 | 2.8424 |
| ENSMUSG00000053522 | *Lgals7* | 1.41E-07 | 2.82395 |
| ENSMUSG00000043472 | *Lce3d* | 5.15E-03 | 2.80644 |
| ENSMUSG00000045539 | *Sprr3* | 1.03E-15 | 2.79819 |
| ENSMUSG00000018862 | *Otop3* | 4.58E-06 | 2.79571 |
| ENSMUSG00000114091 | *Gm40655* | 1.03E-06 | 2.79416 |
| ENSMUSG00000071019 | *Sdr16c6* | 2.08E-04 | 2.79341 |
| ENSMUSG00000099906 | *Gm28653* | 9.77E-04 | 2.78803 |
| ENSMUSG00000004872 | *Pax3* | 6.22E-03 | 2.78741 |
| ENSMUSG00000045019 | *Acer1* | 5.37E-03 | 2.78363 |
| ENSMUSG00000057609 | *Lce1a1* | 1.99E-04 | 2.77319 |
| ENSMUSG00000056078 | *Lipm* | 6.48E-07 | 2.76931 |
| ENSMUSG00000033860 | *Fgg* | 1.23E-05 | 2.7686 |
| ENSMUSG00000047428 | *Dlk2* | 1.85E-02 | 2.76136 |

| | | | |
|---|---|---|---|
| ENSMUSG00000025064 | *Col17a1* | 6.87E-17 | 2.75765 |
| ENSMUSG00000034774 | *Dsg1c* | 3.15E-03 | 2.75484 |
| ENSMUSG00000017607 | *Tns4* | 1.32E-09 | 2.74798 |
| ENSMUSG00000030484 | *Lypd5* | 7.90E-11 | 2.74454 |
| ENSMUSG00000074156 | *Ces1h* | 1.86E-07 | 2.73844 |
| ENSMUSG00000026983 | *Il1f5* | 2.08E-04 | 2.73336 |
| ENSMUSG00000059668 | *Krt4* | 1.06E-11 | 2.72465 |
| ENSMUSG00000050359 | *Sprr1a* | 2.66E-11 | 2.71649 |
| ENSMUSG00000105160 | *A530030E21Rik* | 3.10E-02 | 2.71344 |
| ENSMUSG00000050808 | *Muc15* | 3.46E-03 | 2.71087 |
| ENSMUSG00000050463 | *Krt78* | 1.01E-21 | 2.70916 |
| ENSMUSG00000029154 | *Cwh43* | 3.03E-03 | 2.6846 |
| ENSMUSG00000044041 | *Krt13* | 1.85E-11 | 2.68386 |
| ENSMUSG00000103243 | *Lce1d* | 2.22E-04 | 2.66703 |
| ENSMUSG00000078964 | *Ces1b* | 5.60E-04 | 2.66571 |
| ENSMUSG00000070719 | *Pla2g4d* | 4.58E-03 | 2.66266 |
| ENSMUSG00000027224 | *Duoxa1* | 1.53E-02 | 2.65474 |
| ENSMUSG00000062826 | *Ces2f* | 5.60E-13 | 2.64193 |
| ENSMUSG00000074199 | *Krtdap* | 6.33E-07 | 2.64017 |
| ENSMUSG00000050108 | *Bpifc* | 2.11E-03 | 2.63956 |
| ENSMUSG00000055561 | *Spink5* | 2.42E-19 | 2.63531 |
| ENSMUSG00000063651 | *Cnfn* | 1.48E-13 | 2.63424 |
| ENSMUSG00000042124 | *Lce1f* | 4.49E-03 | 2.6302 |
| ENSMUSG00000044594 | *Serpinb3a* | 3.67E-09 | 2.62271 |
| ENSMUSG00000028001 | *Fga* | 6.07E-04 | 2.61124 |
| ENSMUSG00000074489 | *Bglap3* | 1.19E-04 | 2.60397 |
| ENSMUSG00000017204 | *Gsdma* | 1.43E-07 | 2.603 |
| ENSMUSG00000033196 | *Myh2* | 4.95E-14 | 2.58757 |
| ENSMUSG00000030834 | *Abcc6* | 1.07E-03 | 2.57913 |
| ENSMUSG00000081163 | *Defa-ps12* | 6.10E-03 | 2.57856 |
| ENSMUSG00000034634 | *Ly6d* | 1.25E-08 | 2.56812 |
| ENSMUSG00000061928 | *Dsg1b* | 3.62E-11 | 2.56262 |
| ENSMUSG00000106871 | *Gm3289* | 3.48E-02 | 2.55403 |
| ENSMUSG00000022491 | *Glycam1* | 5.93E-08 | 2.552 |
| ENSMUSG00000057454 | *Lypd3* | 9.96E-06 | 2.54396 |
| ENSMUSG00000084228 | *Gm8080* | 3.66E-02 | 2.53954 |
| ENSMUSG00000061259 | *Tmprss11d* | 1.47E-04 | 2.53596 |
| ENSMUSG00000027376 | *Prom2* | 7.87E-09 | 2.53276 |
| ENSMUSG00000022860 | *Chodl* | 3.30E-03 | 2.53009 |
| ENSMUSG00000036960 | *Clca2* | 3.16E-04 | 2.52337 |
| ENSMUSG00000045475 | *Lce3c* | 9.57E-08 | 2.52299 |
| ENSMUSG00000063130 | *Calml3* | 1.43E-10 | 2.51428 |
| ENSMUSG00000040154 | *Wfdc5* | 1.42E-04 | 2.51022 |
| ENSMUSG00000117222 | *AC060761.1* | 1.75E-10 | 2.50439 |
| ENSMUSG00000059956 | *Serpinb12* | 1.32E-07 | 2.48356 |
| ENSMUSG00000021506 | *Pitx1* | 2.41E-04 | 2.46822 |
| ENSMUSG00000043029 | *Trpv3* | 6.01E-03 | 2.45237 |
| ENSMUSG00000034362 | *Csta1* | 4.02E-03 | 2.44214 |
| ENSMUSG00000099146 | *0610031O16Rik* | 1.31E-02 | 2.42834 |
| ENSMUSG00000044499 | *Hs3st5* | 4.61E-14 | 2.42328 |
| ENSMUSG00000116659 | *AC158985.1* | 1.31E-02 | 2.41132 |
| ENSMUSG00000027401 | *Tgm3* | 1.67E-09 | 2.41084 |
| ENSMUSG00000037188 | *Grhl3* | 1.79E-08 | 2.41062 |

| | | | |
|---|---|---|---|
| ENSMUSG00000093385 | *A330044P14Rik* | 4.33E-02 | 2.40792 |
| ENSMUSG00000030713 | *Klk7* | 9.32E-03 | 2.40587 |
| ENSMUSG00000043430 | *Psapl1* | 1.18E-08 | 2.40429 |
| ENSMUSG00000044628 | *Rnf208* | 3.62E-02 | 2.40152 |
| ENSMUSG00000039238 | *Zfp750* | 2.57E-05 | 2.39905 |
| ENSMUSG00000037129 | *Tmprss13* | 4.41E-07 | 2.39884 |
| ENSMUSG00000032013 | *Trim29* | 1.24E-12 | 2.39013 |
| ENSMUSG00000109656 | *Gm45548* | 3.25E-02 | 2.38688 |
| ENSMUSG00000051431 | *Gpr87* | 1.20E-02 | 2.38474 |
| ENSMUSG00000044322 | *Dsc1* | 7.89E-05 | 2.38231 |
| ENSMUSG00000069441 | *Dsg1a* | 1.65E-12 | 2.37371 |
| ENSMUSG00000032807 | *Alox12b* | 8.10E-07 | 2.37178 |
| ENSMUSG00000039269 | *2300002M23Rik* | 6.30E-04 | 2.36131 |
| ENSMUSG00000055301 | *Adh7* | 2.74E-13 | 2.352 |
| ENSMUSG00000022510 | *Trp63* | 3.19E-20 | 2.34287 |
| ENSMUSG00000092222 | *Gm20506* | 3.27E-02 | 2.34271 |
| ENSMUSG00000038020 | *Rapgefl1* | 1.54E-11 | 2.33861 |
| ENSMUSG00000028148 | *Them5* | 1.43E-08 | 2.32907 |
| ENSMUSG00000032454 | *Rbp2* | 2.18E-02 | 2.32839 |
| ENSMUSG00000054325 | *Lce3a* | 1.60E-06 | 2.31499 |
| ENSMUSG00000016327 | *Atp1b4* | 9.25E-04 | 2.30528 |
| ENSMUSG00000047884 | *Klk9* | 1.51E-02 | 2.29768 |
| ENSMUSG00000041991 | *Hrnr* | 2.88E-07 | 2.29503 |
| ENSMUSG00000010064 | *Slc38a3* | 9.74E-11 | 2.28605 |
| ENSMUSG00000027913 | *Crct1* | 1.33E-08 | 2.2855 |
| ENSMUSG00000050296 | *Abca12* | 2.98E-09 | 2.27557 |
| ENSMUSG00000039518 | *Cdsn* | 3.75E-08 | 2.27368 |
| ENSMUSG00000079451 | *Tmprss11g* | 1.35E-04 | 2.2653 |
| ENSMUSG00000073602 | *Serpinb3b* | 7.22E-05 | 2.25697 |
| ENSMUSG00000027513 | *Pck1* | 7.41E-18 | 2.24387 |
| ENSMUSG00000043681 | *Fam25c* | 2.27E-04 | 2.24042 |
| ENSMUSG00000047586 | *Nccrp1* | 1.89E-02 | 2.24009 |
| ENSMUSG00000026984 | *Il1f6* | 6.03E-03 | 2.23995 |
| ENSMUSG00000039070 | *Cpa4* | 3.31E-06 | 2.23183 |
| ENSMUSG00000023176 | *Cpn2* | 3.39E-02 | 2.23093 |
| ENSMUSG00000060621 | *Nkpd1* | 1.11E-02 | 2.2285 |
| ENSMUSG00000048764 | *Tmprss11f* | 1.97E-04 | 2.22505 |
| ENSMUSG00000076613 | *Ighg2b* | 8.46E-03 | 2.22301 |
| ENSMUSG00000025329 | *Padi1* | 1.34E-02 | 2.22281 |
| ENSMUSG00000044359 | *P2ry4* | 2.00E-03 | 2.21873 |
| ENSMUSG00000079644 | *Gm1110* | 8.55E-04 | 2.19573 |
| ENSMUSG00000059230 | *Defb4* | 1.15E-06 | 2.17928 |
| ENSMUSG00000112267 | *Gm47903* | 3.79E-02 | 2.17842 |
| ENSMUSG00000021359 | *Tfap2a* | 8.96E-04 | 2.17337 |
| ENSMUSG00000031844 | *Hsd17b2* | 1.45E-04 | 2.16161 |
| ENSMUSG00000067714 | *Lpar5* | 2.70E-02 | 2.15751 |
| ENSMUSG00000068885 | *Lce3f* | 5.02E-05 | 2.15408 |
| ENSMUSG00000031372 | *Trex2* | 1.77E-02 | 2.15218 |
| ENSMUSG00000073601 | *Serpinb3c* | 7.40E-03 | 2.14657 |
| ENSMUSG00000024331 | *Dsc2* | 6.87E-11 | 2.14446 |
| ENSMUSG00000090356 | *Teddm3* | 1.75E-03 | 2.13866 |
| ENSMUSG00000032281 | *Acsbg1* | 5.63E-12 | 2.13831 |
| ENSMUSG00000021509 | *Slc25a48* | 5.40E-03 | 2.13414 |

| | | | |
|---|---|---|---|
| ENSMUSG00000086448 | *9330162012Rik* | 1.57E-02 | 2.12691 |
| ENSMUSG00000047787 | *Flrt1* | 1.70E-02 | 2.12205 |
| ENSMUSG00000068890 | *Lce1a2* | 2.23E-02 | 2.12041 |
| ENSMUSG00000046818 | *Ddit4l* | 1.82E-04 | 2.11443 |
| ENSMUSG00000045776 | *Lrtm1* | 3.90E-10 | 2.10474 |
| ENSMUSG00000038242 | *Aox4* | 1.01E-05 | 2.10471 |
| ENSMUSG00000027923 | *Lce1b* | 2.48E-02 | 2.10373 |
| ENSMUSG00000055333 | *Fat2* | 2.95E-09 | 2.09985 |
| ENSMUSG00000091376 | *Aadacl2* | 4.12E-02 | 2.09943 |
| ENSMUSG00000086585 | *Gm16126* | 2.80E-02 | 2.08292 |
| ENSMUSG00000074625 | *Arhgap40* | 4.55E-07 | 2.08272 |
| ENSMUSG00000025431 | *Crisp1* | 3.94E-04 | 2.0827 |
| ENSMUSG00000003271 | *Sult2b1* | 1.40E-04 | 2.07941 |
| ENSMUSG00000074771 | *Ankef1* | 1.86E-07 | 2.06756 |
| ENSMUSG00000030577 | *Cd22* | 9.58E-12 | 2.0664 |
| ENSMUSG00000042092 | *Lce1c* | 2.56E-02 | 2.06455 |
| ENSMUSG00000075296 | *Aldh3b2* | 3.22E-03 | 2.06241 |
| ENSMUSG00000048078 | *Tenm4* | 4.36E-46 | 2.05431 |
| ENSMUSG00000043165 | *Lor* | 1.66E-12 | 2.053 |
| ENSMUSG00000046056 | *Sbsn* | 6.79E-08 | 2.04981 |
| ENSMUSG00000049336 | *Tenm2* | 1.73E-20 | 2.03138 |
| ENSMUSG00000020061 | *Mybpc1* | 1.29E-26 | 2.0298 |
| ENSMUSG00000038354 | *Ankrd35* | 5.87E-04 | 2.02604 |
| ENSMUSG00000021255 | *Esrrb* | 2.34E-05 | 2.00726 |
| ENSMUSG00000106245 | *Gm43824* | 2.26E-03 | 2.00651 |
| ENSMUSG00000026616 | *Cr2* | 8.19E-09 | 2.0019 |
| ENSMUSG00000087500 | *Gm12426* | 3.86E-02 | 1.99964 |
| ENSMUSG00000074433 | *Lce3e* | 3.42E-03 | 1.99487 |
| ENSMUSG00000086584 | *Gm12002* | 3.67E-05 | 1.99171 |
| ENSMUSG00000104927 | *Gm43388* | 4.85E-03 | 1.99016 |
| ENSMUSG00000033849 | *B3galt2* | 3.89E-03 | 1.98493 |
| ENSMUSG00000035606 | *Ky* | 1.89E-07 | 1.98363 |
| ENSMUSG00000100781 | *Gm29481* | 2.81E-02 | 1.98361 |
| ENSMUSG00000049598 | *Vsig8* | 2.47E-02 | 1.98235 |
| ENSMUSG00000028544 | *Slc5a9* | 3.70E-04 | 1.97676 |
| ENSMUSG00000036731 | *Cysrt1* | 3.63E-03 | 1.97512 |
| ENSMUSG00000038115 | *Ano2* | 1.53E-17 | 1.96639 |
| ENSMUSG00000027833 | *Shox2* | 3.91E-02 | 1.96557 |
| ENSMUSG00000009900 | *Wnt3a* | 1.55E-04 | 1.96506 |
| ENSMUSG00000022416 | *Cacna1i* | 1.15E-06 | 1.96482 |
| ENSMUSG00000032278 | *Paqr5* | 5.13E-09 | 1.96334 |
| ENSMUSG00000043461 | *Sptssb* | 5.50E-04 | 1.96325 |
| ENSMUSG00000026109 | *Tmeff2* | 5.04E-10 | 1.96288 |
| ENSMUSG00000024471 | *Myot* | 1.16E-18 | 1.95937 |
| ENSMUSG00000115919 | *Gm31583* | 2.72E-02 | 1.95861 |
| ENSMUSG00000037469 | *Acp7* | 1.65E-02 | 1.95823 |
| ENSMUSG00000056966 | *Gjc3* | 4.48E-02 | 1.95084 |
| ENSMUSG00000005716 | *Pvalb* | 1.10E-03 | 1.95002 |
| ENSMUSG00000055489 | *Ano5* | 1.63E-13 | 1.94962 |
| ENSMUSG00000043286 | *Pnpla1* | 6.54E-03 | 1.94448 |
| ENSMUSG00000015879 | *Fam184b* | 2.97E-03 | 1.94 |
| ENSMUSG00000001497 | *Pax9* | 7.64E-06 | 1.93926 |
| ENSMUSG00000021565 | *Slc6a19* | 2.86E-04 | 1.93609 |

| | | | |
|---|---|---|---|
| ENSMUSG00000114493 | *Gm47071* | 1.55E-07 | 1.93539 |
| ENSMUSG00000043795 | *Prr33* | 1.22E-02 | 1.93241 |
| ENSMUSG00000024972 | *Lgals12* | 1.56E-04 | 1.92981 |
| ENSMUSG00000037157 | *Il22ra1* | 2.63E-02 | 1.92792 |
| ENSMUSG00000085184 | *4933439K11Rik* | 1.66E-04 | 1.92699 |
| ENSMUSG00000031936 | *Hephl1* | 4.23E-02 | 1.92403 |
| ENSMUSG00000024673 | *Ms4a1* | 1.09E-08 | 1.92013 |
| ENSMUSG00000111556 | *Gm19299* | 2.18E-02 | 1.91814 |
| ENSMUSG00000030178 | *Klra13-ps* | 1.23E-02 | 1.91485 |
| ENSMUSG00000000673 | *Haao* | 2.01E-04 | 1.91426 |
| ENSMUSG00000019102 | *Aldh3a1* | 2.82E-09 | 1.90858 |
| ENSMUSG00000104213 | *Ighd* | 1.37E-08 | 1.90673 |
| ENSMUSG00000034282 | *Evpl* | 4.31E-11 | 1.90263 |
| ENSMUSG00000021223 | *Papln* | 5.43E-04 | 1.89985 |
| ENSMUSG00000041782 | *Lad1* | 1.90E-10 | 1.89447 |
| ENSMUSG00000067889 | *Sptbn2* | 3.18E-09 | 1.89422 |
| ENSMUSG00000056328 | *Myh1* | 5.41E-31 | 1.89409 |
| ENSMUSG00000048142 | *Nat8l* | 2.19E-04 | 1.88665 |
| ENSMUSG00000116290 | *Gm49518* | 4.80E-03 | 1.88407 |
| ENSMUSG00000027359 | *Slc27a2* | 8.34E-03 | 1.88103 |
| ENSMUSG00000061126 | *Cyp4f39* | 1.85E-03 | 1.87931 |
| ENSMUSG00000019853 | *Hebp2* | 2.38E-02 | 1.8788 |
| ENSMUSG00000024347 | *Psd2* | 6.76E-03 | 1.8787 |
| ENSMUSG00000049123 | *Catsperg2* | 1.49E-02 | 1.87815 |
| ENSMUSG00000017300 | *Tnnc2* | 5.40E-15 | 1.87774 |
| ENSMUSG00000026950 | *Neb* | 1.41E-16 | 1.87734 |
| ENSMUSG00000032033 | *Barx2* | 3.20E-08 | 1.87675 |
| ENSMUSG00000046182 | *Gsg1l* | 2.78E-05 | 1.8741 |
| ENSMUSG00000044737 | *Klk14* | 2.20E-06 | 1.87252 |
| ENSMUSG00000046971 | *Pla2g4f* | 1.78E-09 | 1.87121 |
| ENSMUSG00000044086 | *Lmod3* | 1.84E-04 | 1.86547 |
| ENSMUSG00000084162 | *Gm11251* | 4.13E-02 | 1.86137 |
| ENSMUSG00000033508 | *Asprv1* | 4.71E-06 | 1.85815 |
| ENSMUSG00000034579 | *Pla2g3* | 9.27E-03 | 1.85233 |
| ENSMUSG00000106617 | *Gm36266* | 3.40E-02 | 1.85212 |
| ENSMUSG00000037509 | *Arhgef4* | 9.74E-11 | 1.8494 |
| ENSMUSG00000028435 | *Aqp3* | 6.38E-06 | 1.84168 |
| ENSMUSG00000060180 | *Myh13* | 1.20E-05 | 1.8367 |
| ENSMUSG00000030935 | *Acsm3* | 4.56E-02 | 1.83384 |
| ENSMUSG00000028031 | *Dkk2* | 1.01E-07 | 1.83252 |
| ENSMUSG00000039092 | *Sptlc3* | 6.41E-04 | 1.83181 |
| ENSMUSG00000026407 | *Cacna1s* | 2.79E-13 | 1.82909 |
| ENSMUSG00000000031 | *H19* | 2.56E-20 | 1.82543 |
| ENSMUSG00000073492 | *Gm10521* | 2.31E-03 | 1.81983 |
| ENSMUSG00000087382 | *Ctcflos* | 3.63E-03 | 1.81593 |
| ENSMUSG00000014030 | *Pax5* | 1.30E-07 | 1.81402 |
| ENSMUSG00000030724 | *Cd19* | 4.30E-09 | 1.8132 |
| ENSMUSG00000029361 | *Nos1* | 2.87E-14 | 1.81035 |
| ENSMUSG00000069808 | *Fam57a* | 1.84E-11 | 1.80929 |
| ENSMUSG00000061723 | *Tnnt3* | 3.90E-10 | 1.80825 |
| ENSMUSG00000054966 | *Lmntd1* | 3.80E-23 | 1.80655 |
| ENSMUSG00000003863 | *Ppfia3* | 2.28E-02 | 1.80404 |
| ENSMUSG00000060275 | *Nrg2* | 1.27E-04 | 1.80254 |

| | | | |
|---|---|---|---|
| ENSMUSG00000085235 | *Gm12576* | 4.86E-02 | 1.79966 |
| ENSMUSG00000039809 | *Gabbr2* | 3.16E-02 | 1.79942 |
| ENSMUSG00000102305 | *Gm38192* | 1.07E-02 | 1.79922 |
| ENSMUSG00000030510 | *Cers3* | 3.38E-07 | 1.79054 |
| ENSMUSG00000037922 | *Bank1* | 5.44E-06 | 1.7892 |
| ENSMUSG00000027077 | *Smtnl1* | 6.91E-06 | 1.78907 |
| ENSMUSG00000049349 | *Gm5105* | 2.80E-02 | 1.78388 |
| ENSMUSG00000072591 | *5930412G12Rik* | 3.66E-02 | 1.78195 |
| ENSMUSG00000042306 | *S100a14* | 9.20E-08 | 1.77984 |
| ENSMUSG00000042474 | *Fcmr* | 2.09E-05 | 1.77414 |
| ENSMUSG00000067616 | *Klk11* | 7.10E-03 | 1.77343 |
| ENSMUSG00000040283 | *Btnl9* | 1.58E-02 | 1.77128 |
| ENSMUSG00000001095 | *Slc13a2* | 1.63E-03 | 1.76877 |
| ENSMUSG00000095079 | *Igha* | 6.91E-07 | 1.76823 |
| ENSMUSG00000071540 | *3425401B19Rik* | 2.42E-14 | 1.76757 |
| ENSMUSG00000042514 | *Klhl14* | 1.86E-03 | 1.76512 |
| ENSMUSG00000032502 | *Stac* | 1.48E-03 | 1.75181 |
| ENSMUSG00000004885 | *Crabp2* | 1.22E-02 | 1.74533 |
| ENSMUSG00000022871 | *Fetub* | 2.02E-04 | 1.74369 |
| ENSMUSG00000025754 | *Agbl1* | 7.55E-21 | 1.74292 |
| ENSMUSG00000057802 | *Gm10030* | 2.61E-02 | 1.74044 |
| ENSMUSG00000032564 | *Cpne4* | 7.97E-06 | 1.73707 |
| ENSMUSG00000013523 | *Bcas1* | 2.19E-02 | 1.73402 |
| ENSMUSG00000030546 | *Plin1* | 1.25E-07 | 1.73097 |
| ENSMUSG00000085417 | *Gm13919* | 1.02E-02 | 1.72944 |
| ENSMUSG00000079853 | *Klra1* | 3.86E-04 | 1.72252 |
| ENSMUSG00000105646 | *Gm30211* | 5.74E-11 | 1.72189 |
| ENSMUSG00000033182 | *Kbtbd12* | 8.41E-08 | 1.72084 |
| ENSMUSG00000038751 | *Ptk6* | 2.49E-02 | 1.71998 |
| ENSMUSG00000026989 | *Dapl1* | 3.16E-02 | 1.71791 |
| ENSMUSG00000112041 | *9530020I12Rik* | 4.23E-02 | 1.71514 |
| ENSMUSG00000019851 | *Perp* | 5.28E-09 | 1.71321 |
| ENSMUSG00000026418 | *Tnni1* | 1.91E-02 | 1.71016 |
| ENSMUSG00000062515 | *Fabp4* | 9.13E-16 | 1.70935 |
| ENSMUSG00000026834 | *Acvr1c* | 1.49E-04 | 1.70449 |
| ENSMUSG00000028396 | *2310002L09Rik* | 2.22E-02 | 1.70394 |
| ENSMUSG00000060962 | *Dmkn* | 1.01E-09 | 1.69805 |
| ENSMUSG00000074862 | *BC025920* | 2.99E-02 | 1.69659 |
| ENSMUSG00000091712 | *Sec14l5* | 4.66E-04 | 1.69507 |
| ENSMUSG00000025479 | *Cyp2e1* | 3.83E-17 | 1.69172 |
| ENSMUSG00000029055 | *Plch2* | 9.19E-09 | 1.68869 |
| ENSMUSG00000087514 | *Gm16076* | 4.55E-02 | 1.68559 |
| ENSMUSG00000038167 | *Plekhg6* | 3.73E-04 | 1.6825 |
| ENSMUSG00000076617 | *Ighm* | 3.37E-14 | 1.68243 |
| ENSMUSG00000076275 | *Mir675* | 4.98E-04 | 1.68125 |
| ENSMUSG00000021768 | *Dusp13* | 2.66E-02 | 1.68064 |
| ENSMUSG00000027868 | *Tbx15* | 5.26E-11 | 1.67891 |
| ENSMUSG00000014543 | *Klra17* | 3.90E-08 | 1.67493 |
| ENSMUSG00000086564 | *Cd101* | 3.09E-15 | 1.6748 |
| ENSMUSG00000087090 | *Nctc1* | 3.84E-12 | 1.67425 |
| ENSMUSG00000022053 | *Ebf2* | 5.76E-13 | 1.67262 |
| ENSMUSG00000097312 | *Gm26870* | 4.70E-04 | 1.67024 |
| ENSMUSG00000044788 | *Fads6* | 6.63E-03 | 1.66957 |

| | | | |
|---|---|---|---|
| ENSMUSG00000067081 | *Asb18* | 4.14E-04 | 1.66683 |
| ENSMUSG00000067242 | *Lgi1* | 1.29E-02 | 1.66527 |
| ENSMUSG00000025330 | *Padi4* | 2.91E-05 | 1.65776 |
| ENSMUSG00000079015 | *Serpina1c* | 1.52E-03 | 1.65711 |
| ENSMUSG00000040536 | *Necab1* | 2.89E-02 | 1.65665 |
| ENSMUSG00000029603 | *Dtx1* | 2.06E-03 | 1.65369 |
| ENSMUSG00000032311 | *Nrg4* | 1.05E-09 | 1.65365 |
| ENSMUSG00000038457 | *Tmem255b* | 1.95E-03 | 1.65304 |
| ENSMUSG00000098488 | *Pla2g4b* | 8.22E-03 | 1.65198 |
| ENSMUSG00000066366 | *Serpina1a* | 4.02E-03 | 1.64939 |
| ENSMUSG00000029120 | *Ppp2r2c* | 3.19E-03 | 1.64868 |
| ENSMUSG00000027022 | *Xirp2* | 1.28E-22 | 1.64858 |
| ENSMUSG00000037139 | *Myom3* | 8.53E-11 | 1.64798 |
| ENSMUSG00000033544 | *Angptl1* | 1.83E-02 | 1.64513 |
| ENSMUSG00000024136 | *Dnase1l2* | 3.42E-03 | 1.64394 |
| ENSMUSG00000030592 | *Ryr1* | 1.75E-07 | 1.64026 |
| ENSMUSG00000103824 | *Gm38177* | 1.79E-02 | 1.6397 |
| ENSMUSG00000006411 | *Nectin4* | 3.58E-05 | 1.63718 |
| ENSMUSG00000096974 | *Gm26881* | 4.82E-02 | 1.63547 |
| ENSMUSG00000005547 | *Cyp2a5* | 4.57E-06 | 1.63304 |
| ENSMUSG00000113543 | *Gm36264* | 2.07E-02 | 1.63225 |
| ENSMUSG00000026494 | *Kif26b* | 2.39E-14 | 1.63132 |
| ENSMUSG00000070802 | *Pnmal2* | 1.50E-02 | 1.62883 |
| ENSMUSG00000057003 | *Myh4* | 1.66E-10 | 1.62751 |
| ENSMUSG00000028017 | *Egf* | 1.08E-08 | 1.62681 |
| ENSMUSG00000082674 | *Gm11914* | 2.98E-02 | 1.62365 |
| ENSMUSG00000059994 | *Fcrl1* | 1.55E-04 | 1.62354 |
| ENSMUSG00000020169 | *Best3* | 2.24E-05 | 1.62346 |
| ENSMUSG00000046352 | *Gjb2* | 1.75E-05 | 1.62257 |
| ENSMUSG00000058498 | *Rnf207* | 4.30E-03 | 1.6175 |
| ENSMUSG00000033533 | *Acsm1* | 5.95E-04 | 1.61748 |
| ENSMUSG00000017697 | *Ada* | 1.08E-10 | 1.61747 |
| ENSMUSG00000005373 | *Mlxipl* | 2.73E-09 | 1.61612 |
| ENSMUSG00000056755 | *Grm7* | 2.64E-05 | 1.6147 |
| ENSMUSG00000082179 | *Gm11407* | 4.66E-02 | 1.61227 |
| ENSMUSG00000001027 | *Scn4a* | 1.74E-07 | 1.60993 |
| ENSMUSG00000050600 | *Zfp831* | 3.33E-11 | 1.60941 |
| ENSMUSG00000056155 | *Nanos3* | 2.00E-02 | 1.60734 |
| ENSMUSG00000025105 | *Bnc1* | 3.90E-02 | 1.60648 |
| ENSMUSG00000075555 | *Gm10863* | 1.79E-03 | 1.6055 |
| ENSMUSG00000054889 | *Dsp* | 3.74E-45 | 1.60391 |
| ENSMUSG00000074604 | *Mgst2* | 1.19E-07 | 1.60185 |
| ENSMUSG00000036218 | *Pdzrn4* | 1.90E-11 | 1.60165 |
| ENSMUSG00000058975 | *Kcnc1* | 3.09E-03 | 1.60068 |
| ENSMUSG00000000244 | *Tspan32* | 1.03E-08 | 1.5999 |
| ENSMUSG00000031937 | *Vstm5* | 6.27E-03 | 1.5993 |
| ENSMUSG00000034648 | *Lrrn1* | 1.12E-02 | 1.59815 |
| ENSMUSG00000030495 | *Slc7a10* | 2.96E-07 | 1.59742 |
| ENSMUSG00000017639 | *Rab11fip4* | 7.16E-06 | 1.59393 |
| ENSMUSG00000029602 | *Rasal1* | 3.67E-02 | 1.59345 |
| ENSMUSG00000026676 | *Ccdc3* | 1.21E-09 | 1.59159 |
| ENSMUSG00000034308 | *Sdr42e1* | 2.38E-03 | 1.5912 |
| ENSMUSG00000059900 | *Tmem40* | 2.36E-02 | 1.59117 |

| | | | |
|---|---|---|---|
| ENSMUSG00000089941 | *Gm16168* | 3.31E-03 | 1.59041 |
| ENSMUSG00000020892 | *Aloxe3* | 1.31E-02 | 1.58477 |
| ENSMUSG00000115095 | *Gm49311* | 1.45E-02 | 1.58455 |
| ENSMUSG00000031097 | *Tnni2* | 2.64E-07 | 1.58348 |
| ENSMUSG00000031245 | *Hmgn5* | 9.24E-05 | 1.58329 |
| ENSMUSG00000067653 | *Ankrd23* | 7.42E-10 | 1.58165 |
| ENSMUSG00000039264 | *Gimap3* | 5.32E-08 | 1.58068 |
| ENSMUSG00000030730 | *Atp2a1* | 4.52E-14 | 1.57945 |
| ENSMUSG00000083834 | *Gm12577* | 3.00E-02 | 1.57726 |
| ENSMUSG00000076609 | *Igkc* | 1.26E-05 | 1.5742 |
| ENSMUSG00000021579 | *Lrrc14b* | 3.06E-02 | 1.57311 |
| ENSMUSG00000030598 | *Fbxo17* | 1.83E-04 | 1.57292 |
| ENSMUSG00000008999 | *Bmp7* | 9.10E-03 | 1.5718 |
| ENSMUSG00000036214 | *Znrd1as* | 3.27E-02 | 1.57078 |
| ENSMUSG00000030468 | *Siglecg* | 7.97E-06 | 1.56997 |
| ENSMUSG00000085971 | *Gm15411* | 3.59E-04 | 1.56795 |
| ENSMUSG00000024617 | *Camk2a* | 1.83E-04 | 1.56649 |
| ENSMUSG00000021340 | *Gpld1* | 1.71E-06 | 1.56545 |
| ENSMUSG00000031933 | *Izumo1r* | 4.04E-02 | 1.56461 |
| ENSMUSG00000090799 | *Klhl33* | 9.45E-05 | 1.56433 |
| ENSMUSG00000036854 | *Hspb6* | 5.71E-03 | 1.56314 |
| ENSMUSG00000047419 | *Cmya5* | 1.22E-17 | 1.56238 |
| ENSMUSG00000114014 | *Gm48350* | 2.45E-03 | 1.56063 |
| ENSMUSG00000063522 | *2010109I03Rik* | 1.10E-02 | 1.56032 |
| ENSMUSG00000037661 | *Gpr160* | 1.12E-03 | 1.56002 |
| ENSMUSG00000055775 | *Myh8* | 2.99E-07 | 1.55915 |
| ENSMUSG00000097899 | *Gm16894* | 8.00E-03 | 1.55851 |
| ENSMUSG00000039521 | *Foxp3* | 9.57E-03 | 1.55602 |
| ENSMUSG00000070385 | *Ampd1* | 4.23E-05 | 1.55169 |
| ENSMUSG00000050106 | *Tmc8* | 7.90E-05 | 1.5508 |
| ENSMUSG00000035686 | *Thrsp* | 2.97E-07 | 1.54774 |
| ENSMUSG00000041460 | *Cacna2d4* | 4.52E-02 | 1.5477 |
| ENSMUSG00000026824 | *Kcnj3* | 9.29E-10 | 1.54542 |
| ENSMUSG00000020758 | *Itgb4* | 4.89E-09 | 1.54438 |
| ENSMUSG00000032503 | *Arpp21* | 4.76E-03 | 1.54437 |
| ENSMUSG00000043110 | *Lrrn4* | 7.92E-03 | 1.5421 |
| ENSMUSG00000014453 | *Blk* | 1.27E-06 | 1.54192 |
| ENSMUSG00000027419 | *Pcsk2* | 5.87E-03 | 1.54191 |
| ENSMUSG00000034656 | *Cacna1a* | 7.55E-12 | 1.54042 |
| ENSMUSG00000015702 | *Anxa9* | 7.23E-03 | 1.5397 |
| ENSMUSG00000062713 | *Sim2* | 2.58E-05 | 1.53934 |
| ENSMUSG00000068697 | *Myoz1* | 5.79E-04 | 1.5362 |
| ENSMUSG00000038602 | *Slc35f1* | 8.70E-10 | 1.53615 |
| ENSMUSG00000079110 | *Capn3* | 9.20E-06 | 1.5356 |
| ENSMUSG00000036856 | *Wnt4* | 2.59E-06 | 1.53379 |
| ENSMUSG00000000223 | *Drp2* | 1.32E-03 | 1.53365 |
| ENSMUSG00000031710 | *Ucp1* | 7.53E-03 | 1.53359 |
| ENSMUSG00000056656 | *Apol8* | 4.49E-02 | 1.53347 |
| ENSMUSG00000029862 | *Clcn1* | 2.78E-04 | 1.5334 |
| ENSMUSG00000024526 | *Cidea* | 2.67E-02 | 1.53274 |
| ENSMUSG00000042251 | *Pm20d1* | 1.64E-02 | 1.5326 |
| ENSMUSG00000079055 | *Slc8a3* | 2.56E-06 | 1.53124 |
| ENSMUSG00000031377 | *Bmx* | 4.19E-08 | 1.5308 |

| | | | |
|---|---|---|---|
| ENSMUSG00000034570 | *Inpp5j* | 1.71E-02 | 1.52977 |
| ENSMUSG00000048368 | *Omd* | 4.45E-02 | 1.52965 |
| ENSMUSG00000057719 | *Sh3rf2* | 1.60E-09 | 1.52797 |
| ENSMUSG00000035189 | *Ano4* | 1.29E-06 | 1.52212 |
| ENSMUSG00000043932 | *Klri2* | 1.09E-02 | 1.51956 |
| ENSMUSG00000024112 | *Cacna1h* | 1.02E-02 | 1.51951 |
| ENSMUSG00000072812 | *Ahnak2* | 2.64E-06 | 1.51917 |
| ENSMUSG00000024669 | *Cd5* | 8.63E-04 | 1.51847 |
| ENSMUSG00000037989 | *Wnk2* | 1.82E-10 | 1.51835 |
| ENSMUSG00000074497 | *A430078G23Rik* | 3.97E-14 | 1.51676 |
| ENSMUSG00000054708 | *Ankrd24* | 3.01E-04 | 1.51618 |
| ENSMUSG00000032243 | *Itga11* | 6.65E-04 | 1.51519 |
| ENSMUSG00000087150 | *BC064078* | 4.96E-03 | 1.51421 |
| ENSMUSG00000044461 | *Shisa2* | 4.62E-02 | 1.50751 |
| ENSMUSG00000114937 | *Gm30054* | 6.11E-05 | 1.50651 |
| ENSMUSG00000038605 | *Samd10* | 2.26E-03 | 1.50622 |
| ENSMUSG00000040350 | *Trim7* | 2.15E-04 | 1.50563 |
| ENSMUSG00000100551 | *Gm19503* | 3.16E-03 | 1.5047 |
| ENSMUSG00000015314 | *Slamf6* | 1.76E-09 | 1.50457 |
| ENSMUSG00000032262 | *Elovl4* | 5.29E-03 | 1.50436 |
| ENSMUSG00000090166 | *Ear10* | 2.48E-02 | 1.50393 |
| ENSMUSG00000058207 | *Serpina3k* | 2.37E-03 | 1.50282 |
| ENSMUSG00000040694 | *Apobec2* | 2.65E-05 | 1.50189 |
| ENSMUSG00000113447 | *Gm48538* | 4.54E-02 | 1.50191 |
| ENSMUSG00000026646 | *Suv39h2* | 4.57E-02 | 1.5074 |
| ENSMUSG00000030744 | *Rps3* | 1.27E-09 | 1.50807 |
| ENSMUSG00000085025 | *Gm13715* | 3.50E-02 | 1.50829 |
| ENSMUSG00000059835 | *Rpl13-ps3* | 2.53E-03 | 1.51027 |
| ENSMUSG00000032496 | *Ltf* | 3.04E-16 | 1.51096 |
| ENSMUSG00000084104 | *Gm13578* | 2.62E-05 | 1.51286 |
| ENSMUSG00000091845 | *Rpl36-ps12* | 7.00E-03 | 1.51322 |
| ENSMUSG00000100969 | *1700030N03Rik* | 1.54E-02 | 1.51588 |
| ENSMUSG00000082062 | *Ftl2-ps* | 3.39E-09 | 1.51711 |
| ENSMUSG00000076036 | *Gm22133* | 2.46E-11 | 1.51828 |
| ENSMUSG00000083337 | *Gm11539* | 1.80E-03 | 1.52589 |
| ENSMUSG00000040809 | *Chil3* | 1.62E-04 | 1.52703 |
| ENSMUSG00000024774 | *Ankrd22* | 3.56E-08 | 1.53049 |
| ENSMUSG00000052565 | *Hist1h1d* | 7.00E-03 | 1.53071 |
| ENSMUSG00000024026 | *Glo1* | 8.15E-12 | 1.53202 |
| ENSMUSG00000057863 | *Rpl36* | 4.20E-04 | 1.53263 |
| ENSMUSG00000080811 | *Gm14513* | 1.25E-03 | 1.5339 |
| ENSMUSG00000109658 | *Gm45660* | 4.65E-05 | 1.53887 |
| ENSMUSG00000097366 | *Gm8177* | 1.12E-02 | 1.54012 |
| ENSMUSG00000094955 | *Gm3699* | 1.48E-02 | 1.5463 |
| ENSMUSG00000094125 | *Gm13698* | 6.40E-03 | 1.55008 |
| ENSMUSG00000094336 | *Gm13693* | 6.40E-03 | 1.55008 |
| ENSMUSG00000096484 | *Gm13696* | 6.40E-03 | 1.55008 |
| ENSMUSG00000096337 | *Gm13694* | 6.40E-03 | 1.55008 |
| ENSMUSG00000095824 | *Gm13697* | 6.40E-03 | 1.55008 |
| ENSMUSG00000079247 | *Gm13691* | 6.40E-03 | 1.55008 |
| ENSMUSG00000096729 | *Gm13695* | 5.93E-03 | 1.55266 |
| ENSMUSG00000112908 | *Gm7392* | 3.88E-04 | 1.5528 |
| ENSMUSG00000022097 | *Sftpc* | 6.96E-07 | 1.55295 |

| | | | |
|---|---|---|---|
| ENSMUSG00000046031 | *Calhm6* | 1.49E-02 | 1.5541 |
| ENSMUSG00000082978 | *Rpsa-ps11* | 2.04E-02 | 1.55812 |
| ENSMUSG00000085264 | *Gm15581* | 3.86E-02 | 1.56146 |
| ENSMUSG00000028179 | *Cth* | 1.60E-04 | 1.56198 |
| ENSMUSG00000080950 | *Gm7278* | 2.72E-02 | 1.56602 |
| ENSMUSG00000097679 | *Rps19-ps5* | 3.50E-02 | 1.56618 |
| ENSMUSG00000114304 | *Gm48099* | 2.47E-05 | 1.56873 |
| ENSMUSG00000065968 | *Ifitm7* | 3.54E-02 | 1.57151 |
| ENSMUSG00000077391 | *Gm24336* | 3.92E-02 | 1.57527 |
| ENSMUSG00000073940 | *Hbb-bt* | 7.22E-03 | 1.57539 |
| ENSMUSG00000028896 | *Rcc1* | 7.42E-10 | 1.57968 |
| ENSMUSG00000085401 | *Slc39a1-ps* | 5.82E-09 | 1.58067 |
| ENSMUSG00000049932 | *H2afx* | 6.34E-03 | 1.58133 |
| ENSMUSG00000087968 | *Gm25395* | 2.95E-03 | 1.58263 |
| ENSMUSG00000032484 | *Ngp* | 5.93E-21 | 1.58682 |
| ENSMUSG00000083898 | *Gm7340* | 3.68E-05 | 1.58939 |
| ENSMUSG00000097971 | *Gm26917* | 8.77E-04 | 1.59222 |
| ENSMUSG00000050796 | *B3galt6* | 1.23E-02 | 1.59236 |
| ENSMUSG00000048388 | *Fam171b* | 2.37E-10 | 1.59429 |
| ENSMUSG00000024205 | *Rpl36-ps2* | 4.35E-04 | 1.59832 |
| ENSMUSG00000007892 | *Rplp1* | 5.42E-11 | 1.61046 |
| ENSMUSG00000052305 | *Hbb-bs* | 3.42E-03 | 1.61115 |
| ENSMUSG00000065037 | *Rn7sk* | 4.20E-03 | 1.61406 |
| ENSMUSG00000079505 | *Gm11131* | 4.60E-03 | 1.61438 |
| ENSMUSG00000087700 | *Gm15283* | 1.12E-02 | 1.61608 |
| ENSMUSG00000096972 | *Gm26883* | 1.24E-02 | 1.62189 |
| ENSMUSG00000063779 | *Chil4* | 1.09E-03 | 1.62825 |
| ENSMUSG00000062461 | *Gm5453* | 4.23E-05 | 1.62974 |
| ENSMUSG00000083836 | *Ldha-ps2* | 2.54E-03 | 1.63004 |
| ENSMUSG00000084708 | *Gm22988* | 4.90E-03 | 1.63084 |
| ENSMUSG00000106373 | *Gm6522* | 8.32E-03 | 1.64441 |
| ENSMUSG00000075391 | *Gm13443* | 5.63E-08 | 1.6457 |
| ENSMUSG00000077192 | *Snora17* | 1.41E-02 | 1.64917 |
| ENSMUSG00000021071 | *Trim9* | 2.44E-04 | 1.65066 |
| ENSMUSG00000089281 | *Scarna6* | 7.09E-03 | 1.6534 |
| ENSMUSG00000025716 | *Myo3a* | 4.50E-03 | 1.65551 |
| ENSMUSG00000030329 | *Pianp* | 7.04E-03 | 1.66575 |
| ENSMUSG00000097168 | *C230088H06Rik* | 6.39E-08 | 1.67154 |
| ENSMUSG00000063586 | *Gm5513* | 4.76E-03 | 1.67361 |
| ENSMUSG00000070473 | *Cldn3* | 4.42E-09 | 1.67916 |
| ENSMUSG00000083737 | *Prdx6-ps2* | 2.04E-02 | 1.6846 |
| ENSMUSG00000088789 | *Scarna13* | 1.82E-03 | 1.68485 |
| ENSMUSG00000035299 | *Mid1* | 3.52E-03 | 1.71244 |
| ENSMUSG00000070524 | *Fcrlb* | 2.80E-02 | 1.72336 |
| ENSMUSG00000064382 | *Gm26447* | 2.21E-04 | 1.73092 |
| ENSMUSG00000032425 | *Zfp949* | 1.91E-18 | 1.74133 |
| ENSMUSG00000013936 | *Myl2* | 4.07E-03 | 1.74341 |
| ENSMUSG00000104560 | *Gm8115* | 3.84E-04 | 1.74573 |
| ENSMUSG00000089762 | *Ier5l* | 2.27E-04 | 1.74658 |
| ENSMUSG00000087353 | *Gm13727* | 7.20E-06 | 1.75524 |
| ENSMUSG00000085241 | *Snhg3* | 4.49E-06 | 1.75809 |
| ENSMUSG00000113960 | *4933412O06Rik* | 4.05E-03 | 1.76721 |
| ENSMUSG00000075268 | *Gm10819* | 4.64E-05 | 1.77825 |

| | | | |
|---|---|---|---|
| ENSMUSG00000106497 | *Gm43195* | 7.66E-03 | 1.77886 |
| ENSMUSG00000044117 | *2900011O08Rik* | 6.16E-03 | 1.78002 |
| ENSMUSG00000086420 | *Gm8865* | 2.40E-04 | 1.78093 |
| ENSMUSG00000027014 | *Cwc22* | 5.52E-07 | 1.781 |
| ENSMUSG00000077254 | *Gm26079* | 1.93E-02 | 1.79018 |
| ENSMUSG00000062464 | *Cyp4f37* | 2.53E-04 | 1.80716 |
| ENSMUSG00000082755 | *Gm8692* | 2.33E-02 | 1.80988 |
| ENSMUSG00000080542 | *Gm22710* | 3.28E-02 | 1.81091 |
| ENSMUSG00000107722 | *2900060B14Rik* | 2.04E-02 | 1.81334 |
| ENSMUSG00000103103 | *4833445I07Rik* | 5.75E-03 | 1.81998 |
| ENSMUSG00000089542 | *Gm25835* | 8.85E-06 | 1.82629 |
| ENSMUSG00000085599 | *Gm13449* | 1.80E-02 | 1.82911 |
| ENSMUSG00000084744 | *Gm25291* | 2.45E-02 | 1.83791 |
| ENSMUSG00000029372 | *Ppbp* | 1.41E-06 | 1.83838 |
| ENSMUSG00000095098 | *Ccdc85b* | 1.56E-03 | 1.84033 |
| ENSMUSG00000088609 | *Gm24187* | 4.85E-07 | 1.84671 |
| ENSMUSG00000001482 | *Def8* | 5.29E-14 | 1.84897 |
| ENSMUSG00000089617 | *Scarna10* | 6.74E-04 | 1.85948 |
| ENSMUSG00000099587 | *Gm28967* | 6.74E-04 | 1.85948 |
| ENSMUSG00000088246 | *Gm25911* | 1.03E-07 | 1.86065 |
| ENSMUSG00000001983 | *Taco1* | 8.81E-07 | 1.86641 |
| ENSMUSG00000065353 | *Snora73b* | 2.44E-04 | 1.8684 |
| ENSMUSG00000086354 | *Gm13938* | 1.03E-15 | 1.8767 |
| ENSMUSG00000091405 | *Hist2h4* | 4.81E-07 | 1.88798 |
| ENSMUSG00000091957 | *Rps2-ps10* | 4.59E-07 | 1.89014 |
| ENSMUSG00000069273 | *Hist1h3e* | 1.12E-03 | 1.9083 |
| ENSMUSG00000104690 | *Gm47304* | 5.92E-05 | 1.91679 |
| ENSMUSG00000077714 | *Snord17* | 1.66E-04 | 1.91875 |
| ENSMUSG00000088185 | *Scarna2* | 7.71E-07 | 1.9197 |
| ENSMUSG00000049097 | *Ankrd34a* | 1.63E-02 | 1.91985 |
| ENSMUSG00000036264 | *Fstl4* | 2.89E-03 | 1.92434 |
| ENSMUSG00000114456 | *Hist1h2bh* | 1.82E-02 | 1.92447 |
| ENSMUSG00000098178 | *Gm42418* | 1.34E-05 | 1.92525 |
| ENSMUSG00000058385 | *Hist1h2bg* | 1.06E-03 | 1.92534 |
| ENSMUSG00000064387 | *Snora73a* | 5.40E-09 | 1.936 |
| ENSMUSG00000094338 | *Hist1h2bl* | 3.08E-03 | 1.94362 |
| ENSMUSG00000088948 | *Gm23262* | 1.49E-04 | 1.94793 |
| ENSMUSG00000064220 | *Hist2h2aa1* | 3.69E-03 | 1.95446 |
| ENSMUSG00000028461 | *Ccdc107* | 1.54E-09 | 1.96853 |
| ENSMUSG00000065226 | *Gm25791* | 3.14E-06 | 1.98442 |
| ENSMUSG00000029635 | *Cdk8* | 2.29E-09 | 1.98467 |
| ENSMUSG00000092746 | *Rn7s6* | 2.46E-05 | 1.98632 |
| ENSMUSG00000065259 | *Snora30* | 3.24E-02 | 1.99136 |
| ENSMUSG00000060639 | *Hist1h4i* | 1.64E-03 | 2.00154 |
| ENSMUSG00000065822 | *Snord15a* | 5.99E-03 | 2.00285 |
| ENSMUSG00000063954 | *Hist2h2aa2* | 1.91E-05 | 2.0103 |
| ENSMUSG00000077563 | *Snora68* | 6.54E-05 | 2.01347 |
| ENSMUSG00000074398 | *Gm15441* | 7.12E-03 | 2.02986 |
| ENSMUSG00000069308 | *Hist1h2bp* | 3.32E-02 | 2.03521 |
| ENSMUSG00000106649 | *Gm49349* | 2.06E-06 | 2.04015 |
| ENSMUSG00000116237 | *Gm30339* | 1.38E-02 | 2.04542 |
| ENSMUSG00000089655 | *Gm16004* | 2.91E-02 | 2.05643 |
| ENSMUSG00000076281 | *Gm24270* | 2.24E-07 | 2.06413 |

| | | | |
|---|---|---|---|
| ENSMUSG00000104740 | *Mir6516* | 3.24E-03 | 2.06889 |
| ENSMUSG00000097263 | *Gm26804* | 2.12E-03 | 2.10305 |
| ENSMUSG00000089255 | *Snora78* | 2.81E-02 | 2.10781 |
| ENSMUSG00000065911 | *Gm24447* | 2.25E-05 | 2.11307 |
| ENSMUSG00000062727 | *Hist1h2bk* | 3.23E-03 | 2.11378 |
| ENSMUSG00000093378 | *Gm20663* | 7.20E-04 | 2.11505 |
| ENSMUSG00000085348 | *Myhas* | 2.48E-04 | 2.12433 |
| ENSMUSG00000064841 | *Gm26205* | 8.37E-03 | 2.1269 |
| ENSMUSG00000081684 | *Rps2-ps13* | 5.49E-10 | 2.13212 |
| ENSMUSG00000101972 | *Hist1h3i* | 3.84E-02 | 2.13658 |
| ENSMUSG00000064231 | *Gm5321* | 1.89E-02 | 2.13892 |
| ENSMUSG00000087819 | *Gm25117* | 2.03E-06 | 2.14179 |
| ENSMUSG00000065824 | *Gm26315* | 1.47E-05 | 2.1456 |
| ENSMUSG00000106279 | *Gm47287* | 2.67E-06 | 2.1466 |
| ENSMUSG00000105398 | *Gm47305* | 2.67E-06 | 2.14661 |
| ENSMUSG00000097300 | *Gm26835* | 3.09E-02 | 2.15296 |
| ENSMUSG00000104863 | *Gm49347* | 5.00E-04 | 2.15358 |
| ENSMUSG00000065087 | *Snord22* | 1.78E-14 | 2.16707 |
| ENSMUSG00000100210 | *Hist1h3f* | 5.03E-04 | 2.17714 |
| ENSMUSG00000069265 | *Hist1h3a* | 3.80E-02 | 2.18419 |
| ENSMUSG00000064380 | *Gm26448* | 7.15E-09 | 2.18787 |
| ENSMUSG00000064853 | *Gm23442* | 3.71E-06 | 2.1894 |
| ENSMUSG00000065304 | *Gm23245* | 2.59E-03 | 2.20662 |
| ENSMUSG00000110588 | *Gm45774* | 1.44E-02 | 2.22306 |
| ENSMUSG00000064655 | *Gm25788* | 1.17E-02 | 2.22647 |
| ENSMUSG00000064994 | *Gm22422* | 2.44E-03 | 2.23564 |
| ENSMUSG00000099250 | *Rn7s2* | 9.79E-07 | 2.24149 |
| ENSMUSG00000099021 | *Rn7s1* | 9.79E-07 | 2.24153 |
| ENSMUSG00000105460 | *Gm42671* | 4.96E-02 | 2.24303 |
| ENSMUSG00000065118 | *Gm23297* | 4.50E-03 | 2.24685 |
| ENSMUSG00000065254 | *Gm23973* | 2.55E-02 | 2.27365 |
| ENSMUSG00000064999 | *Gm26035* | 2.23E-06 | 2.27841 |
| ENSMUSG00000065686 | *Snora5c* | 1.32E-02 | 2.28658 |
| ENSMUSG00000117074 | *AC121299.1* | 6.16E-03 | 2.29666 |
| ENSMUSG00000086220 | *Gm16599* | 1.82E-03 | 2.30969 |
| ENSMUSG00000096010 | *Hist4h4* | 2.23E-06 | 2.31336 |
| ENSMUSG00000076258 | *Gm23935* | 1.76E-11 | 2.31549 |
| ENSMUSG00000103626 | *Gm37358* | 6.88E-05 | 2.31983 |
| ENSMUSG00000088529 | *Gm26083* | 7.90E-05 | 2.32066 |
| ENSMUSG00000112246 | *Gm40761* | 4.60E-02 | 2.33454 |
| ENSMUSG00000064694 | *Gm24146* | 5.87E-04 | 2.34501 |
| ENSMUSG00000114969 | *Hba-ps3* | 2.31E-03 | 2.35684 |
| ENSMUSG00000086649 | *Gm15286* | 2.21E-02 | 2.36528 |
| ENSMUSG00000064966 | *Snord15b* | 1.96E-06 | 2.36869 |
| ENSMUSG00000091993 | *B930036N10Rik* | 3.93E-04 | 2.3742 |
| ENSMUSG00000053615 | *Gm9913* | 4.61E-02 | 2.38787 |
| ENSMUSG00000108866 | *Gm35082* | 4.48E-02 | 2.40681 |
| ENSMUSG00000068855 | *Hist2h2ac* | 5.77E-05 | 2.40805 |
| ENSMUSG00000099517 | *Hist1h3g* | 3.11E-02 | 2.42777 |
| ENSMUSG00000080365 | *Gm25776* | 6.97E-05 | 2.42824 |
| ENSMUSG00000102189 | *Gm37194* | 5.60E-09 | 2.44452 |
| ENSMUSG00000061482 | *Hist1h4d* | 2.67E-16 | 2.45181 |
| ENSMUSG00000043340 | *6530409C15Rik* | 1.55E-02 | 2.46462 |

| | | | |
|---|---|---|---|
| ENSMUSG00000099583 | *Hist1h3d* | 9.81E-03 | 2.47409 |
| ENSMUSG00000105448 | *Gm25820* | 1.07E-02 | 2.49056 |
| ENSMUSG00000105288 | *Gm25820* | 1.07E-02 | 2.49056 |
| ENSMUSG00000065145 | *Vaultrc5* | 5.90E-06 | 2.49296 |
| ENSMUSG00000088008 | *Gm25492* | 6.25E-04 | 2.49693 |
| ENSMUSG00000064451 | *Snora23* | 6.01E-03 | 2.49877 |
| ENSMUSG00000095217 | *Hist1h2bn* | 8.20E-04 | 2.50304 |
| ENSMUSG00000065649 | *Snora74a* | 9.27E-07 | 2.50464 |
| ENSMUSG00000092267 | *Gm20417* | 2.33E-11 | 2.5151 |
| ENSMUSG00000069274 | *Hist1h4f* | 6.17E-05 | 2.51954 |
| ENSMUSG00000093650 | *Gm20631* | 3.45E-02 | 2.52141 |
| ENSMUSG00000069310 | *Hist1h3c* | 1.27E-03 | 2.52752 |
| ENSMUSG00000087775 | *Rprl2* | 4.48E-05 | 2.53855 |
| ENSMUSG00000071470 | *Ccnb1ip1* | 1.51E-06 | 2.57238 |
| ENSMUSG00000091383 | *Hist1h2al* | 3.90E-12 | 2.58142 |
| ENSMUSG00000092837 | *Rpph1* | 1.54E-06 | 2.5885 |
| ENSMUSG00000092386 | *Gm20536* | 2.30E-02 | 2.58947 |
| ENSMUSG00000019865 | *Nmbr* | 2.79E-02 | 2.5947 |
| ENSMUSG00000102349 | *Gm37376* | 5.22E-06 | 2.59474 |
| ENSMUSG00000060678 | *Hist1h4c* | 1.34E-12 | 2.602 |
| ENSMUSG00000097417 | *Gm26669* | 3.90E-03 | 2.60481 |
| ENSMUSG00000105145 | *Gm47299* | 4.92E-08 | 2.60571 |
| ENSMUSG00000105788 | *Gm47295* | 4.92E-08 | 2.60593 |
| ENSMUSG00000089536 | *Scarna3a* | 6.53E-03 | 2.6208 |
| ENSMUSG00000101355 | *Hist1h3h* | 1.95E-02 | 2.63462 |
| ENSMUSG00000088675 | *Rprl1* | 4.46E-03 | 2.64112 |
| ENSMUSG00000106222 | *Rprl1* | 4.46E-03 | 2.64112 |
| ENSMUSG00000069266 | *Hist1h4b* | 5.61E-04 | 2.64927 |
| ENSMUSG00000069267 | *Hist1h3b* | 1.36E-03 | 2.66856 |
| ENSMUSG00000060981 | *Hist1h4h* | 3.18E-11 | 2.67747 |
| ENSMUSG00000064288 | *Hist1h4k* | 6.70E-06 | 2.68893 |
| ENSMUSG00000064604 | *Snora44* | 4.05E-05 | 2.70815 |
| ENSMUSG00000093355 | *Snora26* | 1.76E-02 | 2.76757 |
| ENSMUSG00000095590 | *Gm24305* | 5.74E-03 | 2.78226 |
| ENSMUSG00000092819 | *Gm23639* | 4.60E-03 | 2.79569 |
| ENSMUSG00000115420 | *AL732506.1* | 1.91E-12 | 2.79623 |
| ENSMUSG00000088088 | *Rmrp* | 1.91E-12 | 2.79623 |
| ENSMUSG00000086953 | *Aknaos* | 4.22E-02 | 2.79896 |
| ENSMUSG00000010342 | *Tex14* | 2.50E-11 | 2.80189 |
| ENSMUSG00000067455 | *Hist1h4j* | 1.31E-04 | 2.80961 |
| ENSMUSG00000084783 | *Gm15419* | 5.16E-07 | 2.81003 |
| ENSMUSG00000111971 | *Gm48678* | 1.46E-04 | 2.81027 |
| ENSMUSG00000064901 | *Snora21* | 1.28E-03 | 2.87377 |
| ENSMUSG00000087943 | *Gm24245* | 1.66E-12 | 2.89193 |
| ENSMUSG00000087963 | *Gm25394* | 8.18E-03 | 2.89624 |
| ENSMUSG00000096243 | *Gm24265* | 1.61E-07 | 2.90069 |
| ENSMUSG00000065905 | *Gm26110* | 9.26E-04 | 2.91055 |
| ENSMUSG00000105399 | *Mir1843b* | 5.37E-03 | 2.9215 |
| ENSMUSG00000097131 | *D230017M19Rik* | 5.32E-12 | 2.94112 |
| ENSMUSG00000092674 | *Gm24105* | 7.90E-11 | 2.95616 |
| ENSMUSG00000077323 | *Rnu11* | 1.61E-07 | 2.98011 |
| ENSMUSG00000096992 | *Gm26788* | 9.33E-03 | 2.98078 |
| ENSMUSG00000117227 | *AC105304.3* | 2.95E-02 | 2.9812 |

| | | | |
|---|---|---|---|
| ENSMUSG00000065778 | *Gm22154* | 2.44E-03 | 2.9846 |
| ENSMUSG00000105790 | *Gm24105* | 8.13E-11 | 2.98894 |
| ENSMUSG00000088025 | *Rprl3* | 6.04E-07 | 2.99575 |
| ENSMUSG00000095580 | *Rnu1b1* | 3.38E-07 | 3.01229 |
| ENSMUSG00000093834 | *Rnu1b2* | 3.38E-07 | 3.01229 |
| ENSMUSG00000064923 | *Gm22042* | 1.44E-07 | 3.01372 |
| ENSMUSG00000095701 | *Gm24830* | 9.14E-08 | 3.03287 |
| ENSMUSG00000096838 | *Gm26232* | 9.14E-08 | 3.03287 |
| ENSMUSG00000065773 | *Rnu1b6* | 9.14E-08 | 3.03287 |
| ENSMUSG00000010362 | *Rdm1* | 9.37E-11 | 3.04878 |
| ENSMUSG00000095260 | *Gm25890* | 1.10E-07 | 3.05099 |
| ENSMUSG00000094812 | *Gm22614* | 1.10E-07 | 3.05099 |
| ENSMUSG00000069300 | *Hist1h2bj* | 7.93E-04 | 3.06257 |
| ENSMUSG00000069305 | *Hist1h4n* | 3.36E-07 | 3.0744 |
| ENSMUSG00000094377 | *Gm24407* | 6.03E-10 | 3.08024 |
| ENSMUSG00000069306 | *Hist1h4m* | 2.82E-06 | 3.08188 |
| ENSMUSG00000060988 | *Galnt13* | 9.80E-33 | 3.13528 |
| ENSMUSG00000065701 | *Rny1* | 1.52E-08 | 3.17627 |
| ENSMUSG00000070167 | *Snora57* | 1.98E-15 | 3.18889 |
| ENSMUSG00000094655 | *Gm25360* | 5.67E-15 | 3.20136 |
| ENSMUSG00000108563 | *Gm44686* | 1.39E-02 | 3.23591 |
| ENSMUSG00000080465 | *Gm22486* | 2.04E-09 | 3.27578 |
| ENSMUSG00000091165 | *Gm17036* | 1.50E-02 | 3.27618 |
| ENSMUSG00000060093 | *Hist1h4a* | 1.46E-12 | 3.32521 |
| ENSMUSG00000092805 | *Gm26461* | 1.31E-10 | 3.34331 |
| ENSMUSG00000096214 | *Gm22634* | 5.90E-18 | 3.48166 |
| ENSMUSG00000096206 | *Gm22317* | 5.90E-18 | 3.48166 |
| ENSMUSG00000027233 | *Patl2* | 1.67E-05 | 3.4999 |
| ENSMUSG00000065845 | *Gm25189* | 7.04E-03 | 3.50714 |
| ENSMUSG00000096205 | *Gm22068* | 8.03E-11 | 3.52569 |
| ENSMUSG00000088252 | *Snord13* | 8.10E-08 | 3.5475 |
| ENSMUSG00000065176 | *Rnu12* | 1.76E-21 | 3.61877 |
| ENSMUSG00000097799 | *Gm26899* | 1.05E-03 | 3.63697 |
| ENSMUSG00000092702 | *Gm24514* | 7.15E-12 | 3.70097 |
| ENSMUSG00000064943 | *Gm23240* | 3.63E-03 | 3.72556 |
| ENSMUSG00000085573 | *Gm15418* | 1.61E-07 | 3.75952 |
| ENSMUSG00000110790 | *Gm47079* | 2.36E-03 | 3.78383 |
| ENSMUSG00000065767 | *Gm23849* | 2.98E-11 | 3.82342 |
| ENSMUSG00000065944 | *Rnu2-10* | 1.90E-11 | 3.84825 |
| ENSMUSG00000094826 | *Gm23804* | 1.21E-18 | 3.84945 |
| ENSMUSG00000064702 | *Gm24950* | 2.50E-11 | 3.85414 |
| ENSMUSG00000065251 | *Gm23971* | 2.50E-11 | 3.85414 |
| ENSMUSG00000064856 | *Gm23444* | 3.12E-11 | 3.87597 |
| ENSMUSG00000093956 | *Gm24497* | 6.05E-11 | 3.87768 |
| ENSMUSG00000064945 | *Rny3* | 1.84E-12 | 3.87863 |
| ENSMUSG00000094050 | *Gm23472* | 2.54E-11 | 3.89363 |
| ENSMUSG00000094306 | *Gm24924* | 1.54E-10 | 3.89902 |
| ENSMUSG00000108171 | *Gm43915* | 7.23E-04 | 3.89946 |
| ENSMUSG00000065820 | *Gm26316* | 9.58E-12 | 3.91062 |
| ENSMUSG00000097346 | *Gm26619* | 2.86E-04 | 3.94457 |
| ENSMUSG00000065870 | *Rnu3a* | 1.51E-11 | 3.95478 |
| ENSMUSG00000106147 | *Rnu3a* | 1.51E-11 | 3.95478 |
| ENSMUSG00000095969 | *Rnu1a1* | 1.19E-11 | 3.95595 |

| | | | |
|---|---|---|---|
| ENSMUSG00000096659 | *Gm25679* | 1.03E-19 | 3.9906 |
| ENSMUSG00000093815 | *Gm26444* | 1.03E-19 | 3.9906 |
| ENSMUSG00000115432 | *D130009I18Rik* | 1.17E-07 | 4.00512 |
| ENSMUSG00000105115 | *Rnu3b2* | 3.78E-19 | 4.06451 |
| ENSMUSG00000098925 | *Rnu3b2* | 3.78E-19 | 4.06451 |
| ENSMUSG00000098943 | *Rnu3b1* | 6.81E-19 | 4.07869 |
| ENSMUSG00000105025 | *Rnu3b1* | 6.81E-19 | 4.07869 |
| ENSMUSG00000099291 | *Rnu3b3* | 1.50E-18 | 4.09008 |
| ENSMUSG00000104856 | *Rnu3b3* | 1.50E-18 | 4.09008 |
| ENSMUSG00000104896 | *Rnu3b4* | 3.78E-19 | 4.1079 |
| ENSMUSG00000098641 | *Rnu3b4* | 4.19E-19 | 4.1081 |
| ENSMUSG00000095616 | *Gm26244* | 5.63E-11 | 4.1472 |
| ENSMUSG00000080538 | *Gm25541* | 1.15E-12 | 4.15571 |
| ENSMUSG00000064941 | *Gm23238* | 2.71E-15 | 4.16236 |
| ENSMUSG00000065232 | *Gm22973* | 6.24E-13 | 4.18652 |
| ENSMUSG00000085393 | *Gm11280* | 4.14E-03 | 4.19886 |
| ENSMUSG00000096349 | *Gm22513* | 1.18E-06 | 4.21069 |
| ENSMUSG00000090625 | *Gm20721* | 7.53E-07 | 4.26306 |
| ENSMUSG00000095676 | *Gm25099* | 1.09E-21 | 4.3711 |
| ENSMUSG00000095738 | *Gm25313* | 2.50E-11 | 4.39293 |
| ENSMUSG00000093843 | *Gm25939* | 6.33E-11 | 4.45452 |
| ENSMUSG00000095892 | *Rnu5g* | 3.84E-12 | 4.48461 |
| ENSMUSG00000064682 | *Gm25813* | 1.88E-09 | 4.50868 |
| ENSMUSG00000085297 | *Gm11651* | 8.54E-10 | 4.5091 |
| ENSMUSG00000085465 | *Gm15347* | 3.44E-06 | 4.85263 |
| ENSMUSG00000090208 | *Gm15851* | 5.99E-04 | 4.92229 |
| ENSMUSG00000023467 | *Tulp2* | 2.27E-05 | 5.12205 |
| ENSMUSG00000064899 | *Snord118* | 2.09E-07 | 5.44474 |
| ENSMUSG00000094668 | *Gm24871* | 6.67E-06 | 5.69726 |
| ENSMUSG00000097815 | *Gm26809* | 9.27E-08 | 5.9877 |
| ENSMUSG00000047061 | *Gm9817* | 3.07E-05 | 7.37847 |
| ENSMUSG00000097554 | *Gm26825* | 2.08E-06 | 11.0581 |
| ENSMUSG00000105361 | *AY036118* | 1.29E-26 | 11.3177 |
